# Benchmarking longitudinal local centiles for radiographic knee osteoarthritis prognosis

**DOI:** 10.64898/2026.10.06.757088

**Authors:** Tengfei Li, Tianyou Luo, Boqi Chen, Chao Huang, Zhengyang Shen, Zhenlin Xu, Daniel Nissman, Yvonne M. Golightly, Amanda E. Nelson, Girish Gandikota, Marc Niethammer, Hongtu Zhu

## Abstract

Cartilage thickness on MRI is a structural biomarker of osteoarthritis (OA), yet anatomical variation and aging can mask early disease, including changes occurring 12-36 months before incident OA. We propose longitudinal local centile maps that use normative growth charts as prior information and integrate a knee’s earlier scans to identify cartilage that is thinner or thicker than expected. We built the reference charts from 5,045 scans of 957 persistently radiographically normal knees in the Osteoarthritis Initiative. Under participant-separated nested cross-validation, longitudinal centile profiles achieved higher 12- and 24-month AUCs for predicting incident radiographic OA than regional PCA, radiomics, cross-sectional centile features, and raw-map deep features across most classifiers. These conventional descriptors added little when combined with centile profiles, and cross-sectional centiles added little beyond longitudinal centiles. Deep centile features achieved the best AUCs (0.954 [95% CI 0.931-0.974] at 12 months and 0.969 [0.949-0.985] at 24 months; balanced accuracy 0.917 and 0.931) and outperformed raw-map deep features in all 12 classifiers after correction for multiple comparisons. No representation consistently improved 36-month prediction. Longitudinal centiles transform cartilage thickness maps into reference-based deviation maps: deep longitudinal centile features provide the highest predictive accuracy, while regional longitudinal centile profiles provide interpretable prognosis with the second-highest accuracy.

## 1. Introduction

Osteoarthritis (OA) affected an estimated 595 million people in 2020, with knee OA cases projected to increase by 75% by 2050 (GBD 2021 Osteoarthritis Collaborators, 2023). Identifying knees likely to progress within one to two years is important for trials of structure-modifying treatments (Hunter and Bierma-Zeinstra, 2019; Hochberg et al., 2019; Guehring et al., 2021). Radiographic OA, defined by Kellgren–Lawrence grade (KLG) ≥2, becomes apparent after substantial joint changes have occurred (Kellgren and Lawrence, 1957). MRI measures cartilage directly, making cartilage thickness a structural biomarker for detecting change before radiographic disease (Peterfy et al., 2008; Eckstein et al., 2014).

Absolute thickness is difficult to interpret in an individual knee. Cartilage varies with age, sex, body size and BMI, so the same thickness can be normal in one person and unusually low in another (Faber et al., 2001; Otterness and Eckstein, 2007). OA-related changes are also focal and bidirectional: central weight-bearing cartilage often thins, while other regions can thicken (Buck et al., 2010; Wirth et al., 2009; Eckstein et al., 2010a). Regional averages obscure these local changes. Location-independent measures improve sensitivity to progression (Buck et al., 2009, 2011), and regional reference values account for some anatomical variation (Eckstein et al., 2010b), but neither gives a local expectation conditioned on both a person’s characteristics and the knee’s previous scans.

Normative modeling addresses both problems in neuroimaging. Growth-chart-style centile models place each individual relative to a reference distribution conditional on covariates (Cole and Green, 1992; Rigby and Stasinopoulos, 2005; Marquand et al., 2016, 2019), and lifespan brain charts have shown that centile scores are stable, heritable and sensitive to disease (Bethlehem et al., 2022). Voxel- and vertex-wise deviation maps reveal heterogeneous, individual-specific abnormalities that group averages conceal (Ziegler et al., 2014; Wolfers et al., 2018; Segal et al., 2023), and the number of outlying regions predicts clinical conversion (Verdi et al., 2023). Conditional growth charts (Wei and He, 2006) and recent longitudinal extensions (Rehak Buckova et al., 2025) quantify change relative to an individual’s own history. A longitudinal reference is particularly important for cartilage: focal loss can remain within the population range, while persistently thin cartilage need not indicate recent deterioration.

At the same time, OA prediction research has moved toward high-dimensional image descriptors: radiomic texture features (Hirvasniemi et al., 2021; Li et al., 2022) and deep learning on radiographs or MRI (Tiulpin et al., 2019; Guan et al., 2020; Leung et al., 2020; Tolpadi et al., 2020). These models can be accurate in internal data but are hard to interpret, and the blinded KNOAP2020 challenge showed that incident radiographic OA remains difficult to predict (best AUC 0.64; Hirvasniemi et al., 2023). Deep transfer-learning features from ImageNet-pretrained networks offer an inexpensive, widely used image representation (Kirchler et al., 2022; Zhao et al., 2024), but whether they capture information beyond simpler, biologically grounded representations is unclear (Raghu et al., 2019).

We establish a longitudinal local centile framework that converts knee cartilage thickness into reference-based deviation maps for early OA prognosis (Figure 1). Our contributions are:

1. **Dense normative references for knee cartilage.** At each of 74,724 atlas pixels, we fit a generalized additive mixed model (GAMM) with a nonlinear age × BMI surface, sex-specific age and BMI slopes, race, height and participant random effects in radiographically normal knees. These charts extend regional reference values to local thickness expectations across age and body size.
2. **Longitudinal local centiles from historical MRI.** Earlier scans personalize the thickness expectation at each location, yielding maps of thinner- and thicker-than-expected cartilage. At equal feature dimensions, longitudinal profiles and map deep features outperform their cross-sectional counterparts in most classifiers at 12 and 24 months and retain nearly all combined-centile performance.
3. **Interpretable regional centile prognosis and higher prediction of deep longitudinal centile profiles.** Regional profiles quantify the burden and extent of connected cartilage deficits relative to the reference population, giving anatomically interpretable prognosis. Deep features of the continuous centile maps give the highest accuracy and outperform raw-map deep features in all 12 classifiers at 12 and 24 months after Holm adjustment.
4. **Benchmark prognostic value beyond conventional and cross-sectional features.** Longitudinal centile profiles achieve higher 12- and 24-month AUCs than regional principal component analysis (PCA), radiomics, cross-sectional centile features and raw-map deep features in at least 11 of 12 classifiers per comparison under participant-separated nested cross-validation. Paired tests further show that combined centile profiles add prognostic information to conventional descriptors, with limited reciprocal benefit.

**Figure 1.**
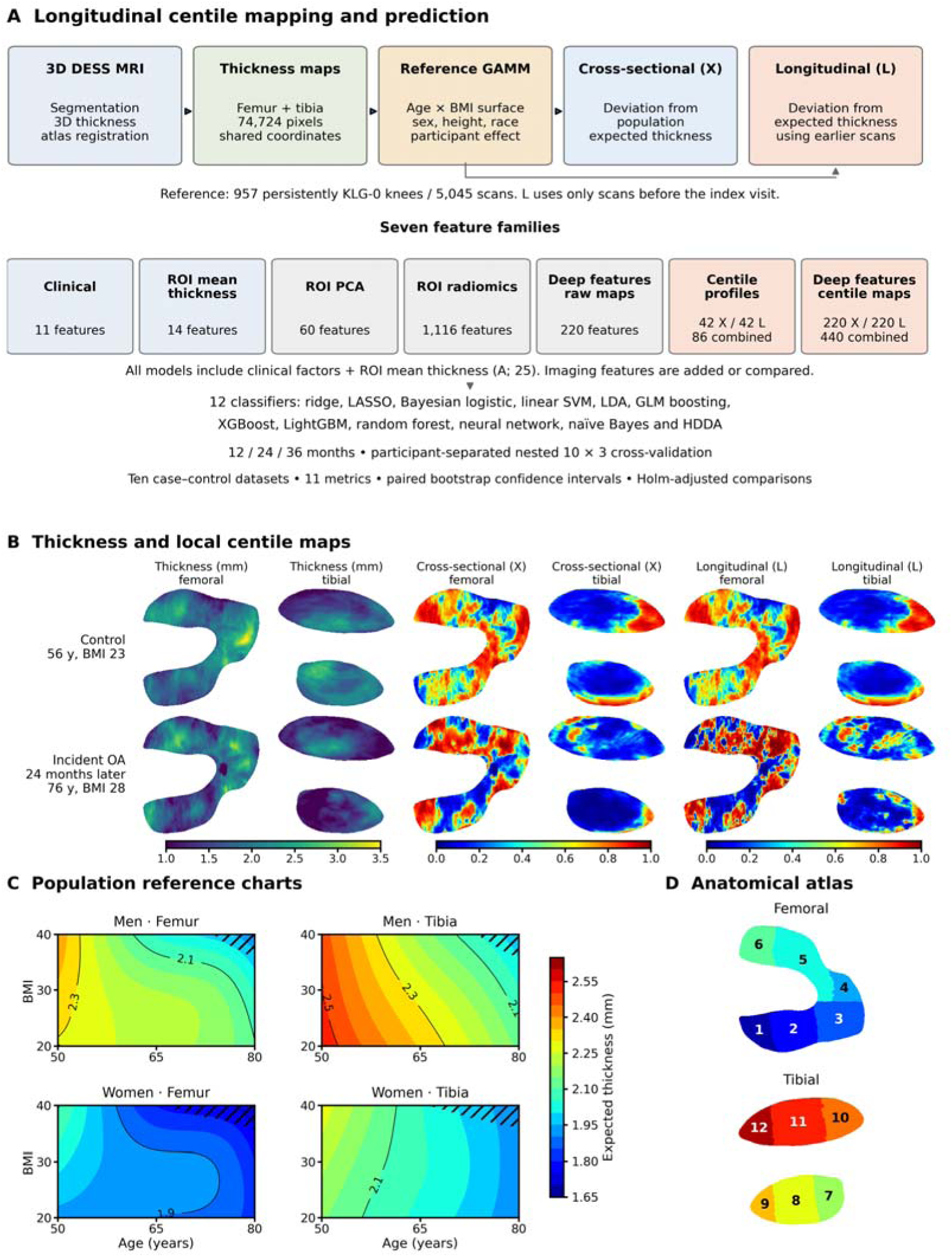
Longitudinal centile mapping and prediction. (A) MRI is converted to atlas-aligned thickness maps and referenced to pixelwise generalized additive mixed models. Cross-sectional centiles (X) use population expectations; longitudinal centiles (L) also use earlier scans of the knee. Seven feature families are compared with the same participants, folds and classifiers. All prediction models include clinical factors and regional mean thickness (A). Combined centile profiles include two pooled maximal-area features. (B) Thickness and centile maps for an illustrative control and a knee developing radiographic OA 24 months later. Centiles range from 0 (thinner than expected) to 1 (thicker). (C) Expected population mean thickness by age and BMI, at fixed height and race. Hatching marks extrapolation beyond observed support; full surfaces and fitted-mean confidence limits are in the supplementary charts. (D) Anatomical ROIs: 1 LPF, 2 LCF, 3 LAF, 4 MAF, 5 MCF, 6 MPF, 7 LPT, 8 LCeT, 9 LAT, 10 MPT, 11 MCT, 12 MAT.

## 2. Related work

### Cartilage thickness and reference values

Quantitative DESS MRI gives reproducible cartilage thickness measurements (Peterfy et al., 2008; Eckstein et al., 2011b). Medial femorotibial thickness loss predicts structural and symptomatic progression (Eckstein et al., 2015a; Wirth et al., 2017), and low baseline medial femoral thickness predicts subsequent thinning (Eckstein et al., 2011a). Both thinning and accelerated thickening can precede incident OA (Iriondo et al., 2021). Regional Z-scores (Eckstein et al., 2010b) and ordered subregional changes (Buck et al., 2009, 2011; Eckstein et al., 2015b) improve interpretation and sensitivity but do not provide a dense, history-adjusted reference map. Our previous atlas method established correspondence between cartilage thickness maps (Huang et al., 2022). The present framework adds local normative references conditioned on population covariates and each knee’s earlier scans.

### Population and longitudinal normative models

GAMLSS, Gaussian-process models and Bayesian regression have established normative modeling in neuroimaging (Rigby and Stasinopoulos, 2005; Dinga et al., 2021; Fraza et al., 2021; Kia et al., 2022; Ge et al., 2024), including models that account for imaging-site differences (Bayer et al., 2022). Conditional growth charts incorporate previous measurements (Wei and He, 2006). Longitudinal z-diff scores use pretrained cross-sectional brain models to quantify change (Rehak Buckova et al., 2025), and recent velocity-centile models predict cognitive decline (Bayer et al., 2026, preprint). Our cartilage model uses a random-intercept update to adjust expected local thickness for the knee’s earlier scans. The prognostic comparison tests the value of this adjustment in both regional profiles and map deep features.

### OA prediction

Clinical models for incident radiographic OA reach AUCs of approximately 0.66–0.79, depending on whether radiographic predictors are included (Kerkhof et al., 2014). Eight-year OAI models reach 0.67 without and 0.72–0.79 with semiquantitative MRI and T2 measures (Joseph et al., 2018, 2022). Radiograph-based deep learning predicts joint-space loss and knee replacement (Guan et al., 2020; Leung et al., 2020; Tolpadi et al., 2020; Rajamohan et al., 2023); MRI deep learning and radiomics have also been evaluated for structural progression and OA outcomes (Schiratti et al., 2021; Hirvasniemi et al., 2021; Li et al., 2022). The blinded KNOAP2020 challenge reported a best AUC of 0.64 for incident symptomatic radiographic OA (Hirvasniemi et al., 2023). Different outcomes, follow-up periods and sampling schemes prevent direct comparison of these AUCs. We therefore assess added value through paired comparisons within the same study samples.

### Interpretability

SHAP explains the contributions of predictors to fitted models (Lundberg and Lee, 2017; Lundberg et al., 2020). Interpretable predictors and additive models can also make the prediction process easier to examine (Rudin, 2019; Lou et al., 2013; Caruana et al., 2015). Our regional centile profiles have a defined anatomical meaning before classifier fitting: each measures the extent of connected, thinner-than-expected cartilage relative to a reference distribution. Deep features of centile maps retain more spatial information but do not have this feature-by-feature interpretation.

## 3. Materials and methods

### 3.1 Data and cohorts

We used retrospective data from the Osteoarthritis Initiative (OAI), a multicenter cohort of 4,796 participants aged 45–79 years (Peterfy et al., 2008). Knee MRI and radiographs were obtained at baseline and months 12, 24, 36, 48, 72 and 96. KLG was read from fixed-flexion posteroanterior radiographs. We analyzed right knees. The normative reference comprised 957 knees that remained KLG 0 throughout observed follow-up, contributing 5,045 MRI scans.

Prediction followed the event-anchored design in Supplementary S1. For each horizon h (12, 24 or 36 months), cases had a first observed KLG ≥2 radiograph preceded by at least one observed radiograph, with all earlier readings ≤1. The index MRI was acquired h months before that event, when KLG was 0 or 1. When the required MRI was unavailable, a scan 12 months before the event was used for 1, 1 and 2 cases at the respective horizons. Index KLG was 1 in 142/196, 78/116 and 48/80 cases. Event visits could occur at different calendar months; the horizon was defined relative to each event.

Controls had an observed KLG-0 radiograph at month h and KLG 0 at every other observed visit; missing readings were allowed. Their median last radiographic reading was at month 48, and their index MRI was the baseline scan. Nonprogressing KLG-1 knees were excluded from controls. Clinical predictors came from the latest visit at or before the index MRI. This design compares future incident OA with persistently radiographically normal knees.

The 12-, 24- and 36-month cohorts contained 196, 116 and 80 cases and 847, 782 and 716 controls. Controls were divided into ten disjoint, BMI-stratified subsets. Each analysis dataset contained all cases and one control subset: 279–282, 194–195 and 150–153 knees at the three horizons. Thus, each case appeared in all ten datasets and each control in one. Models were fitted separately in each dataset; performance was averaged over datasets, and participant-level bootstrapping accounted for repeated appearances of cases.

Prior-scan availability differed between groups. All controls were indexed at baseline, whereas 105/196, 75/116 and 23/80 cases were indexed later at the three horizons. The longitudinal comparison therefore reflects this observation schedule; its implications are discussed in Section 5.4.

Concurrent KLG ≥2 versus ≤1 classification used 2,249 scans from 1,923 participants. These models included 11 clinical predictors (sex, race, BMI, height, age, Alignment, Injury history, Physical activity, Quadriceps strength, Laxity, historical KLG index), 14 regional thickness means and 84 centile profiles. The two maximal-area features were used only for prognosis, so the concurrent baseline and baseline-plus-profile sets contained 19 and 103 predictors (Supplementary Table S6). Cohort sizes and demographic comparisons are given in Supplementary Tables S1–S2.

### 3.2 Cartilage thickness maps

3D DESS images (0.36 × 0.36 × 0.7 mm) were processed with an automated pipeline for segmentation, meshing, three-dimensional thickness computation, registration and projection to a common two-dimensional atlas (Huang et al., 2022), yielding femoral (38,009 pixels) and tibial (36,715 pixels) local thickness maps in correspondence across knees and visits. A musculoskeletal radiologist delineated twelve regions of interest (ROIs): lateral and medial posterior, central and anterior femur (LPF, LCF, LAF, MPF, MCF, MAF) and the corresponding tibial regions (LPT, LCeT, LAT, MPT, MCT, MAT) (Figure 1D).

### 3.3 Normative thickness model

Let *y_i,v_*(*t*,*b*) denote cartilage thickness of knee *i* at pixel *v* when age is *t* and BMI is *b*. In the reference sample we fit, separately at every pixel,

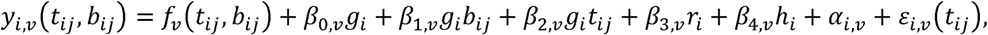

where *f_v_* is a tensor-product smooth surface over age and BMI, *g_i_* is male sex, *r_v_* Black race, *h_i_* height, *α_iv_* ~ 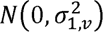 a participant random intercept that accounts for repeated scans, and 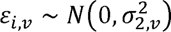. The sex-by-age and sex-by-BMI terms let the chart differ between women and men. Models were estimated by maximum likelihood with the R package mgcv (Wood, 2011, 2017; R 3.6.0 for the full-resolution full-reference charts and mgcv 1.9.3 with R 4.4.0 for the downsampled charts of Supplementary S9). The same model fitted to ROI-mean thickness produced the regional reference charts in Figure 1C; it was compared with a fully linear model (M2) and a model with a one-dimensional age smooth (M3) by maximum likelihood on identical data, using AIC, generalized cross-validation and the marginal likelihood (Supplementary Table S3).

### 3.4 Cross-sectional and longitudinal centile maps

#### Cross-sectional (population; fixed-effect) centile

For a new scan with covariates *x*, the population residual *r_v_* = *y_v_* – *μ*_^v_(*x*) removes the fixed effects. Its local centile is the probability that a matched reference knee has a smaller residual,

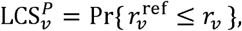

estimated against a wild-bootstrap expansion of a null pool of residuals, in which residuals are multiplied by standard-normal multipliers shared across a participant’s scans (Supplementary Methods S3).

The null pool and number of multipliers depend on compartment and channel: cross-sectional centiles of both compartments use the reference residuals with 100 multipliers; femoral longitudinal centiles use the conditional residuals of the reference training scans, after subtracting the fitted participant random effect, with 100 multipliers (these null residuals differ from the target’s history residual, which subtracts an effect predicted from earlier scans only); and tibial longitudinal centiles use the follow-up (non-baseline) history residuals of KLG 0 scans with 1,000 multipliers, so that extreme centiles are resolved beyond the reference sample size (Supplementary Methods S3).

#### Longitudinal (history; random-effect) centile

When the knee has *N* earlier scans, the expected thickness additionally includes the predicted participant effect, updated incrementally without refitting:

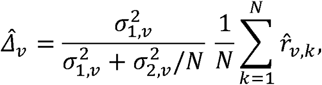

so that the longitudinal residual *r_v_* – ^Δ*_v_* measures deviation from the population trajectory after adjusting for the knee’s persistent offset, shrunk toward the population when history is short or noisy. This weight is the best linear unbiased predictor (empirical-Bayes posterior mean) of the knee’s random effect under the GAMM; its derivation, assumptions and limiting cases are given in Supplementary Methods S3. Its centile 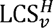 is computed analogously; for a first scan LCS*^H^* = LCS*^P^*. Low centiles indicate thinner-than-expected cartilage and high centiles thicker-than-expected cartilage (Figure 1B).

In Table 1 and figures throughout, feature settings X and L denote cross-sectional and longitudinal centiles in the feature-set names. In feature-set names, P denotes the combined regional profile block. The longitudinal update uses only scans before the index visit. It adjusts a knee-specific offset; the age–BMI trajectory remains the fitted population trajectory. At a first scan, the two local maps are equal, but their regional profiles can differ because subsequent ranking and compression are performed separately. Local centiles are empirical reference ranks; calibration by observation-history length has not been established.

**Table 1.** Prognostic feature sets and their inputs. A contains 11 clinical predictors and 14 regional thickness means (25 total). X and L denote cross-sectional and longitudinal centiles. PX and PL contain 42 profiles each; P combines them with two compartmental maximum-area measures (86). F1 contains 220 raw-map deep features; F2X and F2L contain 220 cross-sectional and longitudinal centile-map deep features each; F2 combines both (440). All centile features in this table use full-resolution full-reference charts. Predictor counts include A. Concurrent classification uses A = 19 and P = 84 predictors (Section 3.1).

| Symbol | Representation | Composition | Predictors |
| --- | --- | --- | --- |
| A | Clinical + regional thickness | 11 clinical + 14 means | 25 |
| B | A + ROI PCA | A + 60 PCs | 85 |
| C | A + ROI radiomics | A + 1,116 radiomics | 1,141 |
| D | A + PCA + radiomics | B + radiomics | 1,201 |
| A+PX | A + cross-sectional profiles | A + PX | 67 |
| A+PL | A + longitudinal profiles | A + PL | 67 |
| A+P | A + combined centile profiles | A + P | 111 |
| E | D + combined centile profiles | D + P | 1,287 |
| G1 | A + raw-map deep features | A + F1 | 245 |
| G2X | A + deep cross-sectional centile features | A + F2X | 245 |
| G2L | A + deep longitudinal centile features | A + F2L | 245 |
| G2 | A + deep combined centile features | A + F2 | 465 |
| G12 | A + all combined raw and centile map deep features | A + F1 + F2 | 685 |
| H1 | G1 + combined centile profiles | G1 + P | 331 |
| H2 | G2 + combined centile profiles | G2 + P | 551 |
| H12 | G12 + combined centile profiles | G12 + P | 771 |

### 3.5 Regional centile profiles

For each ROI, each whole plate and each centile type, we measured the largest eight-connected area whose centile fell below each of five thresholds (*μ_k_* = 0.1exp{–(*k*–1)log(1000)/19}, 0.10 to 0.023), ranked each extent against the extents of reference knees, and combined the five ranks with normalized first-principal-component weights into a regional global centile score G ε [0,1] (GCS), where larger 5 indicates a larger connected deficit (thinner-than-expected area) than in matched reference knees.

The logit *L*(*G*) was split into hinges GCS^+^ = max{*L*(*G*),0} (deficit burden above the reference median rank) and GCS^−^ = min{*L*(*G)*,0}. GCS+ and GCS− are the positive and negative parts of the logit-transformed combined reference-rank score G for connected low-centile extents, split at G=0.5. Longitudinal changes of GCS profiles quantify the spatial burden of cartilage that is unexpectedly thin or thick given the knee’s earlier scans. Increasing GCS+ or decreasing GCS− toward zero indicates a greater burden of these deficits and may signal developing focal cartilage deterioration. Decreasing GCS+ or increasingly negative GCS− indicates a lower burden, compatible with relative cartilage preservation or stabilization. Cross-sectional profiles describe connected low-centile patches relative to population expectations, while longitudinal profiles account for the knee’s earlier scans, which helps distinguish persistently thin cartilage from newly developing deficits, including loss in previously thick cartilage that remains within the population range.

A PCA-weighted extent was extracted from the GCS+ and GCS− across five thresholds for each region. Fourteen regions (12 ROIs and two plates) × two centile types × three summaries, plus the maximal suspicious area (maximal-area) per compartment, give 86 features (Supplementary Methods S3).

GCS+ and GCS− therefore describe different ranges of spatial deficit burden, rather than separate measures of baseline thick and thin cartilage. Changes over time reflect changes in relative deficit burden yet not directly quantify cartilage thickening or thinning. A lower deficit score alone does not establish recovery, which requires corroboration from thickness trajectories and clinical outcomes.

Every feature is referenced to the normative population: it describes how unusual the connected low-centile burden of a region is relative to matched normal knees. Thickening is not encoded by these one-sided features but is retained in the continuous centile maps used for deep features. Supplementary Table S5 lists every feature with its identifier, definition and direction: a higher value of any centile feature always means a larger connected thinner-than-expected area, relative to matched reference knees (P) or to the knee’s own earlier scans (H).

Each centile type gives 42 features: three summaries in each of 14 regions. PX denotes the 42 cross-sectional profiles and PL the 42 longitudinal profiles. The combined profile block P contains both, plus the femoral and tibial maximal-area features pooled across centile types (86 features total). The 42-versus-42 comparison therefore tests longitudinal against cross-sectional profiles at the same dimension. Combined-versus-separate comparisons also include the pooled maximal-area features.

### 3.6 Comparator features

Every prognostic model included the same baseline A: 11 clinical predictors (sex, race, BMI, height, alignment, injury history, physical activity, quadriceps strength, laxity, age and index KLG) and mean thickness in the 12 ROIs and two whole plates. We added the following imaging features to this baseline (Figure 1A; Table 1).

#### Regional PCA

We retained five PCs of the raw thickness map in each ROI, giving 60 scores. PCA was fitted on outer-training rows and applied to held-out rows.

#### Regional radiomics

We extracted 93 PyRadiomics 3.1.0 features from each ROI, giving 1,116 features. These included first-order, GLCM, GLRLM, GLSZM, GLDM and NGTDM measures, with a fixed thickness-bin width of 0.1 mm (van Griethuysen et al., 2017). PyRadiomics follows IBSI definitions with documented deviations, including discretization and kurtosis; an IBSI phantom check was not performed (Zwanenburg et al., 2020). Settings are given in Supplementary Methods S4.

#### Map deep features

Eleven ImageNet-pretrained networks encoded each map at 448 px: ResNet18/34/50, AlexNet, VGG16/19, GoogLeNet, Inception-v3, SqueezeNet, MobileNet-v2 and ShuffleNet-v2 (Kirchler et al., 2022; Zhao et al., 2024). We spatially averaged the last convolutional activations, averaged features across eight rotated or flipped copies, and retained ten PCs per network and map. The networks were frozen. PCA was fitted within each outer-training fold. Raw femoral and tibial maps gave 220 features (11 × 2 × 10). The four centile maps, including cross-sectional and longitudinal for each compartment, gave 440 features. Network weights and preprocessing are documented in Supplementary Table S7 and Methods S4.

Separate cross-sectional and longitudinal models each used two centile maps and 220 deep features. Thus, the comparison used the same networks and feature counts for both centile types. Including A, separate profile models had 67 predictors and separate deep-feature models 245; combined models had 111 and 465. Each imaging block was weighted by the inverse square root of its feature count: 1/√42 or 1/√86 for profiles and 1/√220 or 1/√440 for deep features. Models were fitted separately for every feature set. Combining centile types therefore changes both the number of features and the block weight. Downsampled full-reference and cross-fitted sensitivity sets are defined in Supplementary Table S6.

#### P0 and E0 in the supplementary data

P0 is the combined profile block P: 42 cross-sectional profiles, 42 longitudinal profiles and two compartmental maximum-area measures, derived from full-resolution full-reference charts. The suffix 0 identifies this chart variant. A+P0 adds these 86 profiles to the 25 clinical and regional-thickness predictors (111 total). E0 adds the profiles to the clinical/thickness baseline, 60 regional PCA scores and 1,116 radiomic features (25 + 60 + 1,116 + 86 = 1,287 predictors). The main text and figures denote these blocks and sets as P, A+P and E, respectively. Sensitivity suffixes 1 and 2 are defined in Supplementary Table S6.

### 3.7 Prediction and statistical analysis

We evaluated 12 classifiers: ridge, LASSO and Bayesian logistic regression; linear SVM with Platt calibration; LDA; GLM boosting; XGBoost; LightGBM; random forest; a single-hidden-layer neural network; naïve Bayes; and high-dimensional discriminant analysis (HDDA). Implementations, tuning grids and references are listed in Supplementary Table S8 and Methods S5.

Evaluation used participant-separated nested cross-validation with ten outer and three inner folds. Every feature set used the same folds and random seeds. Imputation and standardization were fitted on the corresponding training rows. PCA bases were fitted on outer-training rows and reused during inner tuning. Hyperparameters minimized the pooled inner out-of-fold Brier score. Most classifiers were fitted in R 4.4.0; ridge used scikit-learn 1.5.1. Deep features were extracted with PyTorch 2.7.1 and torchvision 0.22.1.

We report AUC, Brier score, accuracy, balanced accuracy, case and control precision, recall, and macro-F1 = (F1_case_ + F1_control_)/2. Threshold-based metrics used a probability cutoff of 0.5. Each estimate is the mean over the ten analysis datasets.

Paired participant bootstrapping used 2,000 resamples. Each participant received one multinomial weight shared across datasets, feature sets and classifiers. We calculated 95% confidence intervals and two-sided bootstrap P values for paired differences. Holm adjustment covered the 12 classifiers within each comparison, horizon and metric; additional adjustments covered all comparisons per metric within each statistical suite. Supplementary Table S9 specifies the resampling and multiplicity family for every result. The bootstrap held fitted charts, feature construction and out-of-fold predictions fixed.

### 3.8 Separate-centile comparisons

We compared cross-sectional and longitudinal centiles through 12 paired tests: each centile type versus A, longitudinal versus cross-sectional, combined versus each separate type, and deep features versus profiles within each type. Participants, folds, classifiers and tuning rules were unchanged. The direct comparisons used 42 profiles or 220 deep features per centile type. Figure 4 reports AUC; Supplementary Figures C1–C3 include balanced accuracy and macro-F1, and Data S5 contains all eleven metrics.

**Figure 2.**
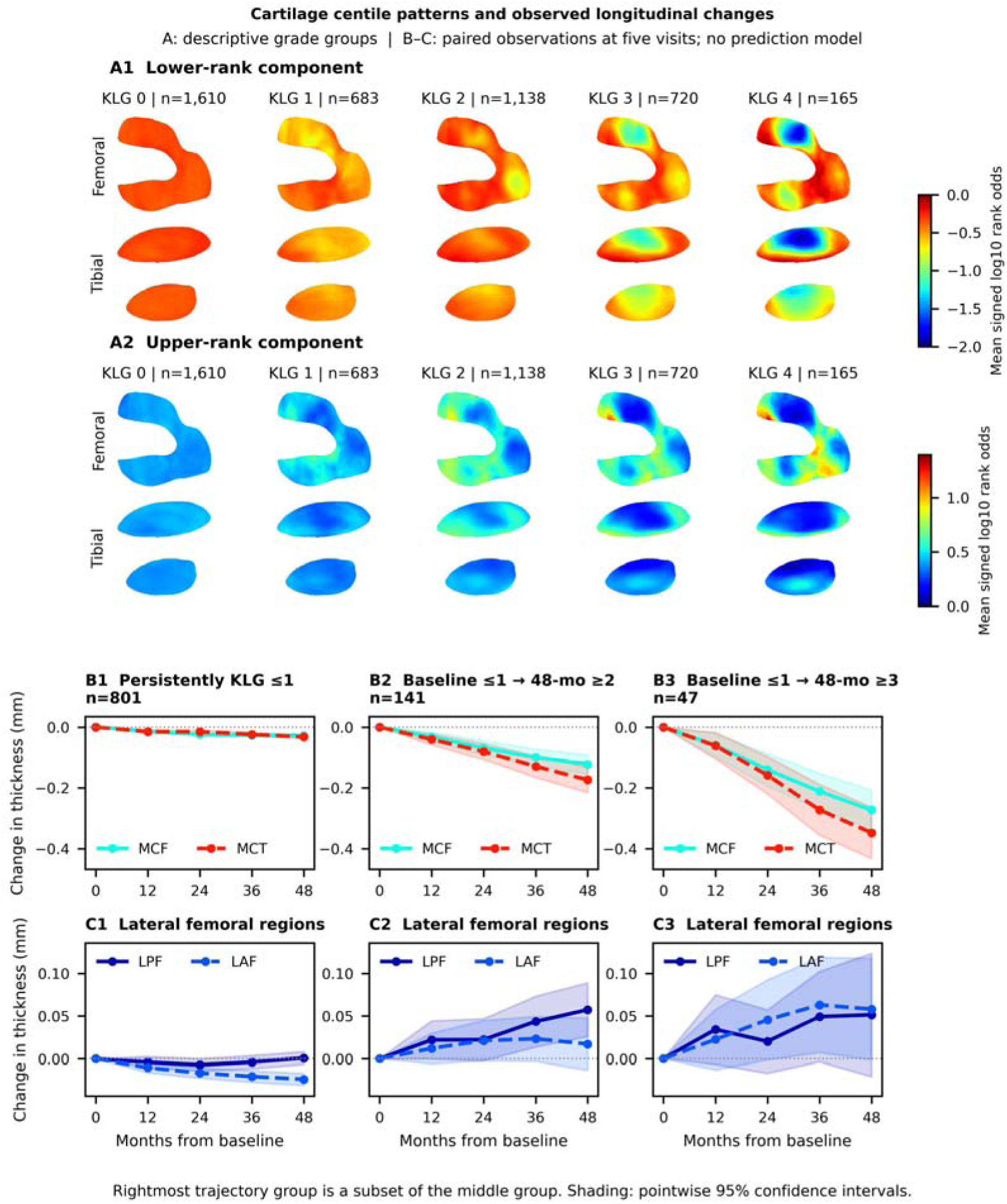
Local centile patterns and cartilage thickness change. (A1–A2) Lower- and upper-rank components of cross-sectional centile maps averaged by radiographic grade, shown as mean signed log10 rank odds; scan counts are given above the maps. Atlas orientation follows Figure 1D: medial plate above lateral plate; posterior-to-anterior runs left-to-right in the femur and right-to-left in the tibia. (B1–B3) Medial central femoral and tibial thickness change over 48 months in knees remaining KLG ≤1 (n = 801), progressing from ≤1 to ≥2 (n = 141), or progressing from ≤1 to ≥3 (n = 47, a subset of the preceding group). (C1–C3) Lateral posterior and anterior femoral change in the same groups. Shading shows pointwise 95% confidence intervals. These descriptive analyses show the anatomical patterns underlying the prediction task.

**Figure 3.**
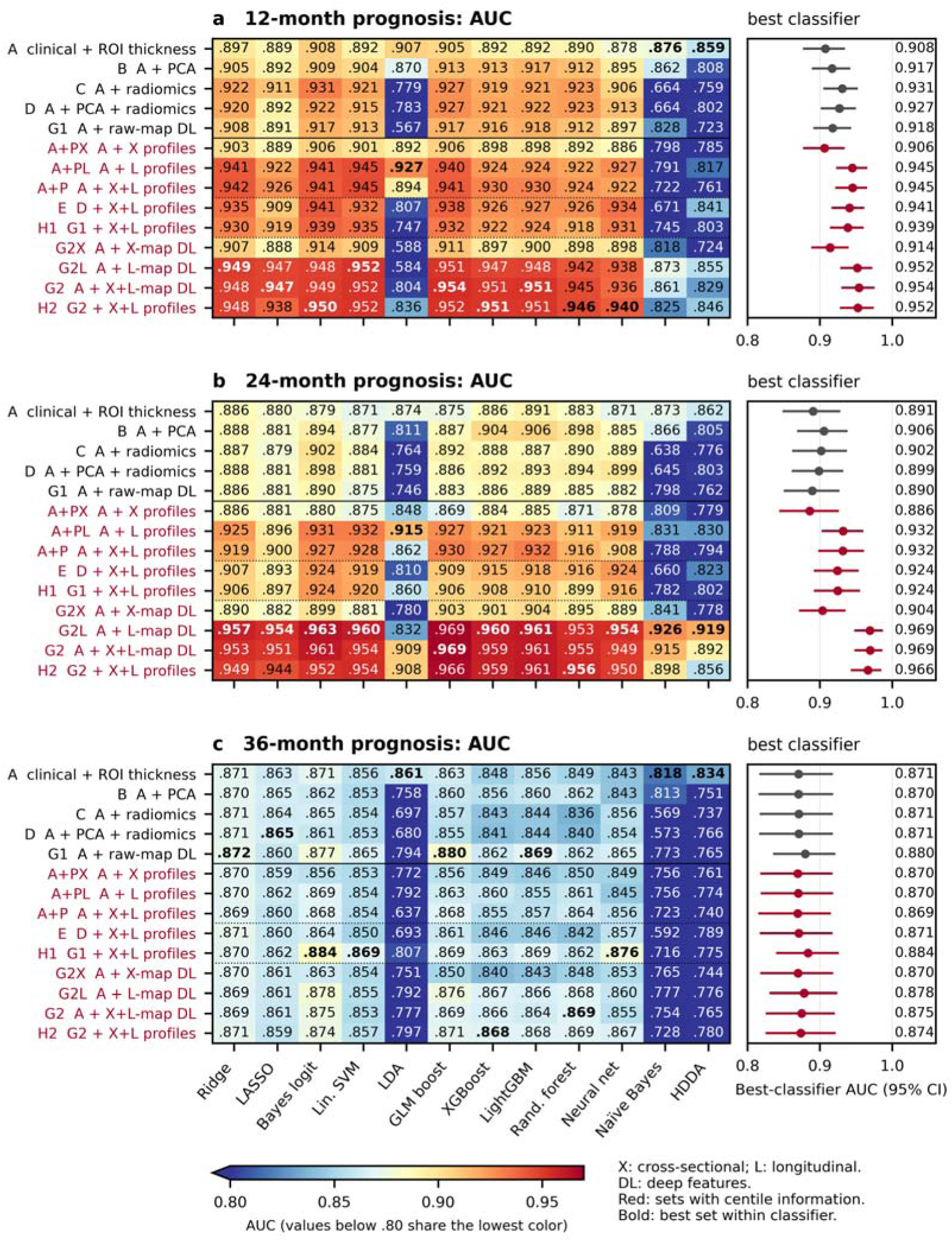
Prediction with clinical, conventional imaging and centile features. AUCs for 12 classifiers at (a) 12, (b) 24 and (c) 36 months, averaged over ten analysis datasets. All 14 displayed feature sets include A and are defined in Table 1. X denotes cross-sectional centiles, L longitudinal centiles and DL map deep features. Red labels identify centile-containing sets; bold values mark the highest unrounded AUC within each classifier and horizon. The color scale saturates below 0.80. Right, the best observed classifier for each set with its participant-bootstrap 95% confidence interval. Classifier selection was not repeated within the bootstrap.

**Figure 4.**
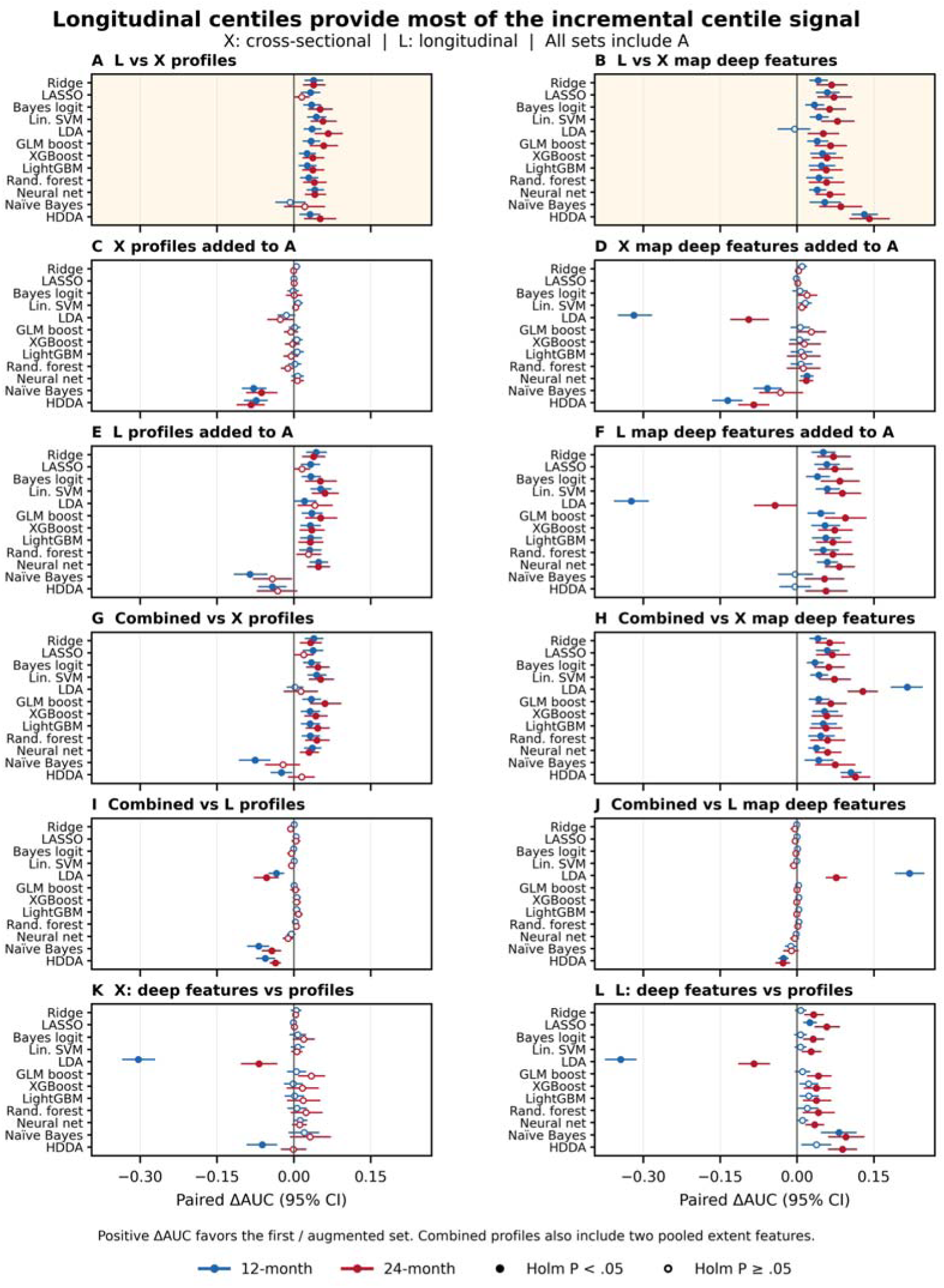
Longitudinal versus cross-sectional centiles at 12 and 24 months. Points show paired AUC differences for 12 classifiers; bars show participant-bootstrap 95% confidence intervals. Blue denotes 12 months and red 24 months. (A–B) Longitudinal versus cross-sectional features at equal dimensions: 42 profiles or 220 deep features. (C–F) Each centile type added to A. (G–J) Combined versus separate centile types. (K–L) Map deep features versus profiles within each centile type. All models include the clinical and regional-thickness baseline A; feature-set names follow Table 1. Positive differences favor the first-named or augmented set. Filled markers indicate Holm P < 0.05 across 12 classifiers within the comparison and horizon; open markers indicate P ≥ 0.05. Combined profiles include two pooled maximal extents. Balanced accuracy and macro-F1 comparisons are shown in Supplementary Figures C1–C3; exact results are in Data S5.

These tests were added after the combined-centile benchmark. Holm adjustment covered 12 classifiers per comparison, horizon and metric; the additional adjustment across the channel analysis covered 432 tests per metric (12 comparisons × 12 classifiers × three horizons). We summarize results by the median paired difference and the number of significant gains or losses. The classifiers are correlated, so these counts describe consistency across algorithms rather than independent replications. Stored cross-sectional-minus-longitudinal differences were reversed for display, with confidence-interval endpoints reversed and two-sided P values unchanged.

### 3.9 Interpretation of centile predictors

Regional centile profiles identify the anatomical region, centile type and extent of a connected cartilage deficit. We examined their contributions with probability-scale SHAP values (Lundberg and Lee, 2017), averaged equally over the five classifiers with the highest observed AUC for each outcome. SHAP describes fitted predictions; it does not identify causal tissue effects. The normative charts show covariate effects, the centile maps show where cartilage differs from its reference, and the regional profiles summarize the connected deficits (Supplementary Table S5).

## 4. Results

### 4.1 Normative charts and local cartilage changes

Expected cartilage thickness decreased with age in all regions, was greater in men than in women at every age and BMI, and followed a nonlinear age × BMI surface shared by women and men, shifted and tilted by sex-specific linear age and BMI slopes (Figure 1C; ROI-level charts for both knees in Supplementary Figures A1– A7 and A13–A14, and covariate-adjusted regional associations in Supplementary Figure A15). In women, femoral thickness declined steeply after the mid-60s at high BMI, whereas tibial thickness declined more uniformly with age; in men, higher BMI was associated with thinner cartilage at older ages. For right knees, with all three models fitted on identical data, the two-dimensional model had the lowest AIC and GCV of the three models in 13 of 14 regions, a lower AIC than the linear model in 13 of 14 regions (by up to 193 units) and the highest marginal likelihood in every region; in the lateral anterior femur the one-dimensional age model fitted best (Supplementary Table S3).

Mean local centile maps showed progressive, spatially organized thinning with increasing KLG, concentrated in the central medial femur and tibia, while upper-centile (thicker-than-expected) components also increased with grade, most visibly in posterior femoral regions (Figure 2A1–A2). In knees followed over five visits, those progressing from KLG ≤1 to ≥2 (n = 141) or ≥3 (n = 47) lost cartilage in the medial central femur and tibia faster than persistently KLG ≤1 knees (n = 801), reaching about −0.17 and −0.35 mm in the medial central tibia at 48 months, whereas the lateral posterior and anterior femur thickened by about 0.02–0.06 mm (Figure 2B–C), consistent with bidirectional early change (Buck et al., 2010). Grade-specific regional thickness, stage-specific annual thinning and thickening, observed grade transitions and attrition-sensitivity summaries are shown in Supplementary Figures A10–A12 and A16–A17.

### 4.2 Centile maps give the best 12- and 24-month prediction

Centile features gave the highest AUCs at 12 and 24 months (Figure 3). The best clinical-plus-thickness models reached 0.908 and 0.891. Adding regional centile profiles increased the best AUCs to 0.945 and 0.932, and centile-map deep features increased them to 0.954 and 0.969. Regional profiles therefore approached the best 12-month performance with 86 anatomically defined features; map deep features gave a larger advantage at 24 months.

GLM boosting with combined centile-map deep features gave the best model at both horizons: AUC 0.954 (95% CI 0.931–0.974) and 0.969 (0.949–0.985), balanced accuracy 0.917 and 0.931, and macro-F1 0.897 and 0.923. With longitudinal maps alone, the same classifier reached AUC 0.951 (0.927–0.973) and 0.969 (0.949–0.986), balanced accuracy 0.922 and 0.935, and macro-F1 0.901 and 0.926 (Table 2). Thus, longitudinal maps retained almost all of the best combined-map AUC. At 24 months, adding cross-sectional maps changed AUC by 0.00011 (95% CI −0.00288 to 0.00290; Holm P = 1.00).

**Table 2.** Performance with GLM boosting across feature sets. Values are means over ten analysis datasets with participant-bootstrap 95% confidence intervals. GLM boosting gave the best combined centile-map AUC at 12 and 24 months and is shown consistently across sets; it was not the best classifier for every set. Feature-set inputs are defined in Table 1. All classifiers and metrics are reported in Data S1 and S5.

| Horizon (mo) | Set | AUC (95% CI) | Balanced accuracy (95% CI) | Macro-F1 (95% CI) |
| --- | --- | --- | --- | --- |
| 12 | A | 0.905 (0.877–0.931) | 0.832 (0.805–0.859) | 0.807 (0.773–0.842) |
| 12 | G1 | 0.917 (0.891–0.941) | 0.851 (0.826–0.877) | 0.827 (0.792–0.861) |
| 12 | A+PX | 0.906 (0.879–0.933) | 0.840 (0.815–0.867) | 0.813 (0.780–0.848) |
| 12 | A+PL | 0.940 (0.918–0.960) | 0.866 (0.843–0.888) | 0.854 (0.824–0.883) |
| 12 | A+P | 0.941 (0.919–0.961) | 0.868 (0.845–0.892) | 0.854 (0.823–0.885) |
| 12 | G2X | 0.911 (0.885–0.937) | 0.841 (0.816–0.867) | 0.817 (0.784–0.850) |
| 12 | **G2L** | 0.951 (0.927–0.973) | **0.922 (0.899–0.942)** | **0.901 (0.869–0.930)** |
| 12 | **G2** | **0.954 (0.931–0.974)** | 0.917 (0.894–0.937) | 0.897 (0.866–0.926) |
| 12 | H2 | 0.952 (0.929–0.973) | 0.910 (0.888–0.930) | 0.893 (0.862–0.922) |
| 24 | A | 0.875 (0.832–0.917) | 0.839 (0.802–0.874) | 0.819 (0.775–0.860) |
| 24 | G1 | 0.883 (0.846–0.919) | 0.833 (0.798–0.869) | 0.815 (0.772–0.855) |
| 24 | A+PX | 0.869 (0.826–0.911) | 0.822 (0.784–0.858) | 0.802 (0.757–0.844) |
| 24 | A+PL | 0.927 (0.897–0.957) | 0.873 (0.843–0.904) | 0.863 (0.827–0.898) |
| 24 | A+P | 0.930 (0.898–0.960) | 0.881 (0.851–0.910) | 0.872 (0.837–0.905) |
| 24 | G2X | 0.903 (0.870–0.935) | 0.834 (0.800–0.870) | 0.818 (0.777–0.860) |
| 24 | **G2L** | **0.969 (0.949–0.986)** | **0.935 (0.909–0.958)** | **0.926 (0.894–0.954)** |
| 24 | **G2** | **0.969 (0.949–0.985)** | 0.931 (0.905–0.954) | 0.923 (0.892–0.950) |
| 24 | H2 | 0.966 (0.944–0.984) | 0.929 (0.902–0.953) | 0.920 (0.887–0.948) |

### 4.3 Longitudinal centiles provide most of the predictive gain

Longitudinal centiles supplied most of the improvement from centile features at 12 and 24 months. They outperformed cross-sectional centiles in both regional profiles and map deep features, and adding cross-sectional features to longitudinal features rarely improved prediction (Figure 4; Table 3).

**Table 3.** Longitudinal and cross-sectional centile comparisons. Δ is the median paired AUC difference across 12 classifiers. +/− gives the number of classifiers with significant gains/losses after Holm adjustment within each comparison and horizon. Medians summarize correlated classifiers and are not pooled effect estimates. Feature-set inputs are defined in Table 1. Combined-profile comparisons include two pooled maximal extents.

| Contrast (first – second) | 12-mo ΔAUC | 12-mo +/- | 24-mo ΔAUC | 24-mo +/- |
| --- | --- | --- | --- | --- |
| (A+PL) – (A+PX) | +0.0332 | 11/0 | +0.0406 | 10/0 |
| G2L – G2X | +0.0433 | 11/0 | +0.0651 | 12/0 |
| (A+PX) – A | +0.0018 | 0/2 | -0.0037 | 0/2 |
| (A+PL) – A | +0.0328 | 10/2 | +0.0368 | 7/0 |
| G2X – A | +0.0065 | 1/3 | +0.0110 | 1/2 |
| G2L – A | +0.0515 | 9/1 | +0.0723 | 11/1 |
| (A+P) – (A+PX) | +0.0332 | 9/2 | +0.0378 | 8/0 |
| (A+P) – (A+PL) | +0.0005 | 0/3 | -0.0042 | 0/3 |
| G2 – G2X | +0.0449 | 12/0 | +0.0650 | 12/0 |
| G2 – G2L | +0.0007 | 1/1 | -0.0024 | 1/1 |
| G2X – (A+PX) | +0.0048 | 0/2 | +0.0141 | 0/1 |
| G2L – (A+PL) | +0.0156 | 2/1 | +0.0384 | 11/1 |

#### Regional profiles

Replacing 42 cross-sectional profiles with 42 longitudinal profiles increased AUC by a median of 0.033 at 12 months and 0.041 at 24 months. Gains were significant in 11/12 and 10/12 classifiers after Holm adjustment, with no significant losses. For GLM boosting, the gains were 0.034 (95% CI 0.019–0.051) and 0.058 (0.033–0.085), both Holm P < 0.001. Longitudinal profiles improved on the clinical-plus-thickness baseline in 10/12 and 7/12 classifiers; cross-sectional profiles did not significantly improve AUC in any classifier at either horizon. Two classifiers lost AUC when longitudinal profiles were added at 12 months.

#### Map deep features

With 220 features per centile type, longitudinal maps outperformed cross-sectional maps in 11/12 classifiers at 12 months and all 12 at 24 months. Median AUC gains were 0.043 and 0.065. Median gains in balanced accuracy were 0.076 and 0.099, significant in 11/12 and 12/12 classifiers; median macro-F1 gains were 0.080 and 0.102, significant in 10/12 and 11/12. Longitudinal deep features improved AUC over baseline in 9/12 and 11/12 classifiers, compared with one classifier at each horizon for cross-sectional deep features. One classifier lost AUC with longitudinal deep features at each horizon.

#### Combining centile types

Adding longitudinal information to cross-sectional features improved AUC in 9/12 and 8/12 classifiers for profiles and in all 12 for map deep features. The reverse addition gave no significant gain for profiles and a gain in only one classifier for map deep features at each horizon. Median combined-minus-longitudinal AUC differences were +0.0005 and −0.0042 for profiles and +0.0007 and −0.0024 for deep features. Three profile models and one deep-feature model lost AUC at each horizon. These results show that the combined models gained little from cross-sectional features once longitudinal features were included.

#### Profiles versus map deep features

Longitudinal map deep features outperformed longitudinal profiles in 2/12 classifiers at 12 months and 11/12 at 24 months, with median AUC gains of 0.016 and 0.038 and one significant loss at each horizon. Cross-sectional deep features did not significantly improve on cross-sectional profiles in any classifier at either horizon. The advantage of map deep features was therefore clearest for longitudinal centiles at 24 months.

### 4.4 Longitudinal profiles outperform conventional and cross-sectional features

Longitudinal centile profiles achieved higher 12- and 24-month AUCs than regional PCA, radiomics, cross-sectional centile features and raw-map deep features in at least 11/12 classifiers per comparison (Figure 3; Data S1 and S5). This held for both regional profiles and deep features from cross-sectional centile maps. All models included clinical factors and regional mean thickness; these counts describe observed AUC differences.

Paired tests of incremental value used combined cross-sectional and longitudinal profiles (Figure 5). Adding combined profiles to the clinical-plus-thickness, PCA and radiomics model significantly increased AUC in 8/12 classifiers at 12 months and 9/12 at 24 months after Holm adjustment, with no significant losses and median gains of 0.016 and 0.023. Ridge gains were 0.015 and 0.018 (Holm P < 0.001 and P = 0.005). Adding profiles to models containing raw-map deep features improved AUC in 8/12 and 10/12 classifiers.

**Figure 5.**
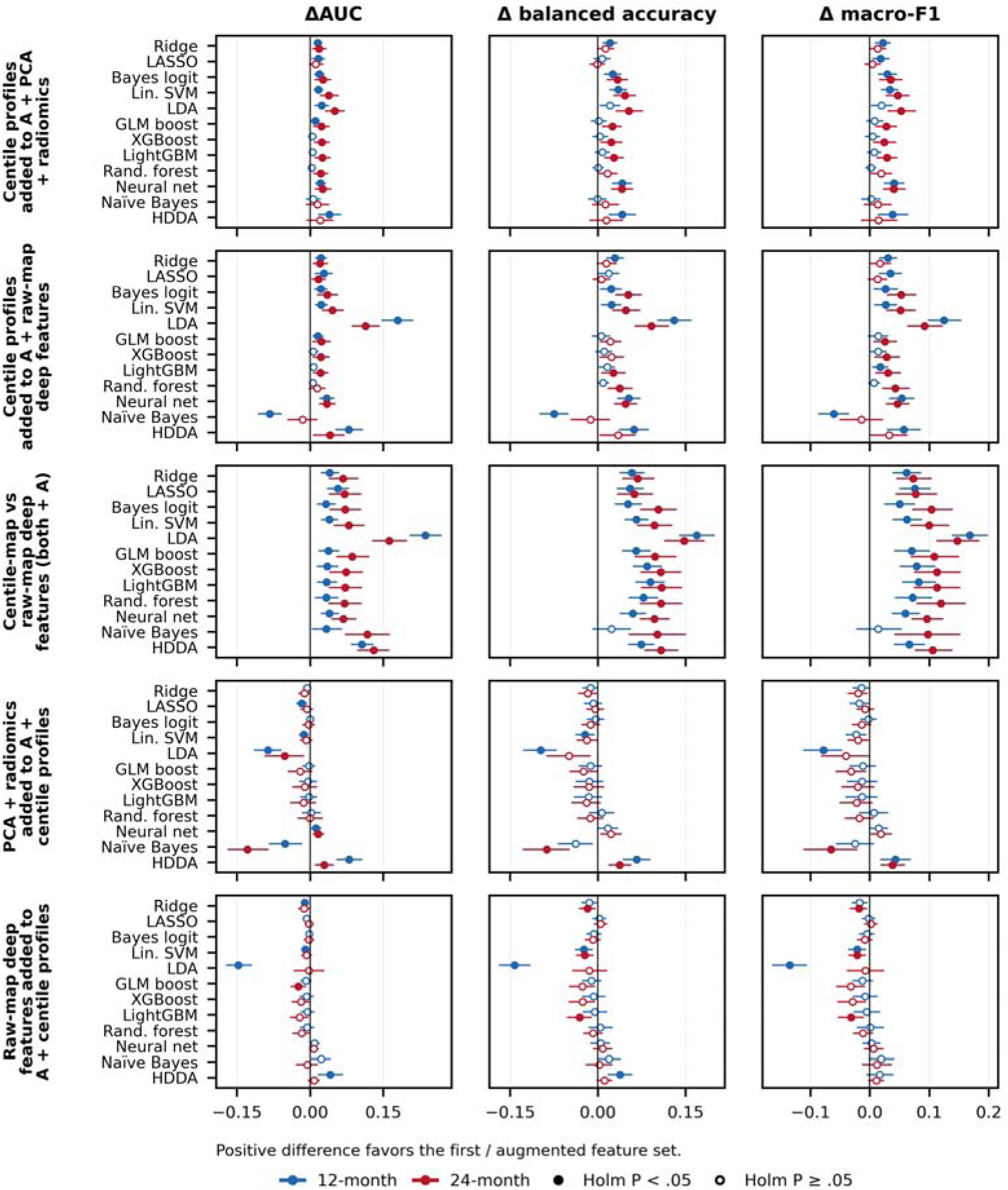
Added value of centiles, PCA, radiomics and raw-map deep features. Paired differences in AUC, balanced accuracy and macro-F1 at 12 months (blue) and 24 months (red), with participant-bootstrap 95% confidence intervals. Filled markers indicate Holm P < 0.05 across 12 classifiers within each comparison, horizon and metric. Rows compare: centile profiles added to A + PCA + radiomics (E − D); profiles added to A + raw-map deep features (H1 − G1); centile-map versus raw-map deep features (G2 − G1); PCA and radiomics added to A + profiles (E − (A+P)); and raw-map deep features added to A + profiles (H1 − (A+P)). Centile sets combine cross-sectional and longitudinal features. A is the clinical and regional-thickness baseline; complete feature-set definitions are in Table 1. Models are refitted for each feature set. G2 and G1 use different feature counts.

The reverse comparisons showed limited, inconsistent benefit. Adding PCA and radiomics to the profile model improved AUC in two classifiers at each horizon, but reduced it in four and two; median differences were −0.004 and −0.010. Ridge differences were −0.006 and −0.012 (Holm P = 0.37 and 0.08). Adding raw-map deep features improved AUC in one classifier at 12 months and none at 24 months, with losses in three and one.

Deep features of combined centile maps outperformed raw-map deep features in all 12 classifiers at both horizons. For GLM boosting, AUC gains were 0.037 and 0.086, balanced-accuracy gains 0.066 and 0.098, and macro-F1 gains 0.070 and 0.108 (all Holm P < 0.001). At 24 months, combined centile-map deep features also outperformed combined profiles in all 12 classifiers; adding profiles to centile-map deep features gave little further benefit.

### 4.5 Thirty-six-month prediction and concurrent classification

No representation consistently improved 36-month prediction over the clinical-plus-thickness baseline (Figure 3c). Neither separate centile type significantly improved baseline AUC in any classifier. Longitudinal deep features exceeded cross-sectional deep features in two classifiers, but this did not translate into a consistent baseline gain. The 36-month cohort had 80 cases, only 23 of which had a post-baseline index scan.

For concurrent KLG ≥2 versus ≤1 classification, combined profiles increased ridge AUC from 0.847 to 0.907 (difference 0.060, 95% CI 0.048–0.072; Holm P < 0.0001 within the 96-test ridge family; Supplementary Table S11). Separate cross-sectional and longitudinal models were not evaluated for concurrent classification.

### 4.6 Longitudinal profiles identify predictive cartilage deficits

Longitudinal features were the leading centile predictors at 12 and 24 months (Figure 6). At 12 months, the largest contributions came from longitudinal GCS− of the femoral plate, lateral central tibia (LCeT) and lateral posterior tibia, followed by longitudinal GCS+ of the medial anterior and central femur. At 24 months, the leading features were LCeT longitudinal GCS− and longitudinal GCS+ of the lateral posterior, medial posterior, medial central and medial anterior femur. Higher values, indicating larger connected cartilage deficits, increased predicted risk.

**Figure 6.**
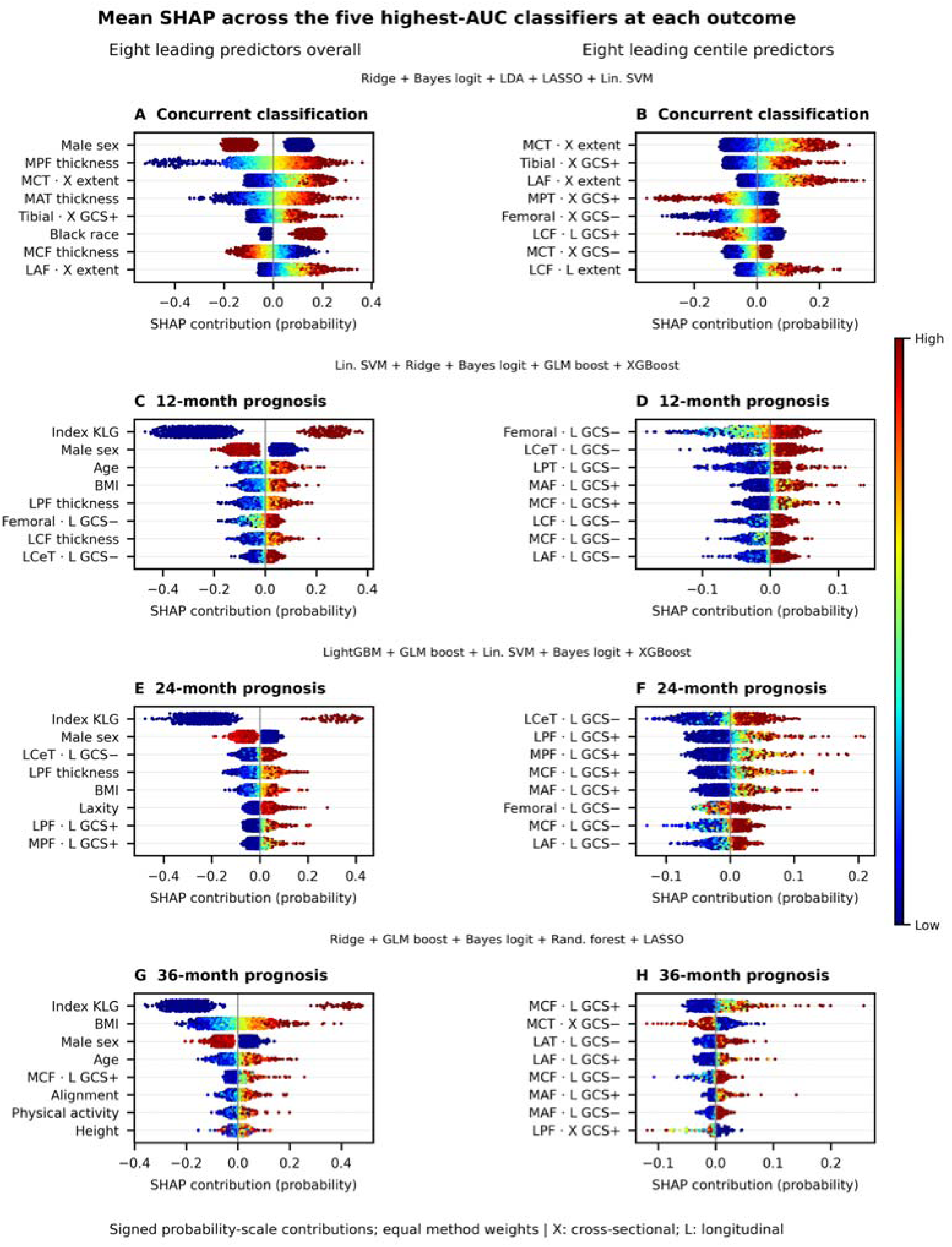
Anatomical interpretation of centile-profile predictions. Probability-scale SHAP values are averaged equally over the five classifiers with the highest observed AUC for each outcome. Left, eight leading predictors overall; right, eight leading centile predictors. Each dot represents a held-out participant, colored by the within-feature value. X and L denote cross-sectional and longitudinal centiles (P and H in the supplementary atlas). GCS+ and GCS− are positive and negative logit hinges of the regional connected-deficit rank; neither measures thickening. Higher values indicate larger or less unusually small connected deficits (Supplementary Table S5). Classifier selection and SHAP summaries are descriptive.

For concurrent classification, the leading centile features were cross-sectional extent in the medial central tibia and lateral anterior femur and cross-sectional GCS+ of the tibial plate. Centile contributions were small at 36 months. Index KLG was the strongest overall predictor: its leading SHAP magnitudes were 0.264, 0.237 and 0.207 probability units at 12, 24 and 36 months, compared with 0.027, 0.034 and 0.021 for the leading centile features. Index KLG separates many cases from controls by design. Longitudinal predominance therefore concerns the added centile information, not the complete predictor set. Per-classifier explanations and feature distributions are provided in Supplementary S10, Figures SH1–SH24 and Figure A8.

## 5. Discussion

### 5.1 Longitudinal centiles improve cartilage-based prediction

Longitudinal local centiles improve the predictive use of cartilage thickness in two ways. Regional profiles identify anatomically defined deficits and give accurate, interpretable prognosis. Deep features of centile maps give the highest discrimination. Both results depend strongly on longitudinal information: longitudinal profiles and map deep features outperform their cross-sectional counterparts, while adding cross-sectional features to longitudinal features rarely improves AUC. The improvement from centiles at 12 and 24 months is therefore largely retained when only longitudinal centiles are used.

Longitudinal centile profiles gave higher AUCs than regional PCA, radiomics, raw-map deep features and both cross-sectional centile representations in at least 11/12 classifiers per comparison at 12 and 24 months. Combined profiles also improved models that already contained conventional descriptors. Adding these descriptors to combined profiles gave limited, inconsistent gains. Moreover, applying the same pretrained networks to centile maps produced better predictions than applying them to raw thickness maps in every classifier at both horizons. The results show the value of converting thickness into reference-based deviation before extracting predictive features.

Population centiles adjust for differences in expected thickness associated with age, sex, BMI and other covariates. Longitudinal centiles also adjust for the knee’s earlier anatomy. A persistently thin region can have a low population centile without showing an unusual new deficit, while loss in an initially thick region can remain within the population range. The random-effect update makes such departures easier to detect by adjusting the local expectation using earlier scans. Conditional growth charts and longitudinal neuroimaging methods use related principles (Wei and He, 2006; Rehak Buckova et al., 2025; Bayer et al., 2026). Our results demonstrate the prognostic value of longitudinal local centiles in both anatomically defined profiles and deep features of cartilage maps.

### 5.2 Interpretable profiles and more accurate map deep features

Regional profiles summarize connected areas of thinner-than-expected cartilage. Each feature has a defined location, centile type and direction, allowing a prediction to be examined against the underlying map. Combined profiles reached best AUCs of 0.945 and 0.932 with 86 imaging features. Longitudinal profiles alone reached 0.945 and 0.932 with 42 features. At 12 months, these profiles approached the best centile-map AUC of 0.954; at 24 months, centile-map deep features increased AUC to 0.969.

The maps retain information that the regional profiles discard. They preserve continuous centile values, spatial arrangement and both thinner- and thicker-than-expected cartilage. The profiles summarize only connected low-centile deficits. This difference may explain why longitudinal map deep features outperformed longitudinal profiles in 11/12 classifiers at 24 months. We did not separately test spatial arrangement, upper-centile information or network architecture, so their individual contributions remain unresolved. The deep features also remain generic network activations; the anatomical interpretation comes from the input maps and regional profiles (Raghu et al., 2019).

The longitudinal advantage was not explained by a larger feature count in the direct comparisons: both centile types used 42 profiles or 220 deep features. The combined centile-map versus raw-map comparison used different counts, however, and combined-profile models also included two pooled extent features. Nonsignificant differences do not establish equivalence. At a first scan, only population information is available.

### 5.3 Anatomical and clinical interpretation

The leading centile predictors agree with known patterns of cartilage change. Deficits in central weight-bearing tibial cartilage and central or posterior femoral regions match the distribution of early medial cartilage loss (Wirth et al., 2009; Eckstein et al., 2010a). Our grade-specific maps and observed trajectories also show local thickening, consistent with earlier studies of early OA (Buck et al., 2010; Iriondo et al., 2021). Continuous centile maps retain both directions of change.

Local centiles rank thickness residuals; regional profiles rank the extent of connected, thinner-than-expected areas against the reference. GCS combines ranks across thresholds; it is not the thickness percentile of the whole region. GCS+ describes the upper half of this deficit-rank distribution. GCS− is negative below the median and zero at or above it, so increasing GCS− indicates a less abnormally small deficit, not cartilage thickening. These definitions make the profiles readable, although their calibration as literal percentiles across covariates and history lengths still requires held-out reference data.

The 12- and 24-month results are relevant to trial enrichment, where participants must be likely to progress during a defined treatment window (Widera et al., 2020; Guehring et al., 2021). Local maps could also support the study of pre-radiographic disease, for which MRI criteria remain under development (Luyten et al., 2018; Mahmoudian et al., 2021). Clinical use will require independent validation, risk calibration and decision analysis. The present case–control probabilities are not estimates of population incidence.

### 5.4 Limitations

#### Observation history and case selection

All controls were indexed at baseline, whereas 105/196 and 75/116 cases at 12 and 24 months had later index scans. Longitudinal centiles can therefore reflect differences in available history as well as disease-related change. Using the same classifier folds does not remove this difference. Controls also had to remain KLG 0, nonprogressing KLG-1 knees were excluded, and index KLG 1 occurred only among cases. Future validation work should include all eligible KLG ≤1 knees at common index visits, with comparable numbers and timing of earlier scans and separate evaluation by index KLG. The current results do not establish the longitudinal gain under matched histories.

#### Normative reference

The full-resolution charts were fitted before outcome partitioning and included the prediction controls. We assessed this overlap in sensitivity analyses, by averaging 3 × 3 pixel blocks to 8,305 pixels and refitting the GAMMs on the full reference or two participant-disjoint halves (Supplementary S9). Scoring each control with a chart that excluded it reduced the ridge AUC gain from profiles by 0.002 at 12 months and 0.003 at 24 months; gains remained 0.029 (95% CI 0.017–0.044) and 0.017 (0.005–0.031). Across classifiers, reference inclusion increased AUC by at most 0.010 and 0.012, significantly in three and one classifiers (Tables S19–S20).

The combined centile-map deep-feature analysis was also repeated with cross-fitted charts and all 12 classifiers (Tables S21–S22). Best AUCs were 0.963 and 0.973. No classifier had a Holm-significant loss relative to full-resolution full-reference charts; median differences were −0.000 and +0.001, and the largest numerical losses were −0.013 and −0.015 for naïve Bayes. Cross-fitted centile-map deep features still outperformed raw-map deep features in 11/12 and 12/12 classifiers. These checks support the combined-centile result. Cross-fitting has not yet been repeated for the separate longitudinal-versus-cross-sectional comparison.

#### Statistical scope

PCA bases were reused within inner tuning rather than refitted in each inner fold. These choices limit separation of the effects of history conditioning, reference construction and feature processing. Independent normative cohorts and calibration by history length are needed.

#### Other limits

The study used right knees from one cohort and an event-anchored case–control design. Segmentation and registration errors can affect local centiles. Frozen ImageNet features do not cover all end-to-end or temporal deep-learning methods. The completed paired tests did not directly compare the 220 longitudinal deep features with the 220 raw-map deep features. Raw longitudinal change and slope predictors were not separately compared, and the random-intercept model does not estimate knee-specific slopes. No representation consistently improved 36-month prediction; the smaller case sample and fewer knees with earlier scans limit interpretation of that result.

## 6. Conclusion

Longitudinal normative centiles turn cartilage thickness maps into reference-based deviation maps. Longitudinal profiles achieved higher 12- and 24-month AUCs than regional PCA, radiomics, cross-sectional centile features and raw-map deep features in most classifiers. Combined centile profiles added prognostic value to conventional descriptors, with little benefit from the reverse addition. Longitudinal centiles retained almost all combined-centile performance. Regional profiles give anatomically interpretable prognosis, approaching the best accuracy at 12 months; centile-map deep features give the highest discrimination at both horizons. Validation with comparable observation histories will determine how this advantage transfers to clinical prediction.

## Data and code availability

OAI data are available from the NIMH Data Archive (https://nda.nih.gov/oai/). Analysis code for normative modeling, centile scoring, feature extraction, classification and statistical comparisons is available from the authors. Data S1–S4 contain the benchmark, sensitivity and SHAP results. Data S5 contains separate-centile performance, paired differences, confidence intervals and adjusted P values.

## Author contributions

TF. L., M. N., and H. Z. designed the study. M. N. supervised the imaging analyses and H. Z. supervised the statistical analyses. TF. L. and TY. L. analyzed the data. B. C., C. H., Z. S. and Z. X. processed imaging data. A. E. N., D. N., Y. M. G. and G. G. provided clinical interpretations of the analyses and reviewed the paper. TF. L. and TY. L. wrote the manuscript with feedback from all authors.

## Acknowledgements

Data and/or research tools used in the preparation of this manuscript were obtained and analyzed from the controlled access datasets distributed from the Osteoarthritis Initiative (OAI), a data repository housed within the NIMH Data Archive (NDA). OAI is a collaborative informatics system created by the National Institute of Mental Health and the National Institute of Arthritis, Musculoskeletal and Skin Diseases (NIAMS) to provide a worldwide resource to quicken the pace of biomarker identification, scientific investigation and OA drug development. Dataset identifier(s): NIMH Data Archive Collection ID: 2343. The OAI is a public–private partnership comprising five contracts (N01-AR-2-2258, N01-AR-2-2259, N01-AR-2-2260, N01-AR-2-2261, N01-AR-2-2262) funded by the National Institutes of Health. We thank the individuals represented in the OAI datasets for their participation and the research teams for their work in collecting, processing and disseminating these datasets for analysis.

Research reported in this publication was partially supported by the National Institutes of Health (NIH) under award numbers R01AR072013 (TY. L., B. C. and M. N.); RF1AG082938, R01AR082684, RF1AG098697, R01AG085581, and R01MH136055 (TF. L. and H. Z.); and R01DE024450, R21HD120911 and K01AG095286 (TF. L.). We also acknowledge support from NIH/NIAMS P30AR072580 (A. E. N. and Y. M. G.). The content is solely the responsibility of the authors and does not necessarily represent the official views of the National Institutes of Health. The funders had no role in study design, data collection and analysis, decision to publish or preparation of the manuscript.

