## Supplementary materials for "Benchmarking longitudinal local centiles for radiographic knee osteoarthritis prognosis"

### Contents

#### Supplementary sections

**S1** — Cohorts and sampling

**S2** — Normative model specification and model comparison

**S3** — Local centiles and connected-deficit features

**S4** — Comparator representations

**S5** — Classifiers, nested cross-validation and tuning

**S6** — Performance metrics and statistical inference

**S7** — Complete performance results

**S8** — Paired comparisons

**S9** — Sensitivity analysis: reference membership (downsampled cross-fitted versus full-reference charts)

**S10** — SHAP atlas with detailed interpretation

**S11** — Reference-chart and descriptive atlas (Figures A1–A17)

**S12** — Separate-channel centile comparison

Supplementary data files

Supplementary references

#### Excel data files (Data S1–S5)

The following Excel workbooks accompany this supplement.

**Data S1 — DataS1.xlsx.** Complete performance results for the PCA/radiomics and deep-feature benchmarks.

**Data S2 — DataS2.xlsx.** Paired feature-set comparisons, confidence intervals and adjusted P values.

**Data S3 — DataS3.xlsx.** Sensitivity to reference membership and chart resolution.

**Data S4 — DataS4.xlsx.** SHAP importance, selected classifiers and explanation stability.

**Data S5 — DataS5.xlsx.** Separate cross-sectional and longitudinal centile results, paired comparisons and figure data.

#### Supplementary tables (Tables S1–S23)

**Table S1** — Participants and outcome sampling

**Table S2** — Characteristics of cases and controls for concurrent classification and each prognostic horizon (unique participants; mean ± SD or n (%))

**Table S3** — AIC (top) and GCV (bottom) of models M1–M3 for ROI-mean thickness, left (L) and right (R) knees; the minimum per knee is in bold

**Table S4** — Null pools of the local centiles by compartment and channel (Algorithm 1)

**Table S5** — Global centile features and accompanying predictors: identifiers, definitions and interpretation

**Table S6** — Prognostic feature blocks and feature sets

**Table S7** — Pretrained networks used for deep transfer-learning features (torchvision 0.22.1 IMAGENET1K_V1 weights; PyTorch 2.7.1)

**Table S8** — The 12 classifiers: abbreviation used in figures and tables, full name, implementation, tuned hyperparameters (grid searched by inner cross-validation) and key references

**Table S9** — Statistical inference and multiplicity adjustments

**Table S10** — Ridge logistic regression, raw-map comparator benchmark (all metrics; estimate and 95% CI)

**Table S11** — Concurrent classification (KLG ≥2 vs ≤1; 2,249 scans, 1,923 participants; five participant-grouped partitions): ridge AUC and Brier score (95% CI)

**Table S12** — AUC for each feature set and classifier

**Table S13** — Balanced accuracy for each feature set and classifier

**Table S14** — Macro-F1 for each feature set and classifier

**Table S15** — Best model (GLM boosting with centile-map deep features, feature set G2): all metrics with 95% CIs

**Table S16** — Paired differences between feature sets for ridge logistic regression, with 95% confidence intervals

**Table S17** — Summary of paired comparisons across the 12 classifiers (AUC): number of classifiers with a Holm-significant gain (+) or loss (−), and median difference

**Table S18** — Per-classifier paired AUC differences (95% CI; Holm P across classifiers) for the key deep-feature comparisons at 12 and 24 months

**Table S19** — Ridge AUC of clinical + ROI mean thickness (feature set A) and with centile profiles (feature set A+P) under each reference arm, and the centile increment

**Table S20** — Across the 12 classifiers: number with Holm-significant gain (+) or loss (−), median and range of differences

**Table S21** — Best classifier per setting for centile-map deep features: AUC, balanced accuracy and macro-F1 (95% CI)

**Table S22** — F2 check across the 12 classifiers: number with Holm-significant gain (+) or loss (−), median and range of paired differences

**Table S23** — Paired performance differences for the separate-centile comparisons

#### Supplementary figures

##### SHAP atlas (Figures SH1–SH24)

**Figure SH1** — Concurrent classification (KLG ≥2 vs ≤1) — Equal-weight average of the five classifiers

**Figure SH2** — Concurrent classification (KLG ≥2 vs ≤1) — Ridge

**Figure SH3** — Concurrent classification (KLG ≥2 vs ≤1) — Bayesian logistic

**Figure SH4** — Concurrent classification (KLG ≥2 vs ≤1) — LDA

**Figure SH5** — Concurrent classification (KLG ≥2 vs ≤1) — LASSO

**Figure SH6** — Concurrent classification (KLG ≥2 vs ≤1) — Linear SVM

**Figure SH7** — 12-month prognosis — Equal-weight average of the five classifiers

**Figure SH8** — 12-month prognosis — Linear SVM

**Figure SH9** — 12-month prognosis — Ridge

**Figure SH10** — 12-month prognosis — Bayesian logistic

**Figure SH11** — 12-month prognosis — GLM boosting

**Figure SH12** — 12-month prognosis — XGBoost

**Figure SH13** — 24-month prognosis — Equal-weight average of the five classifiers

**Figure SH14** — 24-month prognosis — LightGBM

**Figure SH15** — 24-month prognosis — GLM boosting

**Figure SH16** — 24-month prognosis — Linear SVM

**Figure SH17** — 24-month prognosis — Bayesian logistic

**Figure SH18** — 24-month prognosis — XGBoost

**Figure SH19** — 36-month prognosis — Equal-weight average of the five classifiers

**Figure SH20** — 36-month prognosis — Ridge

**Figure SH21** — 36-month prognosis — GLM boosting

**Figure SH22** — 36-month prognosis — Bayesian logistic

**Figure SH23** — 36-month prognosis — Random forest

**Figure SH24** — 36-month prognosis — LASSO

##### Reference-chart and descriptive atlas (Figures A1–A17)

**Figure A1** — Right femoral subregional reference surfaces

**Figure A2** — Right tibial subregional reference surfaces

**Figure A3** — Left femoral subregional reference surfaces

**Figure A4** — Left tibial subregional reference surfaces

**Figure A5** — Left femoral subregional age–BMI contours

**Figure A6** — Left tibial subregional age–BMI contours

**Figure A7** — Left whole-compartment age–BMI contours

**Figure A8** — Group distributions of leading centile predictors

**Figure A9** — Added centile discrimination within twelve classifiers

**Figure A10** — Observed 48-month thickness change by baseline grade

**Figure A11** — Observed radiographic-grade transitions over five visits

**Figure A12** — Complete-visit and available-pair longitudinal summaries

**Figure A13** — Right femoral subregional age–BMI contours

**Figure A14** — Right tibial subregional age–BMI contours

**Figure A15** — Covariate-adjusted regional thickness associations

**Figure A16** — Regional thickness across radiographic grades

**Figure A17** — Stage-specific annual thinning and thickening

##### Separate-channel comparisons (Figures C1–C3)

**Figure C1** — Prognostic value of separate and combined centile profiles

**Figure C2** — Prognostic value of separate and combined centile-map deep features

**Figure C3** — Centile-map deep features compared with regional profiles

### S1. Cohorts and sampling

The Osteoarthritis Initiative (OAI) enrolled 4,796 adults aged 45–79 years and followed them for up to 96 months. The paired femoral and tibial map collection comprised 21,383 scans from 4,323 participants; 21,369 scans entered profile compression. The normative reference comprised 5,045 scans from 957 participants with persistently KLG-0 knees. These samples overlap. Concurrent classification included 450, 449, 450, 450 and 450 scans at KLG 0, 1, 2, 3 and 4, respectively. All scans from a participant were assigned to the same cross-validation fold.

**Prognostic cohorts.** We used observed right-knee Kellgren–Lawrence grades from the central Boston University readings (OAI files kxr_sq_bu), ordered by visit. Missing grades were not imputed. Cases and controls were defined separately for each prediction horizon $h\in\{12,24,36\}$ months:

- *Cases (incident radiographic OA).* The outcome was the first observed KLG ≥2, preceded by at least one radiograph; every earlier observed grade had to be ≤1. Knees with KLG ≥2 at their first reading were excluded. The index MRI was the latest scan acquired at least $h$ months before the outcome, allowing a further 12 months when the scan exactly $h$ months before the outcome was unavailable. The resulting lead time was $h$ months, or $h+12$ months for 1, 1 and 2 cases in the 12-, 24- and 36-month cohorts. Outcomes occurred at months 12/24/36/48 in the 12-month cohort (91/36/47/22 cases), months 24/36/48 in the 24-month cohort (41/52/23) and months 36/48 in the 36-month cohort (55/25). Outcome grades were 2 or 3, except for two KLG-4 cases in the 12-month cohort. Index grades were 0 or 1; KLG 1 accounted for 142/196, 78/116 and 48/80 cases at 12, 24 and 36 months. Index visits were at months 0/12/24/36 for the 12-month cohort (91/37/46/22 cases), months 0/12/24 for the 24-month cohort (41/53/22) and months 0/12 for the 36-month cohort (57/23).
- *Controls (no radiographic OA).* Controls required an observed KLG of 0 at month $h$ and at every other available visit. A single KLG-1 reading excluded the knee. Visits with missing grades were allowed; the median last observed visit was at 48 months, with follow-up extending to 96 months. The index MRI was the baseline scan (month 0), and the outcome assessment was at month $h$, giving a lead time of $h$ months. All 847, 782 and 716 controls in the respective cohorts had an observed KLG of 0 at month $h$.
- *Excluded knees.* The analysis excluded knees whose maximum observed grade was 1 (712 participants) and knees with KLG ≥2 at their first radiograph. The comparison therefore concerns incident radiographic OA versus persistently KLG-0 knees. All controls belonged to the normative reference sample; Section S9 examines the effect of this overlap.
- *Covariates.* Clinical covariates, height and physical-activity variables were taken from the latest visit at or before the index scan.
- *Analysis datasets.* Controls (847, 782 and 716) outnumbered cases (196, 116 and 80). We divided the controls into ten disjoint subsets, stratified by BMI using createFolds in caret 6.0.94 (R 4.4.0), and paired each subset with all cases. The resulting case:control ratios were approximately 2.3:1, 1.5:1 and 1.1:1. Each control appeared in one dataset and each case in all ten. Models were fitted and evaluated by participant-separated nested cross-validation within each dataset (S5), and performance was averaged across datasets.

**Table S1.** Participants and outcome sampling.

| Characteristic | Concurrent | 12 months | 24 months | 36 months |
| --- | --- | --- | --- | --- |
| Unique participants | 1,923 | 1,043 | 898 | 796 |
| Knees per dataset | 2,249 scans | 279–282 | 194–195 | 150–153 |
| Cases per dataset | 1,350 scans | 196 | 116 | 80 |
| Controls per dataset | 899 scans | 83–86 | 78–79 | 70–73 |
| Women, n (%) | 1,125 (58.5) | 596 (57.1) | 513 (57.1) | 454 (57.0) |
| Age (years) | 65.9 ± 9.1 | 59.1 ± 9.0 | 58.7 ± 9.0 | 58.5 ± 9.0 |
| BMI (kg/m²) | 28.7 ± 5.0 | 27.1 ± 4.5 | 26.8 ± 4.5 | 26.8 ± 4.4 |
| Analysis datasets | 5 repetitions of the same 2,249 scans (different random fold partitions) | 10 (all cases + one of 10 disjoint control subsets) | 10 (all cases + one of 10 disjoint control subsets) | 10 (all cases + one of 10 disjoint control subsets) |
| Eligible controls (split into 10 subsets) | not split: all 899 control scans used in every repetition | 847 | 782 | 716 |

Age and BMI are mean ± SD. Demographics use unique participants across samples, with the earliest included scan for classification. Across prognostic horizons, 1,306 distinct index scans represented 1,129 participants; every dataset contains all cases, while each control belongs to exactly one of the ten datasets. Index KLG 1 occurred in 142/196, 78/116 and 48/80 cases; all controls had KLG 0. MRI used sagittal water-excited 3D DESS on 3-T Siemens Trio systems (nominal 384 × 384 × 160 matrix, 0.36 × 0.36 × 0.7 mm). Projected femoral and tibial maps had 38,009 and 36,715 valid pixels.

**Table S2.** Characteristics of cases and controls for concurrent classification and each prognostic horizon (unique participants; mean ± SD or n (%)). P: Welch t test (continuous) or χ² test (Fisher exact test when an expected cell is small), cases versus controls; descriptive, not adjusted for multiplicity.

| Characteristic | Cases | Controls | P |
| --- | --- | --- | --- |
| **Concurrent classification (KLG ≥2 vs ≤1)** | **1,025** | **898** |  |
| Age, years | 66.1 ± 8.9 | 65.5 ± 9.3 | 0.145 |
| Women, n (%) | 610 (59.5) | 515 (57.3) | 0.361 |
| Black race, n (%) | 229 (22.3) | 113 (12.6) | <0.001 |
| BMI, kg/m² | 29.8 ± 5.0 | 27.5 ± 4.7 | <0.001 |
| Height, cm | 167.8 ± 9.1 | 167.6 ± 9.3 | 0.627 |
| **12-month prognosis** | **196** | **847** |  |
| Age, years | 61.9 ± 8.9 | 58.5 ± 8.9 | <0.001 |
| Women, n (%) | 120 (61.2) | 476 (56.2) | 0.230 |
| Black race, n (%) | 27 (13.8) | 93 (11.0) | 0.327 |
| BMI, kg/m² | 29.5 ± 4.6 | 26.5 ± 4.3 | <0.001 |
| Height, cm [missing 15] | 168.1 ± 9.3 | 168.4 ± 9.5 | 0.701 |
| Knee alignment, absolute angle (°) [missing 320] | 2.6 ± 2.0 | 2.1 ± 1.6 | 0.005 |
| Right-knee injury history, n (%) | 56 (28.6) | 193 (22.8) | 0.105 |
| Physical activity, PASE score [missing 7] | 161.2 ± 86.2 | 169.8 ± 82.9 | 0.208 |
| Quadriceps strength (maximum knee extension force), N [missing 191] | 362.5 ± 131.3 | 370.1 ± 137.1 | 0.552 |
| Knee laxity grade (0–2) [missing 247] | 0.4 ± 0.6 | 0.3 ± 0.5 | 0.101 |
| Index KLG 1, n (%) | 142 (72.4) | 0 (0.0) | — |
| **24-month prognosis** | **116** | **782** |  |
| Age, years | 61.7 ± 8.3 | 58.3 ± 9.0 | <0.001 |
| Women, n (%) | 80 (69.0) | 433 (55.4) | 0.008 |
| Black race, n (%) | 16 (13.8) | 84 (10.7) | 0.414 |
| BMI, kg/m² | 29.6 ± 4.9 | 26.4 ± 4.3 | <0.001 |
| Height, cm [missing 13] | 166.6 ± 8.5 | 168.6 ± 9.5 | 0.019 |
| Knee alignment, absolute angle (°) [missing 256] | 2.7 ± 2.1 | 2.1 ± 1.5 | 0.025 |
| Right-knee injury history, n (%) [missing 2] | 33 (28.9) | 186 (23.8) | 0.279 |
| Physical activity, PASE score [missing 6] | 154.7 ± 80.6 | 172.5 ± 82.5 | 0.030 |
| Quadriceps strength (maximum knee extension force), N [missing 167] | 350.7 ± 132.0 | 371.6 ± 139.9 | 0.235 |
| Knee laxity grade (0–2) [missing 153] | 0.5 ± 0.6 | 0.3 ± 0.5 | 0.034 |
| Index KLG 1, n (%) | 78 (67.2) | 0 (0.0) | — |
| **36-month prognosis** | **80** | **716** |  |
| Age, years | 60.9 ± 8.3 | 58.2 ± 9.0 | 0.009 |
| Women, n (%) | 56 (70.0) | 398 (55.6) | 0.019 |
| Black race, n (%) | 10 (12.5) | 78 (10.9) | 0.805 |
| BMI, kg/m² | 29.4 ± 4.8 | 26.5 ± 4.3 | <0.001 |
| Height, cm [missing 11] | 167.4 ± 8.8 | 168.4 ± 9.3 | 0.372 |
| Knee alignment, absolute angle (°) [missing 214] | 2.7 ± 2.2 | 2.2 ± 1.6 | 0.087 |
| Right-knee injury history, n (%) | 25 (31.2) | 173 (24.2) | 0.210 |
| Physical activity, PASE score [missing 5] | 176.3 ± 93.5 | 171.7 ± 82.2 | 0.675 |
| Quadriceps strength (maximum knee extension force), N [missing 129] | 356.3 ± 115.4 | 370.5 ± 139.4 | 0.409 |
| Knee laxity grade (0–2) [missing 164] | 0.5 ± 0.6 | 0.3 ± 0.5 | 0.072 |
| Index KLG 1, n (%) | 48 (60.0) | 0 (0.0) | — |

For concurrent classification, cases had KLG ≥2 and controls KLG ≤1; the table reports each participant’s earliest included scan. Clinical predictors were age, sex, race, BMI and height. For prognosis, each incident case and persistently KLG-0 control is counted once in the table, although cases appeared in all ten analysis datasets. Covariates refer to the index scan. All controls had index KLG 0 by definition. Bracketed values give missing counts; missing covariates were median-imputed within training folds.

### S2. Normative model specification and model comparison

At each pixel $v$ the reference model (M1) was

$$y_{i,v}\left( t_{ij},b_{ij} \right)=f_{v}\left( t_{ij},b_{ij} \right)+\beta_{0,v}g_{i}+\beta_{1,v}g_{i}b_{ij}+\beta_{2,v}g_{i}t_{ij}+\beta_{3,v}r_{i}+\beta_{4,v}h_{ij}+\alpha_{i,v}+\varepsilon_{i,v}\left( t_{ij} \right),$$

with $f_{v}$ a tensor-product smooth of age $t$ and BMI $b$, $g_{i}$ male sex, $r_{i}$ Black race, $h_{ij}$ height, $\alpha_{i,v}\sim N\left( 0,\tau_{v}^{2} \right)$ and $\varepsilon\sim N\left( 0,\sigma_{v}^{2} \right)$. Models were fitted by maximum likelihood with the R package mgcv (R 3.6.0 for the full-resolution full-reference charts; mgcv 1.9.3 with R 4.4.0 for the downsampled charts of S9) using the formula ( y ~ Race + H + te(Age, BMI) + Gender*Age + Gender*BMI + s(ID, bs = "re")). Two alternatives were compared:

$$\text{M2: }y=\beta_{0}g+\beta_{1}gb+\beta_{2}gt+\beta_{3}r+\beta_{4}h+\beta_{5}t+\beta_{6}b+\beta_{7}bt+\alpha+\varepsilon,$$

$$\text{M3: }y=f\left( t \right)+\beta_{0}g+\beta_{1}gb+\beta_{2}gt+\beta_{3}r+\beta_{4}h+\beta_{6}b+\beta_{7}bt+\alpha+\varepsilon.$$

M2 uses linear covariate terms and an age × BMI interaction. M3 replaces the two-dimensional surface with a smooth age term and a linear age × BMI interaction. We compared these models using AIC and generalized cross-validation (GCV) for regional mean thickness (Table S3). M1 had lower AIC than M2 in every region of both knees except the right lateral anterior femur (ROI 3), with differences of up to 193 units. GCV was lower for M1 than M2 throughout. In the right knees used for prediction, M1 had the lowest AIC and GCV in 13 of 14 regions; M3 performed best in ROI 3, with an AIC advantage of 63.6. M1 improved AIC over M3 by more than 10 units in 12 right-knee regions, reaching 121 units for the whole femur. In left knees, M1 had the lowest AIC in 10 regions and M3 in four (ROIs 3, 9, 11 and 12; differences of 1.6–5.8); M1 had the lowest GCV in nine regions and M3 in five. We selected M1 for its joint age–BMI surface and its better fit in 13 of 14 right-knee regions by both criteria. M1 also had the highest marginal likelihood in every region of both knees. M1 and M3 use different smooth bases and penalties and are not formally nested.

**Table S3.** AIC (top) and GCV (bottom) of models M1–M3 for ROI-mean thickness, left (L) and right (R) knees; the minimum per knee is in bold. ROI numbering follows main Figure 1D (1 LPF, 2 LCF, 3 LAF, 4 MAF, 5 MCF, 6 MPF, 7 LPT, 8 LCeT, 9 LAT, 10 MPT, 11 MCT, 12 MAT).

| ROI | M1-L | M1-R | M2-L | M2-R | M3-L | M3-R |
| --- | --- | --- | --- | --- | --- | --- |
| 1 | **−5,060.8** | **−7,003.1** | −4,983.5 | −6,889.1 | −4,983.5 | −6,927.9 |
| 2 | **−8,729.7** | **−11,884.9** | −8,725.5 | −11,807.4 | −8,725.4 | −11,819.8 |
| 3 | −7,702.3 | −7,333.2 | −7,659.3 | −7,338.0 | **−7,705.7** | **−7,396.7** |
| 4 | **−2,547.9** | **−1,851.9** | −2,512.6 | −1,744.5 | −2,546.9 | −1,773.4 |
| 5 | **−6,986.5** | **−3,956.5** | −6,897.7 | −3,857.0 | −6,937.4 | −3,897.0 |
| 6 | **−7,394.5** | **−6,571.4** | −7,382.2 | −6,540.4 | −7,393.8 | −6,559.7 |
| 7 | **931.0** | **−1,450.3** | 1,053.0 | −1,410.4 | 1,053.0 | −1,428.8 |
| 8 | **−2,277.5** | **−6,239.5** | −2,211.1 | −6,118.1 | −2,266.4 | −6,203.2 |
| 9 | −508.1 | **−1,584.2** | −506.9 | −1,542.8 | **−512.2** | −1,563.0 |
| 10 | **−2,328.4** | **1,010.1** | −2,327.6 | 1,047.8 | −2,319.1 | 1,028.7 |
| 11 | −6,570.2 | **−3,478.3** | −6,533.3 | −3,461.3 | **−6,571.8** | −3,469.5 |
| 12 | −3,288.5 | **−3,274.8** | −3,270.8 | −3,249.7 | **−3,294.3** | −3,249.4 |
| Tibia | **−5,235.5** | **−5,455.9** | −5,180.2 | −5,349.0 | −5,180.2 | −5,380.9 |
| Femur | **−8,396.1** | **−7,414.1** | −8,303.0 | −7,221.5 | −8,342.1 | −7,293.4 |

| ROI | M1-L | M1-R | M2-L | M2-R | M3-L | M3-R |
| --- | --- | --- | --- | --- | --- | --- |
| 1 | **0.0208** | **0.0152** | 0.0211 | 0.0155 | 0.0211 | 0.0154 |
| 2 | **0.0096** | **0.0058** | **0.0096** | 0.0059 | **0.0096** | 0.0059 |
| 3 | **0.0119** | **0.0140** | 0.0120 | 0.0142 | **0.0119** | **0.0140** |
| 4 | **0.0356** | **0.0421** | 0.0358 | 0.0430 | **0.0356** | 0.0427 |
| 5 | **0.0139** | **0.0277** | 0.0141 | 0.0282 | 0.0140 | 0.0280 |
| 6 | **0.0127** | **0.0163** | **0.0127** | **0.0163** | **0.0127** | **0.0163** |
| 7 | **0.0743** | **0.0456** | 0.0761 | 0.0460 | 0.0761 | 0.0458 |
| 8 | **0.0376** | **0.0177** | 0.0382 | 0.0181 | 0.0378 | 0.0178 |
| 9 | 0.0548 | **0.0444** | 0.0548 | 0.0448 | **0.0547** | 0.0446 |
| 10 | **0.0371** | **0.0738** | **0.0371** | 0.0743 | 0.0372 | 0.0741 |
| 11 | **0.0151** | **0.0304** | 0.0153 | 0.0305 | **0.0151** | 0.0305 |
| 12 | 0.0304 | **0.0317** | 0.0305 | 0.0319 | **0.0303** | 0.0319 |
| Tibia | **0.0201** | **0.0207** | 0.0204 | 0.0211 | 0.0204 | 0.0210 |
| Femur | **0.0103** | **0.0140** | 0.0105 | 0.0145 | 0.0104 | 0.0143 |

*Note:* M1, M2 and M3 were fitted by maximum likelihood with mgcv 1.9.3 (R 4.4.0) on identical rows (all scans of the reference participants with complete covariates; 14 regions × 2 knees), each with a participant random intercept s(ID, bs = “re”). AIC is the mgcv AIC; GCV = n·RSS/(n − edf)², with RSS from the fitted values including the participant effect and edf the total effective degrees of freedom. M1 also had the highest maximum-likelihood score (marginal likelihood) of the three models in every region and knee, as expected because the linear age–BMI structure of M2 lies within the unpenalized space of the tensor-product surface of M1.

### S3. Local centiles and connected-deficit features

**Residuals.** For participant $i$, visit $j$ and pixel $v$, population residuals are $r_{ij}^{P}\left( v \right)=y_{ij}\left( v \right)-\hat{\mu}_{v}\left( x_{ij} \right)$. Given $m>0$ earlier scans with mean residual $\overline{r}_{i,<j}^{P}\left( v \right)$, the predicted participant effect and the longitudinal (history) residual are

$$\hat{b}_{i,<j}\left( v \right)=\frac{m\hat{\tau}_{v}^{2}}{\hat{\sigma}_{v}^{2}+m\hat{\tau}_{v}^{2}} \overline{r}_{i,<j}^{P}\left( v \right), r_{ij}^{H}\left( v \right)=r_{ij}^{P}\left( v \right)-\hat{b}_{i,<j}\left( v \right).$$

The posterior variance $\tau_{v}^{2}\sigma_{v}^{2}/\left( \sigma_{v}^{2}+m\tau_{v}^{2} \right)$ decreases as more earlier scans become available. The current scan is excluded from this estimate. At the first visit, the longitudinal and cross-sectional residuals are identical.

**Derivation of the longitudinal adjustment.** Under the reference model (S2), the population residual of knee $i$ at pixel $v$ and visit $k$ is $r_{ik}=b_{i}+\varepsilon_{ik}$, with $b_{i}\sim N\left( 0,\tau^{2} \right)$ and $\varepsilon_{ik}\sim N\left( 0,\sigma^{2} \right)$ independent across visits (pixel index dropped). Given $m$ earlier residuals, $\left( b_{i},r_{i1},\ldots,r_{im} \right)$ is jointly Gaussian with $\mathrm{Cov}\left( b_{i},r_{ik} \right)=\tau^{2}$, $\mathrm{Var}\left( r_{ik} \right)=\tau^{2}+\sigma^{2}$ and $\mathrm{Cov}\left( r_{ik},r_{il} \right)=\tau^{2}$ for $k\neq l$. Because the errors are exchangeable, the mean $\overline{r}_{i}$ is sufficient for $b_{i}$, with $\mathrm{Cov}\left( b_{i},\overline{r}_{i} \right)=\tau^{2}$ and $\mathrm{Var}\left( \overline{r}_{i} \right)=\tau^{2}+\sigma^{2}/m$. Gaussian conditioning then gives

$$E\left( b_{i}\mid r_{i1},\ldots,r_{im} \right)=\frac{\tau^{2}}{\tau^{2}+\sigma^{2}/m} \overline{r}_{i}=\frac{m\tau^{2}}{\sigma^{2}+m\tau^{2}} \overline{r}_{i}, \mathrm{Var}\left( b_{i}\mid r_{i1},\ldots,r_{im} \right)=\frac{\tau^{2}\sigma^{2}}{\sigma^{2}+m\tau^{2}}.$$

The conditional mean is the best linear unbiased predictor (BLUP) of the random effect (Henderson, 1975; Robinson, 1991). Substituting maximum-likelihood estimates of the variance components gives the empirical-Bayes posterior mean. Expected thickness conditional on earlier scans is $\hat{\mu}_{v}\left( x_{ij} \right)+\hat{b}_{i,<j}\left( v \right)$. The history residual $r^{H}=r^{P}-\hat{b}$ therefore measures deviation from this individual expectation and has conditional distribution $N\{0,\sigma^{2}+\tau^{2}\sigma^{2}/\left( \sigma^{2}+m\tau^{2} \right)\}$. The weight $m\tau^{2}/\left( \sigma^{2}+m\tau^{2} \right)$ determines how strongly earlier residuals adjust the population prediction. It is zero when no earlier scan is available ($m=0$) and approaches 1 as $m$ increases. Greater between-knee variation $\tau^{2}$ increases the weight, whereas greater residual variation $\sigma^{2}$ reduces it. This construction follows the principle of conditional growth charts (Wei and He, 2006). The main-text notation is $\sigma_{1,v}^{2}=\tau_{v}^{2}$, $\sigma_{2,v}^{2}=\sigma_{v}^{2}$ and $N=m$.

**Assumptions.** Pixelwise variance components are treated as fixed after estimation, so the conditional variance excludes their estimation uncertainty. Earlier residuals use the same fitted population model and are assumed independent across visits conditional on $b_{i}$; serial correlation would reduce the effective number of earlier scans below $m$. Longitudinal centiles rank $r^{H}$ against the empirical null residuals in Algorithm 1 (Table S4). They do not standardize $r^{H}$ by its $m$-specific predictive standard deviation, so all history lengths share the same empirical ranking scale.

**Wild-bootstrap empirical local ranks (Algorithm 1).** We used a participant-level wild bootstrap to construct the empirical reference distribution for local centile ranks, adapting the residual-multiplier approach of Zhu et al. (2007). Independent standard-normal multipliers were shared across each participant’s scans, while the fitted normative models and variance components were held fixed.

For channel $d\in\{P,H\}$ and compartment $c$ (femur or tibia), target scans $q\mathcal{\in T}$ are ranked by their target residuals $r_{q}^{d}\left( v \right)$ (the population residual $r^{P}$ or the history residual $r^{H}$ defined above), against null residuals $e_{q}^{dc}\left( v \right)$ of the null scans $q\in\mathcal{R}_{dc}$. The pooled multiset is $\mathcal{P}_{dcv}=\{r_{q}^{d}\left( v \right):q\mathcal{\in T\}⊎\{}W_{i\left( q \right),b} e_{q}^{dc}\left( v \right):q\in\mathcal{R}_{dc}, b=1,\ldots,B_{dc}\}$, with standard-normal multipliers $W$ shared across a participant’s scans. The null residuals are: for the cross-sectional channel of both compartments, the population residuals of the reference training scans, $e_{q}^{Pc}=r_{q}^{P}$; for the femoral longitudinal channel, the conditional residuals of the reference training scans after subtracting the fitted participant random effect, $e_{q}^{H,fem}=r_{q}^{P}-\hat{\alpha}_{i\left( q \right)}$, where $\hat{\alpha}_{i\left( q \right)}$ is estimated from all of that participant’s reference scans (unlike the target history residual $r^{H}$, which subtracts an effect predicted from earlier scans only); and for the tibial longitudinal channel, the history residuals of the scored scans read KLG 0 at follow-up visits, $e_{q}^{H,tib}=r_{q}^{H}$. The null scans and numbers of multipliers are listed in Table S4. At a participant’s earliest scored visit the longitudinal centile is set to the cross-sectional centile. The local centile of a target residual $z=r_{q}^{d}\left( v \right)$ is its mid-rank in the pooled multiset,

$$C_{dc}\left( z;v \right)=\frac{\sum_{p\in\mathcal{P}_{dcv}} \mathbf{1}\left( p<z \right)+\{1+\sum_{p\in\mathcal{P}_{dcv}} \mathbf{1}\left( p=z \right)\}/2}{\left| \mathcal{P}_{dcv} \right|}.$$

Low centiles indicate thinner-than-expected cartilage and high centiles thicker-than-expected cartilage. The multipliers enlarge the empirical reference pool but do not increase the number of independent participants. The resulting centiles describe relative thickness; they are not calibrated P values.

**Table S4.** Null pools of the local centiles by compartment and channel (Algorithm 1).

| Compartment | Channel | Null residuals $e_{q}^{dc}$, $q\in\mathcal{R}_{dc}$ | Multipliers $B_{dc}$ |
| --- | --- | --- | --- |
| Femur | Cross-sectional (P) | Population residuals of the reference (training) scans | 100 |
| Tibia | Cross-sectional (P) | Population residuals of the reference (training) scans | 100 |
| Femur | Longitudinal (H) | Conditional residuals of the reference training scans, after subtracting the fitted participant random effect | 100 |
| Tibia | Longitudinal (H) | History residuals $r^{H}$ (earlier scans only) of the scored scans read KLG 0 at follow-up (non-baseline) visits | 1,000 |

*Note:* the same specification was used for the full-resolution full-reference charts of the main analysis and for the downsampled charts of S9; reference (training) maps were ranked against their own residuals × 100 multipliers.

**Connected deficit extent (Algorithm 2).** For region $r$ with valid pixels $V_{r}$, we defined the pixels below threshold u as $E_{r}\left( u \right)=\{v\in V_{r}:C\left( v \right)<u\}$. Using eight-neighbour connectivity, $K_{r}\left( u \right)$ counts pixels in the largest connected component of $E_{r}\left( u \right)$. We used five thresholds, $u_{k}=0.1exp\{-\left( k-1 \right)\log\left( 1000 \right)/19\}$ (0.100, 0.070, 0.048, 0.034 and 0.023). Each extent was ranked against reference extents as $Q\left( a \right)=\{1+\sum_{\mathcal{l}} \mathbf{1}\left( a_{\mathcal{l}}^{0}<a \right)+1/2\sum_{\mathcal{l}} \mathbf{1}\left( a_{\mathcal{l}}^{0}=a \right)\}/\left( n_{0}+1 \right)$. A larger $Q$ indicates a larger connected deficit relative to reference knees.

**Regional centile features.** We combined the five extent ranks and the five extents separately using normalized first-principal-component loadings $w=v/\mathbf{1}^{\top}v$ (all positive). This gave the global centile score $G=w_{Q}^{\top}q$ and weighted extent $D=w_{A}^{\top}a$ for each region. We then split the logit $L\left( G \right)=log\{\left( G+\varepsilon\right)/\left( 1-G+\varepsilon\right)\}$ into positive and negative parts, $\mathrm{GCS}^{+}=max\{L,0\}$ and $\mathrm{GCS}^{-}=min\{L,0\}$, with $\varepsilon={10}^{-6}$. Across 14 regions and two centile types, this produced 56 logit features and 28 extents. Prognostic models also included the maximum relative deficit area in each compartment, giving 86 features. Negative GCS− values indicate deficit ranks below the reference median, not cartilage thickening. Compression weights were estimated from all 21,369 scans before splitting by outcome.

**Table S5.** Global centile features and accompanying predictors: identifiers, definitions and interpretation. This dictionary covers every predictor of feature set A+P. Global centile scores (GCS) summarize connected low-centile deficits within each region or whole cartilage plate. The supplementary SHAP atlas retains P/H labels; revised main Figure 6 uses the equivalent X/L labels. Identifiers below follow Figures SH1–SH24; [region] is a region code of main Figure 1D (e.g. 8 LCeT) or Femoral/Tibial for a whole plate. P = cross-sectional (population, fixed-effect) channel; H = longitudinal (prior-scan, random-effect) channel.

| Block (n) | Identifier | Definition | Direction: a higher value means |
| --- | --- | --- | --- |
| Clinical (11; 5 for concurrent classification) | Age, Male sex, Black race, BMI, Height, Alignment, Injury history, Physical activity, Quadriceps strength, Laxity, Index KLG | Baseline covariates (units and coding in Table S2); index KLG 0 or 1 (prognosis only) | A larger covariate value (male sex, Black race and index KLG 1 coded 1) |
| ROI mean thickness (14) | [region] thickness | Mean cartilage thickness (mm) of the region, not adjusted for covariates | Thicker cartilage in absolute terms |
| Cross-sectional GCS+ (14) | [region] · P GCS+ | max{logit *G*, 0}, *G* = combined rank of the connected low-centile extents of the region among reference knees, from cross-sectional centiles | A larger connected thinner-than-expected area than the reference median, relative to matched reference knees (age, BMI, sex, race, height); zero for *G* ≤ 0.5 |
| Cross-sectional GCS− (14) | [region] · P GCS− | min{logit *G*, 0}, same *G* | A deficit rank closer to the reference median from below; the value reaches zero at *G* = 0.5 and remains zero for larger deficits |
| Cross-sectional extent (14) | [region] · P extent | PCA-weighted combination of the five largest connected low-centile extents (thresholds 0.100–0.023) | A larger connected thinner-than-expected area |
| Longitudinal GCS+ (14) | [region] · H GCS+ | As P GCS+, from longitudinal centiles (residual relative to the thickness predicted from the knee’s earlier scans) | A larger connected area of cartilage thinner than predicted from earlier scans; zero for *G* ≤ 0.5 |
| Longitudinal GCS− (14) | [region] · H GCS− | As P GCS−, from longitudinal centiles | A deficit rank closer to the reference median from below, using longitudinal centiles; zero for *G* ≥ 0.5 |
| Longitudinal extent (14) | [region] · H extent | As P extent, from longitudinal centiles | A larger connected area thinner than predicted from the knee’s own history |
| Maximum extent (2; prognosis only) | Femoral maximum extent, Tibial maximum extent | Largest connected low-centile extent of the compartment over both channels and the five thresholds, divided by the number of compartment pixels | A larger fraction of the compartment in one connected thinner-than-expected area |

*Note:* The 86 features summarize connected areas of thinner-than-expected cartilage. Cross-sectional features (P) use covariate-adjusted population expectations; longitudinal features (H) also use the knee’s earlier scans. Low feature values indicate little connected deficit and do not measure thickening. High local centiles retain information about thicker-than-expected cartilage in the continuous maps (main Figure 2A2) used for deep features (F2). At the first scored visit, the H map is set equal to the P map. Regional profiles can still differ because extent ranking and compression are performed separately for each centile type. A positive SHAP value accompanying a high feature value (main Figure 6) indicates that greater connected deficit burden contributed to a higher predicted probability of incident OA.

**Spatial extent and sensitivity to centile changes.** Extent depends on how low-centile pixels are connected. Four pixels below the threshold give an extent of 4 when connected and 1 when isolated, despite having the same centile histogram.

Lowering centiles cannot reduce the largest connected deficit: if $C'\leq C$ pointwise, then $K_{C}\left( u \right)\leq K_{C'}\left( u \right)$. For two maps that satisfy $\parallel C-C'\parallel_{\infty}\leq\delta$, the extents satisfy $K_{C}\left( u-\delta\right)\leq K_{C'}\left( u \right)\leq K_{C}\left( u+\delta\right)$. The integrated difference is bounded by $\int\left| K_{C}\left( u \right)-K_{C'}\left( u \right) \right|du\leq2\delta\left| V_{r} \right|$.

**Continuous centile maps for deep features.** Deep features were extracted directly from the femoral and tibial rank maps $C_{P}$ and $C_{H}$ (four images per scan). These maps retain both thinner-than-expected and thicker-than-expected cartilage.

### S4. Comparator representations

**Table S6.** Prognostic feature blocks and feature sets. X and L denote cross-sectional and longitudinal centiles. P denotes regional profiles and F map deep features. A is the shared clinical and regional-thickness baseline. Counts for complete feature sets include A. The main-analysis blocks use full-resolution full-reference charts; sensitivity variants are defined below. The population/history labels P/H in centile equations and the SHAP atlas correspond to X/L.

| Block | Content | Features |
| --- | --- | --- |
| Clinical | Sex, race, BMI, height, alignment, injury history, physical activity, quadriceps strength, laxity, age, index KLG | 11 |
| ROI mean thickness | Mean thickness of 12 ROIs and two plates | 14 |
| ROI PCA | First five principal components of the raw thickness map per ROI (fitted in each outer-training fold) | 60 |
| ROI radiomics | 93 PyRadiomics features per ROI | 1,116 |
| Cross-sectional profiles (PX) | Three regional summaries in each of 14 regions, from cross-sectional centiles | 42 |
| Longitudinal profiles (PL) | Three regional summaries in each of 14 regions, from longitudinal centiles | 42 |
| Combined centile profiles (P = P0) | PX + PL + femoral and tibial maximum-area measures (S3), full-resolution full-reference charts | 86 |
| Raw-map deep features (F1) | 11 ImageNet CNNs × 2 raw thickness maps × 10 PCs | 220 |
| Cross-sectional centile-map deep features (F2X) | 11 networks × 2 cross-sectional maps (femur/tibia) × 10 PCs | 220 |
| Longitudinal centile-map deep features (F2L) | 11 networks × 2 longitudinal maps (femur/tibia) × 10 PCs | 220 |
| Combined centile-map deep features (F2 = F2X + F2L) | 11 ImageNet CNNs × 4 local centile maps (fixed-effect cross-sectional and random-effect longitudinal, × femur/tibia) × 10 PCs, full-resolution full-reference charts; twice the size of F1 because each thickness map yields two centile maps | 440 |

| Symbol | Feature set | Blocks | Features |
| --- | --- | --- | --- |
| A | Clinical + ROI mean thickness (baseline) | Clinical + ROI mean thickness | 25 |
| A+PX | Baseline + cross-sectional profiles | A + PX | 67 |
| A+PL | Baseline + longitudinal profiles | A + PL | 67 |
| G2X | Baseline + cross-sectional map deep features | A + F2X | 245 |
| G2L | Baseline + longitudinal map deep features | A + F2L | 245 |
| B | Baseline + ROI PCA | A + ROI PCA | 85 |
| C | Baseline + ROI radiomics | A + ROI radiomics | 1,141 |
| D | Baseline + ROI PCA + ROI radiomics | A + ROI PCA + ROI radiomics | 1,201 |
| A+P | Baseline + combined centile profiles | A + P | 111 |
| E | Baseline + ROI PCA + ROI radiomics + combined centile profiles | D + P | 1,287 |
| G1 | Baseline + raw-map deep features | A + F1 | 245 |
| G2 | Baseline + centile-map deep features | A + F2 | 465 |
| G12 | Baseline + raw-map and centile-map deep features | A + F1 + F2 | 685 |
| H1 | Baseline + raw-map deep features + centile profiles | G1 + P | 331 |
| H2 | Baseline + centile-map deep features + centile profiles | G2 + P | 551 |
| H12 | Baseline + both deep feature sets + centile profiles | G12 + P | 771 |

**P0 and E0 in the supplementary data.** P0 is the combined profile block P: 42 cross-sectional profiles, 42 longitudinal profiles and two compartmental maximum-area measures, derived from full-resolution full-reference charts. The suffix 0 identifies this chart variant. A+P0 adds these 86 profiles to the 25 clinical and regional-thickness predictors (111 total). E0 adds the profiles to the clinical/thickness baseline, 60 regional PCA scores and 1,116 radiomic features (25 + 60 + 1,116 + 86 = 1,287 predictors). The main text and figures denote these blocks and sets as P, A+P and E, respectively. Sensitivity suffixes 1 and 2 are defined below.

**Concurrent classification (Table S11).** These models used five clinical predictors (sex, race, BMI, height and age), 14 regional thickness means and 84 centile profiles. Alignment, injury history, physical activity, quadriceps strength, laxity and index KLG were used only for prognosis, as were the two maximum-area features. For concurrent classification, A therefore contained 19 predictors and A+P contained 103, compared with 25 and 111 for prognosis.

**Sensitivity profile blocks (S9).** P1 contains 86 combined cross-sectional and longitudinal profiles recomputed from downsampled cross-fitted charts (method 1; the crossfit_avg scoring variant). P2 contains the same 86 profile definitions recomputed from downsampled full-reference charts (method 2). Downsampling averages each 3 × 3 pixel block. Method 1 scores each reference participant with charts fitted on the other reference half; method 2 fits the charts to the full reference. Thus, A+P1 = A + P1 and A+P2 = A + P2 (111 predictors each); E1 = D + P1 and E2 = D + P2 (1,287 each). The suffix identifies reference construction; all three profile blocks P0, P1 and P2 combine both centile types. Section S9 also reports a separate single-chart cross-fitted variant.

**Sensitivity deep-feature sets.** The suffixes m1 and m2 identify the same downsampled cross-fitted and downsampled full-reference charts, respectively. G2m1 and G2m2 combine A with 440 deep features from the two centile types (465 predictors). G12m1 and G12m2 additionally contain the 220 raw-map deep features (685). H2m1 = G2m1 + P1 and H2m2 = G2m2 + P2 (551). H12m1 = G12m1 + P1 and H12m2 = G12m2 + P2 (771). Unsuffixed G2, G12, H2 and H12 use full-resolution full-reference charts. The feature-set numeral in G1/G2 identifies raw-map/centile-map inputs; the appended m1/m2 identifies chart construction.

**ROI mean thickness (14).** Mean thickness in the 12 ROIs and the two plates.

**ROI PCA (60).** We represented each full-resolution right-knee thickness map as a vector of thickness values in millimetres on the common two-dimensional atlas (38,009 femoral and 36,715 tibial pixels). The 12 radiologist-defined ROI masks (main Figure 1D) were intersected with the compartment masks. Pixels within ROI $r$ formed a scan-by-pixel matrix $X_{r}$. Within each outer-training fold, we centred $X_{r}$ by the training mean and used singular value decomposition to obtain five principal components without variance scaling. Component signs were chosen so that loadings summed to a positive value. Training and held-out scans were projected onto these components, yielding 60 scores (5 × 12). Held-out scans contributed to neither the means nor the components. Inner tuning and SVM calibration used the components estimated from the corresponding outer-training fold. Scores were median-imputed, standardized and weighted using training-fold statistics.

**ROI radiomics (1,116).** Each thickness map was rendered as a 310 × 310 image with intensities in millimetres, pixel spacing of 1 atlas unit and zeros outside the compartment. We applied the same 12 ROI masks used for PCA. PyRadiomics 3.1.0 (Python; van Griethuysen et al., 2017) extracted two-dimensional features (force2D) from the original image, without filtering, resampling or intensity normalization. Intensity bins had a fixed width of 0.1 mm of thickness; this value does not specify anatomical pixel spacing. PyRadiomics implements IBSI definitions with documented deviations, including fixed-width discretization and the kurtosis convention (Zwanenburg et al., 2020). Full IBSI compliance was not verified with a phantom. We extracted 93 features per ROI: 18 first-order statistics, 24 grey-level co-occurrence matrix features, 16 grey-level run-length matrix features, 16 grey-level size-zone matrix features, 14 grey-level dependence matrix features and five neighbouring grey-tone difference matrix features. Shape features were excluded because ROI masks were identical across scans. The resulting 1,116 features were extracted independently for each scan without outcome information, then median-imputed, standardized and weighted within training folds.

**Deep transfer-learning features.** Following Kirchler et al. (2022) and Zhao et al. (2024), we rendered each map at 448 × 448 pixels, set the masked background to zero, applied per-image min–max scaling, replicated the image across three channels and used ImageNet normalization. Each image was passed through 11 ImageNet-pretrained networks from torchvision 0.22.1 (PyTorch 2.7.1; Table S7). Activations from the last convolutional block were spatially averaged, then averaged across eight augmented views (four 90° rotations with and without horizontal flipping). Within each outer-training fold, we retained ten principal components per network and image type. Raw-map features comprised 220 scores (F1: 11 networks × 2 maps × 10 components). The combined centile representation comprised 440 scores (F2: 11 × 4 × 10), using cross-sectional and longitudinal maps for each compartment (S3).

**Table S7.** Pretrained networks used for deep transfer-learning features (torchvision 0.22.1 IMAGENET1K_V1 weights; PyTorch 2.7.1).

| Network | Weights | ImageNet top-1 (%) |
| --- | --- | --- |
| resnet50 | ResNet50_Weights.IMAGENET1K_V1 | 76.1 |
| alexnet | AlexNet_Weights.IMAGENET1K_V1 | 56.5 |
| vgg16 | VGG16_Weights.IMAGENET1K_V1 | 71.6 |
| vgg19 | VGG19_Weights.IMAGENET1K_V1 | 72.4 |
| googlenet | GoogLeNet_Weights.IMAGENET1K_V1 | 69.8 |
| inception_v3 | Inception_V3_Weights.IMAGENET1K_V1 | 77.3 |
| resnet18 | ResNet18_Weights.IMAGENET1K_V1 | 69.8 |
| resnet34 | ResNet34_Weights.IMAGENET1K_V1 | 73.3 |
| squeezenet1_0 | SqueezeNet1_0_Weights.IMAGENET1K_V1 | 58.1 |
| mobilenet_v2 | MobileNet_V2_Weights.IMAGENET1K_V1 | 71.9 |
| shufflenet_v2_x1_0 | ShuffleNet_V2_X1_0_Weights.IMAGENET1K_V1 | 69.4 |

### S5. Classifiers, nested cross-validation and tuning

All 12 classifiers used the same cohorts, folds and random seeds, with ten outer and three inner folds separated by participant. Hyperparameters minimized the pooled inner out-of-fold Brier score; ties were resolved in favour of greater regularization. The selected model was refitted on the full outer-training fold. Missing values were imputed with training medians, and predictors were standardized with training means and standard deviations. Clinical predictors were multiplied by $\sqrt{10}$, and each imaging block was divided by the square root of its feature count to balance its contribution to penalized models. PCA for regional and deep features was fitted within each outer-training fold. Linear SVM probabilities were calibrated by Platt scaling on additional participant-separated training folds.

**Table S8.** The 12 classifiers: abbreviation used in figures and tables, full name, implementation, tuned hyperparameters (grid searched by inner cross-validation) and key references.

| Abbreviation | Full name | Implementation | Tuned hyperparameters (grid) | References |
| --- | --- | --- | --- | --- |
| Ridge | Ridge (L2-penalized) logistic regression | scikit-learn 1.5.1 LogisticRegression (Python), L-BFGS | penalty λ = 10⁻⁵ … 10⁻¹ | Hoerl and Kennard (1970); le Cessie and van Houwelingen (1992); Pedregosa et al. (2011) |
| LASSO | Least absolute shrinkage and selection operator (L1-penalized) logistic regression | glmnet::glmnet (glmnet 4.1.8, R 4.4.0) | λ = 10⁻¹, 10⁻², 10⁻³, 10⁻⁴, 10⁻⁵ | Tibshirani (1996); Friedman et al. (2010) |
| Bayes logit | Bayesian logistic regression with weakly informative Cauchy priors | arm::bayesglm (arm 1.14.4, R 4.4.0) | prior scale 2.5 (fixed) | Gelman et al. (2008) |
| Lin. SVM | Linear support vector machine with Platt-calibrated probabilities | kernlab::ksvm (kernlab 0.9.33, R 4.4.0; C-svc, linear kernel); calibration on participant-separated folds | C = 0.001, 0.01, 0.1, 1, 10 | Cortes and Vapnik (1995); Platt (1999); Karatzoglou et al. (2004) |
| LDA | Linear discriminant analysis | MASS::lda (MASS 7.3.65, R 4.4.0) | one discriminant (no tuning) | Fisher (1936); Venables and Ripley (2002) |
| GLM boost | Component-wise gradient boosting of a logistic generalized linear model | mboost::glmboost (mboost 2.9.10, R 4.4.0; step size 0.1) | boosting iterations 50, 100, 200, 400, 800 | Bühlmann and Hothorn (2007) |
| XGBoost | Extreme gradient boosting of decision trees | xgboost::xgb.train (xgboost 1.7.7.1, R 4.4.0) | depth 1/3/6; learning rate 0.03/0.1; 100/300 rounds | Chen and Guestrin (2016) |
| LightGBM | Light gradient boosting machine | lightgbm::lgb.train (lightgbm 4.4.0, R 4.4.0) | leaves 7/15/31; learning rate 0.03/0.1; 100/300 rounds | Ke et al. (2017) |
| Rand. forest | Random forest | randomForest::randomForest (randomForest 4.7.1.1, R 4.4.0; 500 trees) | mtry ⌊√p⌋, ⌊p/3⌋, p; node size 1/5/10 | Breiman (2001) |
| Neural net | Single-hidden-layer feed-forward neural network (cross-entropy loss) | nnet::nnet (nnet 7.3.20, R 4.4.0) | hidden units 3/5/10; weight decay 0.001/0.01/0.1 | Ripley (1996); Venables and Ripley (2002) |
| Naïve Bayes | Naïve Bayes classifier with Gaussian or kernel class-conditional densities | naivebayes::naive_bayes (naivebayes 1.0.0, R 4.4.0) | Gaussian, or kernel density with bandwidth adjustment 0.5/1/2 | Hand and Yu (2001) |
| HDDA | High-dimensional discriminant analysis | HDclassif::hdda (HDclassif 2.2.2, R 4.4.0; all covariance models, chosen by BIC) | intrinsic-dimension threshold 0.05/0.175/0.3 | Bouveyron et al. (2007); Bergé et al. (2012) |

**Additional settings.** Tree boosting used 0.8 row/column subsampling and L2 penalty 1. Non-converged candidates were excluded from inner selection; when a selected outer fit failed to converge, the next-best inner candidate was used, and Bayesian logistic regression was refitted with up to 10,000 iterations where needed. Fits used R 4.4.0 and scikit-learn 1.5.1.

### S6. Performance metrics and statistical inference

Threshold-dependent metrics used a probability cut-off of 0.50. Case precision was TP/(TP+FP), case recall TP/(TP+FN), control precision TN/(TN+FN) and control recall TN/(TN+FP). Control precision and recall correspond to negative predictive value and specificity. Case and control F1 scores were 2TP/(2TP+FP+FN) and 2TN/(2TN+FP+FN). Balanced accuracy was the mean of the two recalls, and macro-F1 was (F1_case_ + F1_control_)/2. Metrics were calculated within each of the ten analysis datasets and then averaged. Precision and F1 depend on the sampled case–control ratio (S1).

We estimated uncertainty with 2,000 paired participant-bootstrap resamples. Each participant received one multinomial weight, shared across datasets, feature sets and classifiers to preserve pairing. We report 95% percentile intervals and two-sided P values, calculated as $2min\{Pr\left( \Delta^{*}\leq0 \right),Pr\left( \Delta^{*}\geq0 \right)\}$. Holm adjustment covered the 12 classifiers within each comparison, horizon and metric. Data S2 also reports global Holm adjustments across comparisons, classifiers and horizons for each metric, separately for the PCA/radiomics/centile-profile, deep-feature and F2-check analyses. These are separate multiplicity families rather than one manuscript-wide adjustment. Table S9 specifies the resampling method and multiplicity family for each analysis. Intervals and tests condition on the fitted charts and feature construction; they exclude uncertainty from refitting those steps.

**Table S9.** Statistical inference and multiplicity adjustments. The table identifies the result source, interval or test, multiplicity family and role of each analysis. Unless stated otherwise, intervals and tests use 2,000 paired participant-bootstrap resamples, and classifier families contain 12 classifiers. Primary inference concerns the 12- and 24-month feature-set comparisons in main Figure 5. Other tests are classified as secondary, sensitivity or exploratory analyses.

| Table / figure | Source | Interval / test | Multiplicity family (size) | Role |
| --- | --- | --- | --- | --- |
| Main Table 2; main Figure 3; Tables S12–S15; Data S1 | Nested-CV predictions of all feature sets × 12 classifiers × 10 analysis datasets | 95% percentile bootstrap intervals; no test | — | Primary estimates (descriptive) |
| Main Figure 4; main Tables 2–3 (channel results); Table S23; Figures C1–C3; Data S5 | Same index participants/folds; separate X/L profile and deep-feature predictions | 95% paired bootstrap intervals; two-sided bootstrap P for differences | Holm across 12 classifiers per contrast × horizon × metric; channel-suite global Holm: 12 contrasts × 12 classifiers × 3 horizons = 432 per metric | Additional channel ablation, not prospectively specified; unequal prior-scan availability remains |
| Main Figure 5; Tables S16–S18; Data S2 | Same predictions, paired differences between feature sets | Two-sided bootstrap P | Holm across classifiers within comparison × horizon × metric (12); global Holm within suite per metric: PCA/radiomics/centile profiles 23 comparisons × 12 × 3 horizons (828), deep features 15 × 12 × 3 (540) | Primary: 12- and 24-month contrasts of main Figure 5; other contrasts and 36 months secondary |
| Table S10 | Ridge rows of the same run | 95% bootstrap intervals; no test | — | Descriptive |
| Table S11; Figure A9 | Concurrent-classification analysis (five participant-grouped partitions) and within-classifier centile increments | Paired delete-one-participant jackknife P; 95% intervals from 10,000 participant resamples | Table S11: 96-test primary ridge family (4 outcomes × 2 comparators × 12 metrics); Figure A9: 1,152 tests (classifier × outcome × metric) | Secondary (concurrent) / exploratory (Figure A9) |
| Tables S19–S20; Data S3 (reference arms) | Downsampled-chart refits (S9) | Two-sided bootstrap P | Holm across classifiers within type × comparison × horizon × metric (12); no global family | Sensitivity |
| Tables S21–S22; Data S3 (F2 check) | Centile-map deep features from downsampled charts (S9) | Two-sided bootstrap P | Holm across classifiers (12); global Holm within the F2-check suite per metric (540) | Sensitivity |
| Table S2 | Case and control characteristics | Welch t, χ² or Fisher exact test | Unadjusted | Descriptive |
| Table S3 | Regional reference models M1–M3 | AIC and GCV; no test | — | Model comparison (descriptive) |
| Main Figure 6; Figures SH1–SH24; Data S4 | SHAP of the five highest-AUC classifiers | Monte Carlo stability check; no test | — | Exploratory interpretation |
| Figures A10–A17 | Descriptive longitudinal and regional analyses | Pointwise 95% intervals; Holm P where shown | Holm within the family stated in each caption (e.g. Figure A17: 126 means, 56 contrasts) | Exploratory |

### S7. Complete performance results

**Table S10.** Ridge logistic regression, raw-map comparator benchmark (all metrics; estimate and 95% CI).

*12-month prognosis*

| Metric | A | B | C | D | A+P | E |
| --- | --- | --- | --- | --- | --- | --- |
| AUC | 0.897 (0.866–0.926) | 0.905 (0.875–0.932) | 0.922 (0.896–0.944) | 0.920 (0.893–0.944) | 0.942 (0.918–0.960) | 0.935 (0.910–0.956) |
| Brier | 0.116 (0.098–0.134) | 0.112 (0.094–0.130) | 0.104 (0.087–0.123) | 0.103 (0.087–0.122) | 0.089 (0.072–0.108) | 0.094 (0.077–0.112) |
| Accuracy | 0.817 (0.783–0.849) | 0.826 (0.793–0.859) | 0.842 (0.811–0.870) | 0.841 (0.809–0.871) | 0.874 (0.845–0.899) | 0.861 (0.832–0.890) |
| Balanced accuracy | 0.803 (0.776–0.829) | 0.815 (0.788–0.840) | 0.829 (0.803–0.851) | 0.830 (0.804–0.854) | 0.863 (0.841–0.884) | 0.851 (0.827–0.874) |
| Precision (case) | 0.893 (0.873–0.911) | 0.901 (0.881–0.918) | 0.907 (0.889–0.924) | 0.909 (0.890–0.925) | 0.926 (0.911–0.940) | 0.920 (0.904–0.936) |
| Recall (case) | 0.839 (0.791–0.883) | 0.844 (0.797–0.888) | 0.862 (0.820–0.901) | 0.859 (0.816–0.901) | 0.891 (0.852–0.925) | 0.878 (0.835–0.915) |
| F1 (case) | 0.865 (0.835–0.891) | 0.872 (0.842–0.898) | 0.884 (0.857–0.906) | 0.883 (0.856–0.908) | 0.908 (0.884–0.928) | 0.898 (0.873–0.921) |
| Precision (control) | 0.673 (0.602–0.746) | 0.685 (0.617–0.756) | 0.713 (0.645–0.782) | 0.711 (0.643–0.781) | 0.768 (0.702–0.829) | 0.744 (0.678–0.811) |
| Recall (control) | 0.767 (0.739–0.796) | 0.785 (0.757–0.813) | 0.795 (0.767–0.824) | 0.800 (0.772–0.827) | 0.836 (0.811–0.861) | 0.824 (0.798–0.849) |
| F1 (control) | 0.717 (0.671–0.760) | 0.732 (0.686–0.774) | 0.751 (0.710–0.790) | 0.753 (0.710–0.793) | 0.800 (0.760–0.835) | 0.782 (0.741–0.821) |
| Macro-F1 | 0.791 (0.756–0.823) | 0.802 (0.767–0.833) | 0.818 (0.786–0.846) | 0.818 (0.786–0.849) | 0.854 (0.824–0.880) | 0.840 (0.809–0.869) |

*24-month prognosis*

| Metric | A | B | C | D | A+P | E |
| --- | --- | --- | --- | --- | --- | --- |
| AUC | 0.886 (0.844–0.926) | 0.888 (0.847–0.927) | 0.887 (0.845–0.926) | 0.888 (0.847–0.928) | 0.919 (0.882–0.950) | 0.907 (0.868–0.941) |
| Brier | 0.124 (0.101–0.148) | 0.123 (0.100–0.148) | 0.123 (0.099–0.149) | 0.123 (0.099–0.147) | 0.105 (0.081–0.132) | 0.114 (0.090–0.140) |
| Accuracy | 0.819 (0.777–0.858) | 0.823 (0.781–0.862) | 0.823 (0.782–0.862) | 0.825 (0.784–0.865) | 0.859 (0.822–0.892) | 0.839 (0.800–0.876) |
| Balanced accuracy | 0.825 (0.788–0.859) | 0.828 (0.793–0.862) | 0.830 (0.794–0.864) | 0.832 (0.796–0.865) | 0.862 (0.830–0.891) | 0.845 (0.810–0.877) |
| Precision (case) | 0.891 (0.862–0.915) | 0.892 (0.864–0.917) | 0.896 (0.868–0.920) | 0.897 (0.869–0.920) | 0.910 (0.885–0.932) | 0.906 (0.881–0.928) |
| Recall (case) | 0.794 (0.727–0.858) | 0.800 (0.735–0.864) | 0.797 (0.731–0.860) | 0.800 (0.735–0.863) | 0.847 (0.789–0.901) | 0.815 (0.752–0.875) |
| F1 (case) | 0.839 (0.795–0.880) | 0.844 (0.800–0.883) | 0.843 (0.798–0.883) | 0.846 (0.802–0.885) | 0.878 (0.840–0.909) | 0.858 (0.816–0.896) |
| Precision (control) | 0.737 (0.668–0.809) | 0.743 (0.674–0.814) | 0.741 (0.672–0.813) | 0.744 (0.675–0.816) | 0.795 (0.728–0.859) | 0.761 (0.694–0.830) |
| Recall (control) | 0.855 (0.832–0.879) | 0.857 (0.832–0.880) | 0.863 (0.839–0.885) | 0.863 (0.839–0.886) | 0.876 (0.852–0.899) | 0.875 (0.852–0.897) |
| F1 (control) | 0.792 (0.748–0.834) | 0.796 (0.752–0.837) | 0.797 (0.754–0.839) | 0.799 (0.756–0.840) | 0.833 (0.794–0.869) | 0.814 (0.771–0.853) |
| Macro-F1 | 0.816 (0.774–0.854) | 0.820 (0.778–0.858) | 0.820 (0.779–0.858) | 0.822 (0.781–0.861) | 0.855 (0.819–0.888) | 0.836 (0.797–0.873) |

*36-month prognosis*

| Metric | A | B | C | D | A+P | E |
| --- | --- | --- | --- | --- | --- | --- |
| AUC | 0.871 (0.818–0.916) | 0.870 (0.817–0.917) | 0.871 (0.819–0.916) | 0.871 (0.818–0.917) | 0.869 (0.817–0.914) | 0.871 (0.818–0.916) |
| Brier | 0.136 (0.110–0.167) | 0.137 (0.110–0.167) | 0.136 (0.110–0.166) | 0.136 (0.109–0.166) | 0.138 (0.111–0.169) | 0.136 (0.110–0.167) |
| Accuracy | 0.795 (0.744–0.841) | 0.796 (0.746–0.842) | 0.799 (0.748–0.844) | 0.797 (0.745–0.844) | 0.796 (0.743–0.843) | 0.799 (0.746–0.846) |
| Balanced accuracy | 0.798 (0.750–0.841) | 0.800 (0.751–0.843) | 0.802 (0.755–0.846) | 0.800 (0.752–0.845) | 0.799 (0.750–0.843) | 0.802 (0.754–0.847) |
| Precision (case) | 0.853 (0.810–0.889) | 0.856 (0.814–0.893) | 0.861 (0.819–0.897) | 0.860 (0.818–0.896) | 0.854 (0.810–0.890) | 0.861 (0.820–0.896) |
| Recall (case) | 0.739 (0.648–0.821) | 0.737 (0.646–0.821) | 0.737 (0.645–0.821) | 0.735 (0.640–0.820) | 0.739 (0.645–0.823) | 0.739 (0.644–0.824) |
| F1 (case) | 0.792 (0.725–0.843) | 0.792 (0.726–0.844) | 0.794 (0.727–0.846) | 0.792 (0.724–0.845) | 0.792 (0.723–0.845) | 0.795 (0.728–0.849) |
| Precision (control) | 0.746 (0.669–0.821) | 0.747 (0.668–0.822) | 0.748 (0.669–0.823) | 0.746 (0.666–0.822) | 0.747 (0.667–0.823) | 0.748 (0.668–0.825) |
| Recall (control) | 0.858 (0.830–0.882) | 0.862 (0.834–0.887) | 0.867 (0.841–0.892) | 0.866 (0.840–0.891) | 0.859 (0.832–0.884) | 0.866 (0.839–0.890) |
| F1 (control) | 0.798 (0.749–0.841) | 0.800 (0.750–0.843) | 0.803 (0.751–0.845) | 0.801 (0.750–0.845) | 0.799 (0.748–0.843) | 0.803 (0.752–0.846) |
| Macro-F1 | 0.795 (0.743–0.840) | 0.796 (0.744–0.841) | 0.799 (0.745–0.844) | 0.797 (0.743–0.842) | 0.795 (0.742–0.842) | 0.799 (0.745–0.845) |

*Note:* feature sets defined in Table S6.

**Table S11.** Concurrent classification (KLG ≥2 vs ≤1; 2,249 scans, 1,923 participants; five participant-grouped partitions): ridge AUC and Brier score (95% CI). Feature sets for this task: clinical = sex, race, BMI, height and age (5 predictors); A = clinical + 14 ROI mean thicknesses (19); A+P = A + 84 centile profiles (103; the two maximal-area features are prognosis-only; Table S6).

| Feature set | AUC (95% CI) | Brier (95% CI) |
| --- | --- | --- |
| Clinical | 0.634 (0.607–0.661) | 0.227 (0.220–0.233) |
| Clinical + ROI mean thickness (feature set A) | 0.847 (0.829–0.863) | 0.156 (0.147–0.166) |
| Clinical + ROI mean thickness + centile profiles (feature set A+P) | 0.907 (0.894–0.919) | 0.121 (0.112–0.130) |

*Note:* feature sets defined in Table S6.

Adding centile profiles to A increased concurrent-classification AUC by 0.060 (95% CI 0.048–0.072), balanced accuracy by 0.070 (0.053–0.088) and macro-F1 by 0.068 (0.051–0.085); all Holm-adjusted P values were <0.0001. Concurrent-classification P values were obtained by paired delete-one-participant jackknife tests, with Holm adjustment across the 96-test primary ridge family (4 outcomes × 2 comparators × 12 metrics). Confidence intervals used 10,000 participant resamples. Prognostic comparisons used the paired bootstrap described in S6.

**Table S12.** AUC for each feature set and classifier. Values are averaged across the ten analysis datasets; 95% confidence intervals are provided in Data S1. Bold indicates the highest value within each classifier.

*12-month prognosis*

| Classifier | B | C | D | E | A | G1 | A+P | G2 | G12 | H1 | H2 | H12 |
| --- | --- | --- | --- | --- | --- | --- | --- | --- | --- | --- | --- | --- |
| Ridge | 0.905 | 0.922 | 0.920 | 0.935 | 0.897 | 0.908 | 0.942 | **0.948** | 0.941 | 0.930 | 0.948 | 0.945 |
| LASSO | 0.892 | 0.911 | 0.892 | 0.909 | 0.889 | 0.891 | 0.926 | **0.947** | 0.936 | 0.919 | 0.938 | 0.932 |
| Bayesian logistic | 0.909 | 0.931 | 0.922 | 0.941 | 0.908 | 0.917 | 0.941 | 0.949 | 0.947 | 0.939 | **0.950** | 0.949 |
| Linear SVM | 0.904 | 0.921 | 0.915 | 0.932 | 0.892 | 0.913 | 0.945 | **0.952** | 0.948 | 0.935 | 0.952 | 0.945 |
| LDA | 0.870 | 0.779 | 0.783 | 0.807 | **0.907** | 0.567 | 0.894 | 0.804 | 0.855 | 0.747 | 0.836 | 0.859 |
| GLM boosting | 0.913 | 0.927 | 0.927 | 0.938 | 0.905 | 0.917 | 0.941 | **0.954** | 0.952 | 0.932 | 0.952 | 0.951 |
| XGBoost | 0.913 | 0.919 | 0.921 | 0.926 | 0.892 | 0.916 | 0.930 | 0.951 | 0.949 | 0.922 | **0.951** | 0.949 |
| LightGBM | 0.917 | 0.921 | 0.922 | 0.927 | 0.892 | 0.918 | 0.930 | **0.951** | 0.950 | 0.924 | 0.951 | 0.950 |
| Random forest | 0.912 | 0.923 | 0.923 | 0.926 | 0.890 | 0.912 | 0.924 | 0.945 | **0.948** | 0.918 | 0.946 | 0.948 |
| Neural network | 0.895 | 0.906 | 0.913 | 0.934 | 0.878 | 0.897 | 0.922 | 0.936 | 0.940 | 0.931 | **0.940** | 0.940 |
| Naïve Bayes | 0.862 | 0.664 | 0.664 | 0.671 | **0.876** | 0.828 | 0.722 | 0.861 | 0.855 | 0.745 | 0.825 | 0.820 |
| HDDA | 0.808 | 0.759 | 0.802 | 0.841 | 0.859 | 0.723 | 0.761 | 0.829 | 0.852 | 0.803 | 0.846 | **0.876** |

*24-month prognosis*

| Classifier | B | C | D | E | A | G1 | A+P | G2 | G12 | H1 | H2 | H12 |
| --- | --- | --- | --- | --- | --- | --- | --- | --- | --- | --- | --- | --- |
| Ridge | 0.888 | 0.887 | 0.888 | 0.907 | 0.886 | 0.886 | 0.919 | **0.953** | 0.943 | 0.906 | 0.949 | 0.941 |
| LASSO | 0.881 | 0.879 | 0.881 | 0.893 | 0.880 | 0.881 | 0.900 | **0.951** | 0.947 | 0.897 | 0.944 | 0.935 |
| Bayesian logistic | 0.894 | 0.902 | 0.898 | 0.924 | 0.879 | 0.890 | 0.927 | **0.961** | 0.948 | 0.924 | 0.952 | 0.947 |
| Linear SVM | 0.877 | 0.884 | 0.881 | 0.919 | 0.871 | 0.875 | 0.928 | 0.954 | 0.949 | 0.920 | **0.954** | 0.947 |
| LDA | 0.811 | 0.764 | 0.759 | 0.810 | 0.874 | 0.746 | 0.862 | 0.909 | **0.912** | 0.860 | 0.908 | 0.906 |
| GLM boosting | 0.887 | 0.892 | 0.886 | 0.909 | 0.875 | 0.883 | 0.930 | **0.969** | 0.964 | 0.906 | 0.966 | 0.962 |
| XGBoost | 0.904 | 0.888 | 0.892 | 0.915 | 0.886 | 0.886 | 0.927 | **0.959** | 0.955 | 0.908 | 0.959 | 0.957 |
| LightGBM | 0.906 | 0.887 | 0.893 | 0.918 | 0.891 | 0.889 | 0.932 | 0.961 | 0.958 | 0.910 | **0.961** | 0.960 |
| Random forest | 0.898 | 0.890 | 0.894 | 0.916 | 0.883 | 0.885 | 0.916 | 0.955 | 0.953 | 0.899 | **0.956** | 0.954 |
| Neural network | 0.885 | 0.889 | 0.899 | 0.924 | 0.871 | 0.882 | 0.908 | 0.949 | 0.945 | 0.916 | **0.950** | 0.948 |
| Naïve Bayes | 0.866 | 0.638 | 0.645 | 0.660 | 0.873 | 0.798 | 0.788 | **0.915** | 0.908 | 0.782 | 0.898 | 0.892 |
| HDDA | 0.805 | 0.776 | 0.803 | 0.823 | 0.862 | 0.762 | 0.794 | 0.892 | **0.903** | 0.802 | 0.856 | 0.872 |

*36-month prognosis*

| Classifier | B | C | D | E | A | G1 | A+P | G2 | G12 | H1 | H2 | H12 |
| --- | --- | --- | --- | --- | --- | --- | --- | --- | --- | --- | --- | --- |
| Ridge | 0.870 | 0.871 | 0.871 | 0.871 | 0.871 | 0.872 | 0.869 | 0.869 | 0.871 | 0.870 | 0.871 | **0.872** |
| LASSO | 0.865 | 0.864 | **0.865** | 0.860 | 0.863 | 0.860 | 0.860 | 0.861 | 0.860 | 0.862 | 0.859 | 0.863 |
| Bayesian logistic | 0.862 | 0.865 | 0.861 | 0.864 | 0.871 | 0.877 | 0.868 | 0.875 | 0.878 | **0.884** | 0.874 | 0.882 |
| Linear SVM | 0.853 | 0.854 | 0.853 | 0.850 | 0.856 | 0.865 | 0.854 | 0.853 | 0.858 | **0.869** | 0.857 | 0.862 |
| LDA | 0.758 | 0.697 | 0.680 | 0.693 | **0.861** | 0.794 | 0.637 | 0.777 | 0.759 | 0.807 | 0.797 | 0.766 |
| GLM boosting | 0.860 | 0.857 | 0.855 | 0.861 | 0.863 | **0.880** | 0.868 | 0.869 | 0.878 | 0.869 | 0.871 | 0.873 |
| XGBoost | 0.856 | 0.843 | 0.841 | 0.846 | 0.848 | 0.862 | 0.855 | 0.866 | **0.871** | 0.863 | 0.868 | 0.865 |
| LightGBM | 0.860 | 0.844 | 0.844 | 0.846 | 0.856 | 0.869 | 0.857 | 0.864 | **0.873** | 0.869 | 0.868 | 0.869 |
| Random forest | 0.862 | 0.836 | 0.840 | 0.842 | 0.849 | 0.862 | 0.864 | 0.869 | **0.874** | 0.862 | 0.869 | 0.871 |
| Neural network | 0.843 | 0.856 | 0.854 | 0.857 | 0.843 | 0.865 | 0.856 | 0.855 | 0.872 | **0.876** | 0.867 | 0.876 |
| Naïve Bayes | 0.813 | 0.569 | 0.573 | 0.592 | **0.818** | 0.773 | 0.723 | 0.754 | 0.754 | 0.716 | 0.728 | 0.736 |
| HDDA | 0.751 | 0.737 | 0.766 | 0.789 | **0.834** | 0.765 | 0.740 | 0.765 | 0.808 | 0.775 | 0.780 | 0.814 |

*Note:* feature sets defined in Table S6.

**Table S13.** Balanced accuracy for each feature set and classifier. Values are averaged across the ten analysis datasets; 95% confidence intervals are provided in Data S1. Bold indicates the highest value within each classifier.

*12-month prognosis*

| Classifier | B | C | D | E | A | G1 | A+P | G2 | G12 | H1 | H2 | H12 |
| --- | --- | --- | --- | --- | --- | --- | --- | --- | --- | --- | --- | --- |
| Ridge | 0.815 | 0.829 | 0.830 | 0.851 | 0.803 | 0.819 | 0.863 | **0.878** | 0.871 | 0.849 | 0.878 | 0.874 |
| LASSO | 0.826 | 0.827 | 0.826 | 0.834 | 0.825 | 0.825 | 0.841 | **0.880** | 0.858 | 0.844 | 0.859 | 0.854 |
| Bayesian logistic | 0.803 | 0.835 | 0.825 | 0.850 | 0.810 | 0.824 | 0.854 | **0.876** | 0.872 | 0.847 | 0.874 | 0.872 |
| Linear SVM | 0.795 | 0.821 | 0.809 | 0.844 | 0.780 | 0.819 | 0.866 | **0.885** | 0.877 | 0.843 | 0.878 | 0.866 |
| LDA | 0.797 | 0.716 | 0.715 | 0.736 | **0.845** | 0.561 | 0.834 | 0.730 | 0.764 | 0.692 | 0.750 | 0.767 |
| GLM boosting | 0.847 | 0.848 | 0.854 | 0.855 | 0.832 | 0.851 | 0.868 | **0.917** | 0.914 | 0.857 | 0.910 | 0.914 |
| XGBoost | 0.825 | 0.828 | 0.825 | 0.829 | 0.801 | 0.825 | 0.843 | 0.910 | 0.911 | 0.836 | 0.903 | **0.917** |
| LightGBM | 0.821 | 0.817 | 0.817 | 0.824 | 0.785 | 0.819 | 0.840 | **0.909** | 0.909 | 0.835 | 0.905 | 0.906 |
| Random forest | 0.834 | 0.849 | 0.848 | 0.849 | 0.787 | 0.837 | 0.842 | 0.915 | 0.918 | 0.846 | 0.916 | **0.921** |
| Neural network | 0.787 | 0.802 | 0.815 | 0.857 | 0.770 | 0.791 | 0.839 | 0.851 | 0.858 | 0.844 | 0.861 | **0.861** |
| Naïve Bayes | **0.787** | 0.627 | 0.630 | 0.630 | 0.749 | 0.762 | 0.668 | 0.785 | 0.781 | 0.687 | 0.747 | 0.756 |
| HDDA | 0.761 | 0.710 | 0.741 | 0.782 | **0.809** | 0.692 | 0.716 | 0.766 | 0.790 | 0.754 | 0.782 | 0.807 |

*24-month prognosis*

| Classifier | B | C | D | E | A | G1 | A+P | G2 | G12 | H1 | H2 | H12 |
| --- | --- | --- | --- | --- | --- | --- | --- | --- | --- | --- | --- | --- |
| Ridge | 0.828 | 0.830 | 0.832 | 0.845 | 0.825 | 0.829 | 0.862 | 0.898 | 0.877 | 0.844 | **0.902** | 0.884 |
| LASSO | 0.838 | 0.837 | 0.838 | 0.837 | 0.840 | 0.840 | 0.842 | **0.903** | 0.896 | 0.847 | 0.894 | 0.878 |
| Bayesian logistic | 0.812 | 0.823 | 0.822 | 0.856 | 0.813 | 0.807 | 0.868 | **0.911** | 0.887 | 0.859 | 0.905 | 0.888 |
| Linear SVM | 0.810 | 0.809 | 0.812 | 0.859 | 0.807 | 0.807 | 0.877 | 0.903 | 0.892 | 0.855 | **0.905** | 0.891 |
| LDA | 0.749 | 0.705 | 0.693 | 0.746 | 0.844 | 0.689 | 0.795 | 0.837 | **0.845** | 0.781 | 0.829 | 0.835 |
| GLM boosting | 0.834 | 0.831 | 0.832 | 0.857 | 0.839 | 0.833 | 0.881 | **0.931** | 0.926 | 0.855 | 0.929 | 0.927 |
| XGBoost | 0.848 | 0.818 | 0.824 | 0.847 | 0.824 | 0.812 | 0.862 | 0.920 | 0.916 | 0.836 | **0.923** | 0.917 |
| LightGBM | 0.846 | 0.817 | 0.826 | 0.854 | 0.828 | 0.815 | 0.873 | **0.925** | 0.925 | 0.842 | 0.922 | 0.925 |
| Random forest | 0.837 | 0.826 | 0.831 | 0.848 | 0.830 | 0.815 | 0.860 | **0.923** | 0.917 | 0.852 | 0.919 | 0.920 |
| Neural network | 0.797 | 0.809 | 0.816 | 0.857 | 0.784 | 0.796 | 0.835 | 0.893 | 0.888 | 0.843 | **0.901** | 0.892 |
| Naïve Bayes | 0.806 | 0.613 | 0.614 | 0.627 | 0.796 | 0.730 | 0.715 | **0.832** | 0.830 | 0.718 | 0.822 | 0.819 |
| HDDA | 0.751 | 0.720 | 0.749 | 0.764 | 0.807 | 0.703 | 0.727 | 0.811 | **0.833** | 0.738 | 0.786 | 0.795 |

*36-month prognosis*

| Classifier | B | C | D | E | A | G1 | A+P | G2 | G12 | H1 | H2 | H12 |
| --- | --- | --- | --- | --- | --- | --- | --- | --- | --- | --- | --- | --- |
| Ridge | 0.800 | 0.802 | 0.800 | 0.802 | 0.798 | 0.798 | 0.799 | 0.800 | 0.801 | 0.800 | **0.803** | 0.801 |
| LASSO | 0.806 | 0.806 | 0.806 | 0.804 | **0.808** | 0.807 | 0.804 | 0.804 | 0.807 | 0.805 | 0.804 | 0.805 |
| Bayesian logistic | 0.785 | 0.773 | 0.780 | 0.773 | 0.786 | 0.791 | 0.777 | 0.792 | 0.795 | 0.794 | 0.791 | **0.800** |
| Linear SVM | 0.784 | 0.779 | 0.774 | 0.773 | 0.783 | **0.789** | 0.778 | 0.778 | 0.782 | 0.787 | 0.782 | 0.783 |
| LDA | 0.716 | 0.641 | 0.623 | 0.642 | **0.783** | 0.729 | 0.611 | 0.712 | 0.694 | 0.735 | 0.728 | 0.701 |
| GLM boosting | 0.810 | 0.802 | 0.802 | 0.802 | 0.802 | 0.815 | 0.807 | **0.820** | 0.815 | 0.811 | 0.818 | 0.812 |
| XGBoost | 0.799 | 0.784 | 0.785 | 0.798 | 0.787 | 0.798 | 0.795 | **0.825** | 0.810 | 0.808 | 0.820 | 0.816 |
| LightGBM | 0.800 | 0.789 | 0.785 | 0.783 | 0.788 | 0.801 | 0.798 | **0.821** | 0.815 | 0.808 | 0.819 | 0.818 |
| Random forest | 0.805 | 0.792 | 0.799 | 0.799 | 0.794 | 0.807 | 0.800 | 0.835 | 0.831 | 0.802 | **0.835** | 0.828 |
| Neural network | 0.772 | 0.773 | 0.777 | 0.778 | 0.776 | 0.791 | 0.775 | 0.786 | 0.798 | 0.795 | 0.795 | **0.800** |
| Naïve Bayes | 0.750 | 0.543 | 0.553 | 0.579 | **0.771** | 0.708 | 0.684 | 0.704 | 0.698 | 0.678 | 0.689 | 0.686 |
| HDDA | 0.710 | 0.696 | 0.719 | 0.730 | **0.776** | 0.698 | 0.678 | 0.713 | 0.737 | 0.706 | 0.712 | 0.739 |

*Note:* feature sets defined in Table S6.

**Table S14.** Macro-F1 for each feature set and classifier. Values are averaged across the ten analysis datasets; 95% confidence intervals are provided in Data S1. Bold indicates the highest value within each classifier.

*12-month prognosis*

| Classifier | B | C | D | E | A | G1 | A+P | G2 | G12 | H1 | H2 | H12 |
| --- | --- | --- | --- | --- | --- | --- | --- | --- | --- | --- | --- | --- |
| Ridge | 0.802 | 0.818 | 0.818 | 0.840 | 0.791 | 0.807 | 0.854 | 0.868 | 0.861 | 0.837 | **0.869** | 0.867 |
| LASSO | 0.791 | 0.807 | 0.792 | 0.810 | 0.791 | 0.791 | 0.828 | **0.867** | 0.845 | 0.826 | 0.848 | 0.839 |
| Bayesian logistic | 0.796 | 0.828 | 0.818 | 0.847 | 0.799 | 0.819 | 0.850 | 0.868 | 0.867 | 0.845 | **0.869** | 0.869 |
| Linear SVM | 0.791 | 0.817 | 0.805 | 0.839 | 0.776 | 0.814 | 0.861 | **0.876** | 0.870 | 0.840 | 0.872 | 0.862 |
| LDA | 0.779 | 0.707 | 0.707 | 0.727 | **0.815** | 0.546 | 0.805 | 0.714 | 0.757 | 0.671 | 0.739 | 0.760 |
| GLM boosting | 0.822 | 0.828 | 0.834 | 0.842 | 0.807 | 0.827 | 0.854 | **0.897** | 0.897 | 0.841 | 0.893 | 0.896 |
| XGBoost | 0.813 | 0.822 | 0.820 | 0.825 | 0.782 | 0.816 | 0.838 | 0.894 | 0.896 | 0.830 | 0.889 | **0.901** |
| LightGBM | 0.812 | 0.815 | 0.815 | 0.822 | 0.779 | 0.812 | 0.835 | 0.894 | **0.895** | 0.830 | 0.892 | 0.893 |
| Random forest | 0.817 | 0.834 | 0.832 | 0.834 | 0.776 | 0.821 | 0.827 | 0.893 | 0.897 | 0.828 | 0.893 | **0.900** |
| Neural network | 0.782 | 0.797 | 0.810 | 0.851 | 0.764 | 0.785 | 0.836 | 0.845 | 0.853 | 0.839 | **0.857** | 0.856 |
| Naïve Bayes | **0.765** | 0.628 | 0.631 | 0.634 | 0.753 | 0.738 | 0.658 | 0.753 | 0.757 | 0.678 | 0.726 | 0.737 |
| HDDA | 0.747 | 0.693 | 0.724 | 0.762 | **0.792** | 0.679 | 0.719 | 0.745 | 0.771 | 0.736 | 0.764 | 0.787 |

*24-month prognosis*

| Classifier | B | C | D | E | A | G1 | A+P | G2 | G12 | H1 | H2 | H12 |
| --- | --- | --- | --- | --- | --- | --- | --- | --- | --- | --- | --- | --- |
| Ridge | 0.820 | 0.820 | 0.822 | 0.836 | 0.816 | 0.820 | 0.855 | 0.893 | 0.871 | 0.837 | **0.898** | 0.878 |
| LASSO | 0.819 | 0.817 | 0.819 | 0.823 | 0.820 | 0.820 | 0.831 | **0.897** | 0.890 | 0.833 | 0.889 | 0.872 |
| Bayesian logistic | 0.808 | 0.819 | 0.816 | 0.851 | 0.808 | 0.804 | 0.865 | **0.908** | 0.882 | 0.856 | 0.901 | 0.883 |
| Linear SVM | 0.801 | 0.802 | 0.805 | 0.852 | 0.798 | 0.799 | 0.872 | 0.898 | 0.887 | 0.851 | **0.899** | 0.885 |
| LDA | 0.742 | 0.701 | 0.690 | 0.743 | 0.827 | 0.683 | 0.783 | 0.830 | **0.840** | 0.775 | 0.824 | 0.832 |
| GLM boosting | 0.815 | 0.813 | 0.813 | 0.841 | 0.819 | 0.815 | 0.872 | **0.923** | 0.917 | 0.840 | 0.920 | 0.917 |
| XGBoost | 0.836 | 0.807 | 0.814 | 0.838 | 0.814 | 0.800 | 0.858 | 0.913 | 0.910 | 0.829 | **0.916** | 0.910 |
| LightGBM | 0.834 | 0.807 | 0.815 | 0.844 | 0.820 | 0.804 | 0.866 | **0.917** | 0.917 | 0.835 | 0.914 | 0.917 |
| Random forest | 0.823 | 0.810 | 0.813 | 0.833 | 0.817 | 0.796 | 0.850 | **0.916** | 0.911 | 0.839 | 0.912 | 0.913 |
| Neural network | 0.793 | 0.805 | 0.811 | 0.851 | 0.781 | 0.792 | 0.832 | 0.888 | 0.882 | 0.839 | **0.897** | 0.887 |
| Naïve Bayes | 0.792 | 0.612 | 0.613 | 0.627 | 0.794 | 0.718 | 0.692 | 0.815 | **0.820** | 0.704 | 0.807 | 0.809 |
| HDDA | 0.743 | 0.715 | 0.743 | 0.758 | 0.798 | 0.698 | 0.720 | 0.804 | **0.825** | 0.731 | 0.777 | 0.786 |

*36-month prognosis*

| Classifier | B | C | D | E | A | G1 | A+P | G2 | G12 | H1 | H2 | H12 |
| --- | --- | --- | --- | --- | --- | --- | --- | --- | --- | --- | --- | --- |
| Ridge | 0.796 | 0.799 | 0.797 | 0.799 | 0.795 | 0.795 | 0.795 | 0.797 | 0.797 | 0.797 | **0.799** | 0.797 |
| LASSO | 0.799 | 0.799 | 0.799 | 0.796 | **0.800** | 0.800 | 0.796 | 0.797 | 0.800 | 0.798 | 0.796 | 0.798 |
| Bayesian logistic | 0.784 | 0.772 | 0.779 | 0.772 | 0.784 | 0.789 | 0.776 | 0.791 | 0.794 | 0.793 | 0.789 | **0.798** |
| Linear SVM | 0.780 | 0.775 | 0.770 | 0.769 | 0.778 | **0.786** | 0.774 | 0.774 | 0.778 | 0.783 | 0.778 | 0.779 |
| LDA | 0.714 | 0.640 | 0.622 | 0.641 | **0.777** | 0.728 | 0.609 | 0.711 | 0.693 | 0.735 | 0.728 | 0.700 |
| GLM boosting | 0.801 | 0.793 | 0.793 | 0.793 | 0.794 | 0.806 | 0.800 | **0.812** | 0.806 | 0.803 | 0.809 | 0.803 |
| XGBoost | 0.793 | 0.779 | 0.780 | 0.792 | 0.781 | 0.793 | 0.790 | **0.820** | 0.805 | 0.802 | 0.815 | 0.811 |
| LightGBM | 0.794 | 0.785 | 0.780 | 0.779 | 0.784 | 0.796 | 0.794 | **0.816** | 0.810 | 0.803 | 0.815 | 0.813 |
| Random forest | 0.798 | 0.785 | 0.792 | 0.792 | 0.788 | 0.798 | 0.794 | 0.828 | 0.823 | 0.793 | **0.828** | 0.820 |
| Neural network | 0.771 | 0.773 | 0.776 | 0.777 | 0.775 | 0.789 | 0.774 | 0.785 | 0.797 | 0.793 | 0.793 | **0.798** |
| Naïve Bayes | 0.738 | 0.541 | 0.551 | 0.577 | **0.765** | 0.700 | 0.671 | 0.695 | 0.692 | 0.670 | 0.683 | 0.681 |
| HDDA | 0.709 | 0.695 | 0.718 | 0.728 | **0.771** | 0.697 | 0.677 | 0.712 | 0.736 | 0.705 | 0.711 | 0.737 |

*Note:* feature sets defined in Table S6.

**Table S15.** Best model (GLM boosting with centile-map deep features, feature set G2): all metrics with 95% CIs.

| Metric | 12 months | 24 months | 36 months |
| --- | --- | --- | --- |
| AUC | 0.954 (0.931–0.974) | 0.969 (0.949–0.985) | 0.869 (0.820–0.916) |
| Brier | 0.074 (0.058–0.091) | 0.064 (0.047–0.083) | 0.136 (0.109–0.162) |
| Accuracy | 0.909 (0.880–0.936) | 0.924 (0.893–0.951) | 0.813 (0.762–0.867) |
| Balanced accuracy | 0.917 (0.894–0.937) | 0.931 (0.905–0.954) | 0.820 (0.773–0.870) |
| Precision (case) | 0.970 (0.960–0.978) | 0.977 (0.966–0.986) | 0.939 (0.912–0.960) |
| Recall (case) | 0.898 (0.857–0.936) | 0.895 (0.844–0.939) | 0.691 (0.599–0.791) |
| F1 (case) | 0.933 (0.908–0.953) | 0.934 (0.904–0.958) | 0.796 (0.728–0.861) |
| Precision (control) | 0.799 (0.733–0.866) | 0.861 (0.801–0.919) | 0.734 (0.662–0.812) |
| Recall (control) | 0.935 (0.918–0.950) | 0.968 (0.955–0.980) | 0.950 (0.934–0.964) |
| F1 (control) | 0.862 (0.820–0.899) | 0.912 (0.876–0.943) | 0.828 (0.779–0.874) |
| Macro-F1 | 0.897 (0.866–0.926) | 0.923 (0.892–0.950) | 0.812 (0.758–0.865) |

*Note:* feature sets defined in Table S6.

### S8. Paired comparisons

**Table S16.** Paired differences between feature sets for ridge logistic regression, with 95% confidence intervals. P values are from two-sided participant-bootstrap tests. Holm P adjusts across the 12 classifiers within each comparison, horizon and metric. Global Holm P adjusts across all comparisons, classifiers and horizons within this analysis, comprising 828 tests per metric (Table S9).

| Comparison | Horizon | Metric | Difference (95% CI) | P | Holm P | Global Holm P |
| --- | --- | --- | --- | --- | --- | --- |
| ROI PCA beyond feature set A (B − A) | 12 | AUC | 0.008 (0.002–0.013) | 0.005 | 0.040 | 1.000 |
| ROI PCA beyond feature set A (B − A) | 12 | Bal. acc. | 0.012 (0.002–0.022) | 0.018 | 0.126 | 1.000 |
| ROI PCA beyond feature set A (B − A) | 12 | Macro-F1 | 0.011 (0.001–0.020) | 0.021 | 0.147 | 1.000 |
| ROI PCA beyond feature set A (B − A) | 24 | AUC | 0.002 (-0.002–0.006) | 0.337 | 1.000 | 1.000 |
| ROI PCA beyond feature set A (B − A) | 24 | Bal. acc. | 0.004 (-0.003–0.010) | 0.305 | 1.000 | 1.000 |
| ROI PCA beyond feature set A (B − A) | 24 | Macro-F1 | 0.004 (-0.002–0.010) | 0.217 | 1.000 | 1.000 |
| ROI PCA beyond feature set A (B − A) | 36 | AUC | -0.000 (-0.003–0.002) | 0.889 | 1.000 | 1.000 |
| ROI PCA beyond feature set A (B − A) | 36 | Bal. acc. | 0.001 (-0.003–0.006) | 0.541 | 1.000 | 1.000 |
| ROI PCA beyond feature set A (B − A) | 36 | Macro-F1 | 0.001 (-0.003–0.006) | 0.588 | 1.000 | 1.000 |
| ROI radiomics beyond feature set A (C − A) | 12 | AUC | 0.024 (0.013–0.036) | <0.001 | <0.001 | <0.001 |
| ROI radiomics beyond feature set A (C − A) | 12 | Bal. acc. | 0.026 (0.011–0.040) | <0.001 | <0.001 | <0.001 |
| ROI radiomics beyond feature set A (C − A) | 12 | Macro-F1 | 0.027 (0.012–0.042) | <0.001 | <0.001 | <0.001 |
| ROI radiomics beyond feature set A (C − A) | 24 | AUC | 0.001 (-0.005–0.006) | 0.730 | 1.000 | 1.000 |
| ROI radiomics beyond feature set A (C − A) | 24 | Bal. acc. | 0.005 (-0.002–0.012) | 0.125 | 0.875 | 1.000 |
| ROI radiomics beyond feature set A (C − A) | 24 | Macro-F1 | 0.005 (-0.001–0.011) | 0.125 | 0.875 | 1.000 |
| ROI radiomics beyond feature set A (C − A) | 36 | AUC | 0.000 (-0.002–0.003) | 0.806 | 1.000 | 1.000 |
| ROI radiomics beyond feature set A (C − A) | 36 | Bal. acc. | 0.004 (-0.001–0.009) | 0.096 | 0.864 | 1.000 |
| ROI radiomics beyond feature set A (C − A) | 36 | Macro-F1 | 0.004 (-0.001–0.009) | 0.128 | 1.000 | 1.000 |
| ROI PCA + radiomics beyond feature set A (D − A) | 12 | AUC | 0.022 (0.013–0.033) | <0.001 | <0.001 | <0.001 |
| ROI PCA + radiomics beyond feature set A (D − A) | 12 | Bal. acc. | 0.027 (0.013–0.040) | <0.001 | <0.001 | <0.001 |
| ROI PCA + radiomics beyond feature set A (D − A) | 12 | Macro-F1 | 0.027 (0.013–0.041) | <0.001 | <0.001 | <0.001 |
| ROI PCA + radiomics beyond feature set A (D − A) | 24 | AUC | 0.002 (-0.003–0.007) | 0.366 | 1.000 | 1.000 |
| ROI PCA + radiomics beyond feature set A (D − A) | 24 | Bal. acc. | 0.007 (-0.001–0.015) | 0.069 | 0.552 | 1.000 |
| ROI PCA + radiomics beyond feature set A (D − A) | 24 | Macro-F1 | 0.007 (-0.000–0.014) | 0.055 | 0.440 | 1.000 |
| ROI PCA + radiomics beyond feature set A (D − A) | 36 | AUC | 0.000 (-0.003–0.003) | 0.897 | 1.000 | 1.000 |
| ROI PCA + radiomics beyond feature set A (D − A) | 36 | Bal. acc. | 0.002 (-0.003–0.008) | 0.435 | 1.000 | 1.000 |
| ROI PCA + radiomics beyond feature set A (D − A) | 36 | Macro-F1 | 0.002 (-0.004–0.008) | 0.520 | 1.000 | 1.000 |
| Centile profiles beyond feature set A ((A+P) − A) | 12 | AUC | 0.044 (0.026–0.064) | <0.001 | <0.001 | <0.001 |
| Centile profiles beyond feature set A ((A+P) − A) | 12 | Bal. acc. | 0.060 (0.041–0.080) | <0.001 | <0.001 | <0.001 |
| Centile profiles beyond feature set A ((A+P) − A) | 12 | Macro-F1 | 0.063 (0.042–0.087) | <0.001 | <0.001 | <0.001 |
| Centile profiles beyond feature set A ((A+P) − A) | 24 | AUC | 0.032 (0.012–0.054) | <0.001 | <0.001 | <0.001 |
| Centile profiles beyond feature set A ((A+P) − A) | 24 | Bal. acc. | 0.037 (0.016–0.058) | <0.001 | <0.001 | <0.001 |
| Centile profiles beyond feature set A ((A+P) − A) | 24 | Macro-F1 | 0.040 (0.018–0.062) | <0.001 | <0.001 | <0.001 |
| Centile profiles beyond feature set A ((A+P) − A) | 36 | AUC | -0.001 (-0.006–0.003) | 0.571 | 1.000 | 1.000 |
| Centile profiles beyond feature set A ((A+P) − A) | 36 | Bal. acc. | 0.001 (-0.006–0.007) | 0.877 | 1.000 | 1.000 |
| Centile profiles beyond feature set A ((A+P) − A) | 36 | Macro-F1 | 0.001 (-0.006–0.007) | 0.889 | 1.000 | 1.000 |
| Centile profiles beyond feature set D (E − D) | 12 | AUC | 0.015 (0.008–0.023) | <0.001 | <0.001 | <0.001 |
| Centile profiles beyond feature set D (E − D) | 12 | Bal. acc. | 0.021 (0.009–0.032) | <0.001 | <0.001 | <0.001 |
| Centile profiles beyond feature set D (E − D) | 12 | Macro-F1 | 0.022 (0.010–0.034) | <0.001 | <0.001 | <0.001 |
| Centile profiles beyond feature set D (E − D) | 24 | AUC | 0.018 (0.006–0.032) | 0.001 | 0.005 | 0.631 |
| Centile profiles beyond feature set D (E − D) | 24 | Bal. acc. | 0.013 (0.000–0.026) | 0.044 | 0.200 | 1.000 |
| Centile profiles beyond feature set D (E − D) | 24 | Macro-F1 | 0.013 (0.001–0.026) | 0.039 | 0.156 | 1.000 |
| Centile profiles beyond feature set D (E − D) | 36 | AUC | 0.000 (-0.003–0.003) | 0.853 | 1.000 | 1.000 |
| Centile profiles beyond feature set D (E − D) | 36 | Bal. acc. | 0.002 (-0.004–0.008) | 0.580 | 1.000 | 1.000 |
| Centile profiles beyond feature set D (E − D) | 36 | Macro-F1 | 0.002 (-0.005–0.009) | 0.569 | 1.000 | 1.000 |
| ROI PCA + radiomics beyond feature set A+P (E − (A+P)) | 12 | AUC | -0.006 (-0.013–0.000) | 0.061 | 0.366 | 1.000 |
| ROI PCA + radiomics beyond feature set A+P (E − (A+P)) | 12 | Bal. acc. | -0.013 (-0.024–-0.000) | 0.048 | 0.336 | 1.000 |
| ROI PCA + radiomics beyond feature set A+P (E − (A+P)) | 12 | Macro-F1 | -0.014 (-0.027–-0.001) | 0.042 | 0.294 | 1.000 |
| ROI PCA + radiomics beyond feature set A+P (E − (A+P)) | 24 | AUC | -0.012 (-0.022–-0.002) | 0.010 | 0.080 | 1.000 |
| ROI PCA + radiomics beyond feature set A+P (E − (A+P)) | 24 | Bal. acc. | -0.017 (-0.032–-0.003) | 0.019 | 0.152 | 1.000 |
| ROI PCA + radiomics beyond feature set A+P (E − (A+P)) | 24 | Macro-F1 | -0.020 (-0.035–-0.005) | 0.005 | 0.050 | 1.000 |
| ROI PCA + radiomics beyond feature set A+P (E − (A+P)) | 36 | AUC | 0.002 (-0.003–0.006) | 0.408 | 1.000 | 1.000 |
| ROI PCA + radiomics beyond feature set A+P (E − (A+P)) | 36 | Bal. acc. | 0.003 (-0.004–0.011) | 0.346 | 1.000 | 1.000 |
| ROI PCA + radiomics beyond feature set A+P (E − (A+P)) | 36 | Macro-F1 | 0.003 (-0.004–0.010) | 0.369 | 1.000 | 1.000 |

*Note:* feature sets defined in Table S6.

**Table S17.** Summary of paired comparisons across the 12 classifiers (AUC): number of classifiers with a Holm-significant gain (+) or loss (−), and median difference.

| Comparison | Horizon | + (Holm) | − (Holm) | Median Δ | Range |
| --- | --- | --- | --- | --- | --- |
| Feature set G2 vs feature set G1: centile-map vs raw-map deep features (G2 − G1) | 12 | 12 | 0 | +0.038 | +0.032 to +0.237 |
| Feature set G2 vs feature set G1: centile-map vs raw-map deep features (G2 − G1) | 24 | 12 | 0 | +0.073 | +0.067 to +0.162 |
| Feature set G2 vs feature set G1: centile-map vs raw-map deep features (G2 − G1) | 36 | 0 | 0 | -0.004 | -0.019 to +0.007 |
| Centile profiles beyond feature set G1 (H1 − G1) | 12 | 8 | 1 | +0.022 | -0.083 to +0.180 |
| Centile profiles beyond feature set G1 (H1 − G1) | 24 | 10 | 0 | +0.023 | -0.016 to +0.114 |
| Centile profiles beyond feature set G1 (H1 − G1) | 36 | 0 | 1 | +0.001 | -0.057 to +0.013 |
| Raw-map deep features beyond feature set A+P (H1 − (A+P)) | 12 | 1 | 3 | -0.007 | -0.147 to +0.041 |
| Raw-map deep features beyond feature set A+P (H1 − (A+P)) | 24 | 0 | 1 | -0.006 | -0.024 to +0.008 |
| Raw-map deep features beyond feature set A+P (H1 − (A+P)) | 36 | 4 | 0 | +0.010 | -0.006 to +0.170 |
| Feature set G2 vs feature set A+P: centile-map deep features vs centile profiles (G2 − (A+P)) | 12 | 5 | 1 | +0.017 | -0.090 to +0.139 |
| Feature set G2 vs feature set A+P: centile-map deep features vs centile profiles (G2 − (A+P)) | 24 | 12 | 0 | +0.039 | +0.026 to +0.127 |
| Feature set G2 vs feature set A+P: centile-map deep features vs centile profiles (G2 − (A+P)) | 36 | 1 | 0 | +0.006 | -0.001 to +0.140 |
| Centile-map deep features beyond feature set A+P (H2 − (A+P)) | 12 | 8 | 1 | +0.015 | -0.058 to +0.102 |
| Centile-map deep features beyond feature set A+P (H2 − (A+P)) | 24 | 12 | 0 | +0.038 | +0.025 to +0.110 |
| Centile-map deep features beyond feature set A+P (H2 − (A+P)) | 36 | 2 | 0 | +0.006 | -0.001 to +0.159 |
| Centile-map deep features beyond feature set A (G2 − A) | 12 | 9 | 1 | +0.053 | -0.103 to +0.060 |
| Centile-map deep features beyond feature set A (G2 − A) | 24 | 9 | 0 | +0.071 | +0.030 to +0.094 |
| Centile-map deep features beyond feature set A (G2 − A) | 36 | 0 | 2 | +0.001 | -0.084 to +0.019 |
| Raw-map deep features beyond feature set A (G1 − A) | 12 | 2 | 3 | +0.011 | -0.340 to +0.026 |
| Raw-map deep features beyond feature set A (G1 − A) | 24 | 0 | 3 | +0.000 | -0.128 to +0.010 |
| Raw-map deep features beyond feature set A (G1 − A) | 36 | 1 | 2 | +0.008 | -0.069 to +0.022 |

*Note:* feature sets defined in Table S6.

**Table S18.** Per-classifier paired AUC differences (95% CI; Holm P across classifiers) for the key deep-feature comparisons at 12 and 24 months.

*Feature set G2 vs feature set G1: centile-map vs raw-map deep features (G2 − G1)*

| Classifier | 12 mo Δ (95% CI) | Holm P | 24 mo Δ (95% CI) | Holm P |
| --- | --- | --- | --- | --- |
| Ridge | 0.040 (0.024–0.058) | <0.001 | 0.067 (0.040–0.098) | <0.001 |
| LASSO | 0.057 (0.036–0.078) | <0.001 | 0.070 (0.040–0.104) | <0.001 |
| Bayesian logistic | 0.032 (0.016–0.050) | <0.001 | 0.072 (0.042–0.103) | <0.001 |
| Linear SVM | 0.039 (0.024–0.056) | <0.001 | 0.079 (0.050–0.110) | <0.001 |
| LDA | 0.237 (0.206–0.268) | <0.001 | 0.162 (0.129–0.197) | <0.001 |
| GLM boosting | 0.037 (0.019–0.058) | <0.001 | 0.086 (0.056–0.120) | <0.001 |
| XGBoost | 0.035 (0.016–0.056) | <0.001 | 0.073 (0.041–0.107) | <0.001 |
| LightGBM | 0.034 (0.016–0.054) | <0.001 | 0.072 (0.040–0.105) | <0.001 |
| Random forest | 0.033 (0.012–0.056) | 0.002 | 0.071 (0.038–0.104) | <0.001 |
| Neural network | 0.039 (0.024–0.057) | <0.001 | 0.068 (0.045–0.093) | <0.001 |
| Naïve Bayes | 0.033 (0.005–0.064) | 0.022 | 0.117 (0.073–0.161) | <0.001 |
| HDDA | 0.106 (0.086–0.129) | <0.001 | 0.130 (0.099–0.161) | <0.001 |

*Centile profiles beyond feature set G1 (H1 − G1)*

| Classifier | 12 mo Δ (95% CI) | Holm P | 24 mo Δ (95% CI) | Holm P |
| --- | --- | --- | --- | --- |
| Ridge | 0.022 (0.012–0.033) | <0.001 | 0.020 (0.007–0.035) | 0.016 |
| LASSO | 0.028 (0.011–0.045) | <0.001 | 0.017 (0.004–0.031) | 0.039 |
| Bayesian logistic | 0.022 (0.010–0.034) | <0.001 | 0.035 (0.016–0.055) | <0.001 |
| Linear SVM | 0.023 (0.012–0.034) | <0.001 | 0.046 (0.026–0.067) | <0.001 |
| LDA | 0.180 (0.148–0.210) | <0.001 | 0.114 (0.087–0.141) | <0.001 |
| GLM boosting | 0.016 (0.006–0.026) | 0.004 | 0.023 (0.006–0.041) | 0.036 |
| XGBoost | 0.006 (-0.002–0.015) | 0.117 | 0.022 (0.007–0.038) | 0.036 |
| LightGBM | 0.007 (0.000–0.014) | 0.098 | 0.021 (0.008–0.035) | 0.021 |
| Random forest | 0.006 (0.001–0.011) | 0.069 | 0.014 (-0.002–0.030) | 0.158 |
| Neural network | 0.034 (0.021–0.048) | <0.001 | 0.034 (0.020–0.050) | <0.001 |
| Naïve Bayes | -0.083 (-0.105–-0.060) | <0.001 | -0.016 (-0.045–0.013) | 0.300 |
| HDDA | 0.079 (0.054–0.106) | <0.001 | 0.040 (0.008–0.068) | 0.036 |

*Raw-map deep features beyond feature set A+P (H1 − (A+P))*

| Classifier | 12 mo Δ (95% CI) | Holm P | 24 mo Δ (95% CI) | Holm P |
| --- | --- | --- | --- | --- |
| Ridge | -0.011 (-0.017–-0.006) | <0.001 | -0.013 (-0.023–-0.003) | 0.070 |
| LASSO | -0.007 (-0.013–-0.001) | 0.088 | -0.002 (-0.010–0.005) | 1.000 |
| Bayesian logistic | -0.002 (-0.007–0.003) | 1.000 | -0.003 (-0.011–0.006) | 1.000 |
| Linear SVM | -0.010 (-0.017–-0.003) | 0.018 | -0.007 (-0.016–0.002) | 0.690 |
| LDA | -0.147 (-0.171–-0.121) | <0.001 | -0.002 (-0.032–0.027) | 1.000 |
| GLM boosting | -0.008 (-0.018–0.001) | 0.520 | -0.024 (-0.039–-0.010) | 0.024 |
| XGBoost | -0.008 (-0.020–0.005) | 1.000 | -0.019 (-0.036–-0.003) | 0.225 |
| LightGBM | -0.005 (-0.018–0.007) | 1.000 | -0.022 (-0.039–-0.005) | 0.066 |
| Random forest | -0.006 (-0.019–0.007) | 1.000 | -0.018 (-0.035–-0.002) | 0.232 |
| Neural network | 0.009 (0.000–0.017) | 0.246 | 0.008 (-0.000–0.016) | 0.399 |
| Naïve Bayes | 0.023 (0.004–0.041) | 0.112 | -0.006 (-0.027–0.015) | 1.000 |
| HDDA | 0.041 (0.017–0.065) | 0.010 | 0.008 (-0.003–0.018) | 0.840 |

*Centile-map deep features beyond feature set A+P (H2 − (A+P))*

| Classifier | 12 mo Δ (95% CI) | Holm P | 24 mo Δ (95% CI) | Holm P |
| --- | --- | --- | --- | --- |
| Ridge | 0.007 (-0.001–0.014) | 0.165 | 0.031 (0.016–0.046) | <0.001 |
| LASSO | 0.011 (0.005–0.018) | <0.001 | 0.044 (0.029–0.062) | <0.001 |
| Bayesian logistic | 0.009 (0.003–0.016) | 0.025 | 0.025 (0.014–0.039) | <0.001 |
| Linear SVM | 0.006 (-0.000–0.013) | 0.165 | 0.027 (0.013–0.042) | <0.001 |
| LDA | -0.058 (-0.082–-0.035) | <0.001 | 0.046 (0.016–0.079) | 0.012 |
| GLM boosting | 0.012 (-0.000–0.025) | 0.165 | 0.036 (0.016–0.059) | <0.001 |
| XGBoost | 0.021 (0.007–0.036) | 0.021 | 0.032 (0.010–0.056) | 0.012 |
| LightGBM | 0.021 (0.007–0.037) | 0.024 | 0.029 (0.007–0.054) | 0.012 |
| Random forest | 0.021 (0.005–0.039) | 0.048 | 0.040 (0.012–0.071) | 0.008 |
| Neural network | 0.018 (0.009–0.028) | <0.001 | 0.042 (0.027–0.059) | <0.001 |
| Naïve Bayes | 0.102 (0.073–0.131) | <0.001 | 0.110 (0.083–0.139) | <0.001 |
| HDDA | 0.085 (0.060–0.111) | <0.001 | 0.062 (0.048–0.077) | <0.001 |

*Note:* feature sets defined in Table S6.

Balanced accuracy, macro-F1 and all other metrics for every comparison and classifier, with raw and Holm P values, are provided in the supplementary data files (Data S1–S2).

### S9. Sensitivity analysis: reference membership (downsampled cross-fitted versus full-reference charts)

**Reference overlap.** All prognostic controls belonged to the 957-participant normative reference sample, whereas no case did. Their charts were therefore fitted partly to their own scans. This overlap could favour control knees in the centile representation and inflate discrimination.

**Chart construction.** We downsampled each map by averaging non-overlapping 3 × 3 pixel blocks and assigning each mean to the block centre, giving 4,227 femoral and 4,078 tibial pixels (8,305 in total). Reference participants were randomly divided into two halves. The model in S2 was fitted by maximum likelihood to the full reference and separately to each half (mgcv 1.9.3; R 4.4.0), retaining the fitted fixed effects, smooth coefficients and variance components. The original main-analysis charts are termed **full-resolution full-reference charts**.

**Downsampled cross-fitted charts (method 1).** Each reference participant was scored with the chart fitted to the other half. Other scans were scored by both half-reference charts, and their local centiles were averaged pixel by pixel (crossfit_avg). We also evaluated a variant that scored these scans with one randomly assigned half-reference chart (crossfit_single).

**Downsampled full-reference charts (method 2).** Every scan was scored with the downsampled chart fitted to the full reference. For both methods, local centiles, connected-deficit features and compression weights were recomputed using the main-analysis procedure. Compression weights were then held fixed for classifier fitting.

**Table S19.** Ridge AUC of clinical + ROI mean thickness (feature set A) and with centile profiles (feature set A+P) under each reference arm, and the centile increment.

| Horizon | Reference arm | AUC, feature set A | AUC, feature set A+P (95% CI) | Δ AUC (95% CI) | P (raw) |
| --- | --- | --- | --- | --- | --- |
| 12 | Full-resolution full-reference (original charts) | 0.897 | 0.942 (0.918–0.962) | +0.044 (+0.028, +0.064) | <0.001 |
| 12 | Downsampled full-reference (method 2) | 0.897 | 0.928 (0.902–0.950) | +0.031 (+0.018, +0.046) | <0.001 |
| 12 | Downsampled cross-fitted (method 1) | 0.897 | 0.926 (0.900–0.948) | +0.029 (+0.017, +0.044) | <0.001 |
| 12 | Downsampled cross-fitted, single-chart variant | 0.897 | 0.926 (0.900–0.948) | +0.029 (+0.017, +0.044) | <0.001 |
| 24 | Full-resolution full-reference (original charts) | 0.886 | 0.919 (0.884–0.952) | +0.032 (+0.012, +0.056) | 0.002 |
| 24 | Downsampled full-reference (method 2) | 0.886 | 0.906 (0.870–0.941) | +0.020 (+0.007, +0.035) | <0.001 |
| 24 | Downsampled cross-fitted (method 1) | 0.886 | 0.903 (0.866–0.939) | +0.017 (+0.005, +0.031) | 0.004 |
| 24 | Downsampled cross-fitted, single-chart variant | 0.886 | 0.901 (0.864–0.938) | +0.015 (+0.004, +0.028) | 0.010 |
| 36 | Full-resolution full-reference (original charts) | 0.871 | 0.869 (0.819–0.914) | -0.001 (-0.006, +0.003) | 0.620 |
| 36 | Downsampled full-reference (method 2) | 0.871 | 0.871 (0.820–0.916) | +0.000 (-0.004, +0.004) | 0.952 |
| 36 | Downsampled cross-fitted (method 1) | 0.871 | 0.870 (0.819–0.916) | -0.001 (-0.005, +0.003) | 0.650 |
| 36 | Downsampled cross-fitted, single-chart variant | 0.871 | 0.872 (0.821–0.916) | +0.001 (-0.004, +0.006) | 0.693 |

*Note:* feature sets defined in Table S6.

**Table S20.** Across the 12 classifiers: number with Holm-significant gain (+) or loss (−), median and range of differences.

| Metric | Horizon | Comparison | + | − | Median | Range |
| --- | --- | --- | --- | --- | --- | --- |
| AUC | 12 | Centile profiles beyond feature set A, downsampled cross-fitted charts ((A+P1) − A) | 9 | 3 | +0.025 | -0.102 to +0.037 |
| AUC | 12 | Centile profiles beyond feature set A, downsampled full-reference charts ((A+P2) − A) | 8 | 2 | +0.027 | -0.101 to +0.041 |
| AUC | 12 | Feature set A+P, downsampled cross-fitted − downsampled full-reference charts ((A+P1) − (A+P2)) | 0 | 3 | -0.002 | -0.010 to +0.008 |
| AUC | 24 | Centile profiles beyond feature set A, downsampled cross-fitted charts ((A+P1) − A) | 8 | 2 | +0.028 | -0.072 to +0.054 |
| AUC | 24 | Centile profiles beyond feature set A, downsampled full-reference charts ((A+P2) − A) | 8 | 2 | +0.030 | -0.062 to +0.059 |
| AUC | 24 | Feature set A+P, downsampled cross-fitted − downsampled full-reference charts ((A+P1) − (A+P2)) | 0 | 1 | -0.003 | -0.012 to +0.002 |
| AUC | 36 | Centile profiles beyond feature set A, downsampled cross-fitted charts ((A+P1) − A) | 0 | 3 | +0.002 | -0.182 to +0.011 |
| AUC | 36 | Centile profiles beyond feature set A, downsampled full-reference charts ((A+P2) − A) | 0 | 2 | +0.002 | -0.198 to +0.014 |
| AUC | 36 | Feature set A+P, downsampled cross-fitted − downsampled full-reference charts ((A+P1) − (A+P2)) | 0 | 0 | +0.001 | -0.003 to +0.015 |
| Balanced accuracy | 12 | Centile profiles beyond feature set A, downsampled cross-fitted charts ((A+P1) − A) | 7 | 2 | +0.030 | -0.088 to +0.066 |
| Balanced accuracy | 12 | Centile profiles beyond feature set A, downsampled full-reference charts ((A+P2) − A) | 6 | 1 | +0.030 | -0.088 to +0.066 |
| Balanced accuracy | 12 | Feature set A+P, downsampled cross-fitted − downsampled full-reference charts ((A+P1) − (A+P2)) | 0 | 0 | -0.001 | -0.010 to +0.006 |
| Balanced accuracy | 24 | Centile profiles beyond feature set A, downsampled cross-fitted charts ((A+P1) − A) | 3 | 3 | +0.018 | -0.069 to +0.032 |
| Balanced accuracy | 24 | Centile profiles beyond feature set A, downsampled full-reference charts ((A+P2) − A) | 3 | 3 | +0.023 | -0.069 to +0.043 |
| Balanced accuracy | 24 | Feature set A+P, downsampled cross-fitted − downsampled full-reference charts ((A+P1) − (A+P2)) | 0 | 0 | -0.005 | -0.014 to +0.004 |
| Balanced accuracy | 36 | Centile profiles beyond feature set A, downsampled cross-fitted charts ((A+P1) − A) | 0 | 3 | -0.002 | -0.134 to +0.016 |
| Balanced accuracy | 36 | Centile profiles beyond feature set A, downsampled full-reference charts ((A+P2) − A) | 0 | 3 | -0.002 | -0.130 to +0.018 |
| Balanced accuracy | 36 | Feature set A+P, downsampled cross-fitted − downsampled full-reference charts ((A+P1) − (A+P2)) | 0 | 0 | -0.002 | -0.019 to +0.007 |
| Macro-F1 | 12 | Centile profiles beyond feature set A, downsampled cross-fitted charts ((A+P1) − A) | 7 | 2 | +0.035 | -0.074 to +0.060 |
| Macro-F1 | 12 | Centile profiles beyond feature set A, downsampled full-reference charts ((A+P2) − A) | 7 | 2 | +0.038 | -0.075 to +0.059 |
| Macro-F1 | 12 | Feature set A+P, downsampled cross-fitted − downsampled full-reference charts ((A+P1) − (A+P2)) | 0 | 0 | -0.002 | -0.011 to +0.005 |
| Macro-F1 | 24 | Centile profiles beyond feature set A, downsampled cross-fitted charts ((A+P1) − A) | 3 | 2 | +0.022 | -0.071 to +0.035 |
| Macro-F1 | 24 | Centile profiles beyond feature set A, downsampled full-reference charts ((A+P2) − A) | 4 | 2 | +0.026 | -0.066 to +0.042 |
| Macro-F1 | 24 | Feature set A+P, downsampled cross-fitted − downsampled full-reference charts ((A+P1) − (A+P2)) | 0 | 0 | -0.004 | -0.010 to +0.002 |
| Macro-F1 | 36 | Centile profiles beyond feature set A, downsampled cross-fitted charts ((A+P1) − A) | 0 | 3 | -0.003 | -0.130 to +0.017 |
| Macro-F1 | 36 | Centile profiles beyond feature set A, downsampled full-reference charts ((A+P2) − A) | 0 | 3 | -0.001 | -0.128 to +0.019 |
| Macro-F1 | 36 | Feature set A+P, downsampled cross-fitted − downsampled full-reference charts ((A+P1) − (A+P2)) | 0 | 0 | -0.002 | -0.018 to +0.006 |

*Note:* feature sets defined in Table S6.

**Profile results.** Relative to downsampled full-reference charts, cross-fitting changed ridge AUC with centile profiles by −0.002, −0.003 and −0.001 at 12, 24 and 36 months. Balanced accuracy and macro-F1 did not differ significantly after Holm adjustment. Across 12 classifiers, median AUC differences were −0.002 at 12 months and −0.003 at 24 months. No classifier improved significantly; losses were significant in three classifiers at 12 months and one at 24 months, with maximum losses of 0.010 and 0.012. Centile profiles retained prognostic value under cross-fitting: ridge AUC increased by 0.029 and 0.017 over A at 12 and 24 months, and gains were significant in 9 and 8 of 12 classifiers after Holm adjustment.

Downsampled full-reference charts gave smaller centile increments than full-resolution full-reference charts (+0.031 versus +0.044 AUC at 12 months). Both reference samples included the controls, so this difference reflects resolution and refitting rather than reference membership. At 36 months, the ridge increment ranged from −0.001 to +0.001 across all methods, and no classifier showed a significant gain after Holm adjustment. Data S3 provides all metrics, reference methods and classifiers.

**Centile-map deep features under cross-fitting (F2 check).** We recomputed continuous cross-sectional and longitudinal maps using downsampled cross-fitted charts (G2m1) and downsampled full-reference charts (G2m2). Maps were encoded with the same 11 pretrained networks, eight augmented views and PCA within outer-training folds, then evaluated with all 12 classifiers. G2m1 and G2m2 combine A with centile-map deep features; H2m1 and H2m2 also include the corresponding centile profiles. A+P1 combines A with profiles from downsampled cross-fitted charts.

**Table S21.** Best classifier per setting for centile-map deep features: AUC, balanced accuracy and macro-F1 (95% CI).

| Setting | Horizon | Best classifier | AUC | Balanced accuracy | Macro-F1 |
| --- | --- | --- | --- | --- | --- |
| Feature set G2, full-resolution full-reference charts (main analysis) | 12 | GLM boosting | 0.954 (0.930–0.974) | 0.917 (0.894–0.937) | 0.897 (0.865–0.926) |
| Feature set G2, full-resolution full-reference charts (main analysis) | 24 | GLM boosting | 0.969 (0.948–0.986) | 0.931 (0.903–0.957) | 0.923 (0.889–0.952) |
| Feature set G2, full-resolution full-reference charts (main analysis) | 36 | Bayesian logistic | 0.875 (0.828–0.919) | 0.792 (0.750–0.834) | 0.791 (0.746–0.832) |
| Feature set G2m2, downsampled full-reference charts | 12 | XGBoost | 0.961 (0.940–0.978) | 0.911 (0.890–0.931) | 0.898 (0.868–0.925) |
| Feature set G2m2, downsampled full-reference charts | 24 | GLM boosting | 0.972 (0.953–0.987) | 0.929 (0.900–0.956) | 0.920 (0.884–0.951) |
| Feature set G2m2, downsampled full-reference charts | 36 | GLM boosting | 0.872 (0.821–0.918) | 0.831 (0.781–0.878) | 0.822 (0.766–0.874) |
| Feature set G2m1, downsampled cross-fitted charts | 12 | XGBoost | 0.963 (0.944–0.979) | 0.908 (0.887–0.927) | 0.897 (0.867–0.922) |
| Feature set G2m1, downsampled cross-fitted charts | 24 | GLM boosting | 0.973 (0.953–0.989) | 0.936 (0.908–0.960) | 0.927 (0.893–0.955) |
| Feature set G2m1, downsampled cross-fitted charts | 36 | Ridge | 0.870 (0.821–0.918) | 0.800 (0.753–0.846) | 0.797 (0.746–0.844) |

*Note:* feature sets defined in Table S6.

**Table S22.** F2 check across the 12 classifiers: number with Holm-significant gain (+) or loss (−), median and range of paired differences.

| Comparison | Metric | Horizon | + | − | Median | Range |
| --- | --- | --- | --- | --- | --- | --- |
| Feature set G2m1 vs feature set G2: downsampled cross-fitted vs full-resolution full-reference charts (G2m1 − G2) | AUC | 12 | 1 | 0 | -0.000 | -0.013 to +0.058 |
| Feature set G2m1 vs feature set G2: downsampled cross-fitted vs full-resolution full-reference charts (G2m1 − G2) | AUC | 24 | 0 | 0 | +0.001 | -0.015 to +0.016 |
| Feature set G2m1 vs feature set G2: downsampled cross-fitted vs full-resolution full-reference charts (G2m1 − G2) | AUC | 36 | 0 | 0 | -0.005 | -0.016 to +0.023 |
| Feature set G2m1 vs feature set G2: downsampled cross-fitted vs full-resolution full-reference charts (G2m1 − G2) | Balanced accuracy | 12 | 1 | 0 | +0.001 | -0.004 to +0.047 |
| Feature set G2m1 vs feature set G2: downsampled cross-fitted vs full-resolution full-reference charts (G2m1 − G2) | Balanced accuracy | 24 | 0 | 0 | +0.001 | -0.009 to +0.012 |
| Feature set G2m1 vs feature set G2: downsampled cross-fitted vs full-resolution full-reference charts (G2m1 − G2) | Balanced accuracy | 36 | 0 | 0 | -0.007 | -0.015 to +0.006 |
| Feature set G2m1 vs feature set G2: downsampled cross-fitted vs full-resolution full-reference charts (G2m1 − G2) | Macro-F1 | 12 | 1 | 0 | +0.002 | -0.004 to +0.049 |
| Feature set G2m1 vs feature set G2: downsampled cross-fitted vs full-resolution full-reference charts (G2m1 − G2) | Macro-F1 | 24 | 0 | 0 | +0.001 | -0.012 to +0.014 |
| Feature set G2m1 vs feature set G2: downsampled cross-fitted vs full-resolution full-reference charts (G2m1 − G2) | Macro-F1 | 36 | 0 | 0 | -0.008 | -0.016 to +0.006 |
| Feature set G2m1 vs feature set G2m2: downsampled cross-fitted vs downsampled full-reference charts (G2m1 − G2m2) | AUC | 12 | 0 | 1 | +0.002 | -0.006 to +0.008 |
| Feature set G2m1 vs feature set G2m2: downsampled cross-fitted vs downsampled full-reference charts (G2m1 − G2m2) | AUC | 24 | 0 | 2 | -0.003 | -0.013 to +0.002 |
| Feature set G2m1 vs feature set G2m2: downsampled cross-fitted vs downsampled full-reference charts (G2m1 − G2m2) | AUC | 36 | 0 | 0 | -0.001 | -0.011 to +0.030 |
| Feature set G2m1 vs feature set G2m2: downsampled cross-fitted vs downsampled full-reference charts (G2m1 − G2m2) | Balanced accuracy | 12 | 0 | 0 | -0.002 | -0.013 to +0.009 |
| Feature set G2m1 vs feature set G2m2: downsampled cross-fitted vs downsampled full-reference charts (G2m1 − G2m2) | Balanced accuracy | 24 | 0 | 0 | -0.002 | -0.008 to +0.010 |
| Feature set G2m1 vs feature set G2m2: downsampled cross-fitted vs downsampled full-reference charts (G2m1 − G2m2) | Balanced accuracy | 36 | 0 | 0 | -0.003 | -0.018 to +0.016 |
| Feature set G2m1 vs feature set G2m2: downsampled cross-fitted vs downsampled full-reference charts (G2m1 − G2m2) | Macro-F1 | 12 | 0 | 0 | -0.002 | -0.012 to +0.014 |
| Feature set G2m1 vs feature set G2m2: downsampled cross-fitted vs downsampled full-reference charts (G2m1 − G2m2) | Macro-F1 | 24 | 0 | 0 | -0.002 | -0.009 to +0.007 |
| Feature set G2m1 vs feature set G2m2: downsampled cross-fitted vs downsampled full-reference charts (G2m1 − G2m2) | Macro-F1 | 36 | 0 | 0 | -0.004 | -0.020 to +0.016 |
| Feature set G2m1 vs feature set G1: cross-fitted centile-map vs raw-map deep features (G2m1 − G1) | AUC | 12 | 11 | 0 | +0.043 | +0.020 to +0.294 |
| Feature set G2m1 vs feature set G1: cross-fitted centile-map vs raw-map deep features (G2m1 − G1) | AUC | 24 | 12 | 0 | +0.077 | +0.063 to +0.178 |
| Feature set G2m1 vs feature set G1: cross-fitted centile-map vs raw-map deep features (G2m1 − G1) | AUC | 36 | 0 | 0 | -0.012 | -0.020 to +0.004 |
| Feature set G2m1 vs feature set G1: cross-fitted centile-map vs raw-map deep features (G2m1 − G1) | Balanced accuracy | 12 | 11 | 0 | +0.066 | +0.020 to +0.217 |
| Feature set G2m1 vs feature set G1: cross-fitted centile-map vs raw-map deep features (G2m1 − G1) | Balanced accuracy | 24 | 12 | 0 | +0.103 | +0.059 to +0.160 |
| Feature set G2m1 vs feature set G1: cross-fitted centile-map vs raw-map deep features (G2m1 − G1) | Balanced accuracy | 36 | 0 | 0 | -0.001 | -0.013 to +0.020 |
| Feature set G2m1 vs feature set G1: cross-fitted centile-map vs raw-map deep features (G2m1 − G1) | Macro-F1 | 12 | 11 | 0 | +0.067 | +0.012 to +0.217 |
| Feature set G2m1 vs feature set G1: cross-fitted centile-map vs raw-map deep features (G2m1 − G1) | Macro-F1 | 24 | 12 | 0 | +0.106 | +0.074 to +0.161 |
| Feature set G2m1 vs feature set G1: cross-fitted centile-map vs raw-map deep features (G2m1 − G1) | Macro-F1 | 36 | 0 | 0 | -0.001 | -0.019 to +0.021 |
| Cross-fitted centile-map deep features beyond feature set A (G2m1 − A) | AUC | 12 | 9 | 2 | +0.055 | -0.045 to +0.071 |
| Cross-fitted centile-map deep features beyond feature set A (G2m1 − A) | AUC | 24 | 10 | 0 | +0.074 | +0.028 to +0.098 |
| Cross-fitted centile-map deep features beyond feature set A (G2m1 − A) | AUC | 36 | 0 | 2 | -0.003 | -0.082 to +0.011 |
| Cross-fitted centile-map deep features beyond feature set A (G2m1 − A) | Balanced accuracy | 12 | 9 | 2 | +0.082 | -0.068 to +0.132 |
| Cross-fitted centile-map deep features beyond feature set A (G2m1 − A) | Balanced accuracy | 24 | 9 | 0 | +0.094 | +0.006 to +0.112 |
| Cross-fitted centile-map deep features beyond feature set A (G2m1 − A) | Balanced accuracy | 36 | 0 | 3 | +0.000 | -0.079 to +0.033 |
| Cross-fitted centile-map deep features beyond feature set A (G2m1 − A) | Macro-F1 | 12 | 9 | 2 | +0.081 | -0.052 to +0.123 |
| Cross-fitted centile-map deep features beyond feature set A (G2m1 − A) | Macro-F1 | 24 | 9 | 0 | +0.096 | +0.009 to +0.110 |
| Cross-fitted centile-map deep features beyond feature set A (G2m1 − A) | Macro-F1 | 36 | 0 | 2 | -0.000 | -0.085 to +0.030 |
| Feature set G2m1 vs feature set A+P1: cross-fitted centile-map deep features vs cross-fitted centile profiles (G2m1 − (A+P1)) | AUC | 12 | 11 | 0 | +0.037 | -0.014 to +0.066 |
| Feature set G2m1 vs feature set A+P1: cross-fitted centile-map deep features vs cross-fitted centile profiles (G2m1 − (A+P1)) | AUC | 24 | 12 | 0 | +0.048 | +0.031 to +0.100 |
| Feature set G2m1 vs feature set A+P1: cross-fitted centile-map deep features vs cross-fitted centile profiles (G2m1 − (A+P1)) | AUC | 36 | 1 | 0 | +0.000 | -0.015 to +0.112 |
| Feature set G2m1 vs feature set A+P1: cross-fitted centile-map deep features vs cross-fitted centile profiles (G2m1 − (A+P1)) | Balanced accuracy | 12 | 11 | 1 | +0.050 | -0.038 to +0.084 |
| Feature set G2m1 vs feature set A+P1: cross-fitted centile-map deep features vs cross-fitted centile profiles (G2m1 − (A+P1)) | Balanced accuracy | 24 | 12 | 0 | +0.072 | +0.057 to +0.089 |
| Feature set G2m1 vs feature set A+P1: cross-fitted centile-map deep features vs cross-fitted centile profiles (G2m1 − (A+P1)) | Balanced accuracy | 36 | 1 | 0 | +0.008 | -0.003 to +0.069 |
| Feature set G2m1 vs feature set A+P1: cross-fitted centile-map deep features vs cross-fitted centile profiles (G2m1 − (A+P1)) | Macro-F1 | 12 | 9 | 0 | +0.045 | -0.028 to +0.077 |
| Feature set G2m1 vs feature set A+P1: cross-fitted centile-map deep features vs cross-fitted centile profiles (G2m1 − (A+P1)) | Macro-F1 | 24 | 12 | 0 | +0.073 | +0.064 to +0.080 |
| Feature set G2m1 vs feature set A+P1: cross-fitted centile-map deep features vs cross-fitted centile profiles (G2m1 − (A+P1)) | Macro-F1 | 36 | 1 | 0 | +0.005 | -0.004 to +0.070 |
| Cross-fitted centile profiles beyond feature set G2m1 (H2m1 − G2m1) | AUC | 12 | 0 | 1 | -0.000 | -0.020 to +0.018 |
| Cross-fitted centile profiles beyond feature set G2m1 (H2m1 − G2m1) | AUC | 24 | 0 | 0 | -0.002 | -0.022 to +0.004 |
| Cross-fitted centile profiles beyond feature set G2m1 (H2m1 − G2m1) | AUC | 36 | 0 | 0 | +0.007 | -0.012 to +0.030 |
| Cross-fitted centile profiles beyond feature set G2m1 (H2m1 − G2m1) | Balanced accuracy | 12 | 0 | 0 | +0.000 | -0.018 to +0.015 |
| Cross-fitted centile profiles beyond feature set G2m1 (H2m1 − G2m1) | Balanced accuracy | 24 | 0 | 0 | -0.000 | -0.022 to +0.006 |
| Cross-fitted centile profiles beyond feature set G2m1 (H2m1 − G2m1) | Balanced accuracy | 36 | 0 | 0 | +0.002 | -0.003 to +0.015 |
| Cross-fitted centile profiles beyond feature set G2m1 (H2m1 − G2m1) | Macro-F1 | 12 | 0 | 0 | +0.002 | -0.013 to +0.017 |
| Cross-fitted centile profiles beyond feature set G2m1 (H2m1 − G2m1) | Macro-F1 | 24 | 0 | 0 | +0.000 | -0.021 to +0.005 |
| Cross-fitted centile profiles beyond feature set G2m1 (H2m1 − G2m1) | Macro-F1 | 36 | 0 | 0 | +0.004 | -0.003 to +0.015 |

*Note:* feature sets defined in Table S6.

**Deep-feature results.** With downsampled cross-fitted charts, the best models reached AUC 0.963 at 12 months (XGBoost) and 0.973 at 24 months (GLM boosting), compared with 0.954 and 0.969 for full-resolution full-reference charts. Across the 12 classifiers, median cross-fitted-minus-full-resolution AUC differences were −0.000 and +0.001. No classifier showed a significant loss after Holm adjustment; LDA improved significantly at 12 months. The largest numerical losses were 0.013 and 0.015 for naïve Bayes, with both 95% confidence intervals including zero. Comparing the two downsampled methods gave median cross-fitted-minus-full-reference differences of +0.002 and −0.003. Losses were significant for HDDA at both horizons and LASSO at 24 months, with a maximum loss of 0.013.

Under cross-fitting, centile-map deep features outperformed raw-map deep features in 11 of 12 classifiers at 12 months and all 12 at 24 months after Holm adjustment (median ΔAUC +0.043 and +0.077). They also improved on A in 9 and 10 classifiers (median +0.055 and +0.074), although two classifiers showed significant losses at 12 months. Compared with cross-fitted centile profiles, gains were significant in 11 and 12 classifiers (median +0.037 and +0.048). Adding profiles to centile-map deep features changed AUC little (median −0.000 and −0.002; one significant loss). Balanced accuracy and macro-F1 showed similar results. No method consistently improved 36-month prediction. These comparisons suggest that including controls in chart fitting does not explain the prognostic advantage of centile-map deep features. Complete results are in Data S3, sheets F2check_perf and F2check_cmp.

### S10. SHAP atlas with detailed interpretation

Feature identifiers, definitions and directions are listed in Table S5.

**SHAP analysis.** For each outcome, we selected the five classifiers with complete results and the highest mean held-out AUC for A+P (Table S6). This selection was post hoc. Probability-scale SHAP values (Lundberg and Lee, 2017) were estimated by permutation with interventional masking, using 32 outer-training background observations and 128 forward/reverse permutation paths shared across classifiers. Reconstructed predictions from all 1,750 fits agreed with saved probabilities to within $2.7\times{10}^{-6}$. Additivity error was below $1.4\times{10}^{-10}$, and feature-importance correlations between permutation halves were ≥0.9993.

Each point represents one participant’s earliest included scan, with SHAP values averaged across held-out appearances. Sample sizes were 1,923 for concurrent classification and 1,043, 898 and 796 for 12-, 24- and 36-month prognosis. Colour denotes the feature value, clipped at its 5th and 95th percentiles. Positive SHAP values indicate contributions to a higher predicted case probability. Spearman correlation $\rho$ summarizes the association between feature values and SHAP values.

P and H denote cross-sectional and longitudinal centiles. GCS+ and GCS− are the positive and negative parts of the logit-transformed deficit rank (S3). GCS+ increases above the reference median; GCS− increases towards zero below the median and stays zero for ranks at or above it. Extent measures the weighted connected pixel count. SHAP describes model contributions in the presence of correlated predictors. It does not establish causality or added discrimination, which is evaluated separately in S7–S8.

#### S10.1 Concurrent classification (KLG ≥2 vs ≤1)

Centile features accounted for 60–78% of total mean absolute SHAP across the five classifiers and 66% for their average, exceeding their share at any prognostic horizon. Male sex was associated with lower predicted probability (ρ = −0.70). The leading imaging contributions included larger cross-sectional deficits in the medial central tibia (MCT · P extent, ρ ≈ 0.85) and tibial plate (Tibial · P GCS+, ρ ≈ 0.94), together with lower medial central femoral thickness (MCF thickness, ρ = −0.99). This medial distribution is consistent with cartilage loss in central weight-bearing regions associated with joint-space narrowing (Eckstein et al., 2010).

Greater medial posterior femoral and medial anterior tibial thickness also contributed to higher predicted probability (MPF and MAT thickness, ρ = 0.99). These associations were estimated alongside correlated centile features and may reflect suppression effects as well as the relative anterior and posterior thickening seen at higher grades (Figure A16). Larger lateral anterior femoral deficits (LAF · P extent) contributed positively, whereas MPT · P GCS+ and LCF · H GCS+ had negative associations in the linear models. These directions describe the fitted multivariable models, not isolated regional effects.

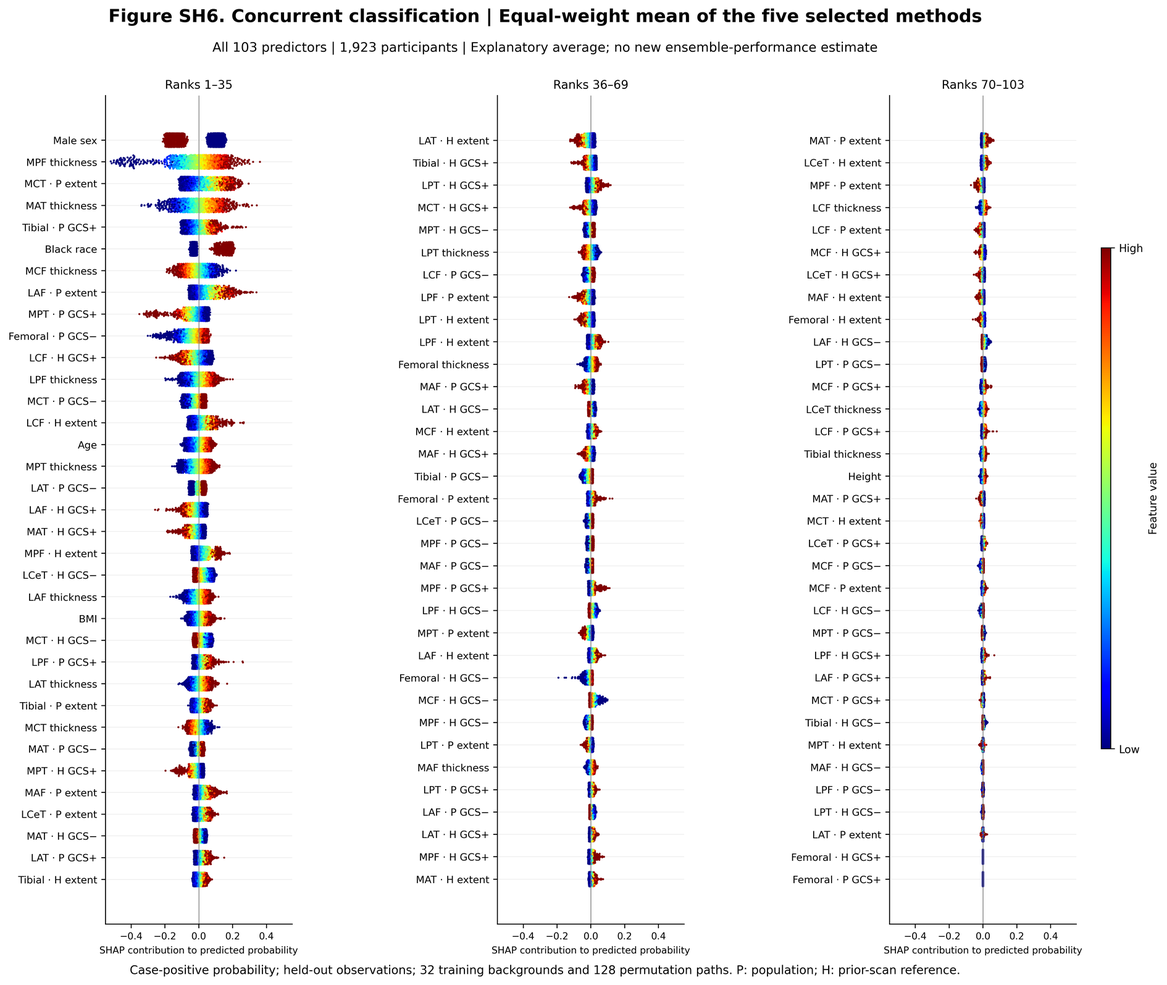

**Figure SH1.** Concurrent classification (KLG ≥2 vs ≤1) — Equal-weight average of the five classifiers. Probability-scale SHAP values for all predictors, ordered by mean absolute SHAP (n = 1,923 participants). Centile features account for 66% of the total mean absolute SHAP.

*Leading predictors (mean |SHAP|, direction)*

| Rank | Feature | Mean | SHAP |
| --- | --- | --- | --- |
| 1 | Male sex | 0.124 | men lower |
| 2 | MPF thickness | 0.090 | higher → ↑ risk (ρ = 0.99) |
| 3 | MCT · P extent | 0.078 | higher → ↑ risk (ρ = 0.85) |
| 4 | MAT thickness | 0.070 | higher → ↑ risk (ρ = 0.99) |
| 5 | Tibial · P GCS+ | 0.056 | higher → ↑ risk (ρ = 0.94) |
| 6 | Black race | 0.052 | Black race higher |
| 7 | MCF thickness | 0.047 | higher → ↓ risk (ρ = -0.99) |
| 8 | LAF · P extent | 0.046 | higher → ↑ risk (ρ = 0.88) |

*Leading centile predictors*

| Feature | Mean | SHAP |
| --- | --- | --- |
| MCT · P extent | 0.078 | higher → ↑ risk (ρ = 0.85) |
| Tibial · P GCS+ | 0.056 | higher → ↑ risk (ρ = 0.94) |
| LAF · P extent | 0.046 | higher → ↑ risk (ρ = 0.88) |
| MPT · P GCS+ | 0.046 | higher → ↓ risk (ρ = -0.85) |
| Femoral · P GCS− | 0.045 | higher → ↑ risk (ρ = 0.88) |

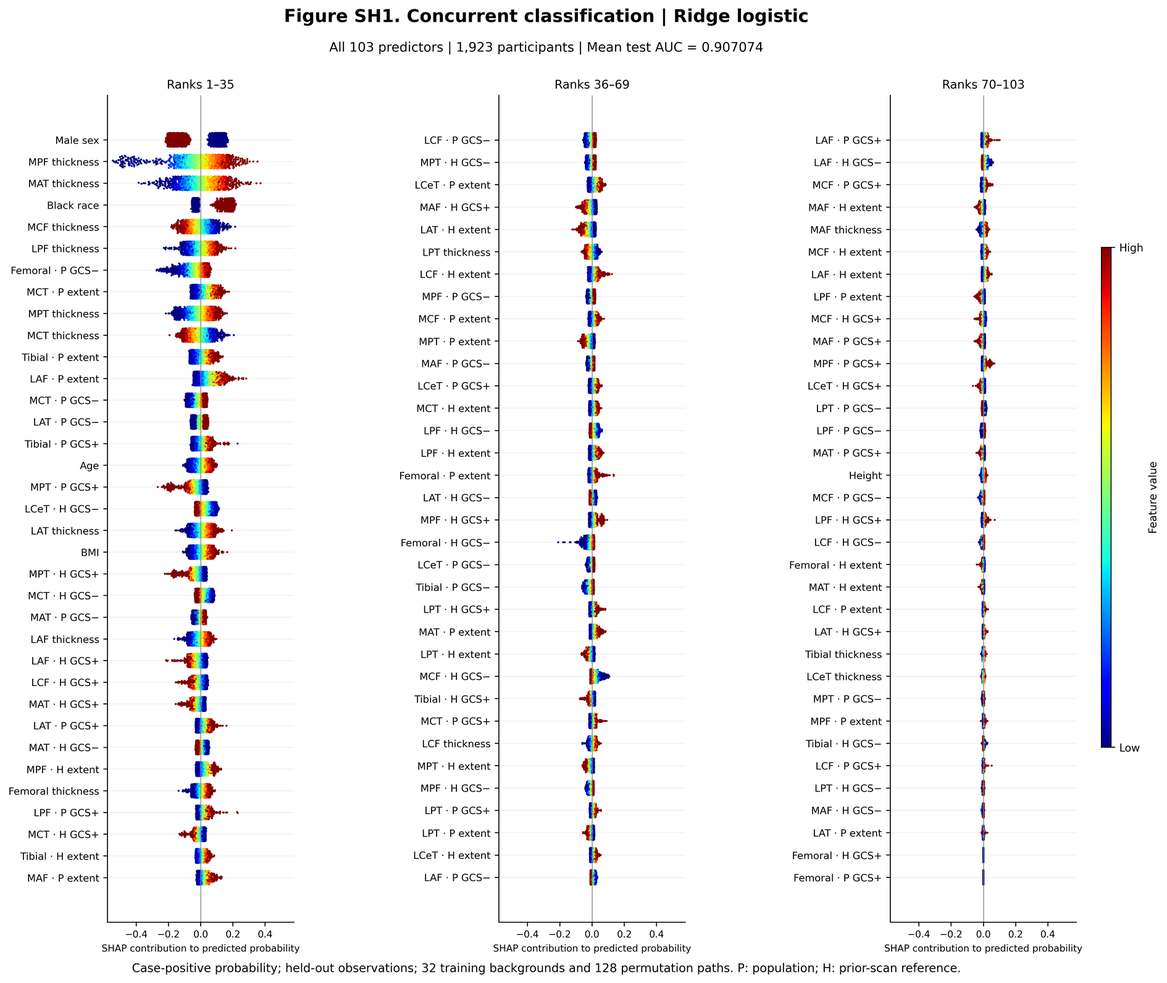

**Figure SH2.** Concurrent classification (KLG ≥2 vs ≤1) — Ridge. Probability-scale SHAP values for all predictors, ordered by mean absolute SHAP (n = 1,923 participants). Centile features account for 61% of the total mean absolute SHAP.

*Leading predictors (mean |SHAP|, direction)*

| Rank | Feature | Mean | SHAP |
| --- | --- | --- | --- |
| 1 | Male sex | 0.129 | men lower |
| 2 | MPF thickness | 0.092 | higher → ↑ risk (ρ = 0.99) |
| 3 | MAT thickness | 0.076 | higher → ↑ risk (ρ = 0.99) |
| 4 | Black race | 0.054 | Black race higher |
| 5 | MCF thickness | 0.046 | higher → ↓ risk (ρ = -0.99) |
| 6 | LPF thickness | 0.043 | higher → ↑ risk (ρ = 0.98) |
| 7 | Femoral · P GCS− | 0.042 | higher → ↑ risk (ρ = 0.89) |
| 8 | MCT · P extent | 0.042 | higher → ↑ risk (ρ = 0.85) |

*Leading centile predictors*

| Feature | Mean | SHAP |
| --- | --- | --- |
| Femoral · P GCS− | 0.042 | higher → ↑ risk (ρ = 0.89) |
| MCT · P extent | 0.042 | higher → ↑ risk (ρ = 0.85) |
| Tibial · P extent | 0.039 | higher → ↑ risk (ρ = 0.95) |
| LAF · P extent | 0.036 | higher → ↑ risk (ρ = 0.88) |
| MCT · P GCS− | 0.034 | higher → ↑ risk (ρ = 0.76) |

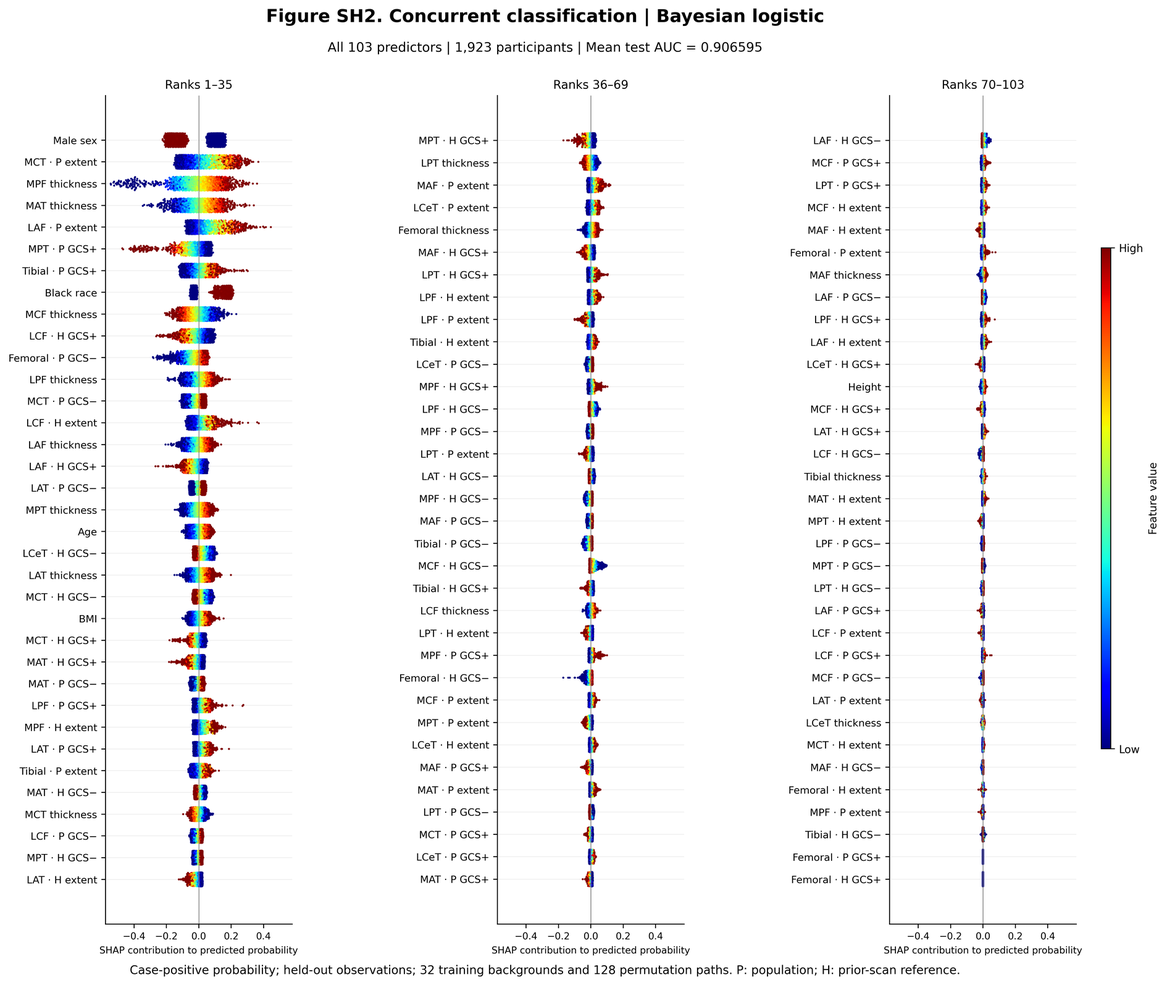

**Figure SH3.** Concurrent classification (KLG ≥2 vs ≤1) — Bayesian logistic. Probability-scale SHAP values for all predictors, ordered by mean absolute SHAP (n = 1,923 participants). Centile features account for 65% of the total mean absolute SHAP.

*Leading predictors (mean |SHAP|, direction)*

| Rank | Feature | Mean | SHAP |
| --- | --- | --- | --- |
| 1 | Male sex | 0.128 | men lower |
| 2 | MCT · P extent | 0.095 | higher → ↑ risk (ρ = 0.84) |
| 3 | MPF thickness | 0.091 | higher → ↑ risk (ρ = 0.99) |
| 4 | MAT thickness | 0.070 | higher → ↑ risk (ρ = 0.99) |
| 5 | LAF · P extent | 0.061 | higher → ↑ risk (ρ = 0.88) |
| 6 | MPT · P GCS+ | 0.060 | higher → ↓ risk (ρ = -0.84) |
| 7 | Tibial · P GCS+ | 0.059 | higher → ↑ risk (ρ = 0.93) |
| 8 | Black race | 0.052 | Black race higher |

*Leading centile predictors*

| Feature | Mean | SHAP |
| --- | --- | --- |
| MCT · P extent | 0.095 | higher → ↑ risk (ρ = 0.84) |
| LAF · P extent | 0.061 | higher → ↑ risk (ρ = 0.88) |
| MPT · P GCS+ | 0.060 | higher → ↓ risk (ρ = -0.84) |
| Tibial · P GCS+ | 0.059 | higher → ↑ risk (ρ = 0.93) |
| LCF · H GCS+ | 0.047 | higher → ↓ risk (ρ = -0.94) |

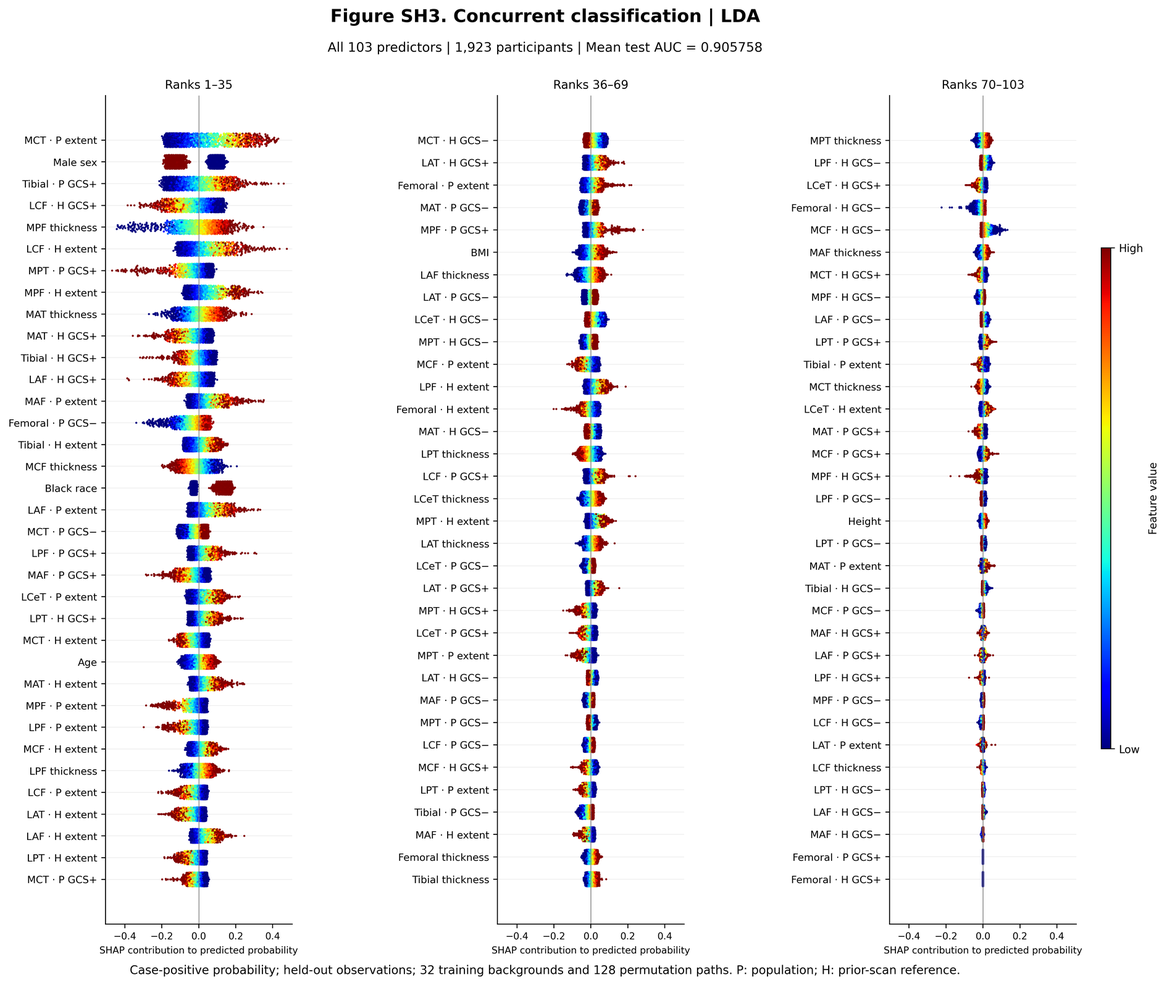

**Figure SH4.** Concurrent classification (KLG ≥2 vs ≤1) — LDA. Probability-scale SHAP values for all predictors, ordered by mean absolute SHAP (n = 1,923 participants). Centile features account for 78% of the total mean absolute SHAP.

*Leading predictors (mean |SHAP|, direction)*

| Rank | Feature | Mean | SHAP |
| --- | --- | --- | --- |
| 1 | MCT · P extent | 0.132 | higher → ↑ risk (ρ = 0.86) |
| 2 | Male sex | 0.111 | men lower |
| 3 | Tibial · P GCS+ | 0.108 | higher → ↑ risk (ρ = 0.95) |
| 4 | LCF · H GCS+ | 0.081 | higher → ↓ risk (ρ = -0.95) |
| 5 | MPF thickness | 0.080 | higher → ↑ risk (ρ = 0.99) |
| 6 | LCF · H extent | 0.075 | higher → ↑ risk (ρ = 0.93) |
| 7 | MPT · P GCS+ | 0.061 | higher → ↓ risk (ρ = -0.85) |
| 8 | MPF · H extent | 0.058 | higher → ↑ risk (ρ = 0.90) |

*Leading centile predictors*

| Feature | Mean | SHAP |
| --- | --- | --- |
| MCT · P extent | 0.132 | higher → ↑ risk (ρ = 0.86) |
| Tibial · P GCS+ | 0.108 | higher → ↑ risk (ρ = 0.95) |
| LCF · H GCS+ | 0.081 | higher → ↓ risk (ρ = -0.95) |
| LCF · H extent | 0.075 | higher → ↑ risk (ρ = 0.93) |
| MPT · P GCS+ | 0.061 | higher → ↓ risk (ρ = -0.85) |

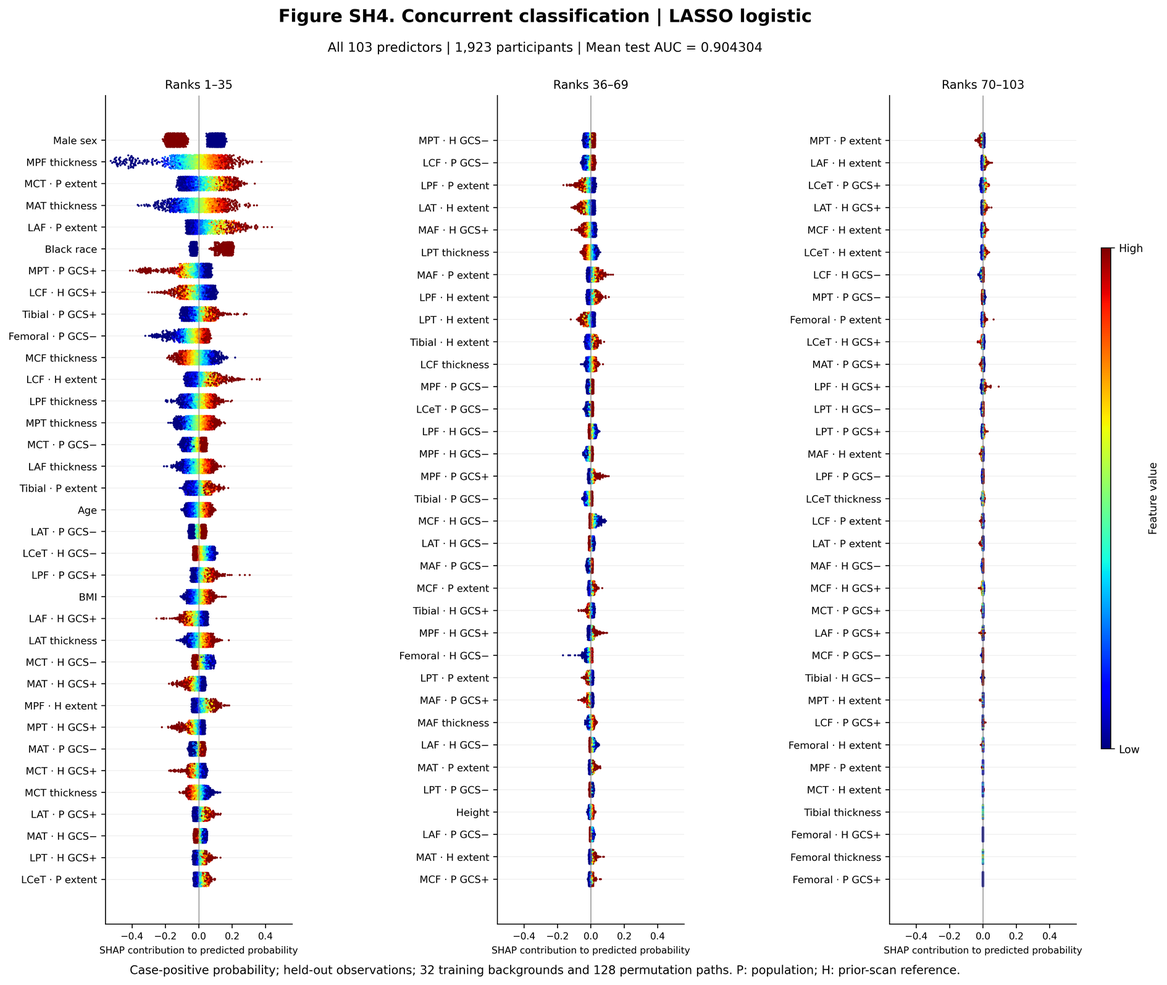

**Figure SH5.** Concurrent classification (KLG ≥2 vs ≤1) — LASSO. Probability-scale SHAP values for all predictors, ordered by mean absolute SHAP (n = 1,923 participants). Centile features account for 63% of the total mean absolute SHAP.

*Leading predictors (mean |SHAP|, direction)*

| Rank | Feature | Mean | SHAP |
| --- | --- | --- | --- |
| 1 | Male sex | 0.122 | men lower |
| 2 | MPF thickness | 0.091 | higher → ↑ risk (ρ = 0.99) |
| 3 | MCT · P extent | 0.079 | higher → ↑ risk (ρ = 0.84) |
| 4 | MAT thickness | 0.074 | higher → ↑ risk (ρ = 0.99) |
| 5 | LAF · P extent | 0.056 | higher → ↑ risk (ρ = 0.87) |
| 6 | Black race | 0.052 | Black race higher |
| 7 | MPT · P GCS+ | 0.051 | higher → ↓ risk (ρ = -0.85) |
| 8 | LCF · H GCS+ | 0.049 | higher → ↓ risk (ρ = -0.93) |

*Leading centile predictors*

| Feature | Mean | SHAP |
| --- | --- | --- |
| MCT · P extent | 0.079 | higher → ↑ risk (ρ = 0.84) |
| LAF · P extent | 0.056 | higher → ↑ risk (ρ = 0.87) |
| MPT · P GCS+ | 0.051 | higher → ↓ risk (ρ = -0.85) |
| LCF · H GCS+ | 0.049 | higher → ↓ risk (ρ = -0.93) |
| Tibial · P GCS+ | 0.048 | higher → ↑ risk (ρ = 0.92) |

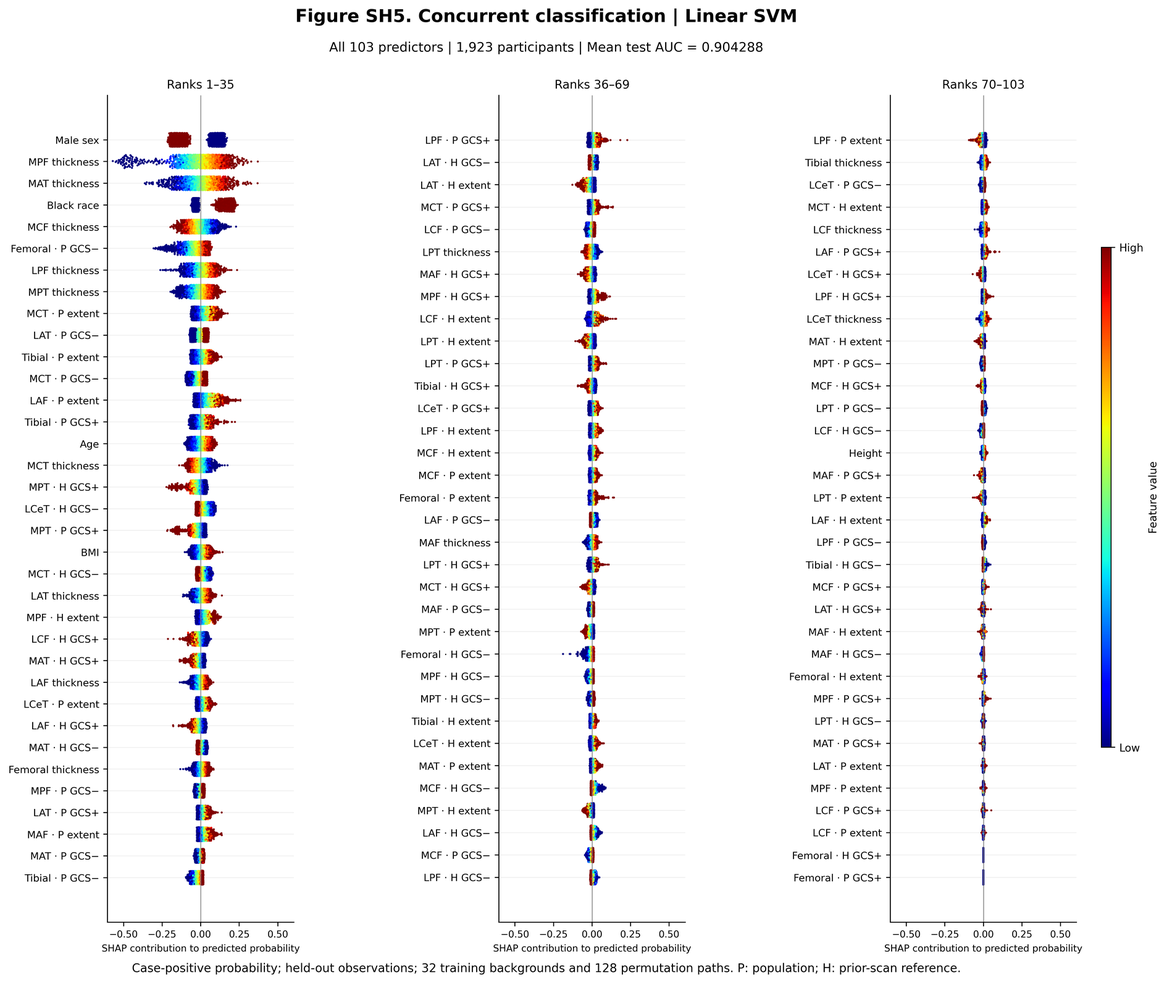

**Figure SH6.** Concurrent classification (KLG ≥2 vs ≤1) — Linear SVM. Probability-scale SHAP values for all predictors, ordered by mean absolute SHAP (n = 1,923 participants). Centile features account for 60% of the total mean absolute SHAP.

*Leading predictors (mean |SHAP|, direction)*

| Rank | Feature | Mean | SHAP |
| --- | --- | --- | --- |
| 1 | Male sex | 0.127 | men lower |
| 2 | MPF thickness | 0.095 | higher → ↑ risk (ρ = 0.99) |
| 3 | MAT thickness | 0.075 | higher → ↑ risk (ρ = 0.99) |
| 4 | Black race | 0.057 | Black race higher |
| 5 | MCF thickness | 0.048 | higher → ↓ risk (ρ = -0.99) |
| 6 | Femoral · P GCS− | 0.047 | higher → ↑ risk (ρ = 0.90) |
| 7 | LPF thickness | 0.044 | higher → ↑ risk (ρ = 0.98) |
| 8 | MPT thickness | 0.040 | higher → ↑ risk (ρ = 0.99) |

*Leading centile predictors*

| Feature | Mean | SHAP |
| --- | --- | --- |
| Femoral · P GCS− | 0.047 | higher → ↑ risk (ρ = 0.90) |
| MCT · P extent | 0.040 | higher → ↑ risk (ρ = 0.85) |
| LAT · P GCS− | 0.037 | higher → ↑ risk (ρ = 0.82) |
| Tibial · P extent | 0.037 | higher → ↑ risk (ρ = 0.95) |
| MCT · P GCS− | 0.035 | higher → ↑ risk (ρ = 0.78) |

#### S10.2 12-month prognosis

Index KLG had the largest contribution (mean |SHAP| 0.26 probability units), with KLG 1 associated with higher predicted probability. Other leading predictors were sex, age, BMI, physical activity and lateral posterior femoral thickness. Predicted probability was higher for women, older participants, higher BMI (age and BMI ρ ≈ 0.92) and greater LPF thickness (ρ ≈ 0.9). The LPF association was consistent with the thickening observed in progressors (main Figure 2C). Centile features accounted for 31–48% of total attribution.

The leading centile predictors were longitudinal: H GCS− in the femoral plate, lateral central tibia (LCeT), lateral posterior tibia (LPT) and medial and lateral central femur (ρ = 0.76–0.92), and H GCS+ in the medial anterior and medial central femur (ρ ≈ 0.6–0.7). For GCS−, values approaching zero from below contributed to higher predicted probability, indicating increasing deficit ranks relative to the knee’s earlier scans. In XGBoost, smaller cross-sectional LPF deficits also contributed to higher probability (LPF · P extent, ρ = −0.79), alongside the positive contribution of LPF mean thickness.

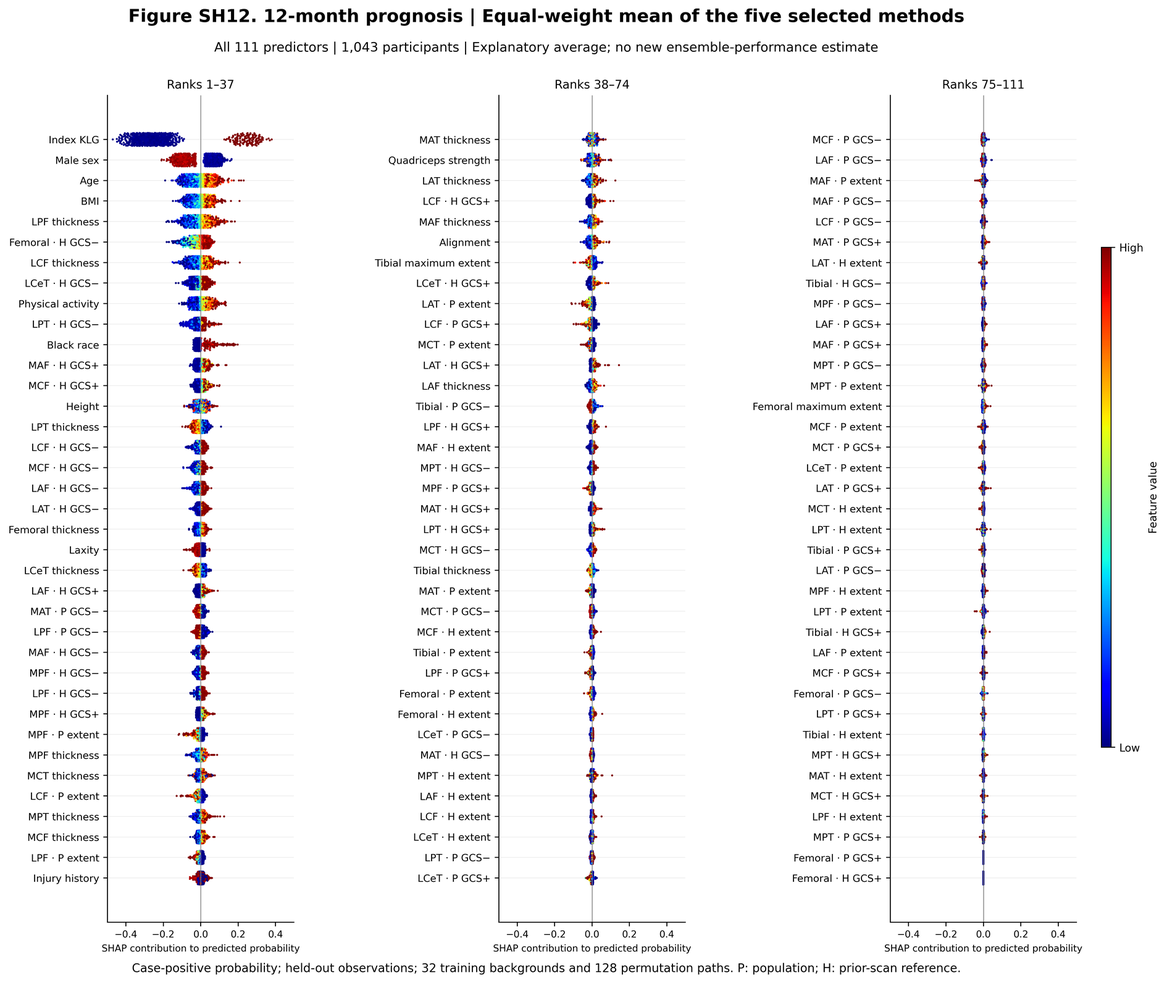

**Figure SH7.** 12-month prognosis — Equal-weight average of the five classifiers. Probability-scale SHAP values for all predictors, ordered by mean absolute SHAP (n = 1,043 participants). Centile features account for 38% of the total mean absolute SHAP.

*Leading predictors (mean |SHAP|, direction)*

| Rank | Feature | Mean | SHAP |
| --- | --- | --- | --- |
| 1 | Index KLG | 0.264 | higher → ↑ risk (ρ = 0.47) |
| 2 | Male sex | 0.073 | men lower |
| 3 | Age | 0.044 | higher → ↑ risk (ρ = 0.92) |
| 4 | BMI | 0.034 | higher → ↑ risk (ρ = 0.93) |
| 5 | LPF thickness | 0.034 | higher → ↑ risk (ρ = 0.94) |
| 6 | Femoral · H GCS− | 0.028 | higher → ↑ risk (ρ = 0.92) |
| 7 | LCF thickness | 0.026 | higher → ↑ risk (ρ = 0.92) |
| 8 | LCeT · H GCS− | 0.026 | higher → ↑ risk (ρ = 0.88) |

*Leading centile predictors*

| Feature | Mean | SHAP |
| --- | --- | --- |
| Femoral · H GCS− | 0.028 | higher → ↑ risk (ρ = 0.92) |
| LCeT · H GCS− | 0.026 | higher → ↑ risk (ρ = 0.88) |
| LPT · H GCS− | 0.018 | higher → ↑ risk (ρ = 0.82) |
| MAF · H GCS+ | 0.015 | higher → ↑ risk (ρ = 0.60) |
| MCF · H GCS+ | 0.014 | higher → ↑ risk (ρ = 0.66) |

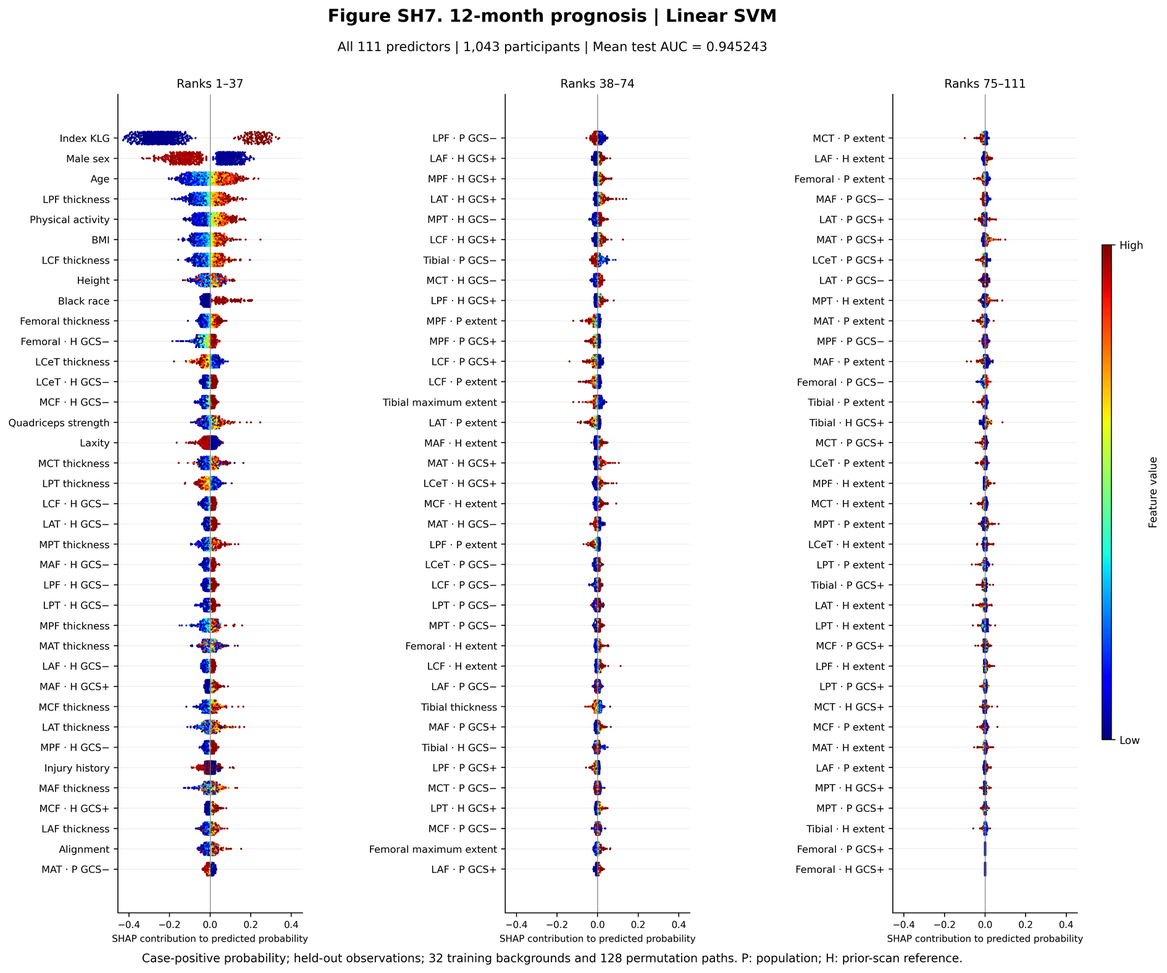

**Figure SH8.** 12-month prognosis — Linear SVM. Probability-scale SHAP values for all predictors, ordered by mean absolute SHAP (n = 1,043 participants). Centile features account for 38% of the total mean absolute SHAP.

*Leading predictors (mean |SHAP|, direction)*

| Rank | Feature | Mean | SHAP |
| --- | --- | --- | --- |
| 1 | Index KLG | 0.246 | higher → ↑ risk (ρ = 0.45) |
| 2 | Male sex | 0.107 | men lower |
| 3 | Age | 0.056 | higher → ↑ risk (ρ = 0.92) |
| 4 | LPF thickness | 0.036 | higher → ↑ risk (ρ = 0.91) |
| 5 | Physical activity | 0.034 | higher → ↑ risk (ρ = 0.93) |
| 6 | BMI | 0.030 | higher → ↑ risk (ρ = 0.92) |
| 7 | LCF thickness | 0.028 | higher → ↑ risk (ρ = 0.92) |
| 8 | Height | 0.020 | non-monotone (ρ = 0.29) |

*Leading centile predictors*

| Feature | Mean | SHAP |
| --- | --- | --- |
| Femoral · H GCS− | 0.017 | higher → ↑ risk (ρ = 0.92) |
| LCeT · H GCS− | 0.016 | higher → ↑ risk (ρ = 0.89) |
| MCF · H GCS− | 0.014 | higher → ↑ risk (ρ = 0.88) |
| LCF · H GCS− | 0.013 | higher → ↑ risk (ρ = 0.87) |
| LAT · H GCS− | 0.013 | higher → ↑ risk (ρ = 0.81) |

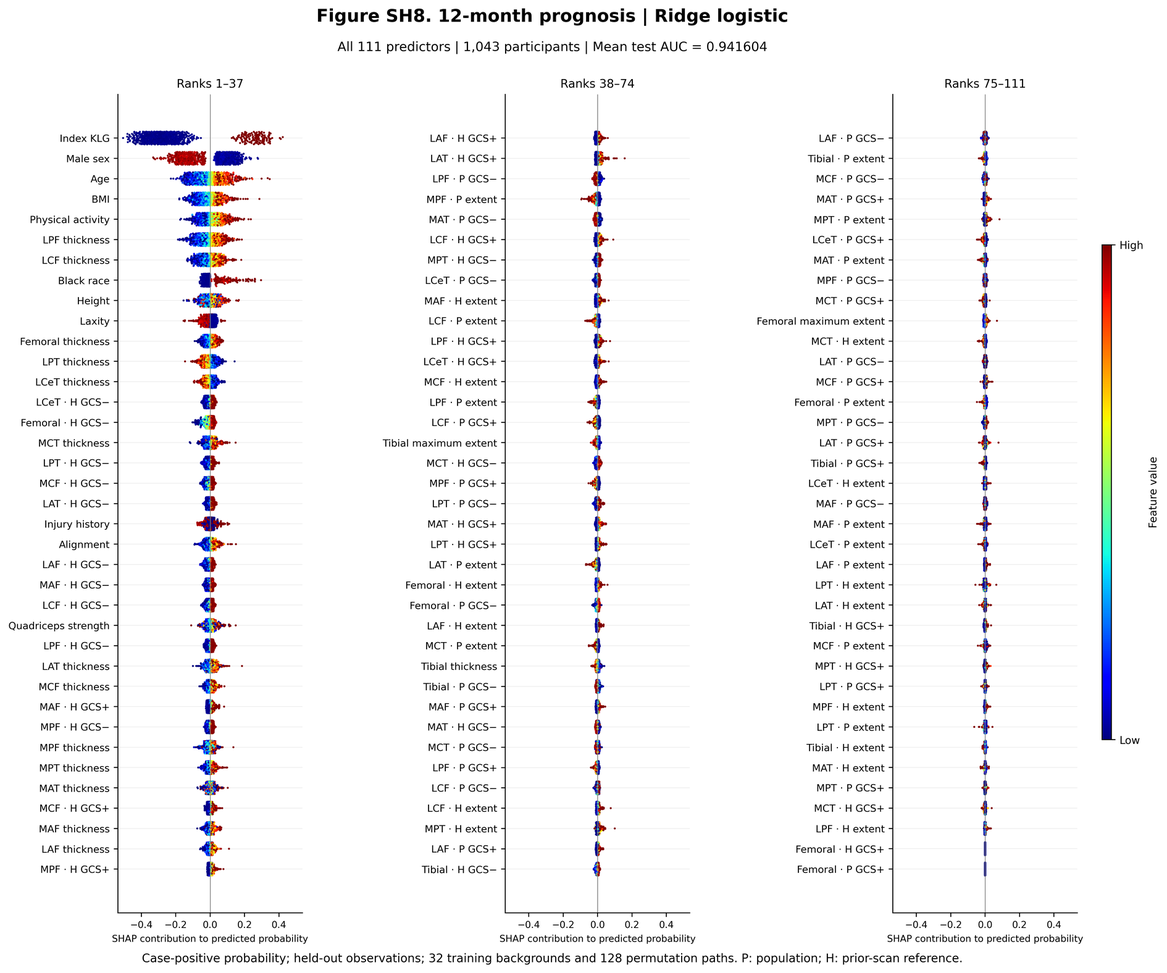

**Figure SH9.** 12-month prognosis — Ridge. Probability-scale SHAP values for all predictors, ordered by mean absolute SHAP (n = 1,043 participants). Centile features account for 31% of the total mean absolute SHAP.

*Leading predictors (mean |SHAP|, direction)*

| Rank | Feature | Mean | SHAP |
| --- | --- | --- | --- |
| 1 | Index KLG | 0.279 | higher → ↑ risk (ρ = 0.47) |
| 2 | Male sex | 0.113 | men lower |
| 3 | Age | 0.061 | higher → ↑ risk (ρ = 0.92) |
| 4 | BMI | 0.041 | higher → ↑ risk (ρ = 0.94) |
| 5 | Physical activity | 0.038 | higher → ↑ risk (ρ = 0.92) |
| 6 | LPF thickness | 0.037 | higher → ↑ risk (ρ = 0.93) |
| 7 | LCF thickness | 0.030 | higher → ↑ risk (ρ = 0.92) |
| 8 | Black race | 0.026 | Black race higher |

*Leading centile predictors*

| Feature | Mean | SHAP |
| --- | --- | --- |
| LCeT · H GCS− | 0.015 | higher → ↑ risk (ρ = 0.88) |
| Femoral · H GCS− | 0.015 | higher → ↑ risk (ρ = 0.89) |
| LPT · H GCS− | 0.012 | higher → ↑ risk (ρ = 0.84) |
| MCF · H GCS− | 0.012 | higher → ↑ risk (ρ = 0.87) |
| LAT · H GCS− | 0.012 | higher → ↑ risk (ρ = 0.80) |

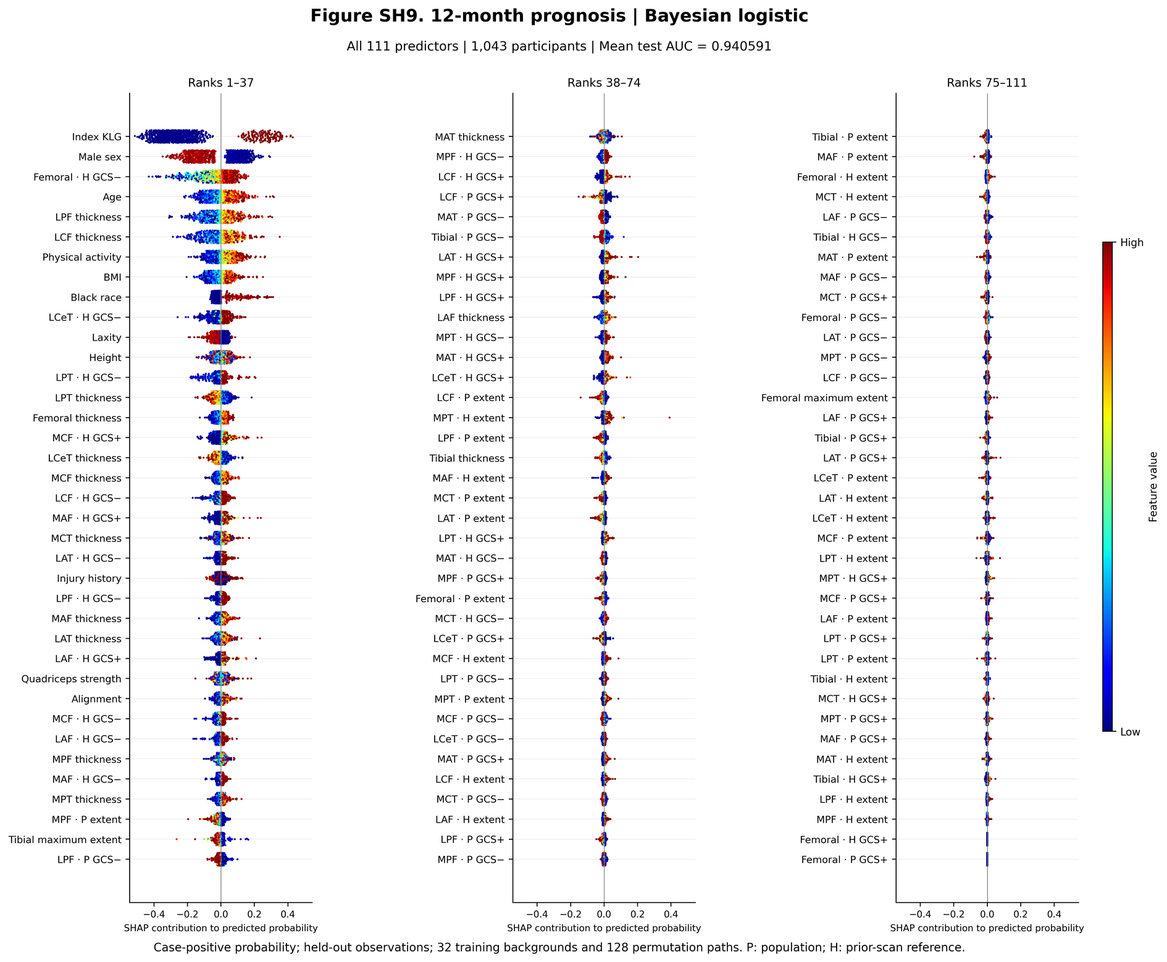

**Figure SH10.** 12-month prognosis — Bayesian logistic. Probability-scale SHAP values for all predictors, ordered by mean absolute SHAP (n = 1,043 participants). Centile features account for 36% of the total mean absolute SHAP.

*Leading predictors (mean |SHAP|, direction)*

| Rank | Feature | Mean | SHAP |
| --- | --- | --- | --- |
| 1 | Index KLG | 0.274 | higher → ↑ risk (ρ = 0.46) |
| 2 | Male sex | 0.120 | men lower |
| 3 | Femoral · H GCS− | 0.056 | higher → ↑ risk (ρ = 0.88) |
| 4 | Age | 0.053 | higher → ↑ risk (ρ = 0.91) |
| 5 | LPF thickness | 0.045 | higher → ↑ risk (ρ = 0.90) |
| 6 | LCF thickness | 0.044 | higher → ↑ risk (ρ = 0.89) |
| 7 | Physical activity | 0.038 | higher → ↑ risk (ρ = 0.92) |
| 8 | BMI | 0.035 | higher → ↑ risk (ρ = 0.93) |

*Leading centile predictors*

| Feature | Mean | SHAP |
| --- | --- | --- |
| Femoral · H GCS− | 0.056 | higher → ↑ risk (ρ = 0.88) |
| LCeT · H GCS− | 0.025 | higher → ↑ risk (ρ = 0.86) |
| LPT · H GCS− | 0.020 | higher → ↑ risk (ρ = 0.80) |
| MCF · H GCS+ | 0.018 | higher → ↑ risk (ρ = 0.66) |
| LCF · H GCS− | 0.015 | higher → ↑ risk (ρ = 0.84) |

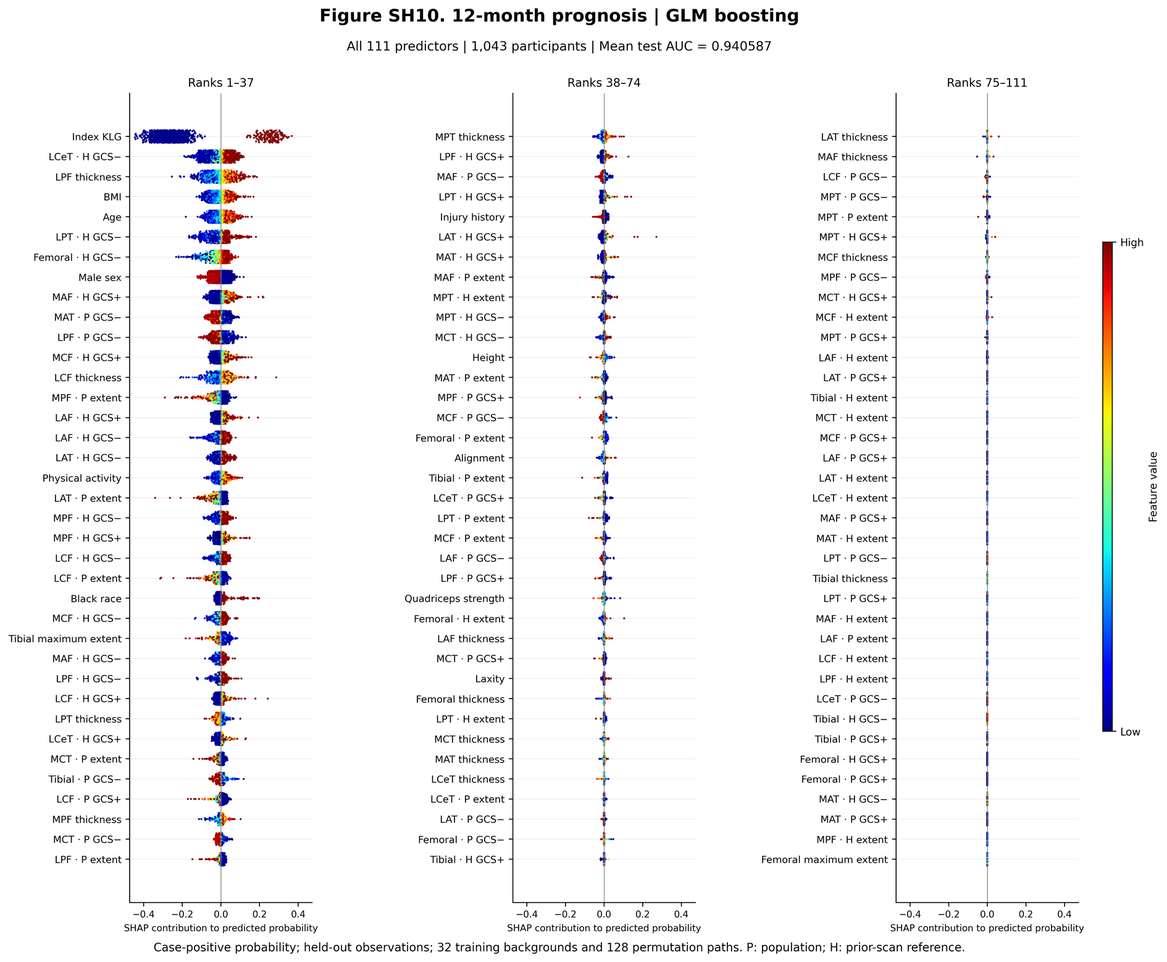

**Figure SH11.** 12-month prognosis — GLM boosting. Probability-scale SHAP values for all predictors, ordered by mean absolute SHAP (n = 1,043 participants). Centile features account for 46% of the total mean absolute SHAP.

*Leading predictors (mean |SHAP|, direction)*

| Rank | Feature | Mean | SHAP |
| --- | --- | --- | --- |
| 1 | Index KLG | 0.267 | higher → ↑ risk (ρ = 0.46) |
| 2 | LCeT · H GCS− | 0.048 | higher → ↑ risk (ρ = 0.88) |
| 3 | LPF thickness | 0.034 | higher → ↑ risk (ρ = 0.90) |
| 4 | BMI | 0.029 | higher → ↑ risk (ρ = 0.93) |
| 5 | Age | 0.026 | higher → ↑ risk (ρ = 0.87) |
| 6 | LPT · H GCS− | 0.026 | higher → ↑ risk (ρ = 0.76) |
| 7 | Femoral · H GCS− | 0.025 | higher → ↑ risk (ρ = 0.87) |
| 8 | Male sex | 0.024 | men lower |

*Leading centile predictors*

| Feature | Mean | SHAP |
| --- | --- | --- |
| LCeT · H GCS− | 0.048 | higher → ↑ risk (ρ = 0.88) |
| LPT · H GCS− | 0.026 | higher → ↑ risk (ρ = 0.76) |
| Femoral · H GCS− | 0.025 | higher → ↑ risk (ρ = 0.87) |
| MAF · H GCS+ | 0.023 | higher → ↑ risk (ρ = 0.64) |
| MAT · P GCS− | 0.018 | higher → ↓ risk (ρ = -0.75) |

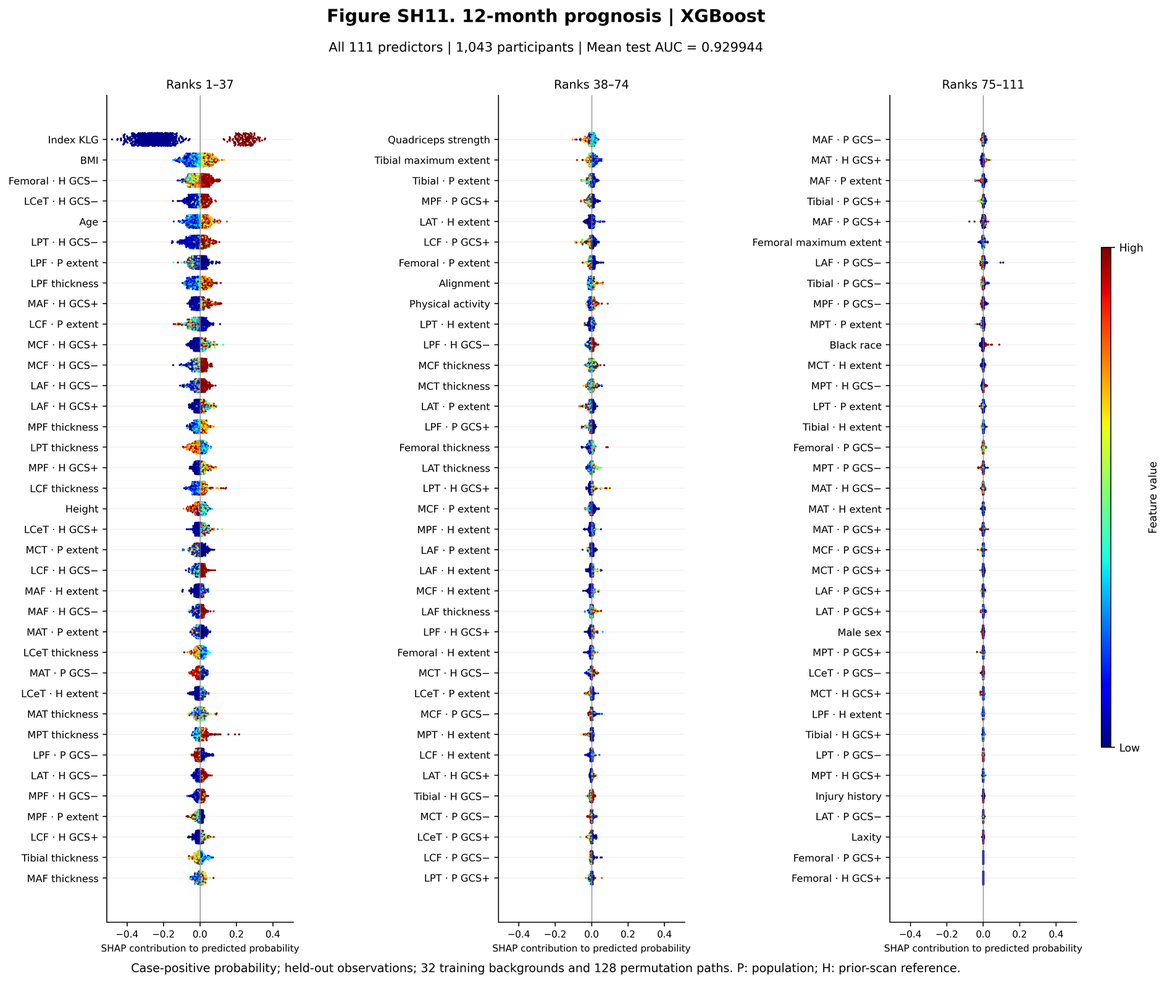

**Figure SH12.** 12-month prognosis — XGBoost. Probability-scale SHAP values for all predictors, ordered by mean absolute SHAP (n = 1,043 participants). Centile features account for 48% of the total mean absolute SHAP.

*Leading predictors (mean |SHAP|, direction)*

| Rank | Feature | Mean | SHAP |
| --- | --- | --- | --- |
| 1 | Index KLG | 0.252 | higher → ↑ risk (ρ = 0.47) |
| 2 | BMI | 0.037 | higher → ↑ risk (ρ = 0.83) |
| 3 | Femoral · H GCS− | 0.028 | higher → ↑ risk (ρ = 0.82) |
| 4 | LCeT · H GCS− | 0.025 | higher → ↑ risk (ρ = 0.85) |
| 5 | Age | 0.024 | higher → ↑ risk (ρ = 0.80) |
| 6 | LPT · H GCS− | 0.022 | higher → ↑ risk (ρ = 0.77) |
| 7 | LPF · P extent | 0.019 | higher → ↓ risk (ρ = -0.79) |
| 8 | LPF thickness | 0.019 | higher → ↑ risk (ρ = 0.79) |

*Leading centile predictors*

| Feature | Mean | SHAP |
| --- | --- | --- |
| Femoral · H GCS− | 0.028 | higher → ↑ risk (ρ = 0.82) |
| LCeT · H GCS− | 0.025 | higher → ↑ risk (ρ = 0.85) |
| LPT · H GCS− | 0.022 | higher → ↑ risk (ρ = 0.77) |
| LPF · P extent | 0.019 | higher → ↓ risk (ρ = -0.79) |
| MAF · H GCS+ | 0.018 | higher → ↑ risk (ρ = 0.46) |

#### S10.3 24-month prognosis

Index KLG and sex remained the leading predictors. Centile features accounted for 37–53% of total attribution (45% for the average), their largest share among the prognostic horizons. The longitudinal lateral central tibial feature LCeT · H GCS− (ρ ≈ 0.86) was the strongest imaging predictor in four of five classifiers. Longitudinal GCS+ features in the lateral posterior, medial posterior, medial central and medial anterior femur also contributed positively (ρ ≈ 0.5–0.6); these features measure deficit ranks above the reference median. Greater laxity contributed to higher probability in Bayesian logistic regression and SVM, as did lower height in tree models. LPF · P extent had a negative association in GLM boosting (ρ = −0.67). The same lateral tibial and femoral longitudinal features ranked highly across SVM, Bayesian logistic regression, GLM boosting, XGBoost and LightGBM.

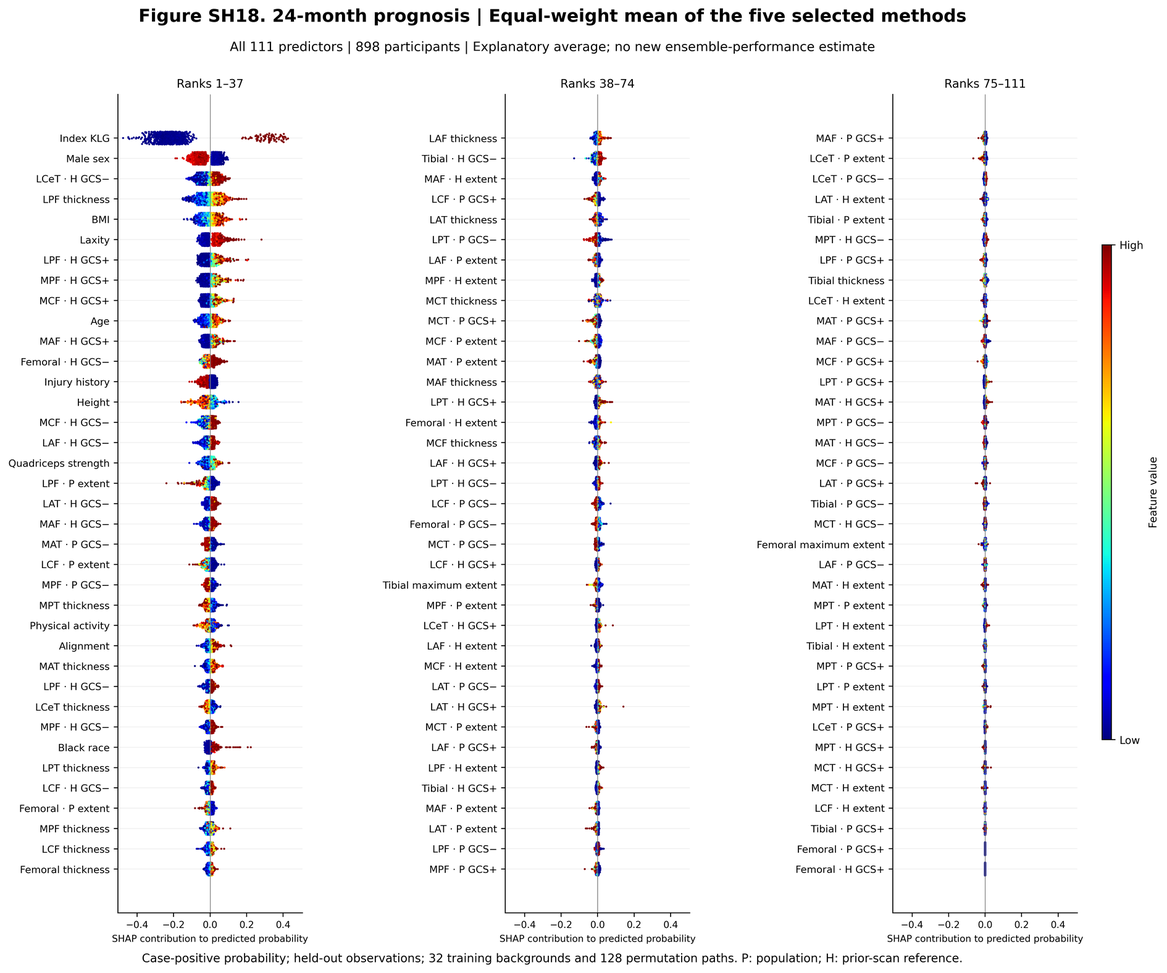

**Figure SH13.** 24-month prognosis — Equal-weight average of the five classifiers. Probability-scale SHAP values for all predictors, ordered by mean absolute SHAP (n = 898 participants). Centile features account for 45% of the total mean absolute SHAP.

*Leading predictors (mean |SHAP|, direction)*

| Rank | Feature | Mean | SHAP |
| --- | --- | --- | --- |
| 1 | Index KLG | 0.236 | higher → ↑ risk (ρ = 0.42) |
| 2 | Male sex | 0.045 | men lower |
| 3 | LCeT · H GCS− | 0.034 | higher → ↑ risk (ρ = 0.86) |
| 4 | LPF thickness | 0.033 | higher → ↑ risk (ρ = 0.90) |
| 5 | BMI | 0.030 | higher → ↑ risk (ρ = 0.86) |
| 6 | Laxity | 0.029 | higher → ↑ risk (ρ = 0.45) |
| 7 | LPF · H GCS+ | 0.025 | higher → ↑ risk (ρ = 0.59) |
| 8 | MPF · H GCS+ | 0.024 | higher → ↑ risk (ρ = 0.52) |

*Leading centile predictors*

| Feature | Mean | SHAP |
| --- | --- | --- |
| LCeT · H GCS− | 0.034 | higher → ↑ risk (ρ = 0.86) |
| LPF · H GCS+ | 0.025 | higher → ↑ risk (ρ = 0.59) |
| MPF · H GCS+ | 0.024 | higher → ↑ risk (ρ = 0.52) |
| MCF · H GCS+ | 0.022 | higher → ↑ risk (ρ = 0.61) |
| MAF · H GCS+ | 0.020 | higher → ↑ risk (ρ = 0.56) |

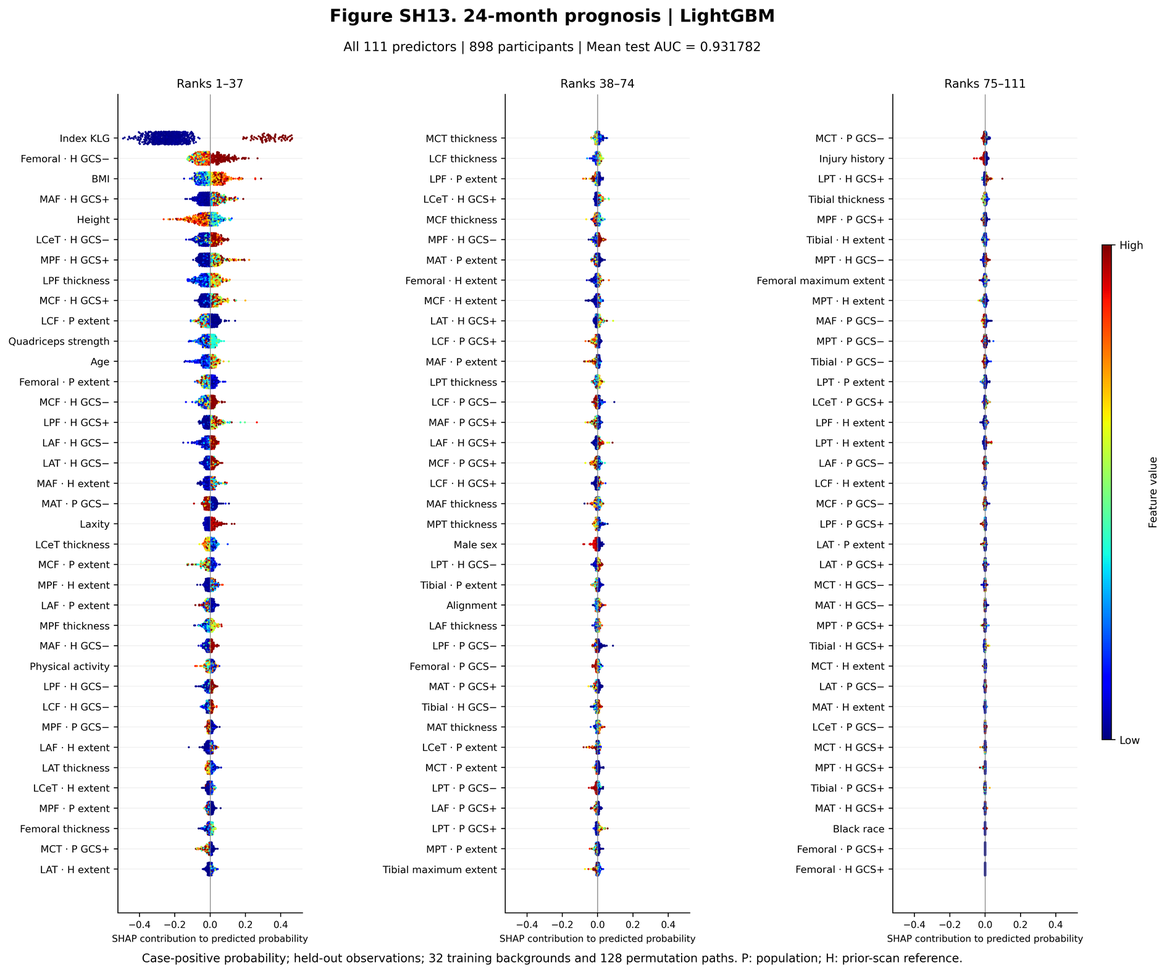

**Figure SH14.** 24-month prognosis — LightGBM. Probability-scale SHAP values for all predictors, ordered by mean absolute SHAP (n = 898 participants). Centile features account for 51% of the total mean absolute SHAP.

*Leading predictors (mean |SHAP|, direction)*

| Rank | Feature | Mean | SHAP |
| --- | --- | --- | --- |
| 1 | Index KLG | 0.248 | higher → ↑ risk (ρ = 0.40) |
| 2 | Femoral · H GCS− | 0.043 | higher → ↑ risk (ρ = 0.50) |
| 3 | BMI | 0.035 | higher → ↑ risk (ρ = 0.72) |
| 4 | MAF · H GCS+ | 0.028 | higher → ↑ risk (ρ = 0.50) |
| 5 | Height | 0.027 | higher → ↓ risk (ρ = -0.52) |
| 6 | LCeT · H GCS− | 0.026 | higher → ↑ risk (ρ = 0.81) |
| 7 | MPF · H GCS+ | 0.025 | higher → ↑ risk (ρ = 0.47) |
| 8 | LPF thickness | 0.023 | higher → ↑ risk (ρ = 0.77) |

*Leading centile predictors*

| Feature | Mean | SHAP |
| --- | --- | --- |
| Femoral · H GCS− | 0.043 | higher → ↑ risk (ρ = 0.50) |
| MAF · H GCS+ | 0.028 | higher → ↑ risk (ρ = 0.50) |
| LCeT · H GCS− | 0.026 | higher → ↑ risk (ρ = 0.81) |
| MPF · H GCS+ | 0.025 | higher → ↑ risk (ρ = 0.47) |
| MCF · H GCS+ | 0.022 | higher → ↑ risk (ρ = 0.58) |

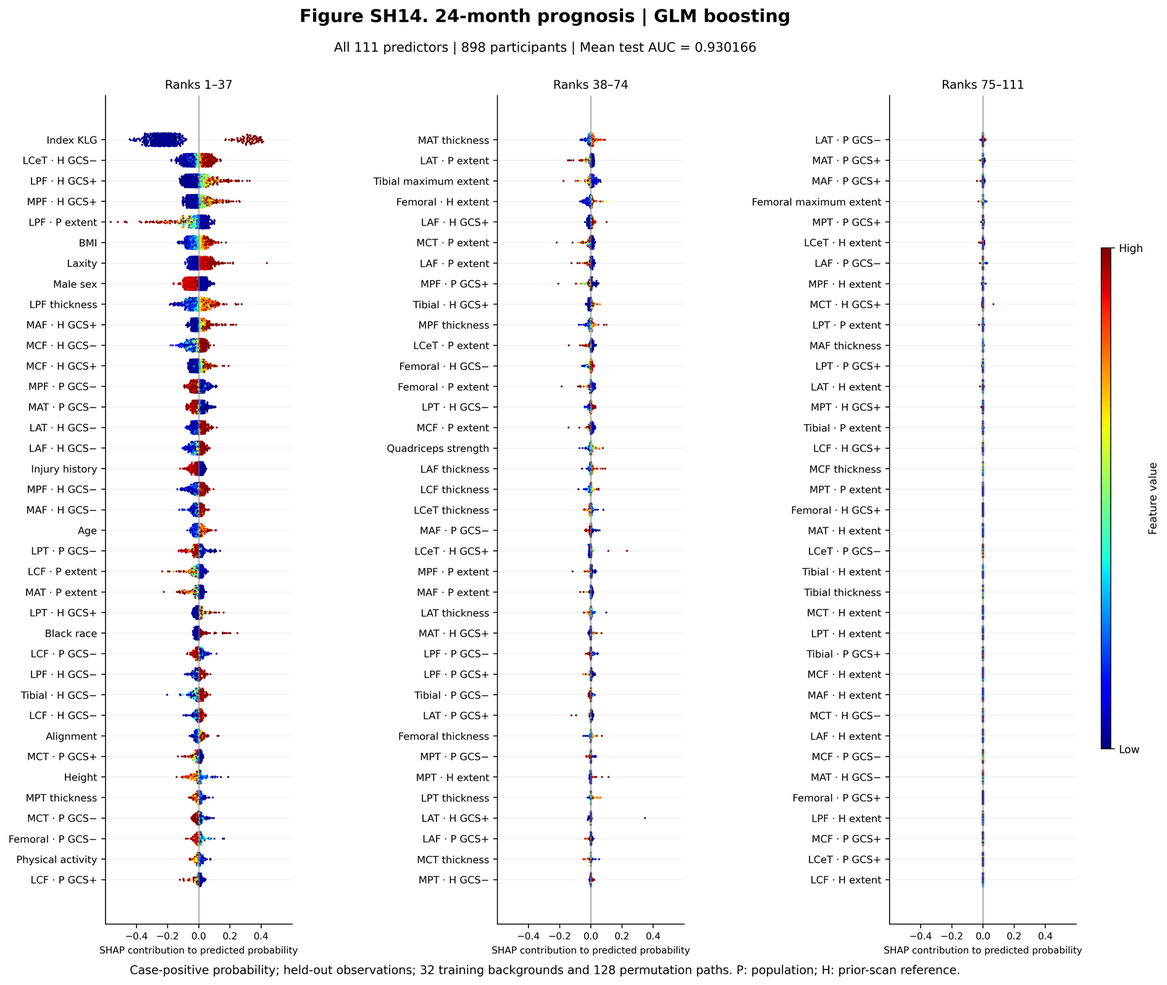

**Figure SH15.** 24-month prognosis — GLM boosting. Probability-scale SHAP values for all predictors, ordered by mean absolute SHAP (n = 898 participants). Centile features account for 51% of the total mean absolute SHAP.

*Leading predictors (mean |SHAP|, direction)*

| Rank | Feature | Mean | SHAP |
| --- | --- | --- | --- |
| 1 | Index KLG | 0.238 | higher → ↑ risk (ρ = 0.42) |
| 2 | LCeT · H GCS− | 0.048 | higher → ↑ risk (ρ = 0.88) |
| 3 | LPF · H GCS+ | 0.044 | higher → ↑ risk (ρ = 0.62) |
| 4 | MPF · H GCS+ | 0.037 | higher → ↑ risk (ρ = 0.51) |
| 5 | LPF · P extent | 0.037 | higher → ↓ risk (ρ = -0.67) |
| 6 | BMI | 0.032 | higher → ↑ risk (ρ = 0.92) |
| 7 | Laxity | 0.031 | higher → ↑ risk (ρ = 0.42) |
| 8 | Male sex | 0.031 | men lower |

*Leading centile predictors*

| Feature | Mean | SHAP |
| --- | --- | --- |
| LCeT · H GCS− | 0.048 | higher → ↑ risk (ρ = 0.88) |
| LPF · H GCS+ | 0.044 | higher → ↑ risk (ρ = 0.62) |
| MPF · H GCS+ | 0.037 | higher → ↑ risk (ρ = 0.51) |
| LPF · P extent | 0.037 | higher → ↓ risk (ρ = -0.67) |
| MAF · H GCS+ | 0.022 | higher → ↑ risk (ρ = 0.62) |

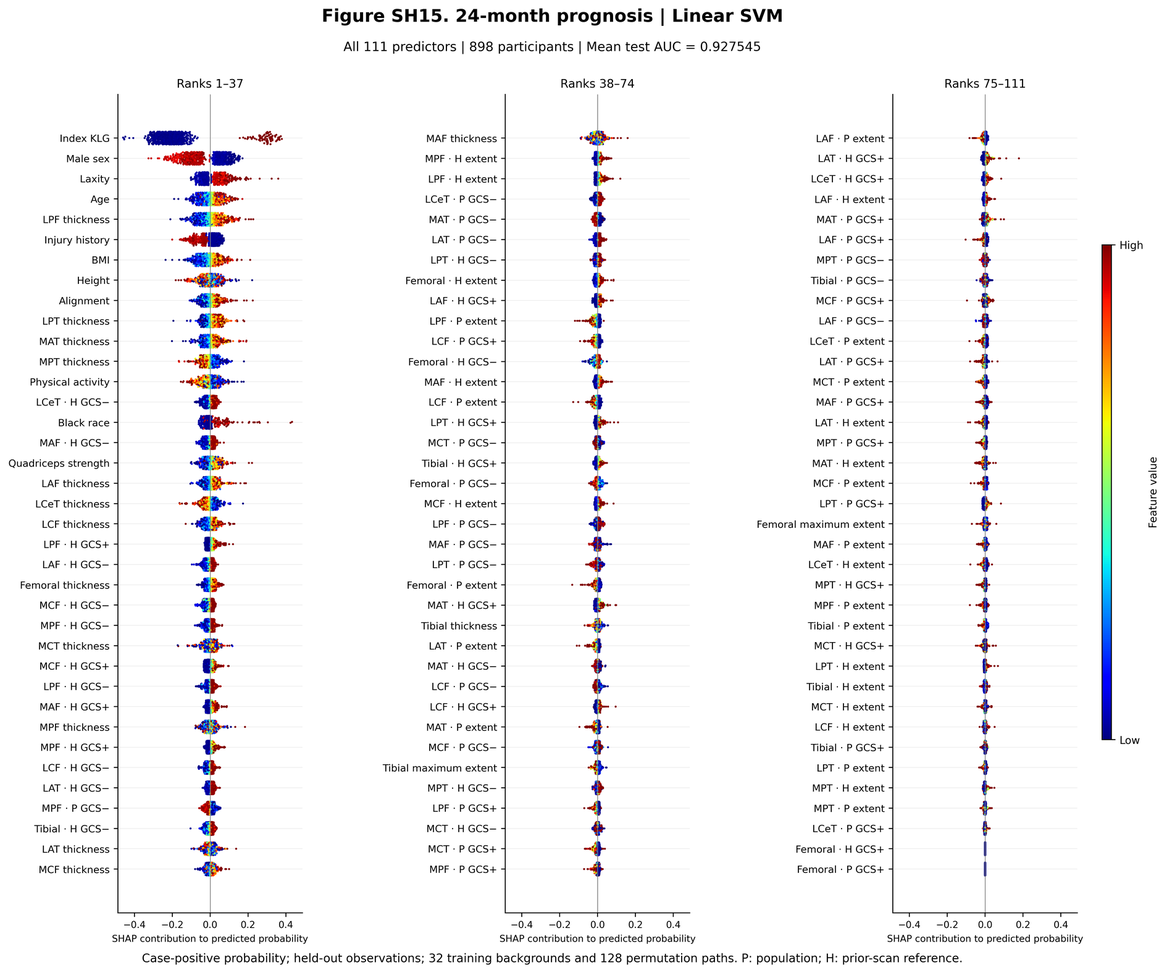

**Figure SH16.** 24-month prognosis — Linear SVM. Probability-scale SHAP values for all predictors, ordered by mean absolute SHAP (n = 898 participants). Centile features account for 39% of the total mean absolute SHAP.

*Leading predictors (mean |SHAP|, direction)*

| Rank | Feature | Mean | SHAP |
| --- | --- | --- | --- |
| 1 | Index KLG | 0.218 | higher → ↑ risk (ρ = 0.41) |
| 2 | Male sex | 0.080 | men lower |
| 3 | Laxity | 0.041 | higher → ↑ risk (ρ = 0.49) |
| 4 | Age | 0.035 | higher → ↑ risk (ρ = 0.73) |
| 5 | LPF thickness | 0.033 | higher → ↑ risk (ρ = 0.88) |
| 6 | Injury history | 0.031 | higher → ↓ risk (ρ = -0.74) |
| 7 | BMI | 0.029 | higher → ↑ risk (ρ = 0.82) |
| 8 | Height | 0.020 | non-monotone (ρ = -0.11) |

*Leading centile predictors*

| Feature | Mean | SHAP |
| --- | --- | --- |
| LCeT · H GCS− | 0.018 | higher → ↑ risk (ρ = 0.89) |
| MAF · H GCS− | 0.017 | higher → ↑ risk (ρ = 0.87) |
| LPF · H GCS+ | 0.014 | higher → ↑ risk (ρ = 0.62) |
| LAF · H GCS− | 0.014 | higher → ↑ risk (ρ = 0.87) |
| MCF · H GCS− | 0.013 | higher → ↑ risk (ρ = 0.87) |

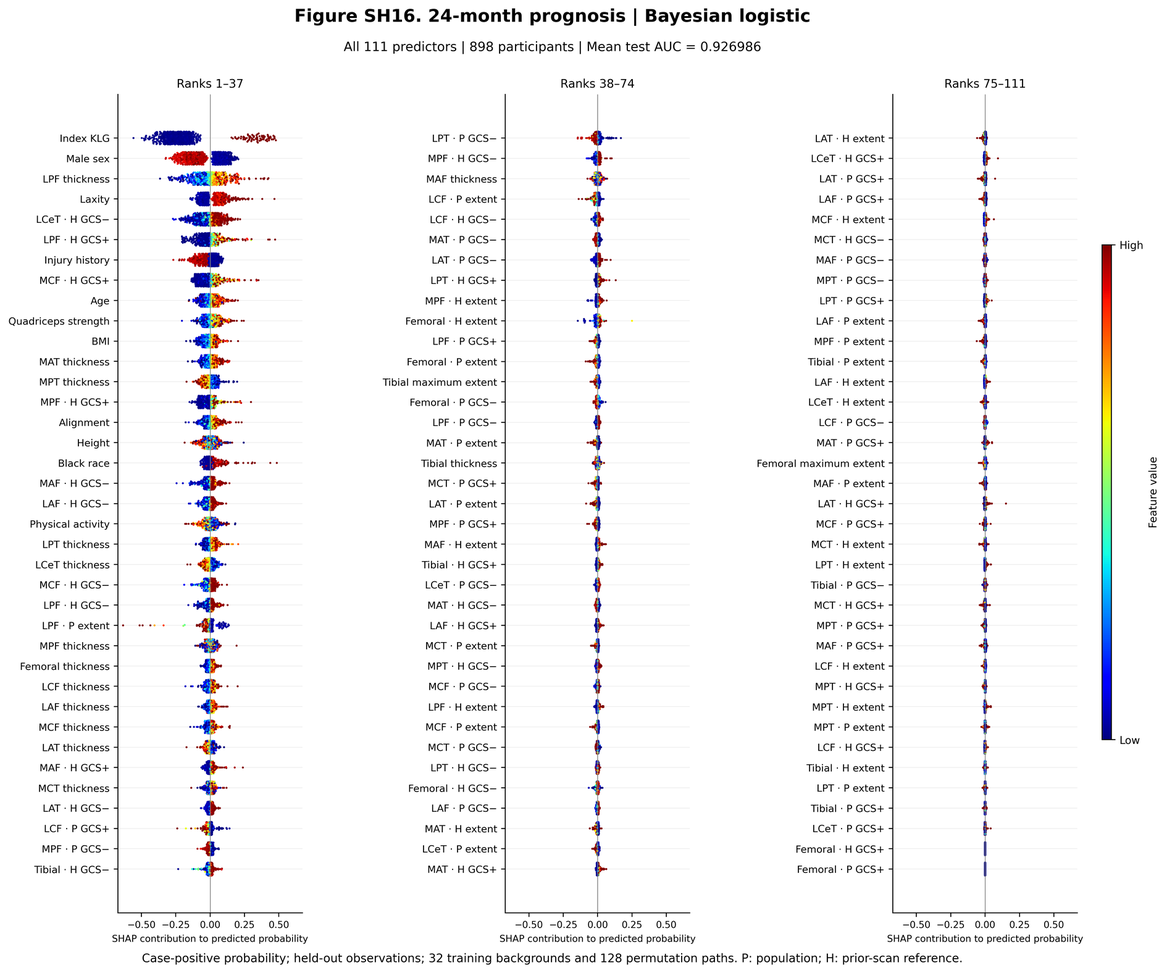

**Figure SH17.** 24-month prognosis — Bayesian logistic. Probability-scale SHAP values for all predictors, ordered by mean absolute SHAP (n = 898 participants). Centile features account for 37% of the total mean absolute SHAP.

*Leading predictors (mean |SHAP|, direction)*

| Rank | Feature | Mean | SHAP |
| --- | --- | --- | --- |
| 1 | Index KLG | 0.254 | higher → ↑ risk (ρ = 0.38) |
| 2 | Male sex | 0.103 | men lower |
| 3 | LPF thickness | 0.060 | higher → ↑ risk (ρ = 0.91) |
| 4 | Laxity | 0.052 | higher → ↑ risk (ρ = 0.53) |
| 5 | LCeT · H GCS− | 0.051 | higher → ↑ risk (ρ = 0.84) |
| 6 | LPF · H GCS+ | 0.038 | higher → ↑ risk (ρ = 0.58) |
| 7 | Injury history | 0.036 | higher → ↓ risk (ρ = -0.71) |
| 8 | MCF · H GCS+ | 0.035 | higher → ↑ risk (ρ = 0.59) |

*Leading centile predictors*

| Feature | Mean | SHAP |
| --- | --- | --- |
| LCeT · H GCS− | 0.051 | higher → ↑ risk (ρ = 0.84) |
| LPF · H GCS+ | 0.038 | higher → ↑ risk (ρ = 0.58) |
| MCF · H GCS+ | 0.035 | higher → ↑ risk (ρ = 0.59) |
| MPF · H GCS+ | 0.021 | higher → ↑ risk (ρ = 0.51) |
| MAF · H GCS− | 0.018 | higher → ↑ risk (ρ = 0.86) |

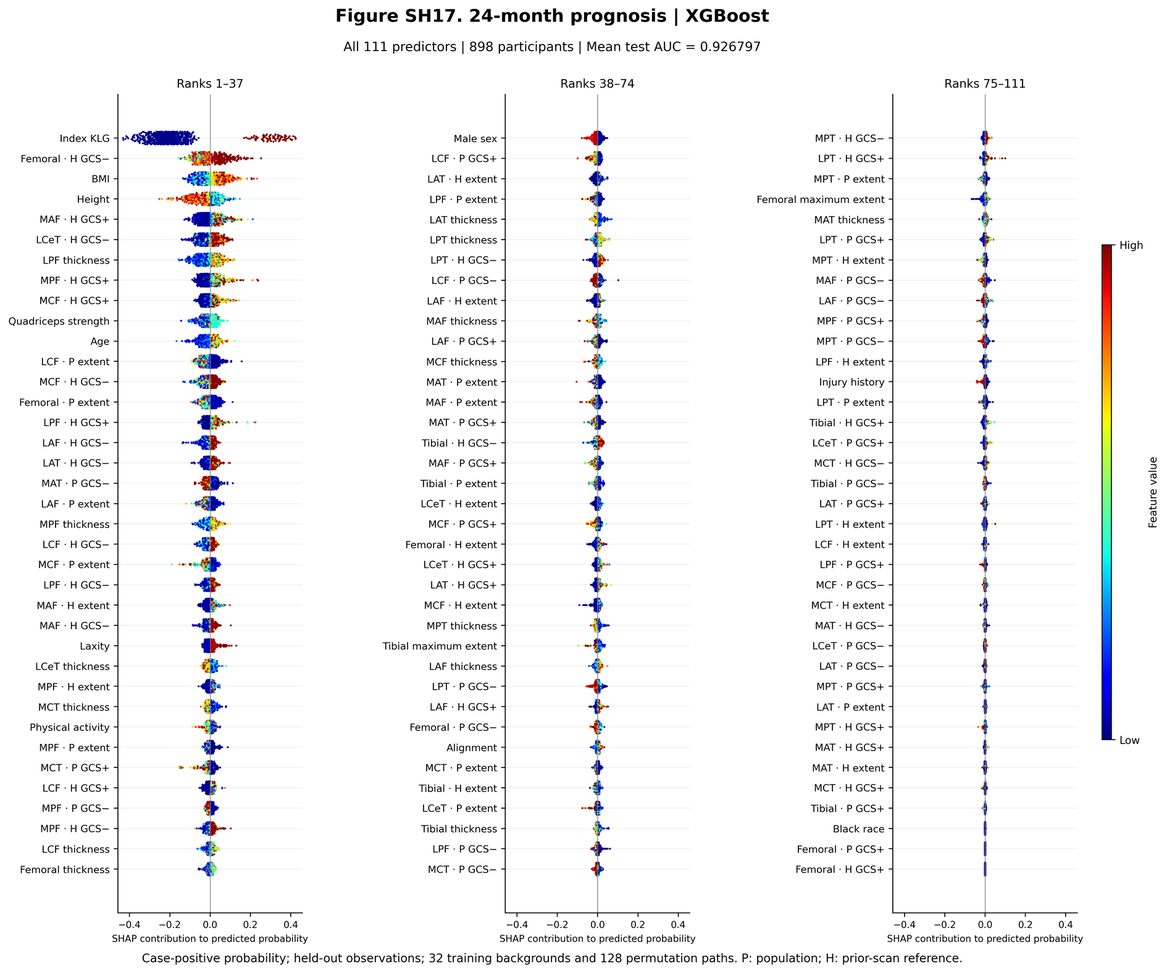

**Figure SH18.** 24-month prognosis — XGBoost. Probability-scale SHAP values for all predictors, ordered by mean absolute SHAP (n = 898 participants). Centile features account for 53% of the total mean absolute SHAP.

*Leading predictors (mean |SHAP|, direction)*

| Rank | Feature | Mean | SHAP |
| --- | --- | --- | --- |
| 1 | Index KLG | 0.224 | higher → ↑ risk (ρ = 0.42) |
| 2 | Femoral · H GCS− | 0.043 | higher → ↑ risk (ρ = 0.55) |
| 3 | BMI | 0.039 | higher → ↑ risk (ρ = 0.76) |
| 4 | Height | 0.031 | higher → ↓ risk (ρ = -0.58) |
| 5 | MAF · H GCS+ | 0.031 | higher → ↑ risk (ρ = 0.51) |
| 6 | LCeT · H GCS− | 0.029 | higher → ↑ risk (ρ = 0.80) |
| 7 | LPF thickness | 0.028 | higher → ↑ risk (ρ = 0.77) |
| 8 | MPF · H GCS+ | 0.027 | higher → ↑ risk (ρ = 0.52) |

*Leading centile predictors*

| Feature | Mean | SHAP |
| --- | --- | --- |
| Femoral · H GCS− | 0.043 | higher → ↑ risk (ρ = 0.55) |
| MAF · H GCS+ | 0.031 | higher → ↑ risk (ρ = 0.51) |
| LCeT · H GCS− | 0.029 | higher → ↑ risk (ρ = 0.80) |
| MPF · H GCS+ | 0.027 | higher → ↑ risk (ρ = 0.52) |
| MCF · H GCS+ | 0.020 | higher → ↑ risk (ρ = 0.57) |

#### S10.4 36-month prognosis

BMI was the second strongest predictor after index KLG (ρ ≈ 0.95), followed by sex, age and alignment. Centile features accounted for 7% of total attribution in ridge regression and 3% in LASSO, compared with 32–38% in Bayesian logistic regression, GLM boosting and random forest. MCF · H GCS+ was the leading centile predictor. These contributions were not accompanied by improved discrimination: adding centile profiles produced no significant AUC gain after Holm adjustment in any classifier, with significant losses in three (main Figure 3; Table S17).

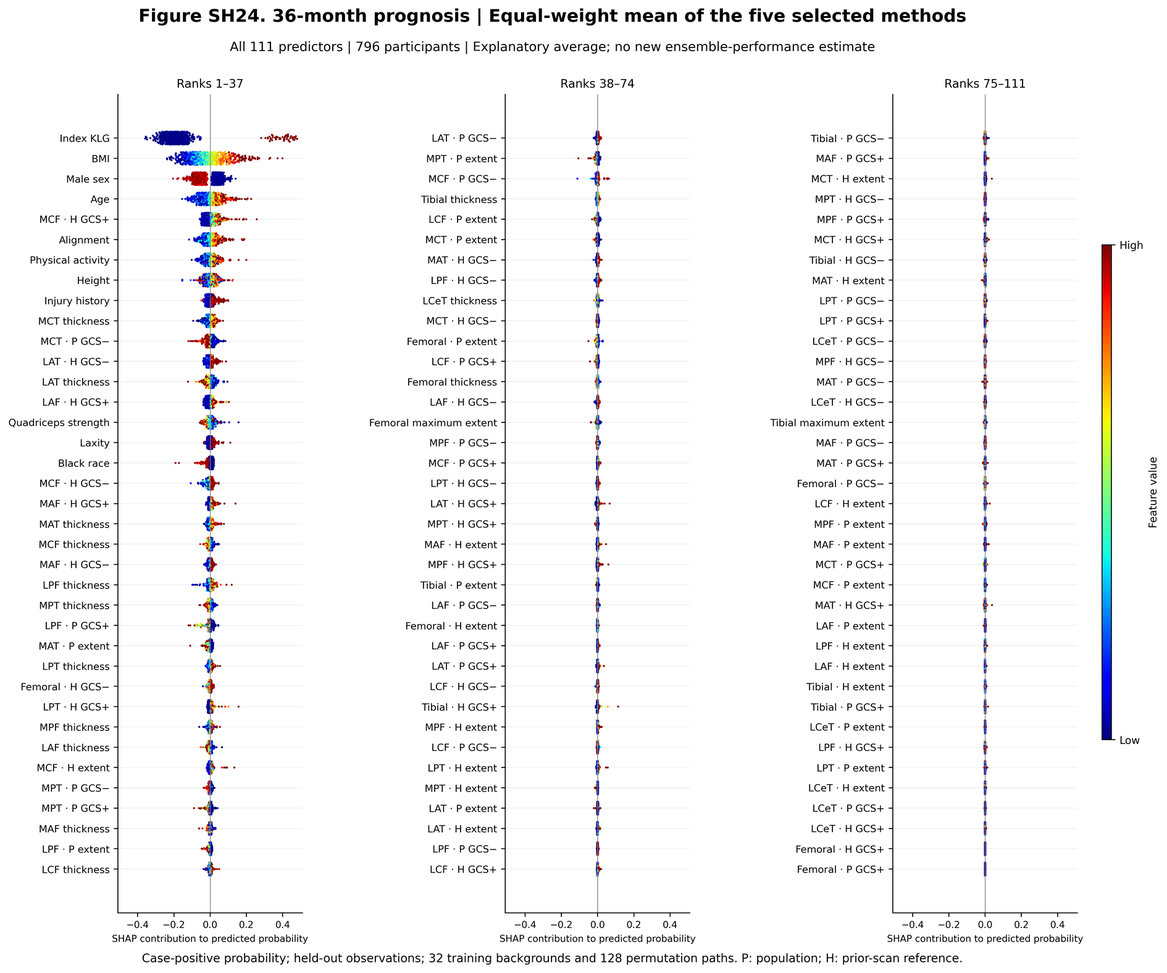

**Figure SH19.** 36-month prognosis — Equal-weight average of the five classifiers. Probability-scale SHAP values for all predictors, ordered by mean absolute SHAP (n = 796 participants). Centile features account for 24% of the total mean absolute SHAP.

*Leading predictors (mean |SHAP|, direction)*

| Rank | Feature | Mean | SHAP |
| --- | --- | --- | --- |
| 1 | Index KLG | 0.207 | higher → ↑ risk (ρ = 0.30) |
| 2 | BMI | 0.080 | higher → ↑ risk (ρ = 0.95) |
| 3 | Male sex | 0.050 | men lower |
| 4 | Age | 0.035 | higher → ↑ risk (ρ = 0.90) |
| 5 | MCF · H GCS+ | 0.021 | higher → ↑ risk (ρ = 0.56) |
| 6 | Alignment | 0.020 | higher → ↑ risk (ρ = 0.81) |
| 7 | Physical activity | 0.015 | higher → ↑ risk (ρ = 0.69) |
| 8 | Height | 0.015 | higher → ↑ risk (ρ = 0.50) |

*Leading centile predictors*

| Feature | Mean | SHAP |
| --- | --- | --- |
| MCF · H GCS+ | 0.021 | higher → ↑ risk (ρ = 0.56) |
| MCT · P GCS− | 0.010 | higher → ↓ risk (ρ = -0.80) |
| LAT · H GCS− | 0.009 | higher → ↑ risk (ρ = 0.72) |
| LAF · H GCS+ | 0.008 | higher → ↑ risk (ρ = 0.47) |
| MCF · H GCS− | 0.006 | higher → ↑ risk (ρ = 0.86) |

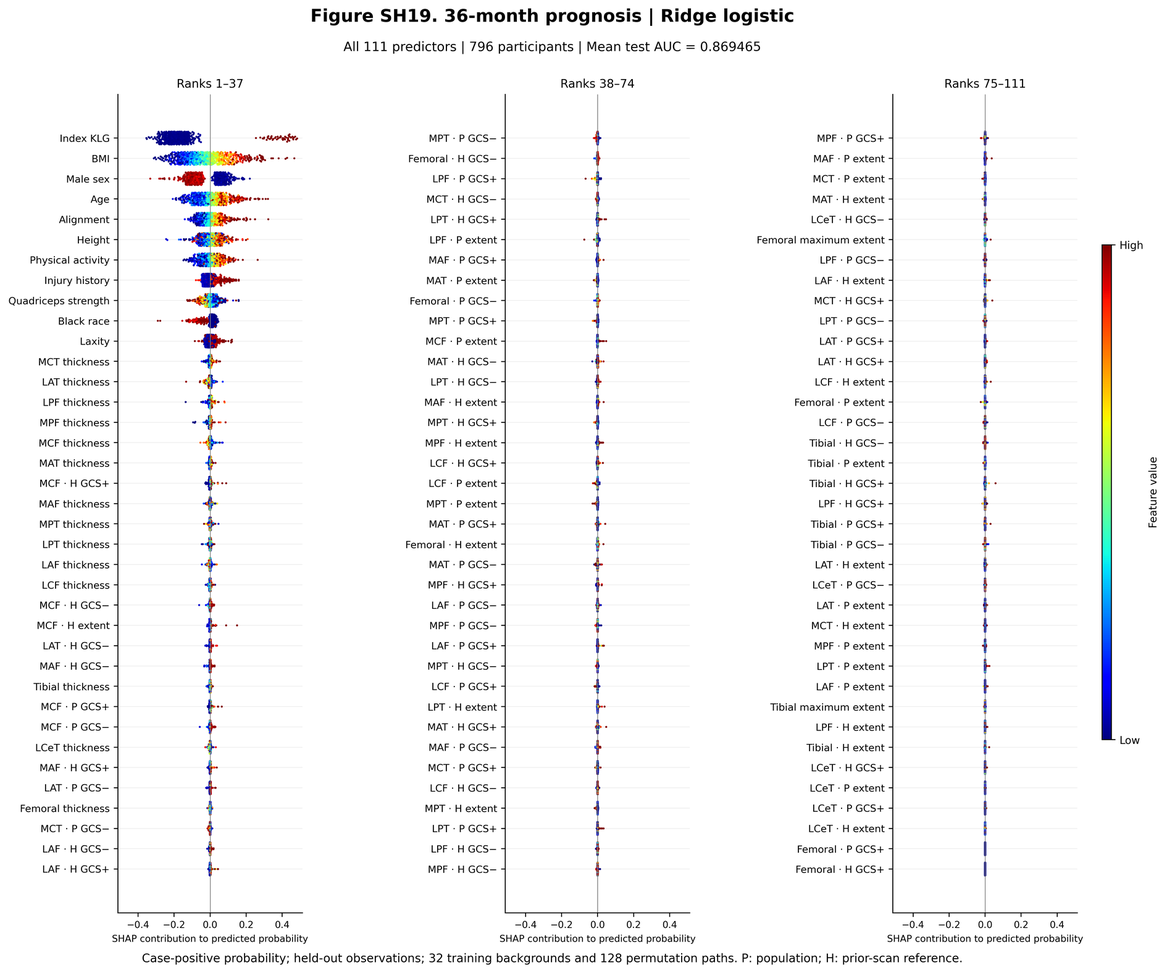

**Figure SH20.** 36-month prognosis — Ridge. Probability-scale SHAP values for all predictors, ordered by mean absolute SHAP (n = 796 participants). Centile features account for 7% of the total mean absolute SHAP.

*Leading predictors (mean |SHAP|, direction)*

| Rank | Feature | Mean | SHAP |
| --- | --- | --- | --- |
| 1 | Index KLG | 0.202 | higher → ↑ risk (ρ = 0.30) |
| 2 | BMI | 0.103 | higher → ↑ risk (ρ = 0.96) |
| 3 | Male sex | 0.077 | men lower |
| 4 | Age | 0.060 | higher → ↑ risk (ρ = 0.91) |
| 5 | Alignment | 0.035 | higher → ↑ risk (ρ = 0.91) |
| 6 | Height | 0.031 | higher → ↑ risk (ρ = 0.63) |
| 7 | Physical activity | 0.031 | higher → ↑ risk (ρ = 0.81) |
| 8 | Injury history | 0.025 | non-monotone (ρ = 0.15) |

*Leading centile predictors*

| Feature | Mean | SHAP |
| --- | --- | --- |
| MCF · H GCS+ | 0.002 | higher → ↑ risk (ρ = 0.63) |
| MCF · H GCS− | 0.001 | higher → ↑ risk (ρ = 0.90) |
| MCF · H extent | 0.001 | higher → ↑ risk (ρ = 0.75) |
| LAT · H GCS− | 0.001 | higher → ↑ risk (ρ = 0.75) |
| MAF · H GCS− | 0.001 | higher → ↑ risk (ρ = 0.87) |

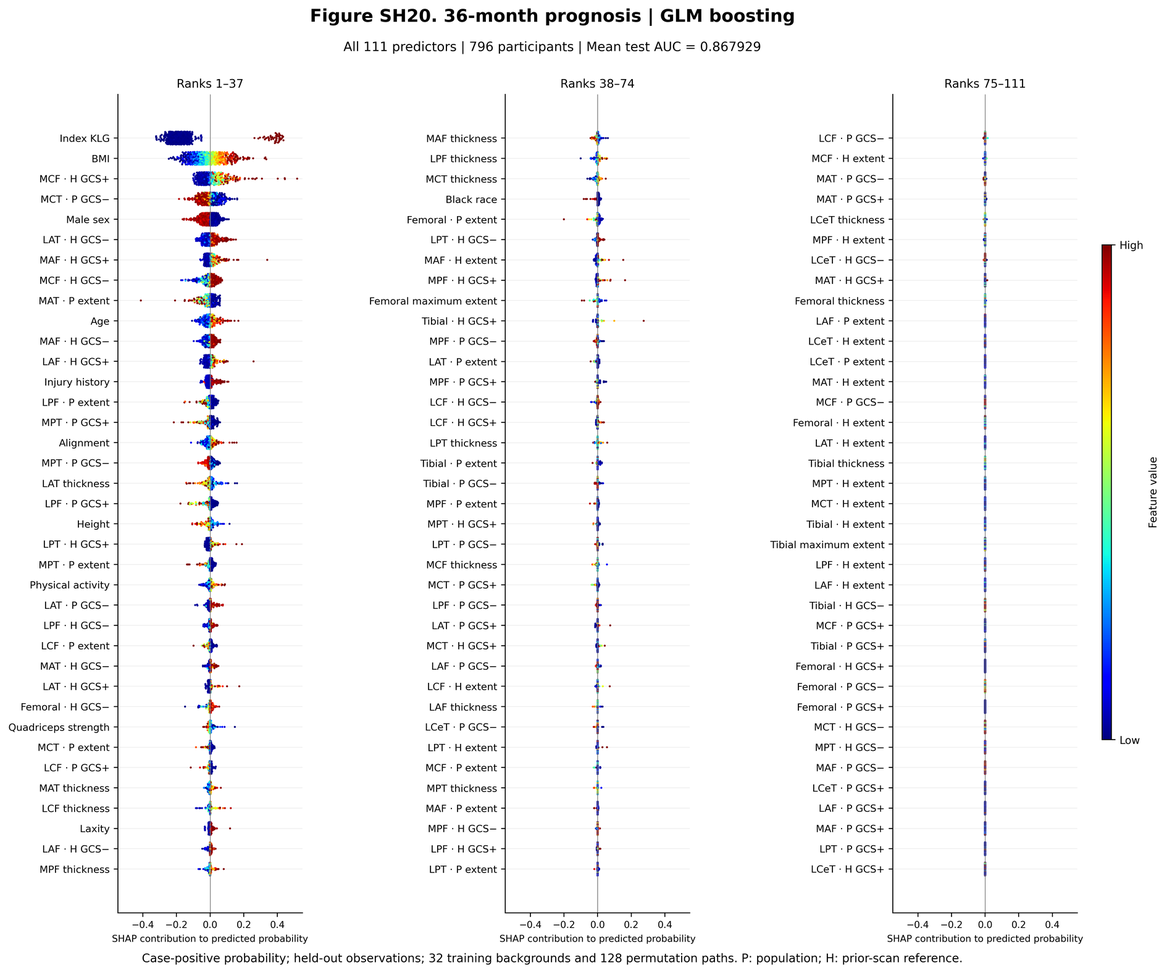

**Figure SH21.** 36-month prognosis — GLM boosting. Probability-scale SHAP values for all predictors, ordered by mean absolute SHAP (n = 796 participants). Centile features account for 38% of the total mean absolute SHAP.

*Leading predictors (mean |SHAP|, direction)*

| Rank | Feature | Mean | SHAP |
| --- | --- | --- | --- |
| 1 | Index KLG | 0.197 | higher → ↑ risk (ρ = 0.31) |
| 2 | BMI | 0.072 | higher → ↑ risk (ρ = 0.96) |
| 3 | MCF · H GCS+ | 0.041 | higher → ↑ risk (ρ = 0.55) |
| 4 | MCT · P GCS− | 0.026 | higher → ↓ risk (ρ = -0.72) |
| 5 | Male sex | 0.025 | men lower |
| 6 | LAT · H GCS− | 0.018 | higher → ↑ risk (ρ = 0.62) |
| 7 | MAF · H GCS+ | 0.013 | higher → ↑ risk (ρ = 0.51) |
| 8 | MCF · H GCS− | 0.013 | higher → ↑ risk (ρ = 0.67) |

*Leading centile predictors*

| Feature | Mean | SHAP |
| --- | --- | --- |
| MCF · H GCS+ | 0.041 | higher → ↑ risk (ρ = 0.55) |
| MCT · P GCS− | 0.026 | higher → ↓ risk (ρ = -0.72) |
| LAT · H GCS− | 0.018 | higher → ↑ risk (ρ = 0.62) |
| MAF · H GCS+ | 0.013 | higher → ↑ risk (ρ = 0.51) |
| MCF · H GCS− | 0.013 | higher → ↑ risk (ρ = 0.67) |

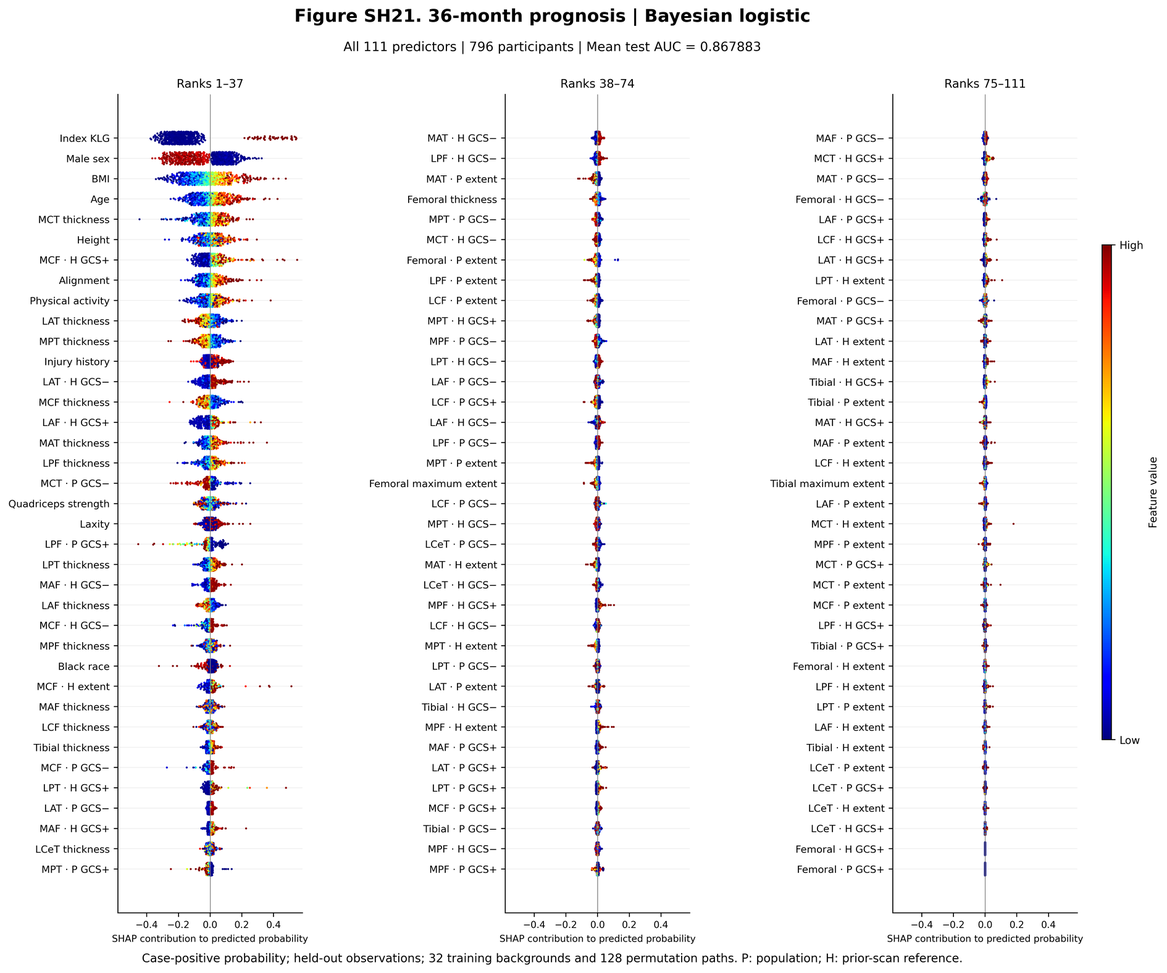

**Figure SH22.** 36-month prognosis — Bayesian logistic. Probability-scale SHAP values for all predictors, ordered by mean absolute SHAP (n = 796 participants). Centile features account for 32% of the total mean absolute SHAP.

*Leading predictors (mean |SHAP|, direction)*

| Rank | Feature | Mean | SHAP |
| --- | --- | --- | --- |
| 1 | Index KLG | 0.206 | non-monotone (ρ = 0.26) |
| 2 | Male sex | 0.117 | men lower |
| 3 | BMI | 0.101 | higher → ↑ risk (ρ = 0.93) |
| 4 | Age | 0.062 | higher → ↑ risk (ρ = 0.89) |
| 5 | MCT thickness | 0.041 | higher → ↑ risk (ρ = 0.92) |
| 6 | Height | 0.037 | higher → ↑ risk (ρ = 0.52) |
| 7 | MCF · H GCS+ | 0.036 | higher → ↑ risk (ρ = 0.65) |
| 8 | Alignment | 0.031 | higher → ↑ risk (ρ = 0.83) |

*Leading centile predictors*

| Feature | Mean | SHAP |
| --- | --- | --- |
| MCF · H GCS+ | 0.036 | higher → ↑ risk (ρ = 0.65) |
| LAT · H GCS− | 0.021 | higher → ↑ risk (ρ = 0.74) |
| LAF · H GCS+ | 0.020 | higher → ↑ risk (ρ = 0.53) |
| MCT · P GCS− | 0.018 | higher → ↓ risk (ρ = -0.82) |
| LPF · P GCS+ | 0.015 | higher → ↓ risk (ρ = -0.77) |

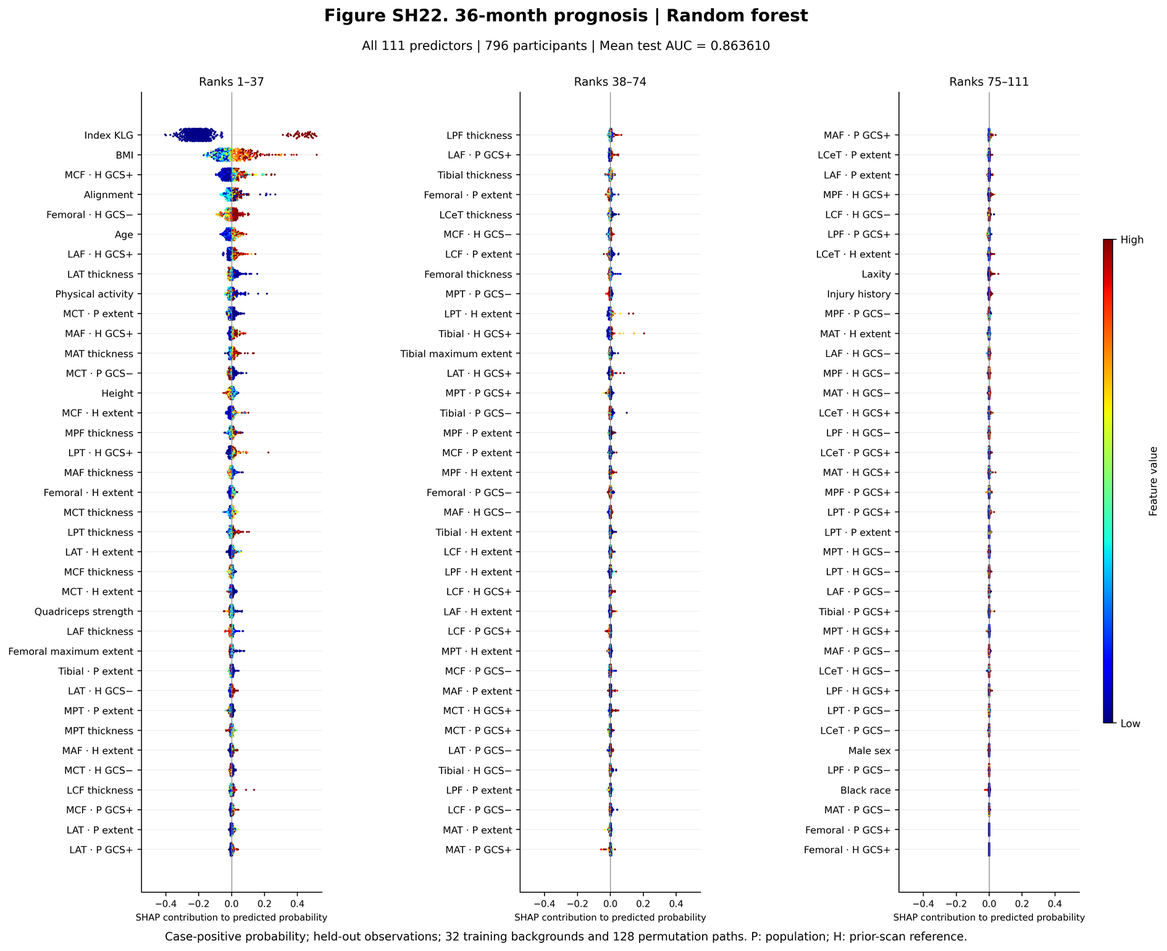

**Figure SH23.** 36-month prognosis — Random forest. Probability-scale SHAP values for all predictors, ordered by mean absolute SHAP (n = 796 participants). Centile features account for 35% of the total mean absolute SHAP.

*Leading predictors (mean |SHAP|, direction)*

| Rank | Feature | Mean | SHAP |
| --- | --- | --- | --- |
| 1 | Index KLG | 0.222 | non-monotone (ρ = 0.27) |
| 2 | BMI | 0.052 | higher → ↑ risk (ρ = 0.57) |
| 3 | MCF · H GCS+ | 0.024 | higher → ↑ risk (ρ = 0.48) |
| 4 | Alignment | 0.016 | non-monotone (ρ = 0.25) |
| 5 | Femoral · H GCS− | 0.015 | higher → ↑ risk (ρ = 0.67) |
| 6 | Age | 0.011 | higher → ↑ risk (ρ = 0.77) |
| 7 | LAF · H GCS+ | 0.009 | higher → ↑ risk (ρ = 0.37) |
| 8 | LAT thickness | 0.008 | higher → ↓ risk (ρ = -0.74) |

*Leading centile predictors*

| Feature | Mean | SHAP |
| --- | --- | --- |
| MCF · H GCS+ | 0.024 | higher → ↑ risk (ρ = 0.48) |
| Femoral · H GCS− | 0.015 | higher → ↑ risk (ρ = 0.67) |
| LAF · H GCS+ | 0.009 | higher → ↑ risk (ρ = 0.37) |
| MCT · P extent | 0.007 | higher → ↓ risk (ρ = -0.69) |
| MAF · H GCS+ | 0.007 | higher → ↑ risk (ρ = 0.48) |

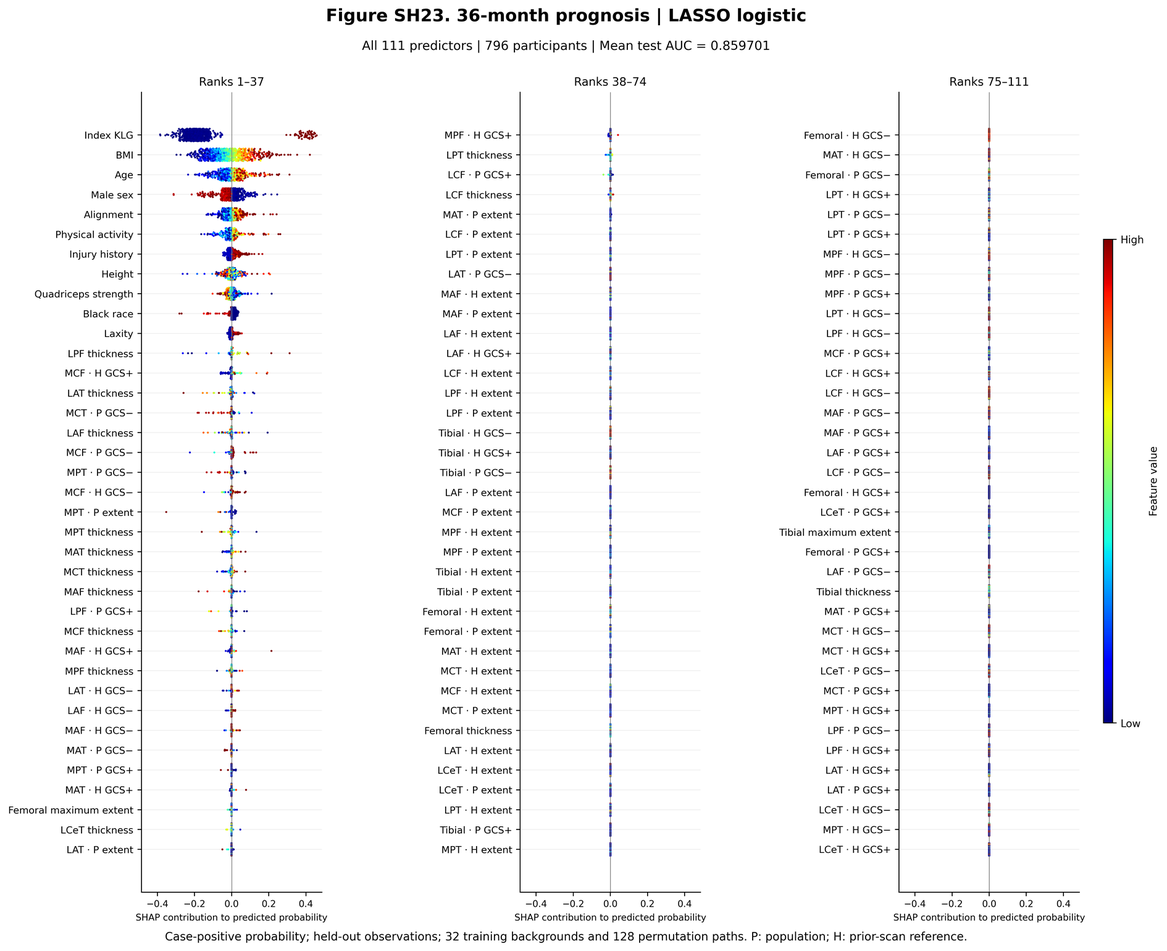

**Figure SH24.** 36-month prognosis — LASSO. Probability-scale SHAP values for all predictors, ordered by mean absolute SHAP (n = 796 participants). Centile features account for 3% of the total mean absolute SHAP.

*Leading predictors (mean |SHAP|, direction)*

| Rank | Feature | Mean | SHAP |
| --- | --- | --- | --- |
| 1 | Index KLG | 0.206 | higher → ↑ risk (ρ = 0.33) |
| 2 | BMI | 0.080 | higher → ↑ risk (ρ = 0.96) |
| 3 | Age | 0.029 | higher → ↑ risk (ρ = 0.77) |
| 4 | Male sex | 0.028 | men lower |
| 5 | Alignment | 0.018 | higher → ↑ risk (ρ = 0.83) |
| 6 | Physical activity | 0.010 | higher → ↑ risk (ρ = 0.56) |
| 7 | Injury history | 0.008 | non-monotone (ρ = 0.21) |
| 8 | Height | 0.008 | non-monotone (ρ = -0.15) |

*Leading centile predictors*

| Feature | Mean | SHAP |
| --- | --- | --- |
| MCF · H GCS+ | 0.002 | non-monotone (ρ = 0.14) |
| MCT · P GCS− | 0.002 | non-monotone (ρ = -0.14) |
| MCF · P GCS− | 0.001 | non-monotone (ρ = 0.26) |
| MPT · P GCS− | 0.001 | non-monotone (ρ = -0.20) |
| MCF · H GCS− | 0.001 | non-monotone (ρ = 0.21) |

**Group distributions of leading centile predictors** are shown in Figure A8.

### S11. Reference-chart and descriptive atlas (Figures A1–A17)

Figures A1–A7 and A13–A14 show regional reference models fitted as described in S2. The surfaces represent population mean thickness conditional on age, BMI and sex, with height fixed at 1678.5 mm, race at the reference category and participant intercepts excluded. Left-knee charts provide anatomical comparisons; prediction and longitudinal analyses used right knees. Hatching on contour plots marks locations outside the observed sex-specific age–BMI convex hull. Confidence limits describe uncertainty in the fitted mean, conditional on the smoothing parameters; they are not individual prediction intervals or centile envelopes. The surfaces describe covariate associations rather than measured within-person change or causal BMI effects. The remaining figures report feature distributions, prognostic increments, observed thickness and grade changes, and sensitivity to follow-up availability.

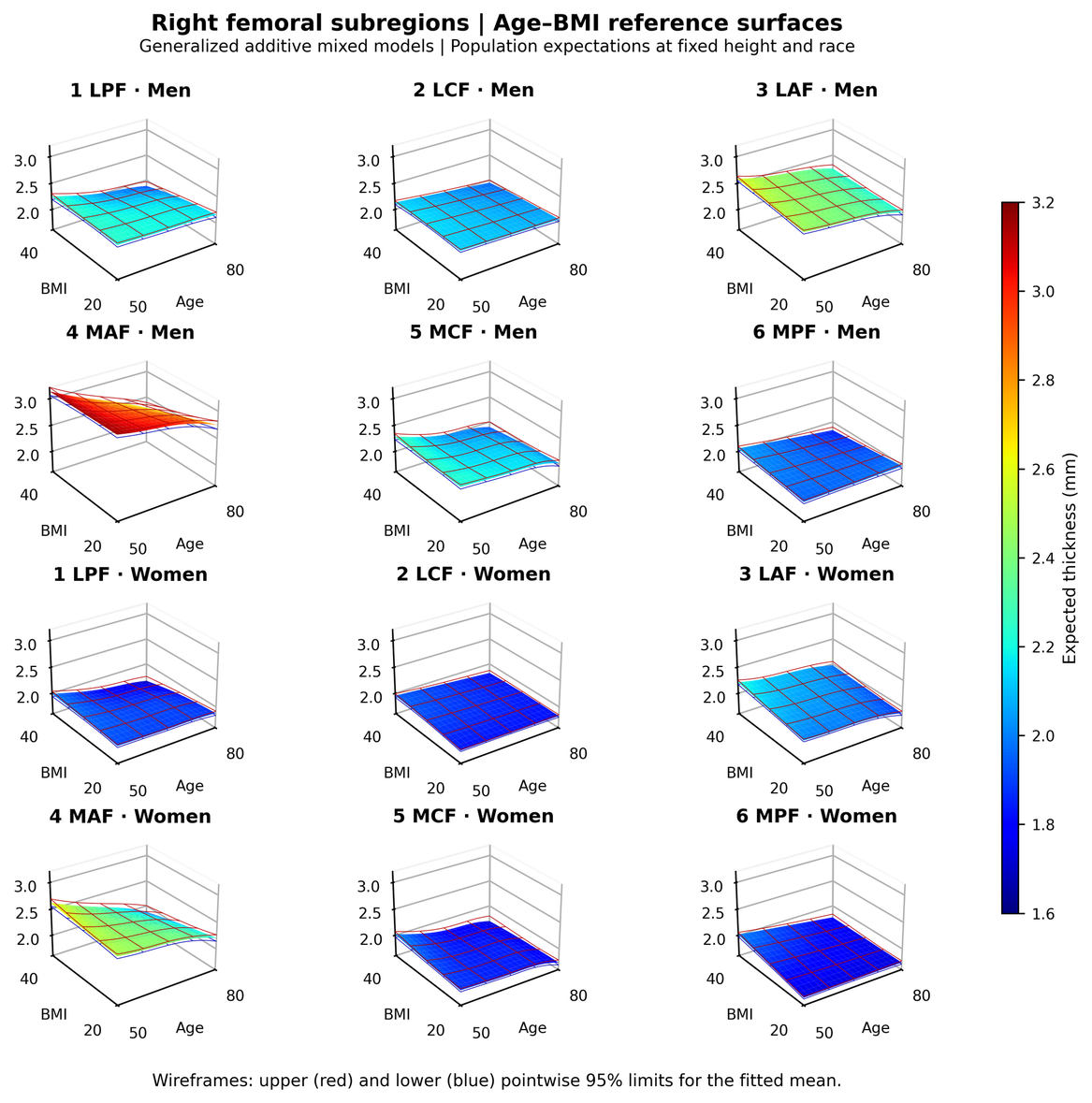

**Figure A1.** Right femoral subregional reference surfaces. Fitted mean thickness is shown for all six subregions in men and women over ages 50–80 years and BMI 20–40 kg/m². Height is fixed at 1678.5 mm and race at the reference category; participant intercepts are excluded. Red and blue wireframes show the upper and lower pointwise 95% confidence limits for the mean (±1.96 model standard errors), conditional on the fitted smoothing parameters. Each compartment uses a common scale across knees and sexes. Region abbreviations follow main Figure 1D. The corresponding contours are shown in Figure A13.

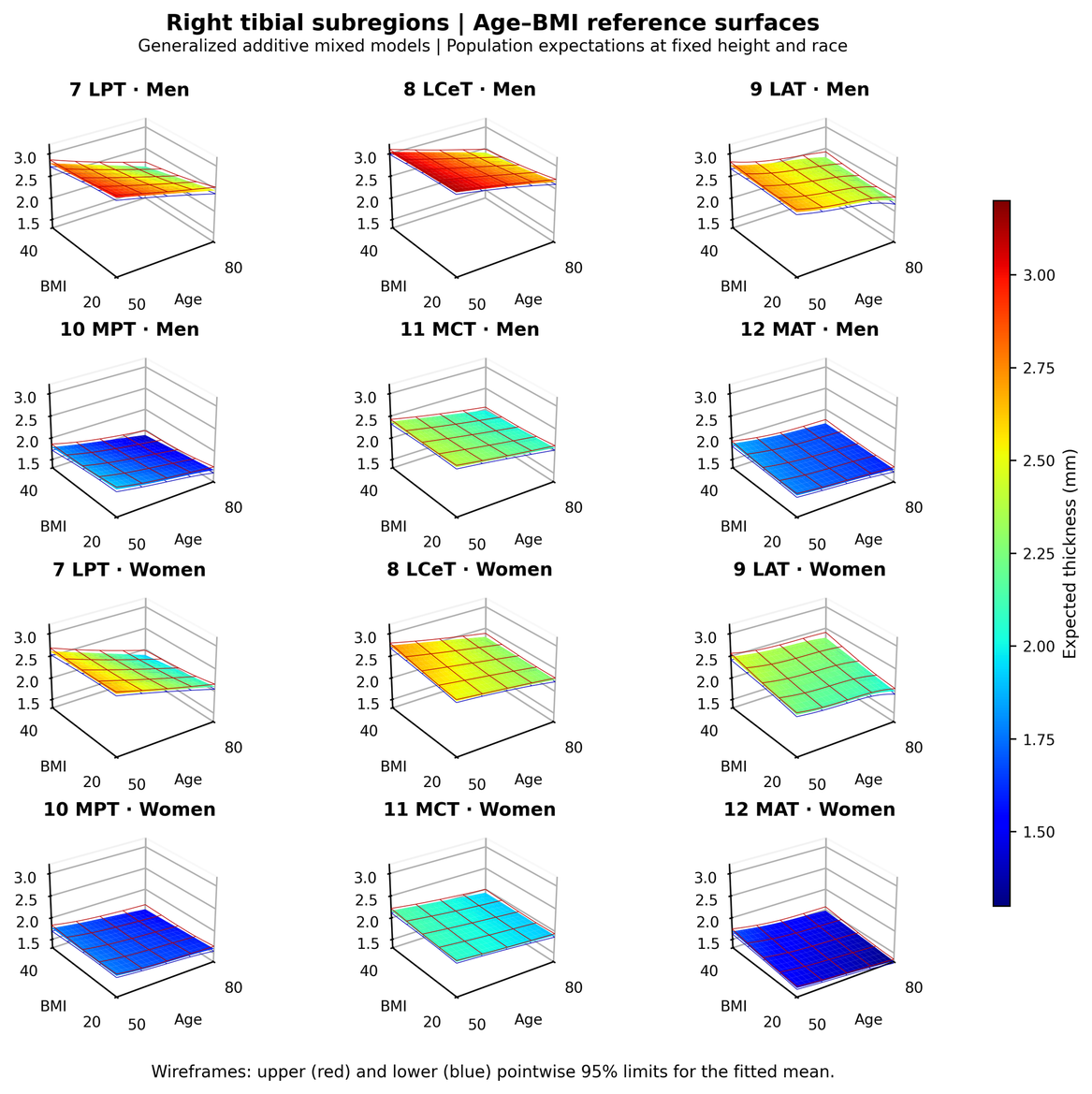

**Figure A2.** Right tibial subregional reference surfaces. Fitted mean thickness is shown for all six subregions in men and women over ages 50–80 years and BMI 20–40 kg/m². Height is fixed at 1678.5 mm and race at the reference category; participant intercepts are excluded. Red and blue wireframes show the upper and lower pointwise 95% confidence limits for the mean (±1.96 model standard errors), conditional on the fitted smoothing parameters. Each compartment uses a common scale across knees and sexes. Region abbreviations follow main Figure 1D. The corresponding contours are shown in Figure A14.

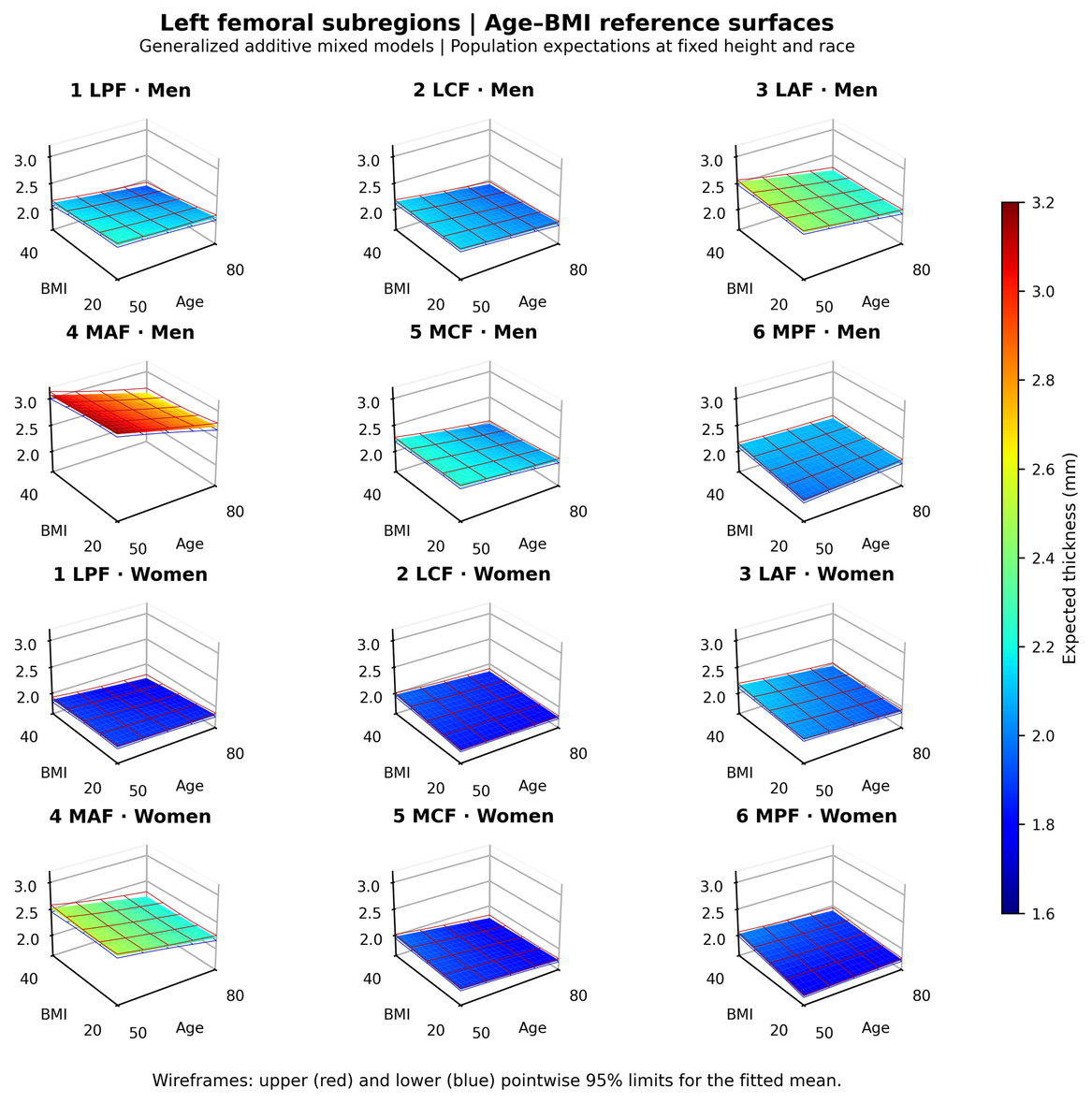

**Figure A3.** Left femoral subregional reference surfaces. Fitted mean thickness is shown for all six subregions in men and women over ages 50–80 years and BMI 20–40 kg/m². Height is fixed at 1678.5 mm and race at the reference category; participant intercepts are excluded. Red and blue wireframes show the upper and lower pointwise 95% confidence limits for the mean (±1.96 model standard errors), conditional on the fitted smoothing parameters. Each compartment uses a common scale across knees and sexes. Region abbreviations follow main Figure 1D. The corresponding contours are shown in Figure A5.

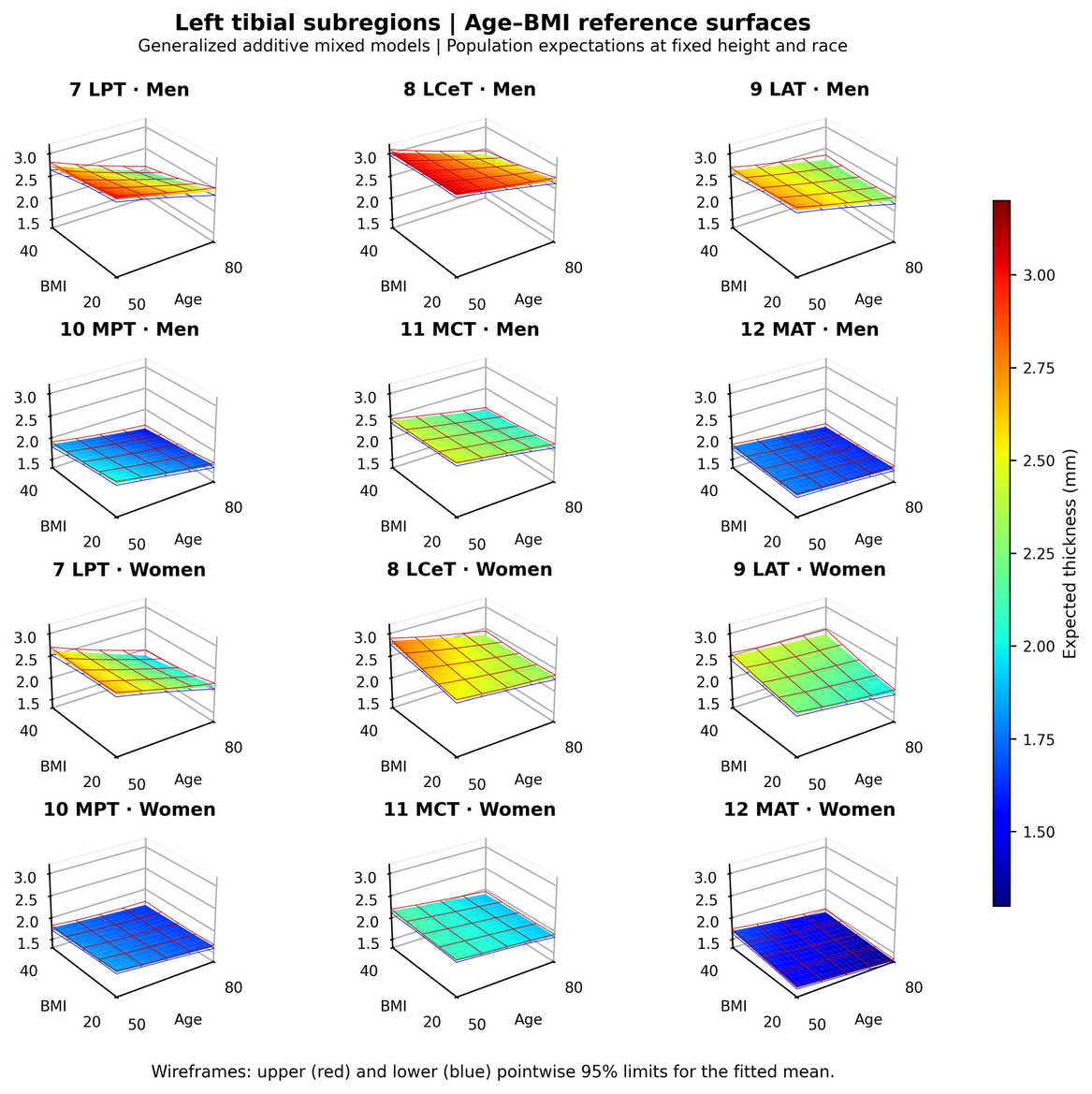

**Figure A4.** Left tibial subregional reference surfaces. Fitted mean thickness is shown for all six subregions in men and women over ages 50–80 years and BMI 20–40 kg/m². Height is fixed at 1678.5 mm and race at the reference category; participant intercepts are excluded. Red and blue wireframes show the upper and lower pointwise 95% confidence limits for the mean (±1.96 model standard errors), conditional on the fitted smoothing parameters. Each compartment uses a common scale across knees and sexes. Region abbreviations follow main Figure 1D. The corresponding contours are shown in Figure A6.

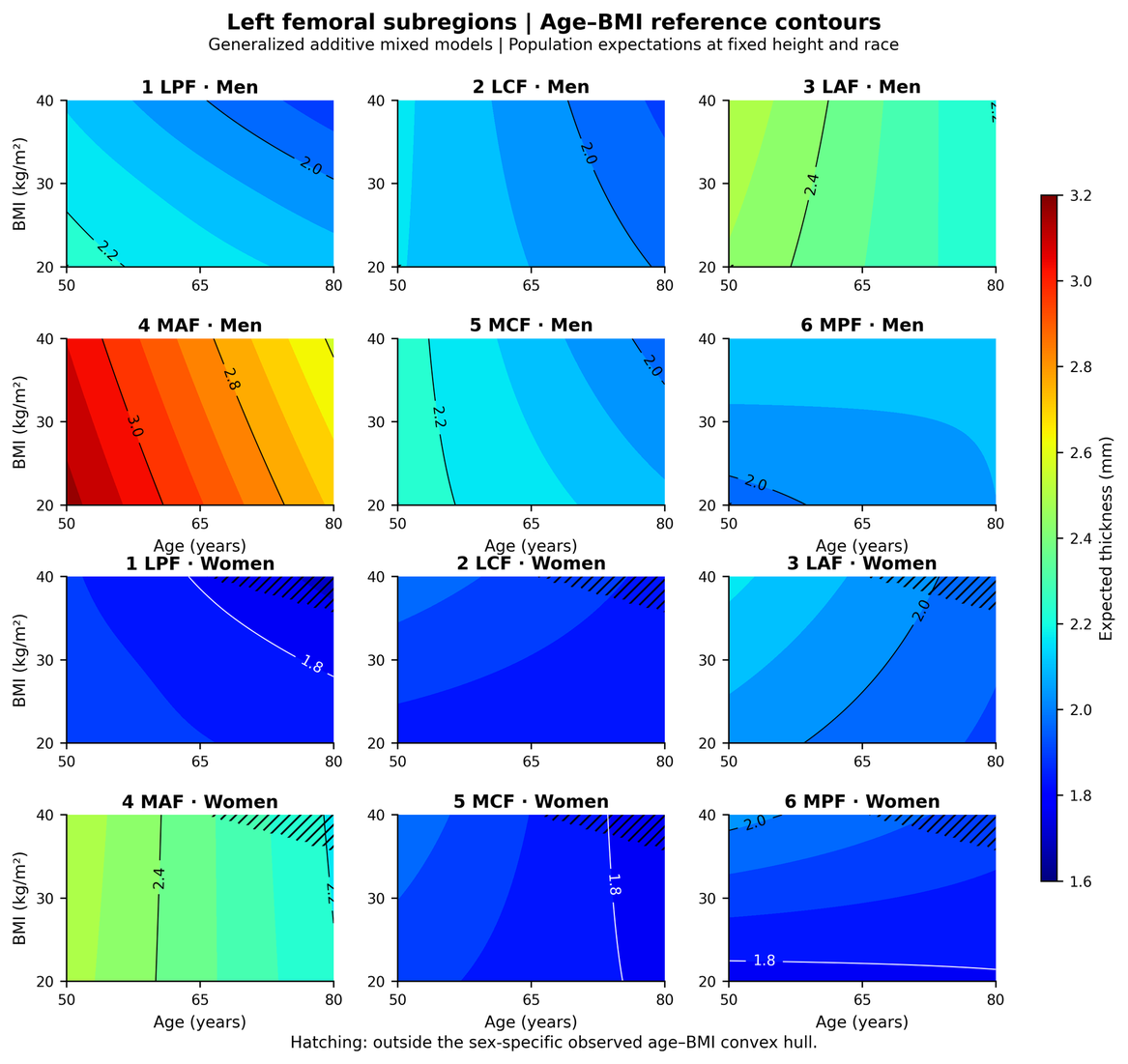

**Figure A5.** Left femoral subregional age–BMI contours. Colours and labelled contours show fitted mean thickness in millimetres for all six subregions in men and women. Predictions cover ages 50–80 years and BMI 20–40 kg/m², with height fixed at 1678.5 mm, race at the reference category and participant intercepts excluded. Hatching marks predictions outside the observed sex-specific age–BMI convex hull. Each compartment uses a common scale across knees and sexes. Region abbreviations follow main Figure 1D. Figure A3 shows the same fitted surfaces in perspective.

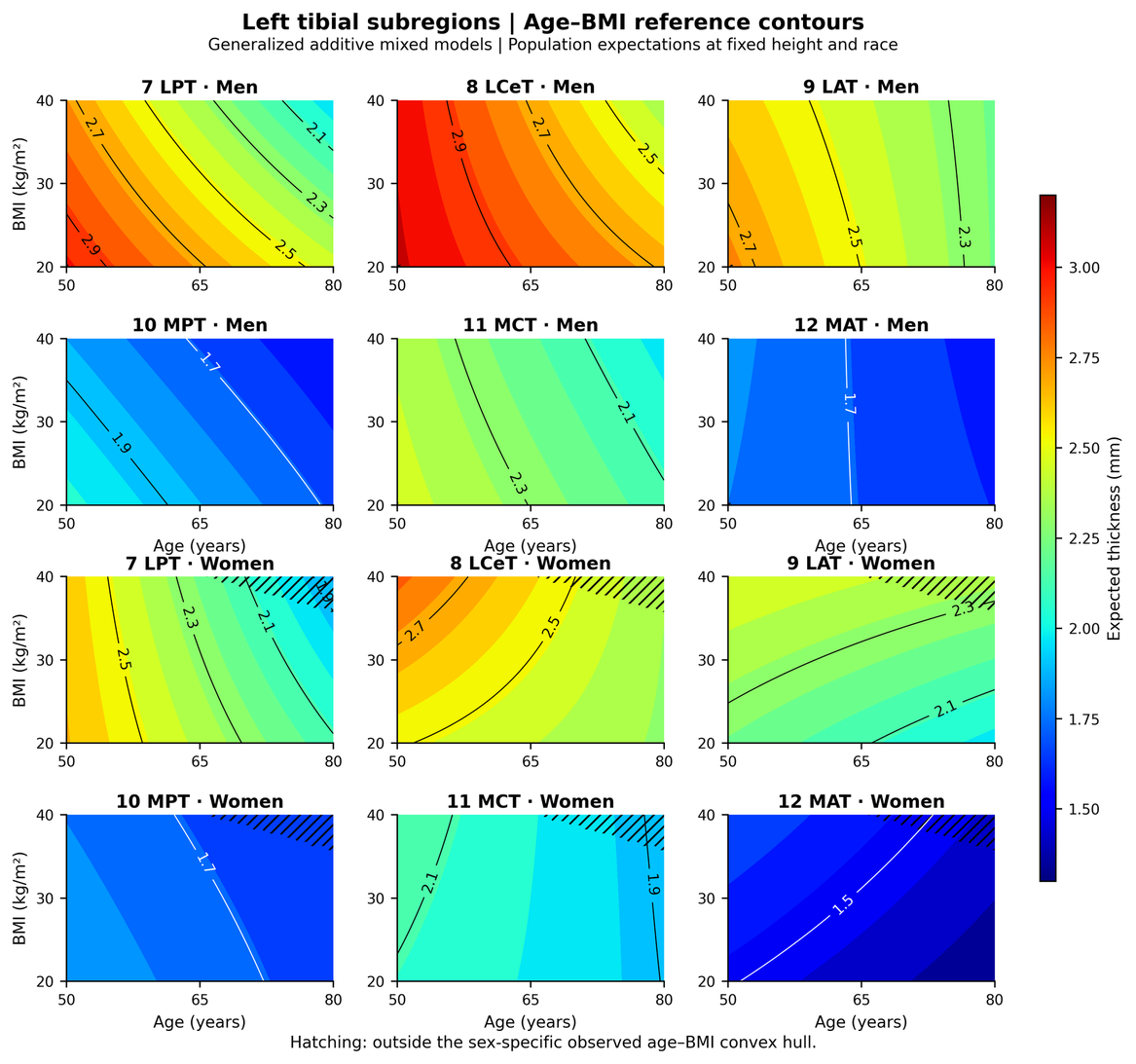

**Figure A6.** Left tibial subregional age–BMI contours. Colours and labelled contours show fitted mean thickness in millimetres for all six subregions in men and women. Predictions cover ages 50–80 years and BMI 20–40 kg/m², with height fixed at 1678.5 mm, race at the reference category and participant intercepts excluded. Hatching marks predictions outside the observed sex-specific age–BMI convex hull. Each compartment uses a common scale across knees and sexes. Region abbreviations follow main Figure 1D. Figure A4 shows the same fitted surfaces in perspective.

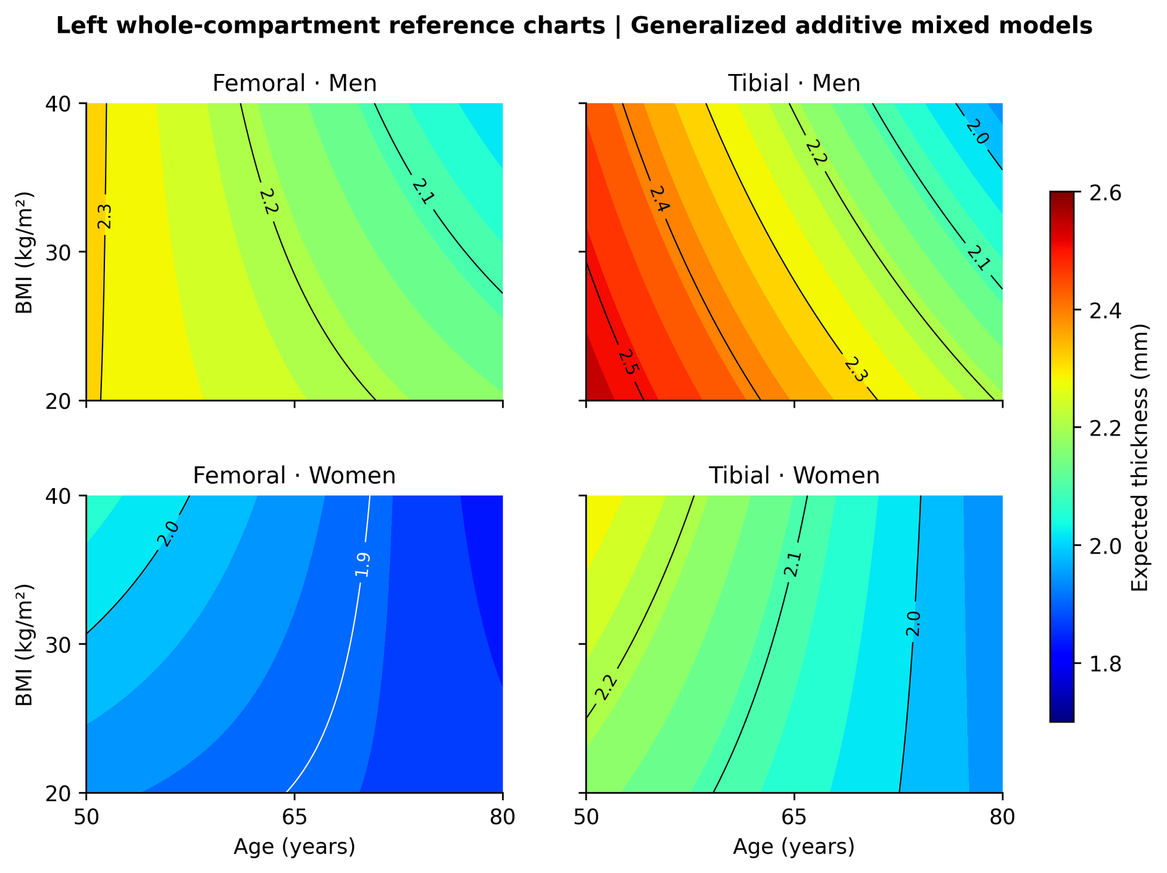

**Figure A7.** Left whole-compartment age–BMI contours. Colours and labelled contours show fitted mean thickness in millimetres for the femoral and tibial plates in men and women. Covariate settings are the same as in Figures A1–A6. The corresponding right-knee charts appear in main Figure 1C; models were fitted separately for the two knees.

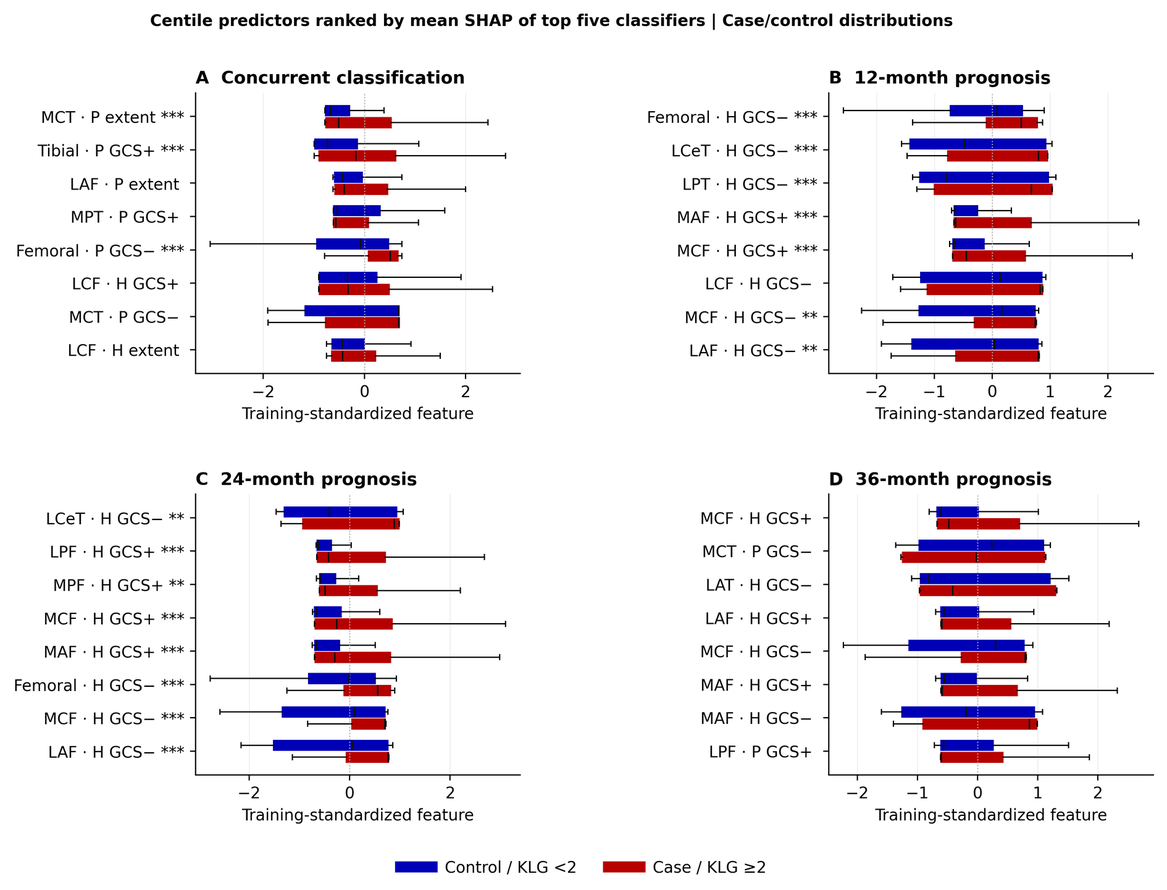

**Figure A8.** Group distributions of leading centile predictors. For each outcome, the eight centile features with the highest mean absolute SHAP were selected from the equal-weight average of the five highest-AUC classifiers (main Figure 6). Boxes show the median and interquartile range (IQR), with whiskers extending to 1.5 IQR. Outliers are omitted from the display but retained in the analysis. Blue denotes controls and red cases. Each participant contributes the earliest included scan. Displayed values were standardized using outer-training statistics and averaged across held-out appearances. Stars indicate two-sided Mann–Whitney P values below .05, .01 and .001 after Holm adjustment across 436 predictor/outcome tests, calculated from the original available values. Exact P values are provided in the supplementary data files. P and H denote cross-sectional and longitudinal centiles; GCS+, GCS− and extent are defined in Table S5. These distributions describe features selected from the fitted models and do not estimate independent biological effects.

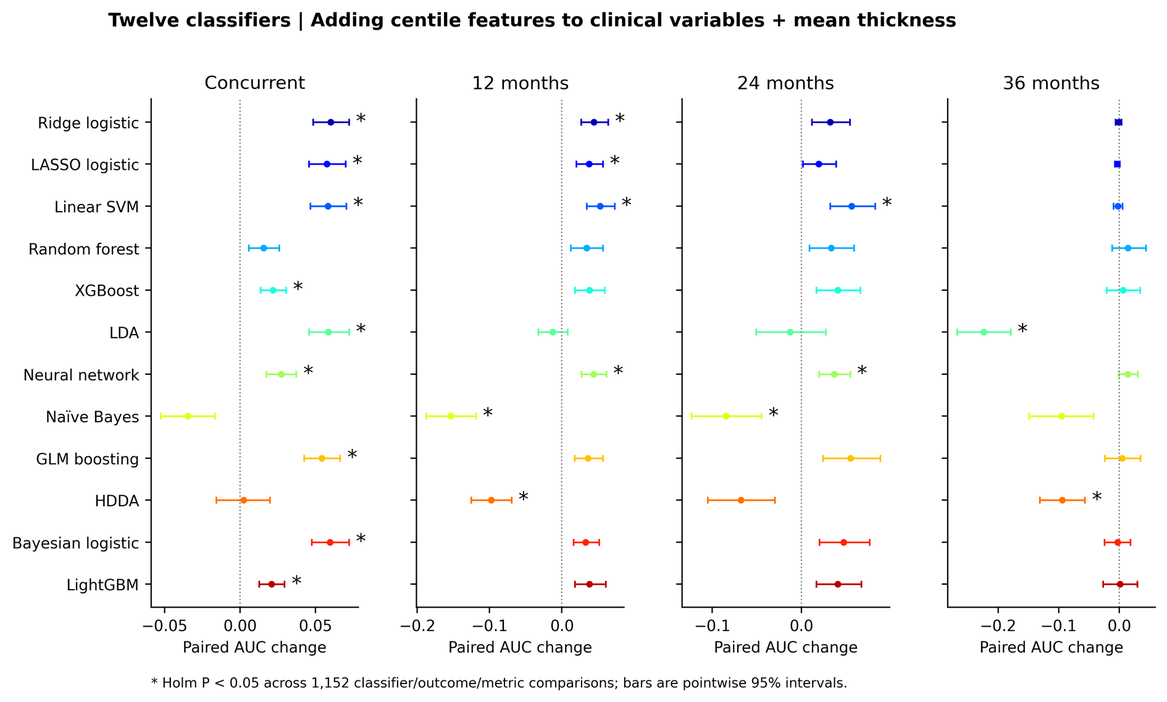

**Figure A9.** Added centile discrimination within twelve classifiers. Points show the change in mean held-out AUC after adding centile profiles to clinical predictors and 14 regional thickness means. Rows identify classifiers; columns show concurrent classification and 12-, 24- and 36-month prognosis. Bars are pointwise 95% participant-bootstrap intervals, with shared weights across models and repeated participant appearances. Intervals condition on the saved predictions and charts and exclude uncertainty from refitting the full analysis. Stars indicate paired participant-jackknife P < .05 after Holm adjustment across 1,152 classifier/outcome/metric tests. Significance is based on these adjusted tests. Exact differences, intervals and P values are provided in the supplementary data files. Main Figure 3 reports absolute discrimination.

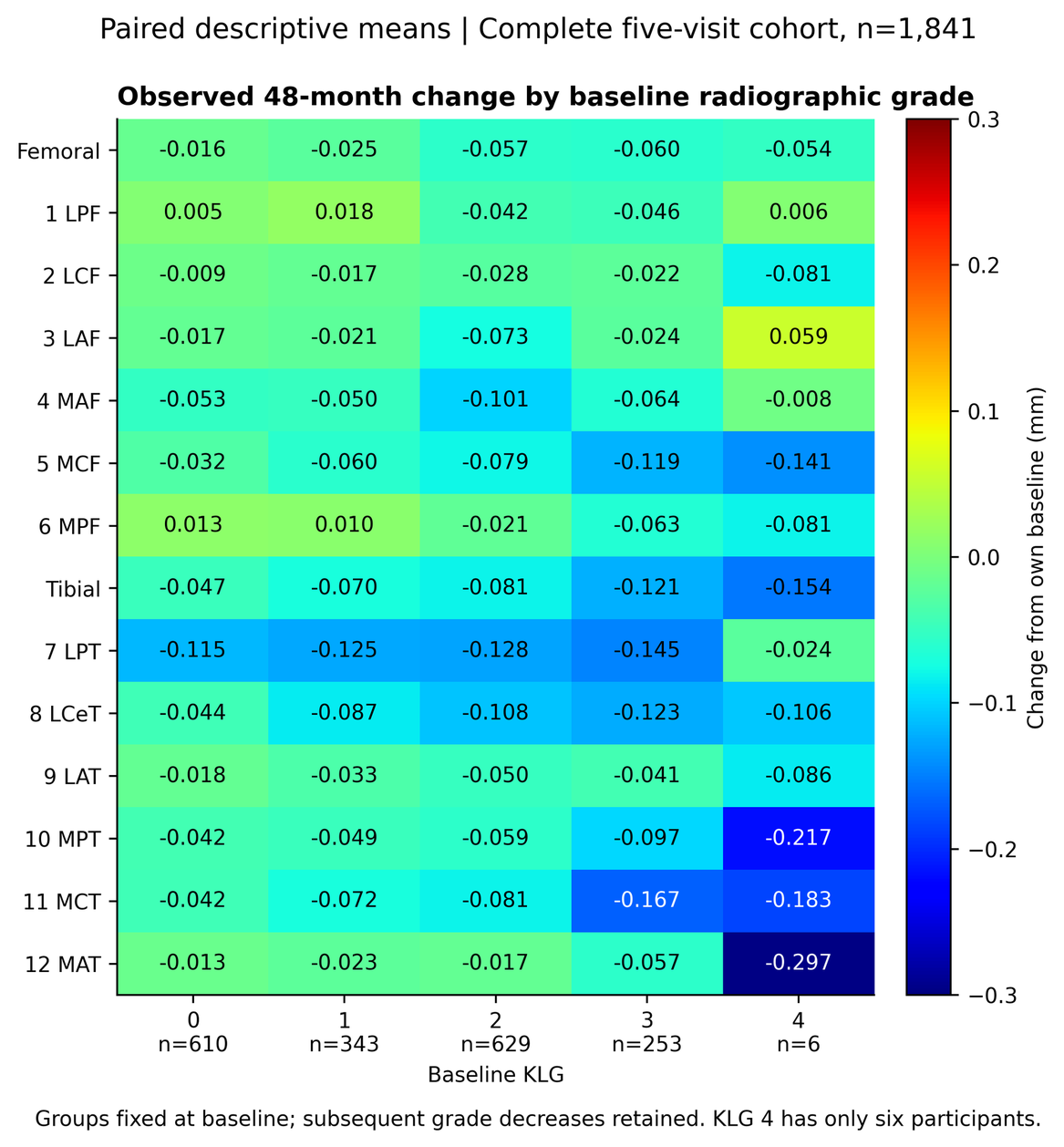

**Figure A10.** Observed 48-month thickness change by baseline grade. Cells show unadjusted mean change from baseline in millimetres for all 14 regions. The complete five-visit cohort contains 1,841 participants, with 610, 343, 629, 253 and six at baseline KLG 0–4. Groups are defined by baseline grade and retained regardless of later increases or decreases in KLG. Estimates for KLG 4 are imprecise because this group contains only six participants. Colours use a common symmetric scale. The supplementary data provide confidence intervals for mean changes and Welch comparisons between baseline-grade groups, with Holm adjustment across 140 contrasts. These descriptive comparisons remain subject to selection and covariate differences.

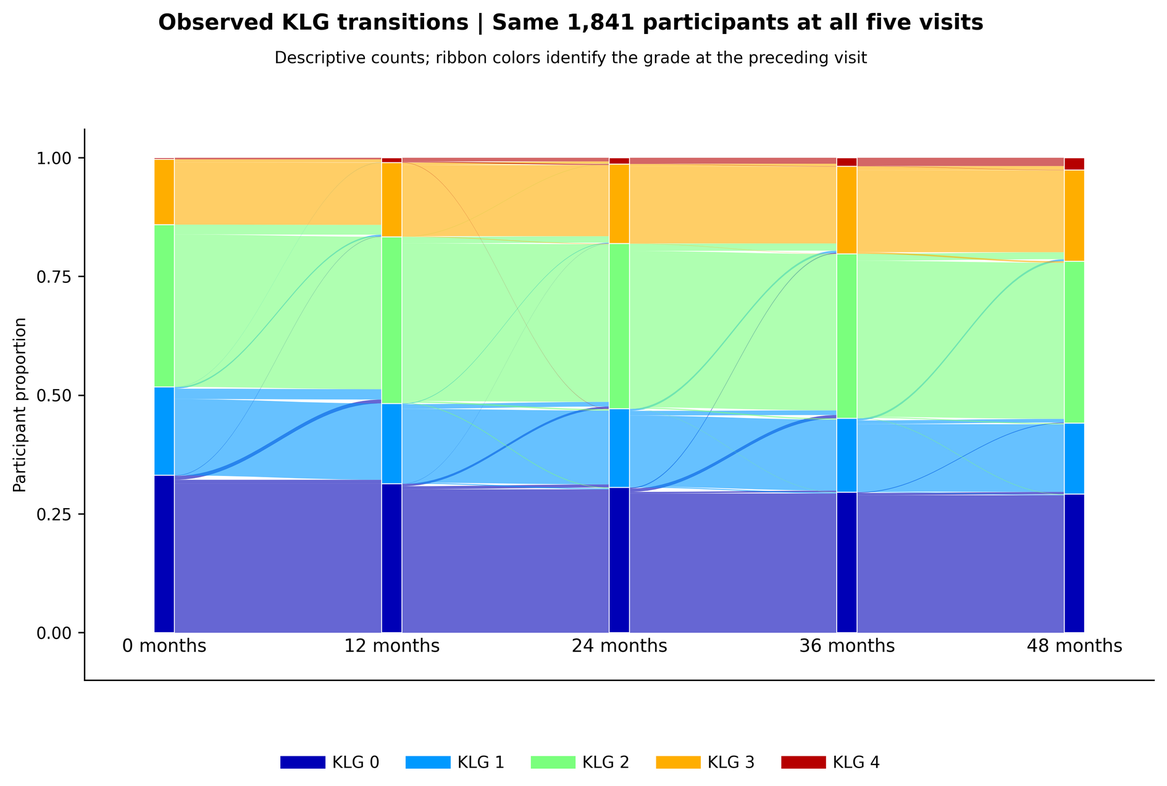

**Figure A11.** Observed radiographic-grade transitions over five visits. The complete five-visit cohort includes 1,841 participants assessed at baseline and 12, 24, 36 and 48 months. Node heights show observed KLG frequencies; ribbon widths show consecutive-visit transitions, coloured by the starting grade. Decreases in grade are retained. Ribbons summarize adjacent visits and do not trace individual trajectories through all five visits. Baseline groups in Figure A10 remain fixed when later grades change.

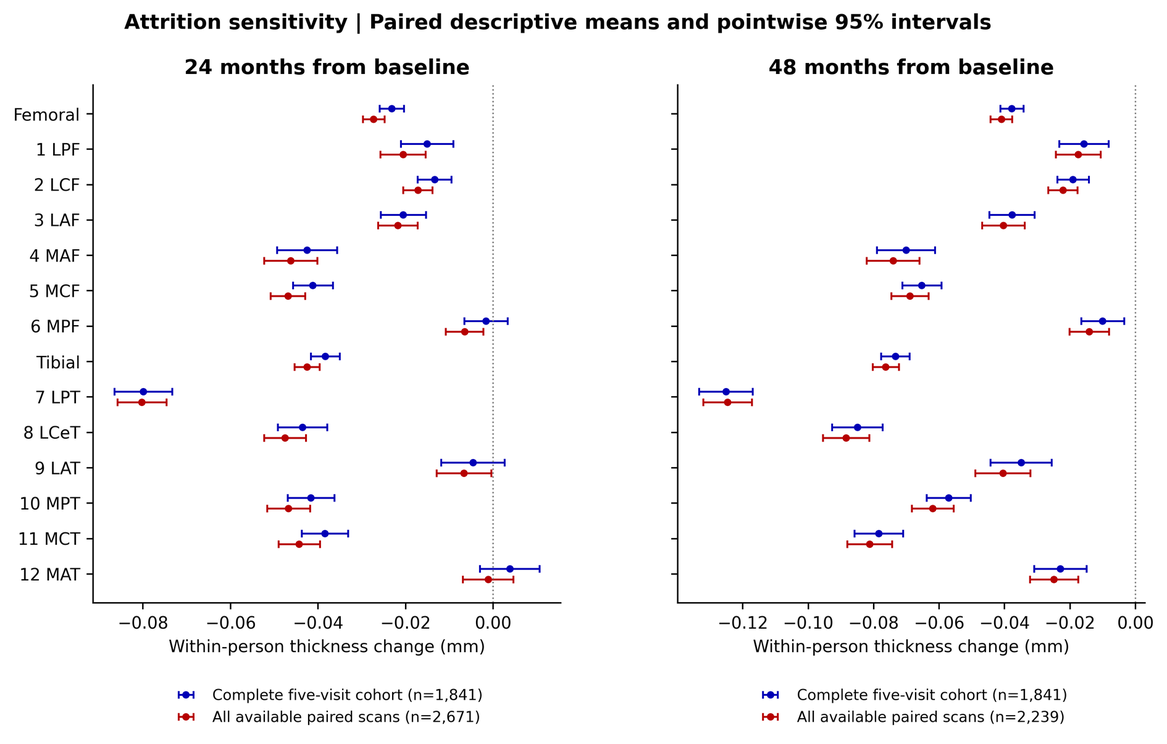

**Figure A12.** Complete-visit and available-pair longitudinal summaries. Points show unadjusted mean thickness change from baseline at 24 and 48 months, in millimetres, with pointwise 95% t intervals. The complete five-visit cohort contains 1,841 participants at both endpoints; the corresponding samples with available baseline–follow-up pairs contain 2,671 and 2,239. Comparisons account for overlapping participants using delete-one-participant jackknife covariance, with Holm adjustment across 28 region/endpoint contrasts. Results are provided in the supplementary data files. This analysis assesses sensitivity to requiring complete follow-up but does not correct selection from missing observations. Both samples include baseline KLG 2–4, whereas the trajectory groups in main Figure 2 are restricted to baseline KLG ≤1.

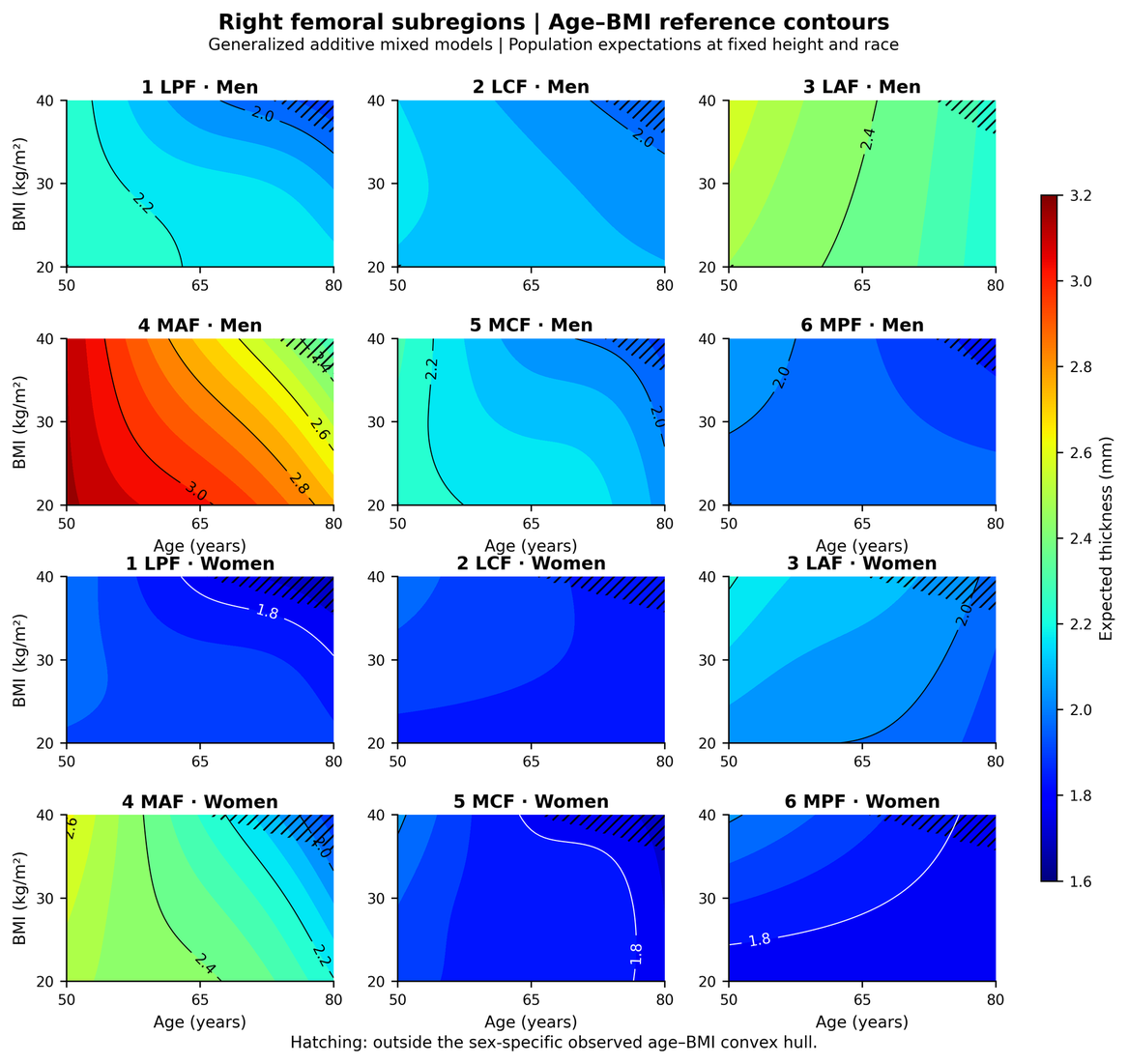

**Figure A13.** Right femoral subregional age–BMI contours. Colours and labelled contours show fitted mean thickness in millimetres for six subregions in men (upper two rows) and women (lower two rows). Predictions cover ages 50–80 years and BMI 20–40 kg/m², with height fixed at 1678.5 mm, race at the reference category and participant intercepts excluded. Hatching marks locations outside the observed sex-specific age–BMI convex hull. Femoral scales are shared across regions, sexes and knees. Region abbreviations follow main Figure 1D. Figure A1 shows the corresponding perspective surfaces and fitted-mean confidence limits.

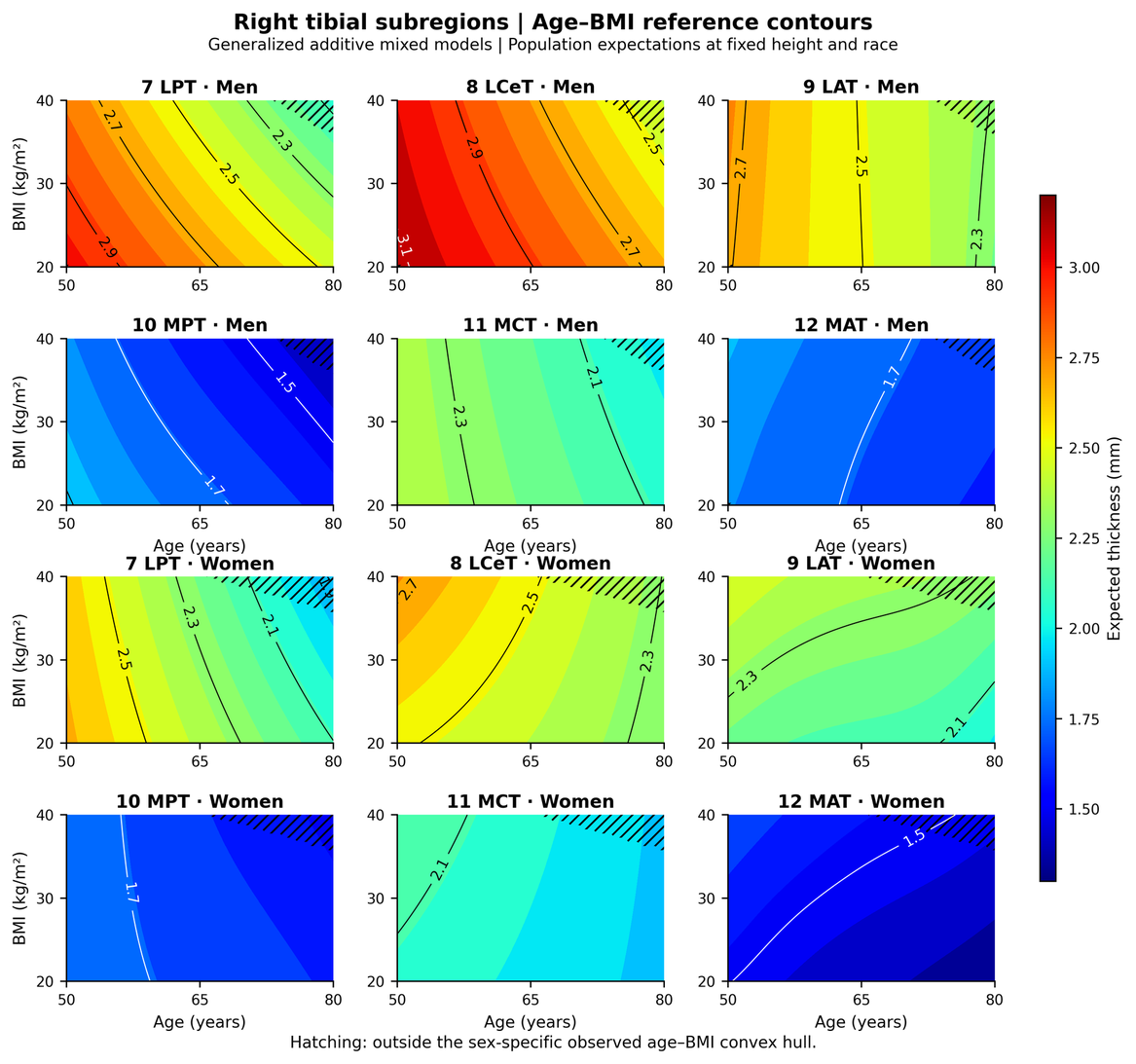

**Figure A14.** Right tibial subregional age–BMI contours. Colours and labelled contours show fitted mean thickness in millimetres for six tibial subregions in men and women. Age, BMI and other covariate settings match Figure A13. Hatching marks predictions outside observed covariate support. Tibial scales are shared across regions, sexes and knees and differ from the femoral scale. Figure A2 shows the corresponding perspective surfaces; main Figure 1C shows whole-compartment right-knee charts.

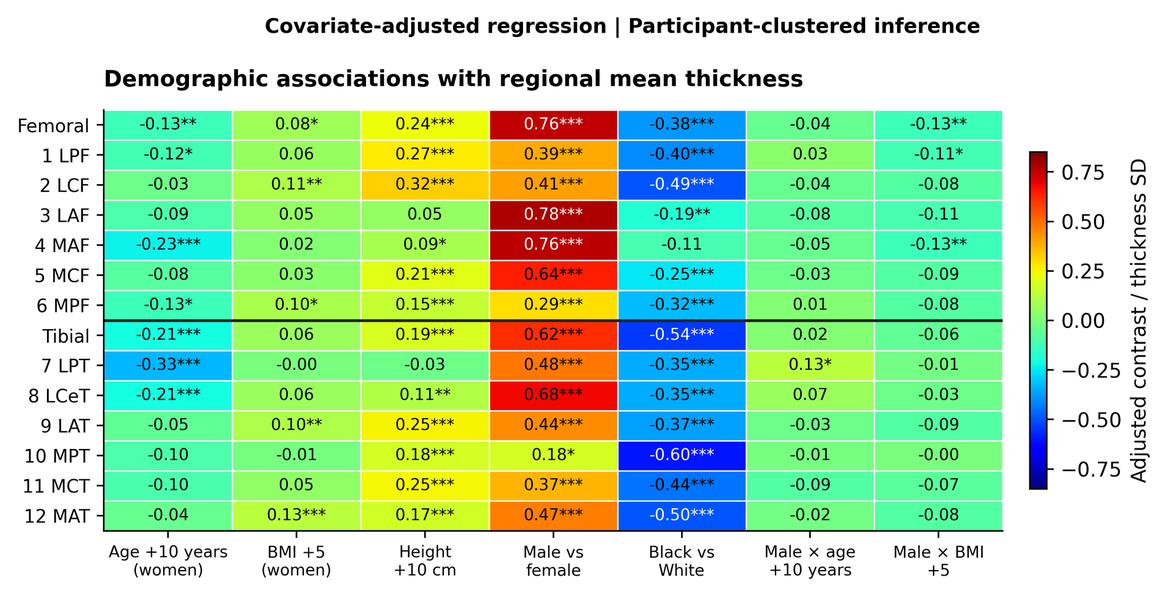

**Figure A15.** Covariate-adjusted regional thickness associations. Contrasts are divided by the regional standard deviation of thickness. Age and BMI contrasts are evaluated for women at age 65 years and BMI 30; sex contrasts use the same age and BMI. Height contrasts span 10 cm. Positive values indicate greater thickness. Stars denote Holm-adjusted $P<.05/.01/.001$. Race refers to recorded categories rather than genetic ancestry.

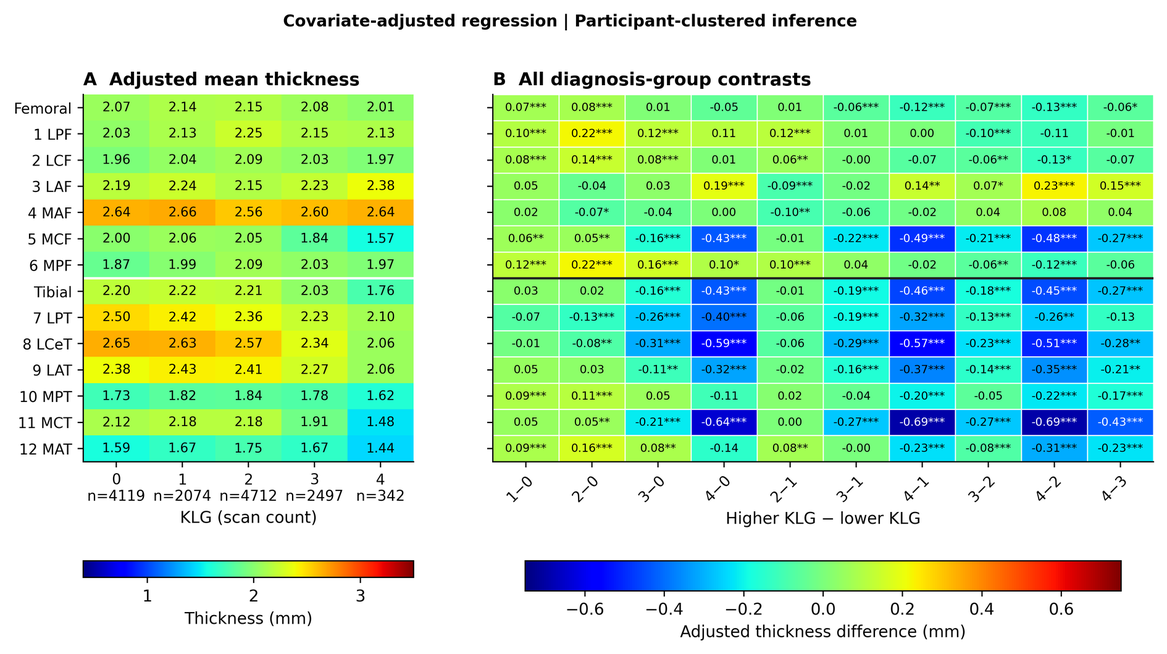

**Figure A16.** Regional thickness across radiographic grades. (A) Adjusted mean thickness in millimetres at each KLG; $n$ denotes the number of scans. (B) Pairwise contrasts between higher and lower grades, with significance symbols as in Figure A15. Advanced grades show lower central medial thickness and greater anterior femoral thickness. These estimates compare grade groups and do not measure within-person change.

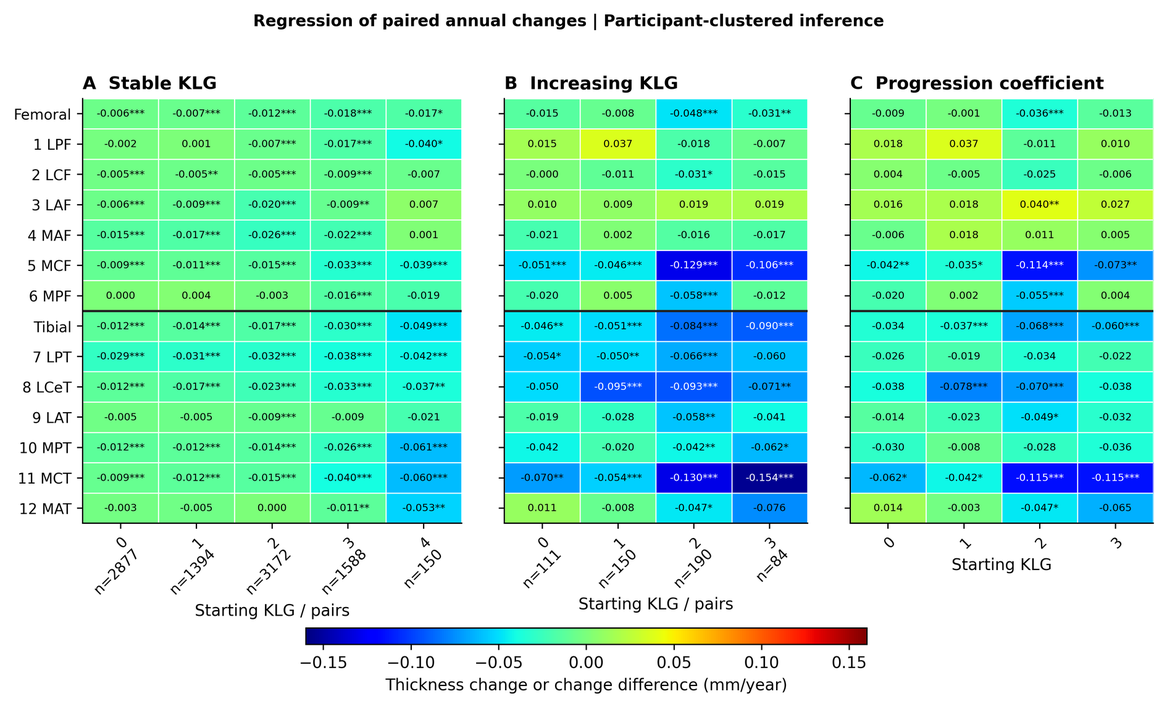

**Figure A17.** Stage-specific annual thinning and thickening. (A) Adjusted annual changes during intervals with stable KLG. (B) Changes during intervals with an increase in KLG. (C) Differences between increasing- and stable-grade intervals at each starting grade. Changes are in mm/year, with blue indicating thinning, green zero change and red thickening. Grade increases may exceed one level; no increase is possible from KLG 4. Counts refer to visit pairs, and covariance estimates account for repeated observations within participants. Stars indicate Holm-adjusted $P<.05$, $.01$ and $.001$, with separate adjustments across 126 means (A/B) and 56 contrasts (C).

### S12. Separate-channel centile comparison

We fitted separate models using cross-sectional centiles (X; superscript P in the equations) and longitudinal centiles (L; superscript H). At 12 and 24 months, longitudinal profiles achieved higher AUC than cross-sectional profiles in 11/12 and 10/12 classifiers after Holm adjustment. The corresponding counts for centile-map deep features were 11/12 and 12/12. Combining both centile types rarely improved on longitudinal centiles alone.

All models included A, comprising 11 clinical predictors and 14 regional thickness means. A+PX and A+PL added 42 cross-sectional or longitudinal profiles, giving 67 predictors each. G2X and G2L added 220 map deep features, giving 245 predictors each. Combined profiles P (= P0) contained 86 features: 42 from each centile type and two compartmental maximum-area measures pooled across types. A+P therefore contained 111 predictors. The combined deep-feature block F2 contained 440 features, and G2 = A + F2 contained 465 predictors. Table S6 defines the complete feature sets. In the result files, OLD_R and OLD_F identify X and L, respectively.

All four separate-centile models used the same participants, control subsets, cross-validation folds and tuning procedures. Each profile or deep-feature block was divided by the square root of its feature count; combining centile types therefore changed both dimension and weighting. The 12 contrasts tested each centile type against A, L against X, combined against separate representations, and deep features against profiles within each type. For every horizon and metric, paired participant-bootstrap tests used 2,000 resamples, with Holm adjustment across 12 classifiers. An additional global Holm adjustment covered all 432 tests per metric in this analysis. These comparisons were added after the original benchmark. Counts of significant classifiers summarize consistency across correlated models and should not be treated as independent replications.

The separate-centile analysis used full-resolution full-reference charts. Cross-fitted analyses in S9 assessed combined centiles and did not test the separate longitudinal representation. All controls were indexed at baseline, whereas 105/196, 75/116 and 23/80 cases had later index scans at 12, 24 and 36 months. Thus, cases and controls differed in the availability of earlier scans despite using the same model folds. This design limits attribution of the observed gain to longitudinal modelling alone. A nonsignificant gain from combining centile types also does not establish equivalence to longitudinal centiles alone. These limitations are discussed in main Sections 3.8 and 5.4.

**Table S23.** Paired performance differences for the separate-centile comparisons. Values are the median difference across 12 classifiers and the numbers with a significant gain or loss after Holm adjustment (+/−). Positive differences favour the first or augmented feature set. Medians summarize the classifiers and do not have a pooled confidence interval. Table S6 defines feature sets and X/L labels. Data S5 provides per-classifier intervals, all eleven metrics and both multiplicity adjustments.

| Metric | Contrast | Horizon | Median Δ | Gains | Losses |
| --- | --- | --- | --- | --- | --- |
| AUC | A+PX - A | 12 | +0.0018 | 0 | 2 |
| AUC | A+PL - A | 12 | +0.0328 | 10 | 2 |
| AUC | A+PL - A+PX | 12 | +0.0332 | 11 | 0 |
| AUC | A+P - A+PX | 12 | +0.0332 | 9 | 2 |
| AUC | A+P - A+PL | 12 | +0.0005 | 0 | 3 |
| AUC | G2X - A | 12 | +0.0065 | 1 | 3 |
| AUC | G2L - A | 12 | +0.0515 | 9 | 1 |
| AUC | G2L - G2X | 12 | +0.0433 | 11 | 0 |
| AUC | G2 - G2X | 12 | +0.0449 | 12 | 0 |
| AUC | G2 - G2L | 12 | +0.0007 | 1 | 1 |
| AUC | G2X - A+PX | 12 | +0.0048 | 0 | 2 |
| AUC | G2L - A+PL | 12 | +0.0156 | 2 | 1 |
| AUC | A+PX - A | 24 | -0.0037 | 0 | 2 |
| AUC | A+PL - A | 24 | +0.0368 | 7 | 0 |
| AUC | A+PL - A+PX | 24 | +0.0406 | 10 | 0 |
| AUC | A+P - A+PX | 24 | +0.0378 | 8 | 0 |
| AUC | A+P - A+PL | 24 | -0.0042 | 0 | 3 |
| AUC | G2X - A | 24 | +0.0110 | 1 | 2 |
| AUC | G2L - A | 24 | +0.0723 | 11 | 1 |
| AUC | G2L - G2X | 24 | +0.0651 | 12 | 0 |
| AUC | G2 - G2X | 24 | +0.0650 | 12 | 0 |
| AUC | G2 - G2L | 24 | -0.0024 | 1 | 1 |
| AUC | G2X - A+PX | 24 | +0.0141 | 0 | 1 |
| AUC | G2L - A+PL | 24 | +0.0384 | 11 | 1 |
| AUC | A+PX - A | 36 | -0.0054 | 0 | 4 |
| AUC | A+PL - A | 36 | -0.0006 | 0 | 2 |
| AUC | A+PL - A+PX | 36 | +0.0078 | 0 | 0 |
| AUC | A+P - A+PX | 36 | +0.0032 | 0 | 1 |
| AUC | A+P - A+PL | 36 | -0.0009 | 0 | 3 |
| AUC | G2X - A | 36 | -0.0082 | 0 | 2 |
| AUC | G2L - A | 36 | +0.0032 | 0 | 2 |
| AUC | G2L - G2X | 36 | +0.0176 | 2 | 0 |
| AUC | G2 - G2X | 36 | +0.0156 | 1 | 0 |
| AUC | G2 - G2L | 36 | -0.0029 | 0 | 1 |
| AUC | G2X - A+PX | 36 | -0.0004 | 0 | 0 |
| AUC | G2L - A+PL | 36 | +0.0072 | 0 | 0 |
| Balanced accuracy | A+PX - A | 12 | +0.0065 | 1 | 1 |
| Balanced accuracy | A+PL - A | 12 | +0.0382 | 7 | 1 |
| Balanced accuracy | A+PL - A+PX | 12 | +0.0348 | 8 | 0 |
| Balanced accuracy | A+P - A+PX | 12 | +0.0380 | 8 | 1 |
| Balanced accuracy | A+P - A+PL | 12 | +0.0005 | 1 | 3 |
| Balanced accuracy | G2X - A | 12 | +0.0098 | 2 | 2 |
| Balanced accuracy | G2L - A | 12 | +0.0844 | 10 | 1 |
| Balanced accuracy | G2L - G2X | 12 | +0.0759 | 11 | 0 |
| Balanced accuracy | G2 - G2X | 12 | +0.0731 | 12 | 0 |
| Balanced accuracy | G2 - G2L | 12 | -0.0036 | 1 | 1 |
| Balanced accuracy | G2X - A+PX | 12 | +0.0010 | 1 | 1 |
| Balanced accuracy | G2L - A+PL | 12 | +0.0444 | 11 | 1 |
| Balanced accuracy | A+PX - A | 24 | -0.0042 | 0 | 4 |
| Balanced accuracy | A+PL - A | 24 | +0.0238 | 5 | 2 |
| Balanced accuracy | A+PL - A+PX | 24 | +0.0384 | 7 | 0 |
| Balanced accuracy | A+P - A+PX | 24 | +0.0362 | 8 | 0 |
| Balanced accuracy | A+P - A+PL | 24 | +0.0011 | 2 | 2 |
| Balanced accuracy | G2X - A | 24 | -0.0049 | 0 | 2 |
| Balanced accuracy | G2L - A | 24 | +0.0937 | 10 | 1 |
| Balanced accuracy | G2L - G2X | 24 | +0.0993 | 12 | 0 |
| Balanced accuracy | G2 - G2X | 24 | +0.0934 | 12 | 0 |
| Balanced accuracy | G2 - G2L | 24 | -0.0052 | 1 | 2 |
| Balanced accuracy | G2X - A+PX | 24 | -0.0002 | 0 | 1 |
| Balanced accuracy | G2L - A+PL | 24 | +0.0653 | 11 | 1 |
| Balanced accuracy | A+PX - A | 36 | -0.0083 | 0 | 4 |
| Balanced accuracy | A+PL - A | 36 | -0.0087 | 0 | 3 |
| Balanced accuracy | A+PL - A+PX | 36 | +0.0016 | 0 | 0 |
| Balanced accuracy | A+P - A+PX | 36 | +0.0023 | 0 | 1 |
| Balanced accuracy | A+P - A+PL | 36 | +0.0018 | 1 | 2 |
| Balanced accuracy | G2X - A | 36 | -0.0028 | 0 | 3 |
| Balanced accuracy | G2L - A | 36 | +0.0042 | 0 | 0 |
| Balanced accuracy | G2L - G2X | 36 | +0.0214 | 2 | 0 |
| Balanced accuracy | G2 - G2X | 36 | +0.0160 | 3 | 0 |
| Balanced accuracy | G2 - G2L | 36 | -0.0008 | 0 | 0 |
| Balanced accuracy | G2X - A+PX | 36 | +0.0012 | 0 | 0 |
| Balanced accuracy | G2L - A+PL | 36 | +0.0150 | 1 | 0 |
| Macro-F1 | A+PX - A | 12 | +0.0050 | 0 | 1 |
| Macro-F1 | A+PL - A | 12 | +0.0441 | 10 | 2 |
| Macro-F1 | A+PL - A+PX | 12 | +0.0381 | 9 | 0 |
| Macro-F1 | A+P - A+PX | 12 | +0.0441 | 9 | 1 |
| Macro-F1 | A+P - A+PL | 12 | -0.0003 | 1 | 3 |
| Macro-F1 | G2X - A | 12 | +0.0139 | 4 | 2 |
| Macro-F1 | G2L - A | 12 | +0.0852 | 9 | 1 |
| Macro-F1 | G2L - G2X | 12 | +0.0797 | 10 | 0 |
| Macro-F1 | G2 - G2X | 12 | +0.0768 | 11 | 0 |
| Macro-F1 | G2 - G2L | 12 | -0.0029 | 1 | 1 |
| Macro-F1 | G2X - A+PX | 12 | +0.0051 | 0 | 2 |
| Macro-F1 | G2L - A+PL | 12 | +0.0310 | 5 | 1 |
| Macro-F1 | A+PX - A | 24 | -0.0062 | 0 | 4 |
| Macro-F1 | A+PL - A | 24 | +0.0277 | 5 | 1 |
| Macro-F1 | A+PL - A+PX | 24 | +0.0393 | 7 | 0 |
| Macro-F1 | A+P - A+PX | 24 | +0.0411 | 8 | 0 |
| Macro-F1 | A+P - A+PL | 24 | +0.0011 | 2 | 2 |
| Macro-F1 | G2X - A | 24 | -0.0032 | 0 | 2 |
| Macro-F1 | G2L - A | 24 | +0.0963 | 9 | 1 |
| Macro-F1 | G2L - G2X | 24 | +0.1018 | 11 | 0 |
| Macro-F1 | G2 - G2X | 24 | +0.0958 | 12 | 0 |
| Macro-F1 | G2 - G2L | 24 | -0.0052 | 1 | 2 |
| Macro-F1 | G2X - A+PX | 24 | +0.0023 | 0 | 1 |
| Macro-F1 | G2L - A+PL | 24 | +0.0664 | 11 | 1 |
| Macro-F1 | A+PX - A | 36 | -0.0090 | 0 | 4 |
| Macro-F1 | A+PL - A | 36 | -0.0073 | 0 | 3 |
| Macro-F1 | A+PL - A+PX | 36 | +0.0023 | 0 | 0 |
| Macro-F1 | A+P - A+PX | 36 | +0.0036 | 0 | 1 |
| Macro-F1 | A+P - A+PL | 36 | +0.0020 | 1 | 2 |
| Macro-F1 | G2X - A | 36 | -0.0035 | 0 | 3 |
| Macro-F1 | G2L - A | 36 | +0.0040 | 0 | 0 |
| Macro-F1 | G2L - G2X | 36 | +0.0208 | 3 | 0 |
| Macro-F1 | G2 - G2X | 36 | +0.0158 | 3 | 0 |
| Macro-F1 | G2 - G2L | 36 | -0.0010 | 0 | 0 |
| Macro-F1 | G2X - A+PX | 36 | +0.0009 | 0 | 0 |
| Macro-F1 | G2L - A+PL | 36 | +0.0151 | 0 | 0 |

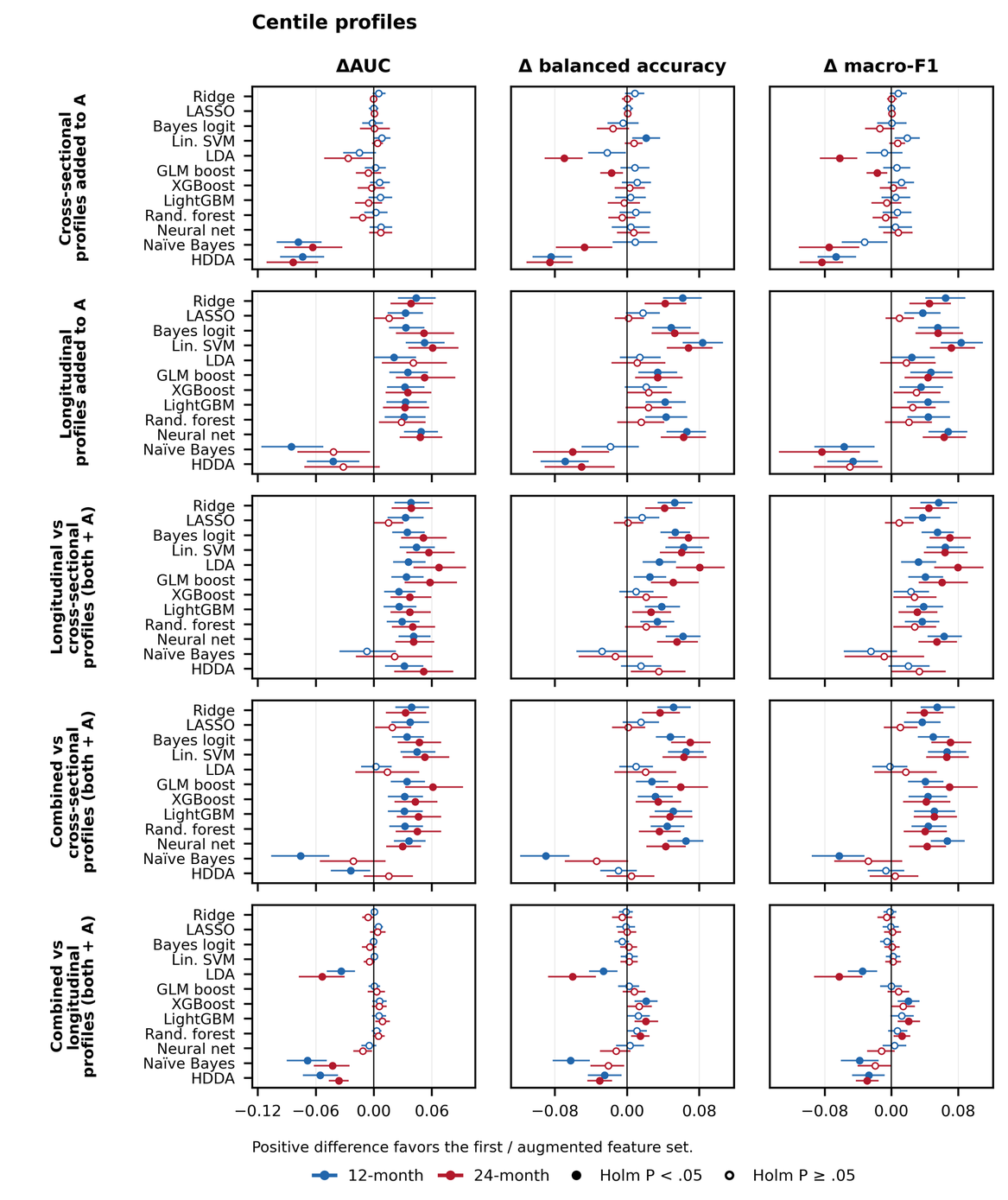

**Figure C1.** Prognostic value of separate and combined centile profiles. Panels compare each centile type with A, L with X, and combined profiles with each separate type. X and L denote cross-sectional and longitudinal centiles. Points show differences in AUC, balanced accuracy and macro-F1 at 12 months (blue) and 24 months (red); bars are paired participant-bootstrap 95% confidence intervals. Filled markers indicate Holm P < 0.05 across 12 classifiers within each contrast, horizon and metric. Positive differences favour the first or augmented set. All models include A, the clinical and regional-thickness baseline; complete definitions are in Table S6. Combined profiles include two pooled maximum-area measures. Main Figure 4 presents the AUC comparisons.

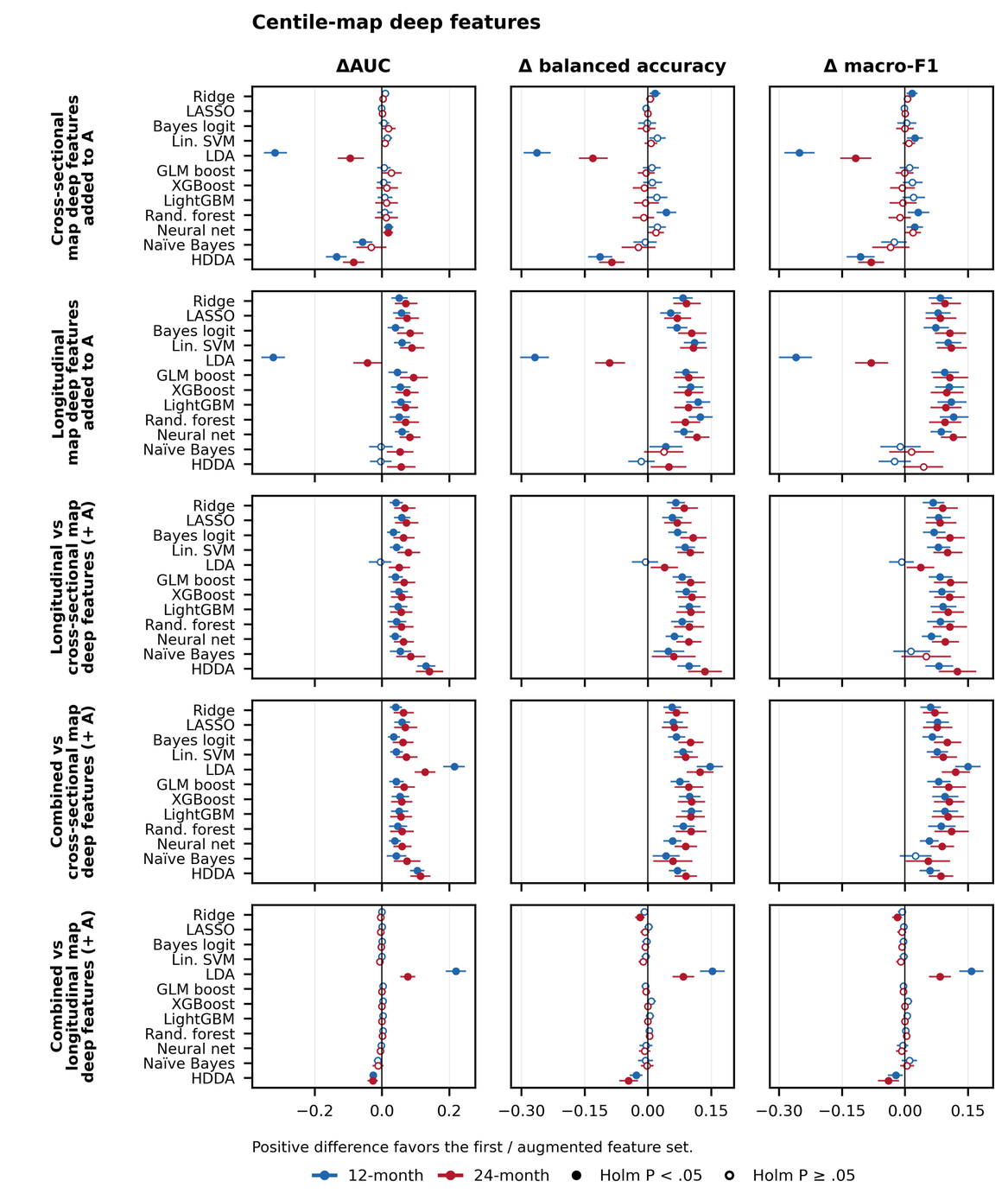

**Figure C2.** Prognostic value of separate and combined centile-map deep features. Panels compare each centile type with A, L with X, and combined maps with each separate type. X and L denote cross-sectional and longitudinal centiles. Points show differences in AUC, balanced accuracy and macro-F1 at 12 months (blue) and 24 months (red); bars are paired participant-bootstrap 95% confidence intervals. Filled markers indicate Holm P < 0.05 across 12 classifiers within each contrast, horizon and metric. Positive differences favour the first or augmented set. All models include A, the clinical and regional-thickness baseline; complete definitions are in Table S6. Combined profiles include two pooled maximum-area measures. Main Figure 4 presents the AUC comparisons.

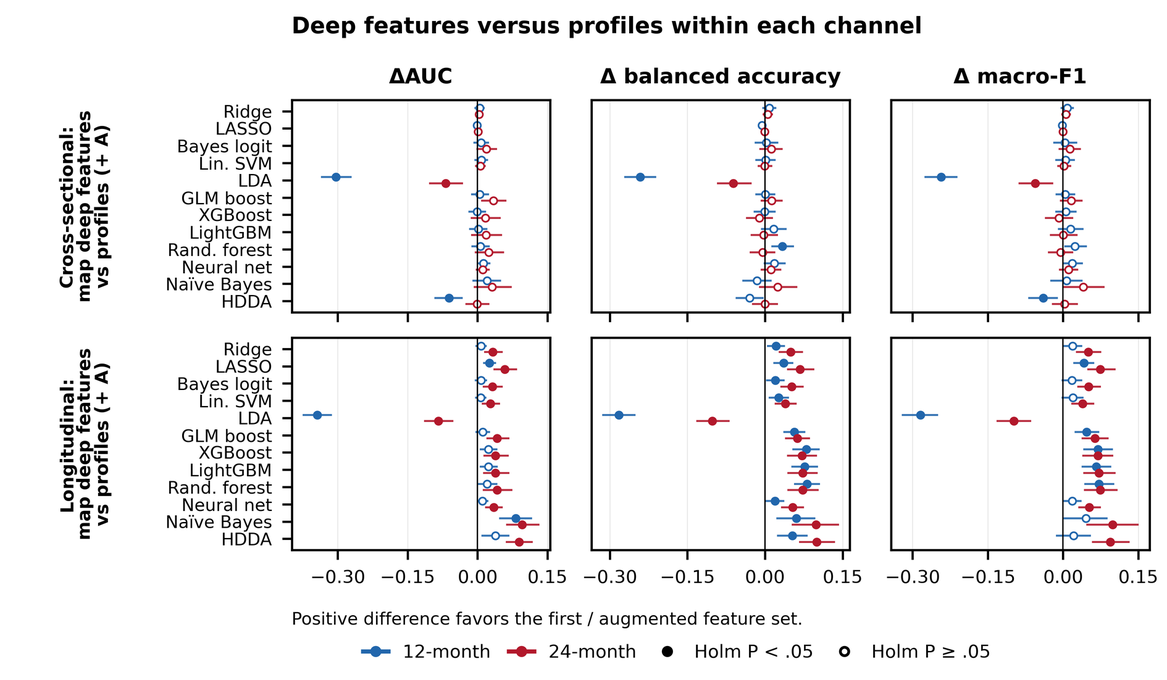

**Figure C3.** Centile-map deep features compared with regional profiles. Comparisons are shown separately for each centile type. X and L denote cross-sectional and longitudinal centiles. Points show differences in AUC, balanced accuracy and macro-F1 at 12 months (blue) and 24 months (red); bars are paired participant-bootstrap 95% confidence intervals. Filled markers indicate Holm P < 0.05 across 12 classifiers within each contrast, horizon and metric. Positive differences favour the first or augmented set. All models include A, the clinical and regional-thickness baseline; complete definitions are in Table S6. Combined profiles include two pooled maximum-area measures. Main Figure 4 presents the AUC comparisons.

### Supplementary data files

Each Excel workbook begins with a README sheet describing its tables, columns and codes. The remaining sheets each contain one data table; CSV copies are also provided.

**Data S1 (DataS1.xlsx).** Performance estimates and 95% confidence intervals for each feature set, classifier, horizon and metric. PCA_radiomics_perf contains the regional PCA, radiomics and centile-profile benchmarks (A, B, C, D, A+P and E), including downsampled cross-fitted and full-reference variants. Deep-feature results are in deep_features_perf (A, A+P, G1, G2, G12, H1, H2 and H12).

**Data S2 (DataS2.xlsx).** Paired differences between feature sets, with 95% confidence intervals and raw, classifier-wise Holm-adjusted and globally Holm-adjusted P values. Results are provided in PCA_radiomics_cmp and deep_features_cmp.

**Data S3 (DataS3.xlsx).** Reference-membership sensitivity analysis (S9). Sheets: ridge_arms_perf, ridge_arms_increment, ridge_arm_differences (ridge, all reference arms), all_metrics_perf and all_metrics_cmp (12 classifiers, all metrics), clf_auc_brier_perf, clf_increments, clf_arm_differences (12 classifiers, AUC and Brier), F2check_perf and F2check_cmp (centile-map deep features from cross-fitted charts).

**Data S4 (DataS4.xlsx).** SHAP explanations (S10). Sheets: shap_importance (mean absolute and signed SHAP of every predictor for each selected classifier and their average), selected_classifiers (the five highest-AUC classifiers per outcome), mc_stability (Monte Carlo stability), fit_validation (per-fit reconstruction and additivity checks).

**Data S5 (DataS5.xlsx).** Separate cross-sectional and longitudinal centile comparisons. The workbook contains performance estimates for seven feature sets, 12 classifiers, three horizons and eleven metrics (2,772 rows), together with twelve paired contrasts (4,752 rows). Additional sheets report median differences, counts of significant gains and losses, and the values plotted in the main and supplementary figures. Raw comparisons retain the X-minus-L direction from the result files; plot-data sheets reverse it to L minus X.
